# Accurate reconstruction of bacterial pan- and core- genomes with PEPPAN

**DOI:** 10.1101/2020.01.03.894154

**Authors:** Zhemin Zhou, Jane Charlesworth, Mark Achtman

**Author notes:** Corresponding author: Z.Z.

## Abstract

Bacterial genomes can contain traces of a complex evolutionary history, including extensive homologous recombination, gene loss, gene duplications and horizontal gene transfer. In order to reconstruct the phylogenetic and population history of a set of multiple bacteria, it is necessary to examine their pan-genome, the composite of all the genes in the set. Here we introduce PEPPAN, a novel pipeline that can reliably construct pan-genomes from thousands of genetically diverse bacterial genomes that represent the diversity of an entire genus. PEPPAN outperforms existing pan-genome methods by providing consistent gene and pseudogene annotations extended by similarity-based gene predictions, and identifying and excluding paralogs by combining tree- and synteny-based approaches. The PEPPAN package additionally includes PEPPAN_parser, which implements additional downstream analyses including the calculation of trees based on accessory gene content or allelic differences between core genes. In order to test the accuracy of PEPPAN, we implemented SimPan, a novel pipeline for simulating the evolution of bacterial pan-genomes. We compared the accuracy and speed of PEPPAN with four state-of-the-art pan-genome pipelines using both empirical and simulated datasets. PEPPAN was more accurate and more specific than any of the other pipelines and was almost as fast as any of them. As a case study, we used PEPPAN to construct a pan-genome of ~40,000 genes from 3052 representative genomes spanning at least 80 species of *Streptococcus*. The resulting gene and allelic trees provide an unprecedented overview of the genomic diversity of the entire *Streptococcus* genus.

## Introduction

Soon after the first bacterial genome was sequenced (Fleischmann *et al*. 1995), it became clear that the genomic contents varied between individual strains within a prokaryotic species. Variable genomic content is caused by the gain or loss of singleton ORFan genes (Daubin and Ochman 2004), genomic islands, selfish DNA (plasmids, bacteriophages, integrative conjugative elements) and/or widespread horizontal gene transfer (Abby *et al*. 2012; Szöllösi *et al*. 2012; Croucher *et al*. 2014). Thus, the designation “pan-genome” was introduced to refer to the entire gene contents of a bacterial species or set of strains (Tettelin *et al*. 2005). Bacterial pan-genomes can be divided into the core genome, which consists of the subset of genes that are present in all genomes and the accessory genome, which consists of genes which are variably present among individual genomes. The core genome often contains phylogenetic signals reflecting the vertical accumulation of mutations, and can be used for assignments of bacterial strains to populations.

An early genomic comparison of eight strains of *Streptococcus agalactiae* indicated that for some bacterial species the total size of the pan-genome may increase indefinitely with the number of genomes sequenced, a concept dubbed an ‘open’ pan-genome (Tettelin *et al*. 2005). The validity of this concept remains questionable because, until recently, few pan-genome analyses have included more than 100 genomes (Vernikos *et al*. 2015), in part because only a limited number of bacterial genomes had been sequenced. Furthermore, initial pan-genome construction algorithms (OrthoMCL (Li *et al*. 2003); Panseq (Laing *et al*. 2010); PGAP (Zhao *et al*. 2012)) were incapable of handling larger numbers of genomes as they rely on an initial all-against-all sequence comparison, which scales computationally with the squared number of gene sequences.

The insufficiency of data no longer exists, as bacterial genome assemblies now number in the 100,000s for some genera (Sanaa *et al*. 2019; Zhou *et al*. 2020). However, such large numbers of genomes exacerbate the scalability problem. Fortunately, at least three recent pipelines (Roary (Page *et al*. 2015); PanX (Ding *et al*. 2018); PIRATE (Bayliss *et al*. 2019)) exist for constructing pan-genomes from large and representative datasets (Ding *et al*. 2018; Alikhan *et al*. 2018).

However, pan-genome construction from large datasets is still hampered by two problems. First, both genome annotations in public repositories and those from automatic annotation pipelines such as PROKKA (Seemann 2014) are incomplete and inconsistent (Wozniak *et al*. 2014; Denton *et al*. 2014; Salzberg 2019). These inconsistencies are propagated into genomic studies and can confound further analyses. Early pan-genome analyses (Tettelin *et al*. 2005; Hogg *et al*. 2007) addressed these problems by running TBLASTN gene-against-genome comparisons, but such inconsistencies between genome annotations are not addressed by the latest generation of pan-genome pipelines, which do not include a reannotation step. Namely, genes which have been fragmented by assembly errors or pseudogenisation may still be relevant to cell function (Goodhead and Darby 2015) and should therefore be included in pan-genomes. The identification of such gene fragments requires comparisons against intact analogs (Lerat and Ochman 2005), but automatic annotation pipelines instead annotate them as multiple intact genes, reducing the size of the estimated core genome and overestimating the overall size of the pan-genome.

The second problem in computing a pan-genome is that of differentiating orthologous genes, which have evolved by vertical descent, from paralogous genes derived from gene duplications or horizontal gene transfer (HGT) events. Paralogous genes can become fixed in populations, but many are gained or lost multiple times. This generates complex patterns of presence/absence along the phylogeny. Therefore, including paralogous genes in a phylogenetic analysis may lead to inaccurate interpretations. State-of-the-art pan-genome pipelines implement either graph- or tree-based algorithms for the identification of paralogous genes (Altenhoff *et al*. 2019). However, tree-based algorithms (used by PanX) which reconcile gene trees with a species tree do not scale well to large datasets. Graph-based algorithms (used by Roary and PIRATE) run faster because they ignore phylogenetic relationships between genomes, but perform poorly on benchmark datasets (Ding *et al*. 2018).

Here we present PEPPAN, a novel pipeline for calculating pan-genomes that specifically deals with the problems described above. We describe the algorithms implemented within PEPPAN, and show that it outperforms other pan-genome methods on both empirical and simulated datasets. As a demonstration of PEPPAN’s capabilities, we present a pan-genome calculated from 3052 representatives of *Streptococcus*, a highly diverse genus.

## Results

### A brief overview of PEPPAN

PEPPAN’s workflow consists of the following five successive groups of operations (Fig. 1 and Supplemental Fig. S1) with additional details in Supplemental Text 1.

1. Identifying representative gene sequences. The inputs for PEPPAN consist of GFF3 formatted genome assemblies (Ensembl Release 98 2019). PEPPAN also accepts inputs of additional nucleotide sequences, which are used to refine gene predictions. To reduce the number of genes used in downstream analyses, PEPPAN iteratively clusters genes using Linclust (Steinegger and Soding 2017), resulting in a single representative gene sequence per 90% nucleotide homology cluster.
2. Identifying all gene candidates in all genomes. Each representative gene is aligned to all genomes using both BLASTn (Altschul *et al*. 1990), which accurately locates short inserts and deletions (INDELs), and DIAMOND (Buchfink *et al*. 2015), which generates amino acid alignments and has greater sensitivity with divergent sequences than BLASTn. Alignments are re-scored and all sequences with homology ≥50% across ≥50% of the representative sequence (Supplemental Text 1.2) are clustered in a NeighbourJoining tree using RapidNJ (Simonsen *et al*. 2011).
3. Identifying clusters of orthologous genes. PEPPAN identifies putative orthologs by calculating a paralogous score for each branch in a gene cluster tree (see Supplemental Text 1.3.2) based on ratio of the pairwise genetic distances of candidate genes within each cluster to the average genetic distances of their host genomes. Using average genetic distances avoids potential errors that can be introduced by using a ‘species’ tree to reconcile individual gene cluster trees (Altenhoff *et al*. 2019). Branches with a paralogous score of >1 are iteratively pruned until none remain. The remaining monophyletic subtrees are treated as putative orthologs. The genomic locations of multiple putative orthologs may overlap in some genomes due to either inconsistent genome annotations or a failure to cluster divergent orthologous sequences in the first stage. These conflicts are resolved by retaining the ortholog with the greatest information score (see Supplemental Text 1.3.3), and eliminating all other gene candidates for that region. The remaining gene candidates from each genome are ordered according to their genomic coordinates, and the final set of orthologous genes is identified based on synteny (see Supplemental Text 1.3.4).
4. Pseudogene prediction and outputs. Each gene candidate in each genome is categorized as either an intact coding sequence (CDS) or a pseudogene, depending on the size of the aligned reading frame relative to its representative gene (Fig. 1C). It is also possible to predict pseudogenes that are disrupted in all genomes by importing their intact analog into PEPPAN as an external representative gene. Finally, the evaluations of all genes, as well as their genomic coordinates and orthologous group are output in GFF3 format, and the extents of the regions that match to their representative genes are saved in FASTA format.
5. Pan-genome analyses. A separate tool, PEPPAN_parser, generates analyses of the estimated pan-genome based on the GFF3 outputs from PEPPAN (details can be found at https://github.com/zheminzhou/PEPPAN/blob/master/docs/source/usage/outputs.rst). Similar to Roary (Page *et al*. 2015) and PIRATE (Bayliss *et al*. 2019), these include rarefaction curves, gene presence matrices and gene presence trees. In addition, PEPPAN_parser can also calculate a core genome tree based on allelic differences of genes which are conserved in most genomes. These core genome trees can scale to 10,000s of genomes, and provide the basis for all core genome MLST schemes in EnteroBase (Zhou *et al*. 2020) (Supplemental Text 3).

**Figure 1.**
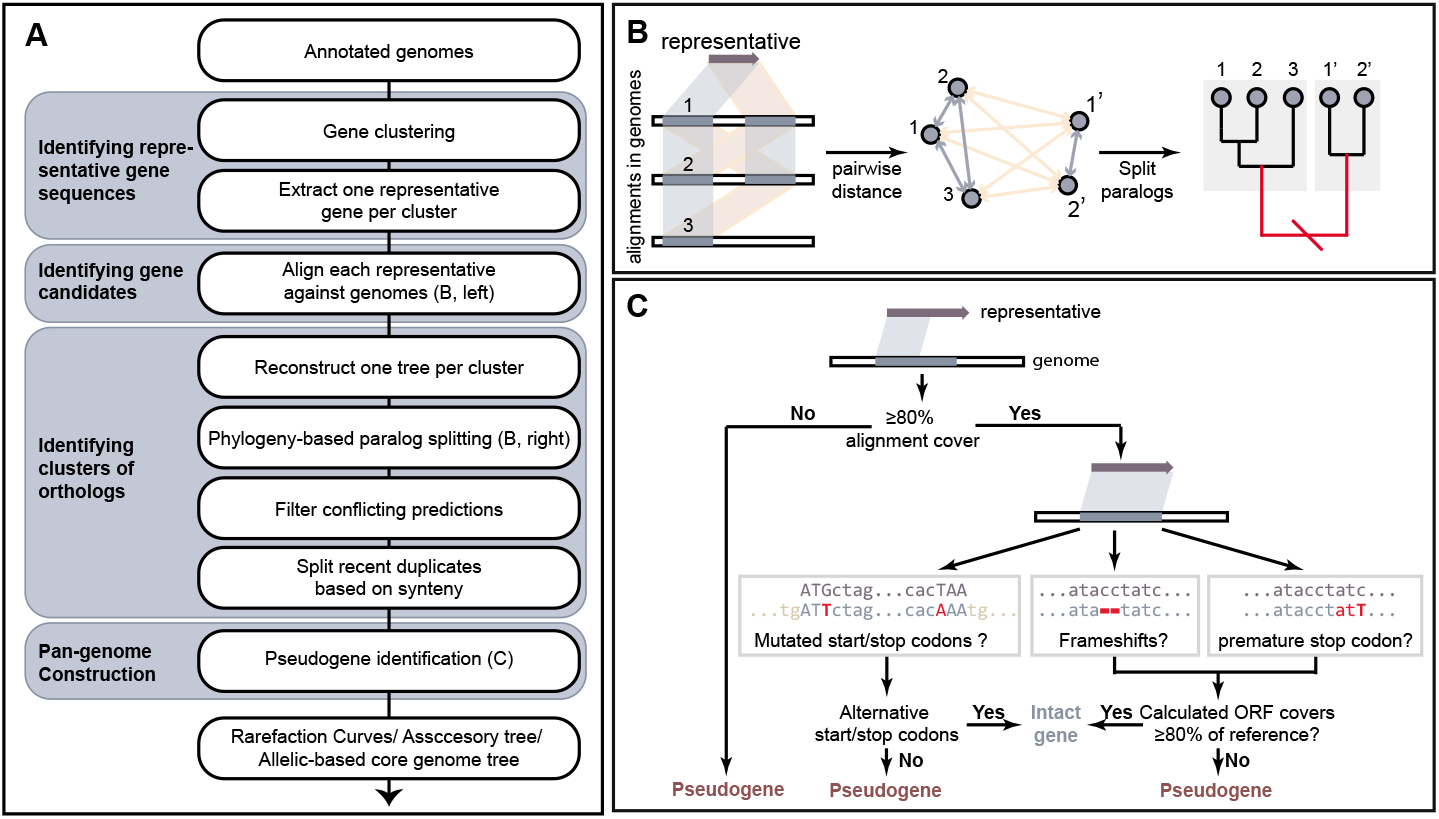
A brief overview of the workflow for PEPPAN. (A) Flow chart indicating the five cascading groups of operations from top to bottom. (B) Cartoon of similarity-based prediction of gene candidates (left) and phylogeny-based paralog splitting (middle and right). The tree was split at the red branch (right) to separate gene candidates into two sub-clusters. The gene pairs in the same sub-cluster had low paralogous scores (blue quadrilaterals and arrows at left) whereas gene pairs between the sub-clusters had high paralogous scores (yellow). (C) Flow chart of the pseudogene identification. The detailed workflow of the algorithm implemented in PEPPAN can be found in Supplemental Fig. S1 and Supplemental Text 1.

### Comparisons of PEPPAN with state-of-the-art pan-genome pipelines

We assessed the absolute performance of PEPPAN, and compared it with other, recently described pipelines for pan-genome construction (Roary (Page *et al*. 2015); PanX (Ding *et al*. 2018); PIRATE (Bayliss *et al*. 2019)) as well as with a classical, smallscale pipeline (OrthoMCL, (Li *et al*. 2003)).

It is important to examine multiple aspects of genomic diversity for these comparisons because the evolutionary history of bacterial pan-genomes can be highly complex. However, we are not aware of any pre-packaged simulation tools that can encompass the entire diversity of bacterial genomic changes, including gene duplications and HGTs (leading to paralogs), homologous recombination and large-scale gene insertions and deletions. We therefore performed our first benchmarks by comparing a pan-genome calculated from 15 manually-curated *Salmonella enterica* genome annotations (Nuccio and Bäumler 2014) with pan-genomes based on automated annotations of the same genome assemblies. Subsequently, we designed a new simulation tool, SimPan, which uses SimBac (Brown *et al*. 2016) to simulate the dynamics of pan-genome evolution *via* recombination, HGT, gene gain and loss as well as the creation of paralogs (Supplemental Text 2).

### Benchmarking pan-genome pipelines on 15 curated genomes

Nuccio and Bäumler re-annotated 15 complete genomes of *S. enterica* (Nuccio and Bäumler 2014). They removed existing annotations for unreliable short genes, performed new BLASTN and TBLASTN alignments to identify previously not annotated genes, corrected the start positions of falsely annotated genes and predicted the existence of pseudogenes based on alignments with orthologous intact CDSs. The result of these efforts is a unique set of consistently annotated genomes from a single species, which we equated with the ‘ground truth’ with which to compare the results from the pan-genome pipelines.

First, we compared the manual re-annotation with three sets of gene annotations for each of the 15 *S*. *enterica* genomes: (1) the original annotation that had been submitted to GenBank (“Submitter”), (2) an automated re-annotation from RefSeq (Haft *et al*. 2018) that was generated with PGAP (Tatusova *et al*. 2016), (3) a novel annotation using PROKKA (Seemann 2014), another popular bacterial annotation pipeline. Genes that had been eliminated by Nuccio and Bäumler as being “unreliable” were removed from all three annotations for consistency. We then examined the degree of concordance between the pan-genome published by Nuccio and Bäumler with the pan-genomes calculated by each of the pipelines. Concordance was estimated by calculating the adjusted Rand index (ARI) (Rand 1971), which is a measure of similarity between clustering results. For Roary or PIRATE we only report results from the run with the greatest ARI among three parallel runs with varying minimum identity (50, 80 or 95%), because the optimal value of this parameter differs for various levels of diversity ((Ding *et al*. 2018) and our own observations).

All pipelines successfully calculated a pan-genome from each of the four annotations, except that “Submitter” annotations never ran to completion with PanX. The PEPPAN pan-genomes consistently yielded ARIs of ~0.98 relative to the manual pan-genome (Fig. 2A, histograms). This is not surprising because PEPPAN re-calculates gene annotations in a fashion which resembles that of the manual curation. All the other pipelines yielded lower ARI values which varied between the annotation methods. The PROKKA annotations yielded ARIs of 0.97 with Roary, PanX, and OrthoMCL and 0.96 with PIRATE. The ARIs were 0.95-0.96 for the PGAP annotations from RefSeq, and 0.93-0.94 for the “Submitter” annotations. We also performed hierarchical clustering using the Neighbor-Joining algorithm on pair-wise comparisons of the ARI scores across all 14 pan-genomes (Fig. 2B). The three pan-genomes predicted by PEPPAN formed a tight cluster with high pair-wise ARI (0.99), which clustered tightly with the curated pan-genome (ARI=0.98). In contrast, pan-genomes generated by the other pipelines clustered according to annotation source rather than pipeline methodology. For each of the three annotation sources, the pan-genome predicted by Roary was the most distinct whereas pan-genomes predicted by OrthoMCL, PanX, and PIRATE clustered more tightly. These results may reflect the fact that Roary differs from the other pipelines by performing an additional splitting of paralogs on the basis of synteny.

**Figure 2.**
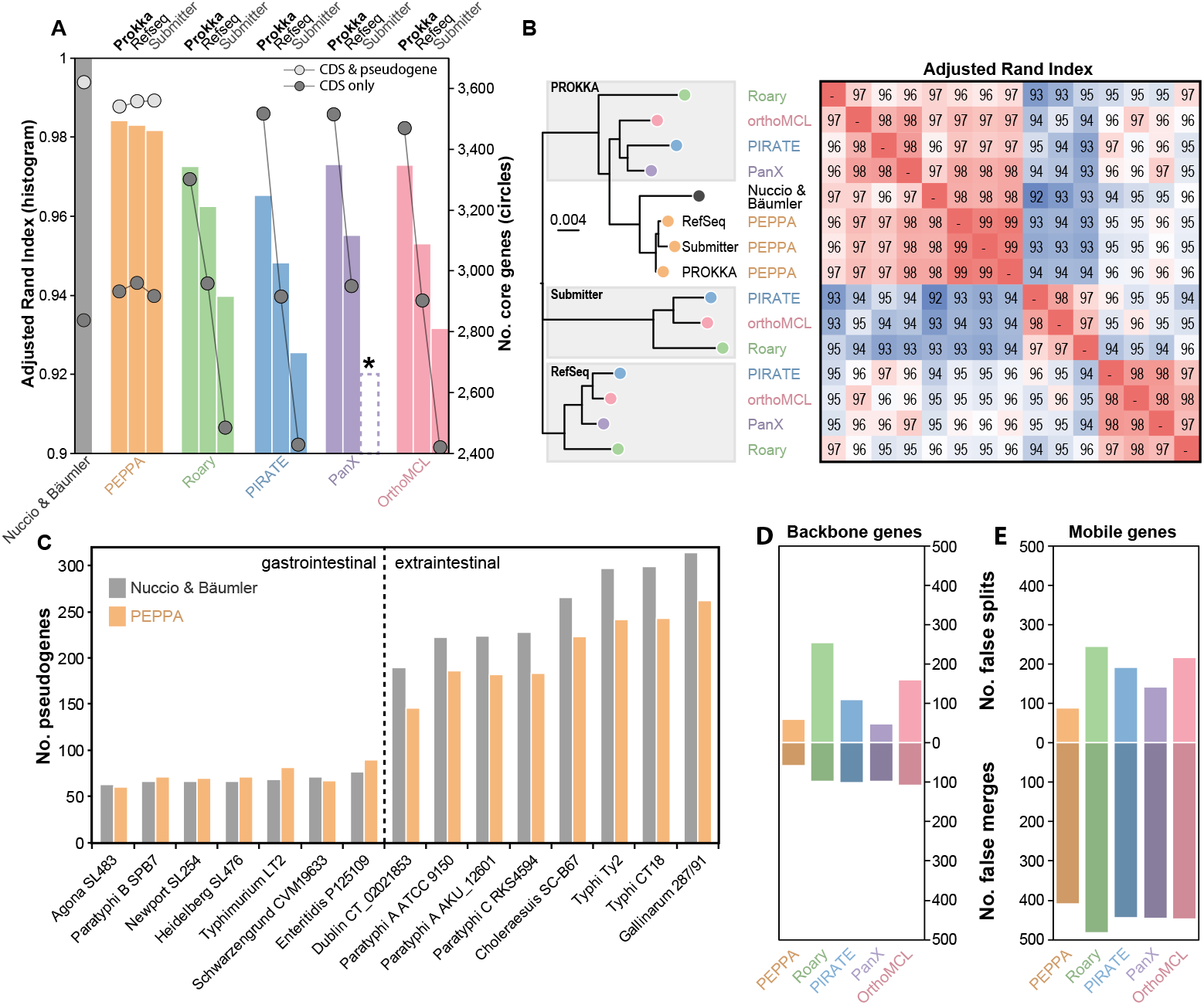
**Comparison of pan-genome predictions for 15 *Salmonella* genomes with a manually curated pan-genome (Nuccio and Bäumler 2014).** (A) The adjusted Rand index *versus* the manual curation (ARI; histogram) and the sizes of core genomes (circles) in each of the pan-genomes after annotation by PROKKA (Seemann 2014), after reannotation in RefSeq with PGAP (Tatusova *et al*. 2016), and as originally submitted to NCBI (Submitter). *indicates that PanX failed to run on the “Submitter” annotations. (B) A Neighbour-joining tree (left) of the pairwise ARI scores (heatmap at the right) between the predicted pan-genomes and the curated pan-genome. The annotation source is indicated within grey shadows at the left except for PEPPAN where it is listed at the tips. Colors are as in part A. (C) Histogram of the numbers of pseudogenes (Y-axis) in each of the genomes (X-axis) in the curated pan-genome (grey) and pan-genome predicted by PEPPAN (orange). A dashed line separates the two *Salmonella* pathovar groups described by Nuccio and Bäumler. (D, E) Histograms of the average numbers of false splits (top) and merges (bottom) of ortholog groups by the individual pipelines (X-axis) in backbone (D) or mobile (E) genes.

#### Pseudogenes prediction

The core genome defined by Nuccio and Bäumler contained 2,838 CDSs that were intact in all 15 genomes and 783 others that were disrupted in at least one genome. PEPPAN predicted marginally more intact CDSs, and slightly fewer pseudogenes, from all three annotations than were present in the manual annotations (Fig. 2A, circles). The number of pseudogenes for each genome was also very similar between the manual curations and PEPPAN’s automated predictions. We note that PEPPAN consistently predicts fewer pseudogenes for extraintestinal strains than those for those linked to gastrointestinal disease (Fig. 2C), This is an interesting observation, as accumulation of pseudogenes has been linked to host specialization in *Salmonella* (Parkhill *et al*. 2001; Holt *et al*. 2008; Nuccio and Bäumler 2014; Zhou *et al*. 2014; Zhou *et al*. 2018b).

Roary, OrthoMCL and PanX do not predict any disrupted genes. PIRATE reports ‘gene diffusion’, a measure of the frequency with which CDSs that are intact in some genomes are split into two or more fragments in others. However, it did not detect any gene diffusion in the RefSeq and GenBank annotations, and only one instance with the PROKKA annotations. PIRATE also failed to predict fragmented genes. Similar to the ARI comparisons described above, the total numbers of predicted core CDSs varied according to annotation source for all pipelines other than PEPPAN. The four pipelines reported 3,301-3,515 core CDSs from PROKKA annotations (Fig. 2A right). These numbers are similar to the total number of intact core CDSs plus pseudogenes within the curated pan-genome, indicating that PROKKA predicted many pseudogenes as intact CDSs. Roary, PIRATE and OrthoMCL only detected 2,418-2,484 core genes in the originally submitted genomes, suggesting inconsistencies between individual genome annotations. In contrast, all four pipelines predicted 2,901-2,957 core CDSs from the RefSeq annotations, and these numbers were similar to the numbers of intact core CDSs in the curated pan-genome (2,838), or as predicted by PEPPAN (2,918-2,961).

#### Inaccurate prediction of orthologs

Inconsistent ortholog calls relative to the manually curated pan-genome (Nuccio and Bäumler 2014) also contributed to variation in the numbers of core CDS predicted by the different pipelines. We designated as ‘false splits’ those cases where a single ortholog cluster in the curated pan-genome was split into multiple ortholog clusters by a pipeline. Similarly, ‘false merges’ occurred when multiple orthologous clusters in the curated pan-genome were assigned to a single orthologous cluster. We identified 4,695 ‘backbone genes’ in the curated pan-genome which were present in the most recent common ancestor (MRCA) and 3,364 ‘mobile’ genes, which were associated in one or more genomes with mobile genetic elements, and which were absent from the MRCA. For backbone genes, PEPPAN made the fewest false splits and false merges of all five pipelines, followed by PanX (Fig. 2D). False merges were made four times as often by all pipelines for mobile genes than for backbone genes, and false splits were up to two times as frequent (Fig. 2E *vs*. 2D). Roary generated the highest number of false calls, while PEPPAN generated the lowest.

### Simulating pan-genome datasets

The analyses above indicate that the backbone and mobile genes might differ in their rates of gain and loss during evolution. In order to test the abilities of pan-genome pipelines to handle varying rates of gene gain and loss, we created SimPan (https://github.com/zheminzhou/SimPan) to simulate the evolution of real bacterial pangenomes (Supplemental Fig. S2 and Table S1; see Supplemental Text 2 for details). In brief, SimPan uses SimBac (Brown *et al*. 2016) to generate a clonal genomic phylogeny. This clonal phylogeny is subjected to random homologous recombination, resulting in different “local trees” that reflect the individual ancestries of backbone and mobile genes. Random INDEL events leading to loss or gain of blocks of genes are simulated along the branches of these local trees until the average number of genes per genome and in the core genome attain user-specified parameters ‘--aveSize’ and ‘--nCore’ (Supplemental Table S1). This results in a presence/absence matrix of all backbone and mobile genes. Finally, sequences of both genes and intergenic regions are subjected to short INDELs, converted into genes with INDELible (Fletcher and Yang 2009) and concatenated into whole genomes.

We simulated five genomic datasets each containing 15 genomes, using parameters derived from the curated *S. enterica* pan-genome, with each genome containing a mean of 3,621 core genes and 879 accessory genes (simulations a-e). We arbitrarily assigned 5% of the backbone genes and 40% of the mobile genes to paralogous clusters, and varied their mean percentage sequence identities between each set of simulations (Fig 3, inset). Simulation c represents the simplest pan-genome construction scenario, with high sequence identity (98%) between genes in an ortholog cluster and low sequence identity (60%) between genes in a paralog cluster. Simulations a and b have decreasing levels of identity between orthologs to simulate more diverse species while simulations d and e have increasing levels of identity between paralogs in order to simulate recent gene duplications.

**Figure 3.**
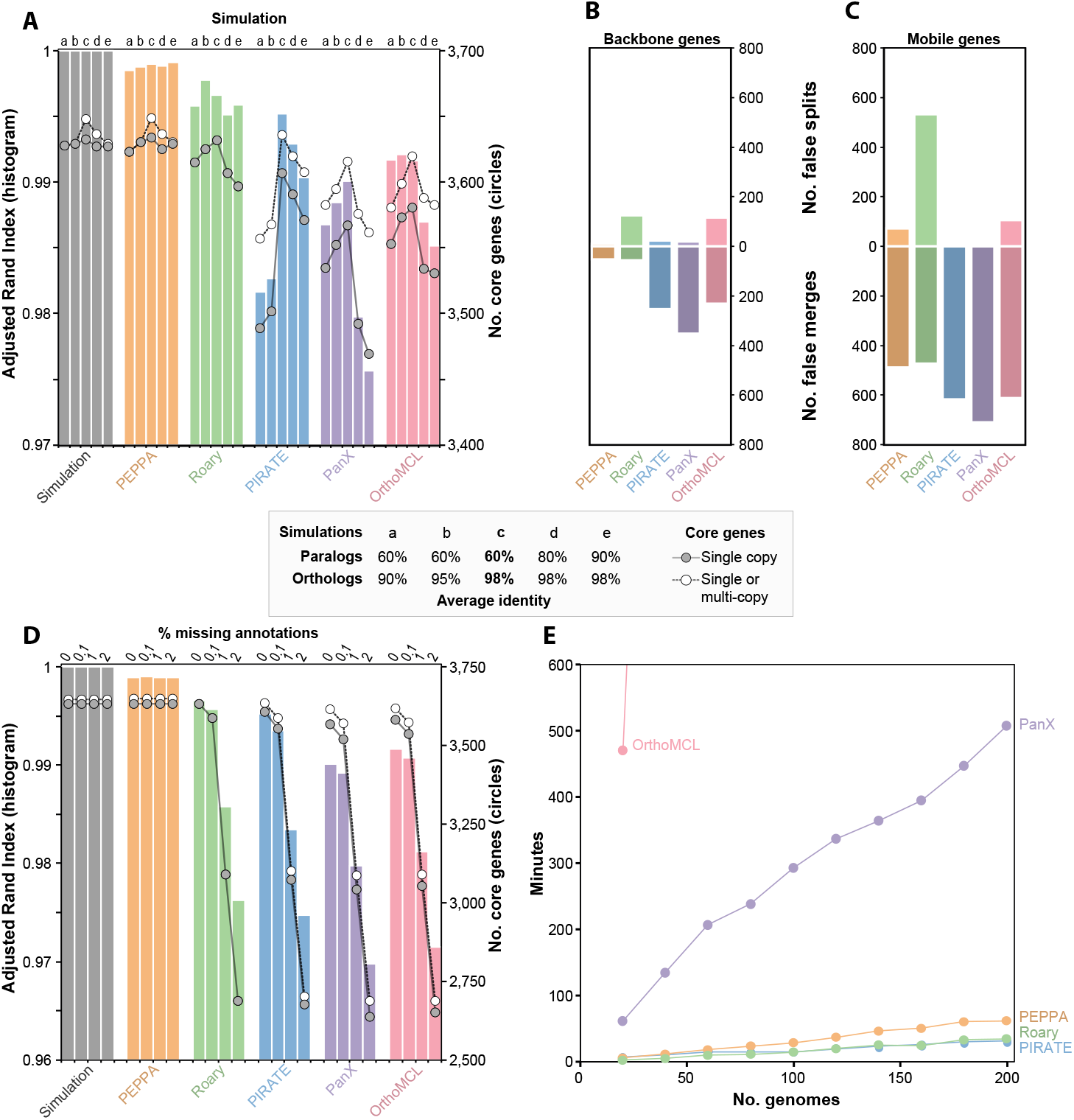
Comparison of the pan-genome pipelines with simulated data generated by SimPan. (A) The adjusted Rand index (ARI; histogram) and the sizes of core genomes (circles) in the pan-genomes produced by SimPan simulations a, b, c, d, e (inset). Left: pan-genome produced by the simulations. Other histograms, pan-genomes calculated by five pipelines. (B) Numbers of failed splits (top) and false merges (bottom) of ortholog groups by five pipelines with backbone genes. (C) Numbers of failed splits (top) and false merges (bottom) of ortholog groups by five pipelines with mobile genes. (D) The adjusted Rand index (ARI; histogram) and the sizes of core genomes (circles) in the pan-genomes produced by SimPan simulation c after random deletions of 0%, 0.1%, 1% and 2% of the gene annotations. Other details as in part A. (E) Runtime for each pipeline (Y-axis) versus number of genomes in simulated datasets (X-axis). Runs which exceed 600 minutes are not shown.

### Pipeline performance on simulated genomes

Pan-genomes calculated from each simulated dataset by PEPPAN, Roary, PIRATE, PanX, and OrthoMCL were compared to the original pan-genomes produced by SimPan (Fig. 3A). Once again, PEPPAN pan-genomes were highly concordant with the known truth (ARI ≥0.998 for all comparisons). Roary performed comparably to PEPPAN on all simulated datasets (ARI ≥ 0.995). PIRATE performed almost as well on simulations c to e, but yielded ARI scores below 0.99 when run on simulations of more diverse genomes (simulations a and b). In contrast, PanX and OrthoMCL yielded ARI scores ≥0.99 when run on simulations a and b, but were less concordant (ARI < 0.99) when run on simulations containing more recent gene duplications (simulations d and e).

PEPPAN correctly predicted all core genes in simulations b, c, and e, and only missed 2-3 core genes in the two remaining datasets (Fig. 3A, circles). Roary correctly predicted all single-copy core genes for simulation c, but failed to identify any multi-copy core genes for any dataset, likely due to its aggressive synteny-based paralog identification step. PIRATE, PanX, and OrthoMCL significantly underestimated the number of core genes when only single-copy core genes were counted, suggesting a high frequency of false splitting of paralog clusters. Indeed, the frequency of false merges was particularly high for backbone genes with these three pipelines, and the frequency of false splits was high with Roary and OrthoMCL (Fig. 3B). All pipelines made multiple false merges of mobile genes, possibly because of their predominance among paralog clusters, and Roary also made large numbers of false splits (Fig. 3C). Overall, PEPPAN made the fewest false calls for both backbone and mobile genes, which explains its higher ARI scores.

#### The effects of missing gene annotations on the pan-genome

As shown above, inconsistent or inaccurate gene annotations are problematic for calculating reliable pangenomes. We simulated this effect by randomly deleting 0.1%, 1% or 2% of the gene annotations from simulation c (Fig 3D). Because PEPPAN re-assigns individual genes to ortholog clusters, it was unaffected by these manipulations. However, the missing annotations yielded drastically reduced ARI scores (Fig. 3D. histograms) and core genome sizes (Fig. 3D, circles) for the other pipelines, and ARI scores became progressively worse with the proportion of missing annotations.

#### Computation time

We generated 10 additional simulations of 20 to 200 genomes with the same parameters as simulation c, and measured the running wall times to calculate a pan-genome for all five pipelines using 4 processors on a server with 1 TB of memory and 40 CPU cores (Fig. 3E). OrthoMCL was the slowest and needed >24 h for ≥60 genomes. PanX was at least eightfold slower than the other three pipelines, and needed 500 min for 200 genomes, despite using a divide-and-conquer algorithm on datasets with >50 genomes. Both Roary and PIRATE scaled very well, and each completed the calculations on 200 genomes within 30 minutes. PEPPAN is about twice as slow as either Roary or PIRATE, and needed 63 minutes for 200 genomes. The good scalability of these pipelines is likely related to the pre-clustering step, which reduces the number of genes used in downstream comparisons. However, this pre-clustering step becomes less efficient with increasing genetic diversity: in an independent simulation of 200 genomes with only 90% sequence identity, the runtime for all three pipelines increased by at least twofold relative to simulation c (PEPPAN: 144 min; Roary: 132; PIRATE: 60).

### A pan-genome for the genus *Streptococcus*

PEPPAN can construct a pan-genome from thousands of genomes with high genetic diversity, and earlier versions of this pipeline were used to generate cgMLST schemes for the genera represented in EnteroBase (Alikhan *et al*. 2018; Frentrup *et al*. 2019; Zhou *et al*. 2020) as well as for ancient DNA analyses (Zhou *et al*. 2018b; Achtman and Zhou 2019). To demonstrate PEPPAN’s capability on genetically diverse datasets, we chose the genus *Streptococcus*, which includes highly significant zoonotic and human pathogens (Gao *et al*. 2014).

We generated a dataset of 3052 high-quality genomes (Supplemental Table S2A) representing the entire taxonomic diversity of *Streptococcus* (see Methods). PEPPAN took 5 days to construct a pan-genome from this dataset. The resulting pan-genome contained 39,042 genes, twice as many as a previous pan-genome based on 138 *Streptococcus* genomes (Gao *et al*. 2014). In agreement with the earlier conclusions by Gao *et al*., the rarefaction curve showed no sign of plateauing, and the pan-genome continued to expand with each new genome added (Fig 4A). Gao *et al*. estimated that the pan-genome would expand by 62 genes for each new genome, whereas we estimate a lower rate of 39 genes per new genome for a randomly sampled set of 138 genomes. However, the growth rate dropped dramatically with the increased number of genomes, and we estimate that the future expansion rate of the pan-genome is only 4.4 new genes for every newly added genome.

**Figure 4.**
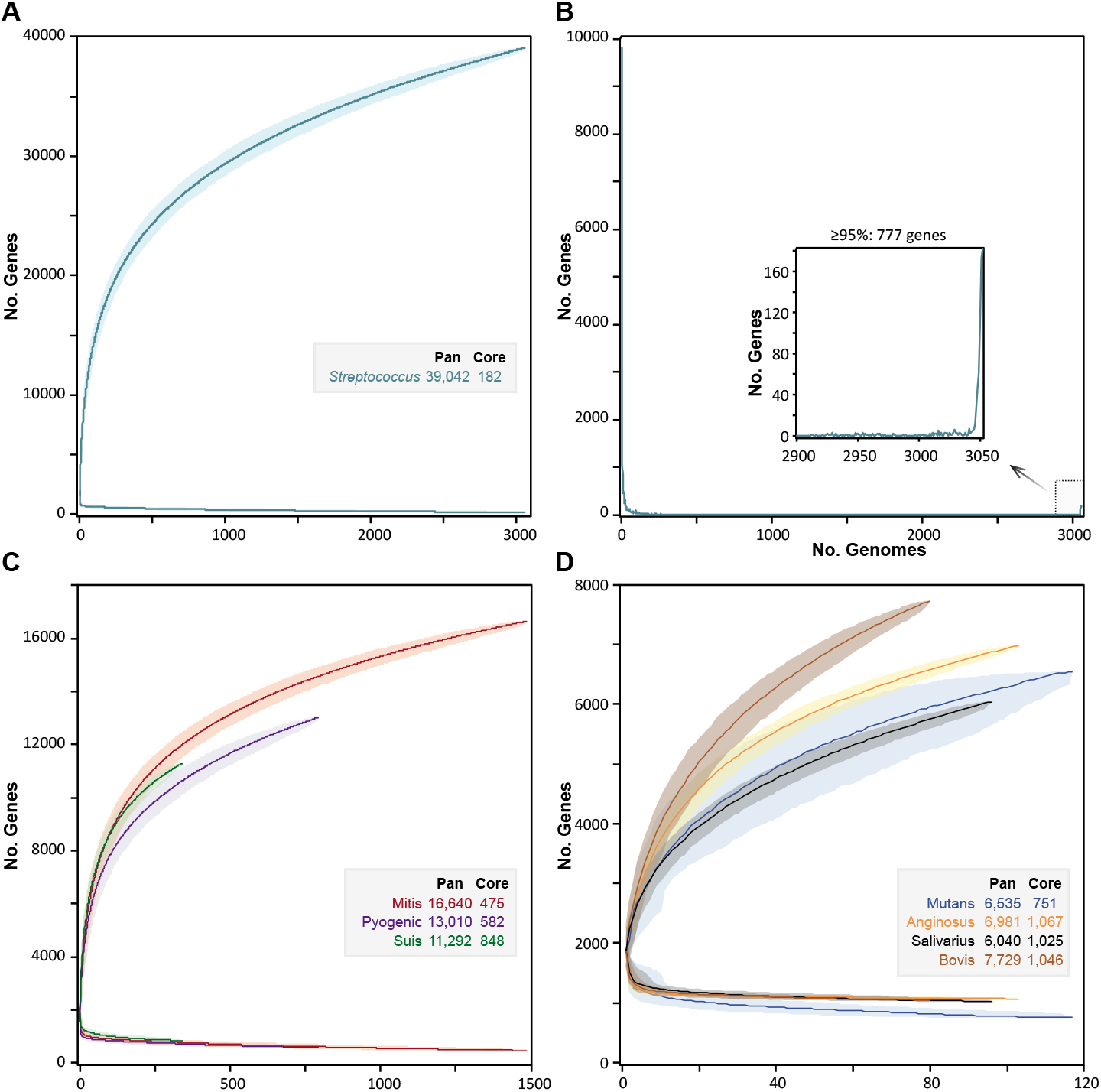
Rarefaction curves of pan- and core-gene numbers in *Streptococcus* and its seven major taxonomic subgroups. (A) Rarefaction curves created with PEPPAN_parser for the accumulations of pan genes and core genes of 3052 *Streptococcus* representative genomes from 1000 random permutations. (B) The frequencies of pan genes (Y-axis) by the numbers of genomes that carried that many genes (X-axis). The inset shows the relaxed core genes present in ≥ 95% of the genomes. (C) Rarefaction curves of genomes in the Mitis, Pyogenic and Suis groups. (D) Rarefaction curves of genomes in the Mutans, Anginosus, Salivarius and Bovis groups. The dark lines in figures A, C and D indicate median values and the shadows indicate 95% confidence intervals.

In contrast to earlier studies (Gao *et al*. 2014), which defined a strict core genome of 278 orthologs, we found only 182 genes that were shared across all *Streptococcus* genomes (Fig 4B, inset). Each of these was disrupted in at least one of the 14,115 *Streptococcus* genomes in RefSeq. This is a common problem for core genome analyses, especially because the multiple contigs within draft genomes can result in the absence of multiple genes from genome assemblies. Core genome schemes used for cgMLST are therefore usually based on a relaxed core, consisting of single-copy genes present in the large majority of representative isolates (Moura *et al*. 2016; Alikhan *et al*. 2018; Zhou *et al*. 2020). Our analyses identified 754 genes that were present in at least 2900 (95%) of the representative streptococcal genomes (Supplemental Table S3, Fig. 4B). However, most of the 754 genes were present in multiple copies in some genomes, leaving a final relaxed core of 292 single-copy genes that are suitable for identifying core genomic relationships and evolutionary history (Table 1; Supplemental Table S3).

**Table 1.** Summary statistics of the pan-genome of *Streptococcus* genus and seven species groups.

|  | Genomes | Number per genome |  |  | 95% of genomes |  | All genomes |  |  |
| --- | --- | --- | --- | --- | --- | --- | --- | --- | --- |
| Traditional Group |  | Genes | CDSs | % pseudogenes | All genes | Single copy | Strict core genes | Strict core CDSs | Pan genes |
| *Streptococcus* | 3052 | 1918 | 1810 | 5.6 | 754 | 292 | 182 | 5 | 39,042 |
| Anginosus | 103 | 1821 | 1720 | 5.5 | 1144 | 648 | 1067 | 722 | 6,981 |
| Bovis | 80 | 1877 | 1768 | 5.8 | 1184 | 559 | 1046 | 583 | 7,729 |
| Mitis | 1485 | 1970 | 1849 | 6.1 | 1087 | 642 | 475 | 36 | 16,640 |
| Mutans | 117 | 1858 | 1747 | 6.0 | 1110 | 380 | 751 | 248 | 6,535 |
| Pyogenic | 792 | 1791 | 1712 | 4.4 | 979 | 439 | 582 | 149 | 13,010 |
| Salivarius | 96 | 1865 | 1727 | 7.4 | 1217 | 595 | 1025 | 666 | 6,040 |
| Suis | 342 | 2059 | 1948 | 5.4 | 1281 | 672 | 848 | 570 | 11,292 |

Strict core refers to the number of genes that were found by DIAMOND and BLASTn in all genomes. Strict core CDSs refers to the number of genes which are not a pseudogene in any genome.

### Taxonomic clusters within *Streptococcus*

*Streptococcus* taxonomy is a highly dynamic area of research (Kikuchi *et al*. 1995; Jensen *et al*. 2016; Dekker and Lau 2016; Velsko *et al*. 2018; Velsko *et al*. 2019; Kilian and Tettelin 2019; Zhou *et al*. 2020). Many *Streptococcus* species are currently defined exclusively by phenotypic markers, and multiple taxonomic assignments in RefSeq are incorrect (Beaz-Hidalgo *et al*. 2015; Gomila *et al*. 2015; Kilian and Tettelin 2019). We therefore initially ignored taxonomic designations, and used the normal cut-off of ANI ≥95% as a proxy for species designations (Konstantinidis *et al*. 2017; Jain *et al*. 2018). Single-linkage agglomerative clustering of pair-wise ANI values calculated from the 3052 representative genomes revealed 223 clusters (Supplemental Table S2). For the 29 clusters containing ≥10 genomes, we also identified a dominant species designation from NCBI metadata, as shown in Supplemental Table S4. Information on each cluster’s pan-genome can be found in Supplemental Text 4 and Supplemental Table S5.

We used PEPPAN_parser to generate two trees of the 3052 representative genomes based on the presence or absence profiles of 39,042 pan genes (Fig. 5A) and on the allelic variation profiles of 292 relaxed core genes (Fig, 5B). The topology of the first tree reflects similarities in pan-genome content and the topology of the second tree reflects sequence similarities within core genes. Unsurprisingly, the details of these two topologies differed somewhat. In particular, the core gene tree contained an unresolved, star-like radiation which we attribute to distinct sequences in all of the core genes from highly diverse species. However, despite these differences in deep branching topology, both trees showed comparable tight clustering of genomes corresponding to each of the 29 common taxonomic groupings. This tight clustering indicates that the topologies of both trees are congruent at the ANI95% level. Both trees also support published taxonomic assignments of subspecies. For example, MG_29 corresponds to *S. gallolyticus* and includes its three subspecies *gallolyticus, macedonicus*, and *pasteurianus* (Dekker and Lau 2016). Similarly, MG_2 corresponds to *S. dysgalactiae* and includes its two subspecies *dysgalactiae* and *equisimilis* (Jensen and Kilian 2012).

**Figure 5.**
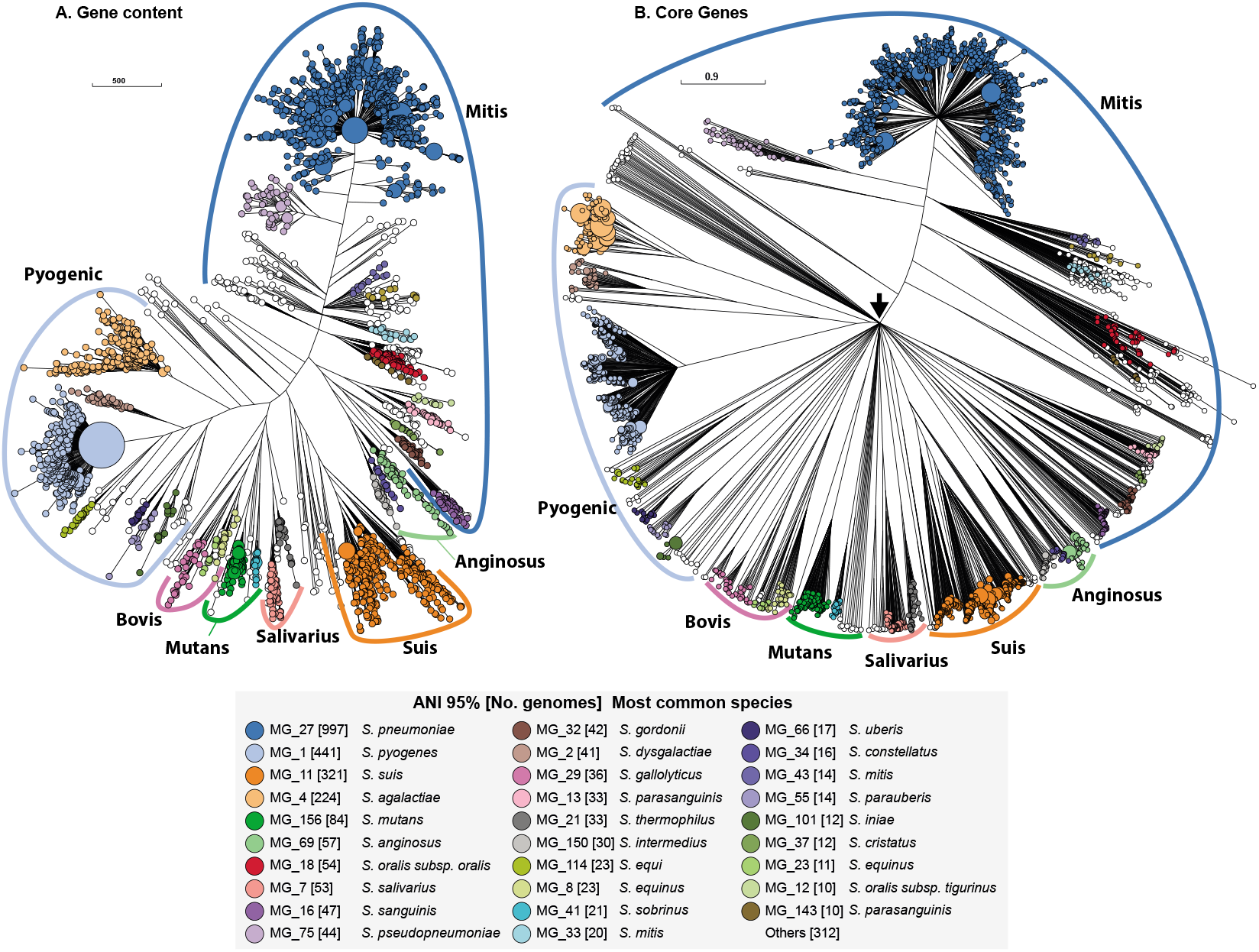
Phylogenies of 3052 *Streptococcus* genomes based on accessory gene content (A) and allelic variation in relaxed core genes (B). (A) A FastTree (Price *et al*. 2010) phylogeny based on binary information of the presence and absence of accessory genes in. (B) A RapidNJ (Simonsen *et al*. 2011) phylogeny based on numbers of identical sequences (alleles) of 292 single copy, relaxed, core genes that are present in ≥95% of *Streptococcus* genomes. These trees are represented in GrapeTree (Zhou *et al*. 2018a). The sizes of the circles in A and B are proportional to the numbers of genomes they encompass, and are color-coded by 29 common ANI95% clusters as shown in the inset. Many *Streptococcus* species have been assigned to one of six traditional taxonomic groups whose names are shown outside colored arcs. These trees define from the Suis group which contains *S. suis*. A black arrow in Figure B shows the root of the tree, where multiple branches radiate directly outwards due to lack of resolution of cgMLST for such distant taxa. All ANI95% cluster information can be found in Supplemental Table S4. Interactive versions of the trees can be found at (A) https://achtman-lab.github.io/GrapeTree/MSTree_holder.html?tree=https://raw.githubusercontent.com/zheminzhou/PEPPAN_data/master/Strep.gene_content.json; (B) https://achtman-lab.github.io/GrapeTree/MSTree_holder.html?tree=https://raw.githubusercontent.com/zheminzhou/PEPPAN_data/master/Strep.CGAV.json

Both *Streptococcus* trees also clustered high order branches according to the traditional taxonomical group names Mitis, Anginosus, Salivarius, Mutans, Bovis, and Pyogenic (Gao *et al*. 2014). They clustered *S. suis* together in a seventh phylogenetic branch, which we designate as Suis, and also clustered *S. acidominimus, S. minor, S. hyovaginalis, S. ovis*, and multiple other taxa into a novel, unnamed neighboring branch. Using PEPPAN_parser, we calculated a pan-genome for each of the seven named taxonomical groups. Similar to the *Streptococcus* pan-genome, each group pangenome is open (Fig. 4C-D), and grows at the rate of 3.5-30.1 new genes for each new representative genome. Unlike the entire genus, these seven named taxonomic groups possess a sizable strict core genome, consisting of 475 to 1067 core genes (Fig. 4C-D, Table 1). After excluding multi-copy genes, the sizes of the group-specific, 95% relaxed core genomes ranged from 380 (Mutans) to 672 (Suis) genes (Fig. 4C-D, Table 1).

In accord with prior observations (Kilian *et al*. 2008; Kilian and Tettelin 2019), numerous discrepancies differentiate the ANI95% groups and the taxonomic designations in RefSeq. Some discrepancies reflect inaccurate metadata, but others reflect true discrepancies between ANI95% clusters and taxonomic designations made by expert microbiologists. For example, *S. mitis* spans 44 distinct ANI95% clusters (Fig. 5, Supplemental Table S4). Similarly, *S. oralis* straddles multiple, distinct ANI95% clusters, as did each of the three *S. oralis* subspecies *oralis, tigurinus* and *dentisani* defined by Jensen *et al*. (Jensen *et al*. 2016). Further investigations will be needed to elucidate how many biological species are truly present within the genus *Streptococcus*. We anticipate that the trees in Fig. 5 might be useful for such analyses.

## Discussion

### Comparison of PEPPAN with other pan-genome pipelines

Pan-genome pipelines must be efficient in order to handle the computational demands of modern, large-scale comparative genomics. Roary (Page *et al*. 2015) and PIRATE (Bayliss *et al*. 2019) were the fastest of all the pipelines tested, likely reflecting their choice of time efficient approaches in every stage of their algorithms. However, this speed comes with tradeoffs in terms of accuracy (Figs. 2, 3). The workflow implemented in PEPPAN requires many more calculations than other pipelines due to its implementation of tree-based splitting of paralogs and similarity-based internal gene prediction, but is only marginally slower because of the care that was taken to implement time efficient algorithms.

Roary, PIRATE and PEPPAN all use a pre-clustering step to reduce the numbers of genes that are analyzed in subsequent, very time-consuming all-against-all comparisons. PEPPAN accelerates this step by using Linclust (Steinegger and Soding 2018). Linclust scales linearly with the number of genes, and is faster than CD-HIT, the clustering package used by Roary and PIRATE.

Roary and Pirate both use MCL, a graph-based clustering approach (Enright *et al*. 2002) to split paralogous clusters. MCL identifies a strict optimal threshold that separates orthologous genes from paralogous genes, and scales well with the numbers of genes. This approach is accurate for closely related genomes, but is error-prone when datasets contain both closely-related and distantly-related genomes, because a single optimal clustering threshold does not exist for both extremes. PIRATE thus failed to split many paralogous clusters from real (Figs. 2D, E) and simulated (Figs. 3B, C) genomes, especially for more diverse datasets (Fig. 3A). Roary implements an additional synteny-based approach to identify and split unresolved paralog clusters, but this approach also failed to correctly split orthologs into multiple clusters (Figs. 2D, E, 3B, C). In contrast, PEPPAN identifies an optimal threshold for each gene and uses that threshold to split paralogous branches in the gene trees. This allows accurate estimates of pan-genomes even in datasets of highly divergent genomes.

PanX uses a “divide and conquer” strategy for the gene comparisons, which is computationally demanding. In addition, PanX constructs a gene tree for every potential gene cluster, which, similar to other tree-based approaches, involves the alignment of gene sequences using MAFFT (Katoh and Standley 2013) followed by a tree construction using FastTree (Price *et al*. 2010). As a result, PanX is substantially slower than PEPPAN, PIRATE or Roary (Fig. 3E). However, PanX was not substantially more accurate than those programs (Figs. 2A, 3A), which might be attributed to its use of raw pairwise genetic distances of genes for paralog splitting. In contrast, inspired by the methods used by large-scale genomics studies (Chewapreecha *et al*. 2014; Banaszkiewicz *et al*. 2019), PEPPAN uses a reference-based approach to generate an alignment for each gene group, which is then used to reconstruct a neighbor-joining gene tree using RapidNJ. These methods are less accurate, but much faster than those in PanX, and scale to thousands of sequences. As a result, although the run time of PEPPAN was approximately twice as long as the run time of Roary or PIRATE, it still scaled linearly with the number of genomes (Fig. 3E).

### Effects of internal annotations by PEPPAN

Our benchmarking analyses on real and simulated genomes revealed the strong impact of inconsistent annotations on the pangenome predictions (Fig. 2A). Indeed, differences in annotation influenced the quality of the pan-genome more than pipeline algorithms (Fig. 2B), and decreased the number of core genes by up to one-third for some pipelines (Fig. 2A). PEPPAN avoids this problem by implementing a similarity-based gene prediction step. Accordingly, pangenomes predicted by PEPPAN varied only slightly with different annotations (Fig. 2A, B). Draft genome assemblies based on 454 or IonTorrent sequencing include elevated numbers of single-base insertions and deletions due to inaccurate sequencing (Shao *et al*. 2013; Zhang *et al*. 2015). Including such genomes in an analysis reduces the quality of the pan-genome for all state-of-the-art pipelines. However, PEPPAN simply scores genes disrupted by artificial INDELs as frameshifts, making such inaccurate genomes easier to identify.

Finally, it is worth noting that the current similarity-based internal annotation algorithm implemented in PEPPAN is optimized for prokaryotes, and does not work for eukaryotic genomes, where multiple exons of a gene can be separated by introns of >10kb. Apart from this limitation, however, the other technological advantages in PEPPAN will also work on eukaryotic genomes. PEPPAN could therefore be extended for use on eukaryotes with collaboration from experts in eukaryotic genomics.

### Relevance to MLST schemes

Alikhan *et al*. (2018) described a pan-genome for all of *Salmonella* based on 537 genomes that had been derived by a precursor of PEPPAN in 2015. That pan-genome was used to develop a wgMLST scheme of 21,065 loci and a cgMLST scheme of 3002 genes. The same publication also described a reference set of 926 genomes that represented the diversity of almost 120,000 *Salmonella* genomes on the basis of rMLST. After completion of this manuscript, we became aware of a new publication (Park and Andam 2020) which used Roary to calculate a pan-genome of 84,041 *S. enterica* genes and 2085 soft-core genes from those 926 representative genomes after re-annotation with PROKKA. Such applications of Roary are strongly discouraged by its documentation, which recommends against using Roary on diverse groups of organisms such as all *Salmonella*. We ran PEPPAN on the same 926 representative genomes. The resulting pan-genome contained 30,000 fewer pan-genes and 1200 more soft-core genes than the calculations by Park and Andam (Supplemental Table S6), confirming that Roary struggled with this task. The high resolution and continued reliability that EnteroBase offers in downstream analyses of phylogenetic relationships between genomes are in part due to the accurate, smaller pan-genome and larger core-genome that was calculated by PEPPAN. The analyses presented here identified a reliable relaxed soft-core genome consisting of 292 single copy genes for *Streptococcus*, which is currently being used to establish an EnteroBase database for this diverse genus.

### Pan-genomes depend on sample size

Early analyses of pan-genomes were based on small numbers of genomic sequences (Tettelin *et al*. 2005), resulting in the conclusion based on 12 genomes that the pan-genome of *Streptococcus pyogenes* was closed (Tettelin *et al*. 2008). The same publication concluded that the pan-genome of *Streptococcus pneumoniae* was open and would continue to expand indefinitely. However, a subsequent study of 44 genomes concluded that the pan-genome of *S. pneumoniae* was also closed (Donati *et al*. 2010). It is only very recently that large numbers of bacterial genomes are available for analysis, and that pipelines exist that can handle such large numbers.

We calculated a pan-genome from 3052 *Streptococcus* genomes that represent the genomic diversity of 14,115 draft and complete genomes. Our pan-genome contains 39,042 genes, is open, and will continue to expand at a rate of 4.4 genes per novel, genome. This rate of expansion is 14fold slower than the original calculations of a pangenome from 138 genomes (Gao *et al*. 2014). We also calculated pan-genomes and their expected growth rates for the 29 most common ANI95% clusters within *Streptococcus* (Supplemental Text 4). All pan-genomes were open, with the single exception of MG_41 (*Streptococcus sobrinus*). These inconsistencies with prior analyses suggest that pan-genome status may be strongly dependent on the number of genomes investigated, sampling strategies used to identify representative genomes, and possibly on pan-genome pipelines.

### Taxonomic insights

It has been clear since 2004 that the strict core genome of all prokaryotes is extremely small. Only 14-30 genes were present in all of 147 diverse genomes (Charlebois and Doolittle 2004), and almost all of those genes encoded ribosomal proteins (Weiss *et al*. 2018). However, it was still unexpected that the strict core genome would be this small for a large collection of *Streptococcus* genomes. We only found 182 strict core genes in the representative set of 3052 genomes, and each of these was absent or incomplete in one or more of the entire set of 14,115 genomes. We therefore recommend using phylogenies based on sequence variation within a relaxed core complement of genes and/or presence/absence of accessory genes for an overview of the phylogenetic relationships of an entire genus instead of relying only on strict core genes. PEPPAN_parser can calculate such phylogenies from the PEPPAN outputs.

As previously noted by others (Kilian *et al*. 2008; Jensen and Kilian 2012; Jensen *et al*. 2013; Jensen *et al*. 2016; Kilian and Tettelin 2019), the taxonomies of multiple *Streptococcus* genomes are misclassified in RefSeq (Supplemental Fig. S3). Misclassification has been ongoing for decades (Kikuchi *et al*. 1995) due to the phenotypic heterogeneity of this species. The Mitis group is particularly heterogeneous (Kilian *et al*. 2008; Jensen *et al*. 2016) and difficult to study (Kilian and Tettelin 2019; Velsko *et al*. 2019). Similar problems also apply to other bacterial genera such as *Pseudomonas* (Gomila *et al*. 2015) and *Aeromonas* (Beaz-Hidalgo *et al*. 2015). The results presented here defined 223 ANI95% clusters which are consistent by independent phylogenetic approaches based on both cgMLST and gene presence. It has been suggested that bacterial diversity does not delinieate species clusters due to extensive HGT (Doolittle and Papke 2006). Our results, instead, revealed congruent clusters between the accessory genome and the core genome at the ANI95% level in *Streptococcus*. Similar congruent clusters have been reported in the *Streptomycetaceae* (Wright and Baum 2018) and we suspect that they will also occur in other genera. Thus, approaches such as those described here may provide a framework for improving future taxonomic assignments. Finally, the test case of *Streptococcus* illustrates the power of PEPPAN, which can now be used for defining the pan-genomes of other diverse genera.

## Methods

### *S. enterica* genomes

We downloaded the assembly_summary_genbank.txt table and the assembly_summary_refseq.txt table from NCBI on 30th May of 2019 (ftp://ftp.ncbi.nlm.nih.gov/genomes/ASSEMBLY_REPORTS/). The first table summarizes all genomes uploaded into GenBank by their original authors and the second summarizes all the genomes in RefSeq. We used these tables as a source of the FTP links for each of the accession codes listed by Nuccio and Bäumler (Nuccio and Bäumler 2014) for genomic sequences of 15 *S. enterica* genomes. These 15 genomes were also annotated *ab initio* with PROKKA 1.12 (Seemann 2014). Nuccio and Bäumler excluded some “unreliable” short genes from their manual re-curation. In order to exclude these genes in our analyses as well, the genomic coordinates of each gene in each of the three annotations (Submitter, RefSeq, PROKKA) were compared with the coordinates of “reliable genes” in Table S1 of Nuccio and Bäumler. Only genes with co-ordinates overlapping those of a reliable gene by ≥90% were used here for further comparisons.

### Preparation of simulated datasets

All simulated datasets were generated using SimPan (Supplemental Text 2) with the input parameters “--genomeNum 15 --aveSize 4500 --pBackbone 4000 --nMobile 10000 --nCore 3621 --pBackbone 0.05 --pMobile 0.40 --rec 0.1”. Datasets a through e were generated with the additional parameters: (a) ‘--idenOrtholog 0.9 --idenParalog 0.6’; (b) ‘--idenOrtholog 0.95 --idenParalog 0.6’; (c) ‘--idenOrtholog 0.98 --idenParalog 0.6’; (d) ‘-- idenOrtholog 0.98 --idenParalog 0.8’; (e) ‘--idenOrtholog 0.98 --idenParalog 0.9’. Ten other sets of simulated genomes that were used to evaluate running times were generated with the same parameters as dataset c but with the additional parameter ‘--genomeNum xxx’, where xxx ranged from 20 to 200 by steps of 20.

### Pan-genome pipelines

The following versions of the individual pipelines and command lines were used for all benchmark datasets.

1. **PEPPAN** with a Git HEAD of f721513 was run in the Python 3.6 environment as:

~~~
python PEPPA.py -t 4 -p PEPPAN --pseudogene 0.9 --min_cds 45 *.gff
~~~
2. **Roary** 3.6.0+dfsg-4 was installed as a Ubuntu APT package and run as:

~~~
roary -p 4 -o roary -f roary -i <identity> -s -v -y *.gff
~~~ Three runs of Roary were performed for each dataset with the additional parameters “-i 50”, “-i 80” or “-i 95”. The data reported here are from the runs with the parameter ‘-i 80’ because that consistently yielded the best ARI values.
3. **PIRATE** with a Git HEAD of effc522 was downloaded from https://github.com/SionBayliss/PIRATE and run as:

~~~
PIRATE -i . -o PIRATE -s <steps> -t 4 -k “--diamond”
~~~ Three runs of PIRATE were performed for each dataset with the additional parameters “-s 50,60,70,80,90,95,98”, “-s 80,90,95,98” or “-s 95,98”. We report the data generated with “-s 80,90,95,98” which had the greatest ARI value, except for simulated dataset e, where “-s 95,98” had the greatest ARI.
4. **PanX** v1.6.0 was downloaded from https://github.com/neherlab/pan-genome-analysis/releases and run in the Python 2.7 environment as:

~~~
panX.py --folder_name panX --species_name panX --threads 4 --diamond_identity 80 --simple_tree --store_locus_tag
~~~
5. **OrthoMCL** v2.0.9 was downloaded from https://orthomcl.org/ and run in multiple steps as described in https://currentprotocols.onlinelibrary.wiley.com/doi/full/10.1002/0471250953.bi0612s35.

### Generating ANI95% clusters of *Streptococcus* genomes

A summary table of all genomes deposited in RefSeq was downloaded on 20 June of 2019 (see *S. enterica* genomes above). 14,115 bacterial records that contained “Streptococcus” in the “organism_name” field were extracted from the table (Supplemental Table S7), and the files for each record were downloaded as described above. MASH (Ondov *et al*. 2016) was used to measure the pairwise distances between the genomes with parameters of ‘-k 19 –s 10000’. The resulting matrix was used to cluster *Streptococcus* genomes with the AgglomerativeClustering function in the scikit-learn package (Pedregosa *et al*. 2011), with parameters linkage=single and distance_threshold=0.002. The function generated 3170 clusters. The genome with the greatest N50 value within each cluster was chosen as its representative genome. Each representative genome was subjected to quality evaluation according to three criteria: (1) carries at least 38 of the 40 single-copy essential genes according to fetchMG (Sunagawa *et al*. 2013); (2) is assigned to *Streptococcus* genus by the ‘Identify species’ function in rmlst.org (Jolley *et al*. 2012); (3) has an N50 value ≥10 kb. 118 genomes failed these criteria and were discarded (Supplemental Table S2B), leaving a dataset of 3052 high-quality genomes (Supplemental Table S2A) that represents the entire taxonomic diversity of *Streptococcus*. Pair-wise ANI values were calculated from the 3052 representative genomes with FastANI v1.2 (Jain *et al*. 2018), and these genomes were grouped into ANI95% clusters using the AgglomerativeClustering function with linkage=single and distance_threshold=0.05.

## Data Access

Source code for PEPPAN is accessible at https://github.com/zheminzhou/PEPPAN and as Supplemental Code S1. Source code for SimPan is accessible at https://github.com/zheminzhou/SimPan and as Supplemental Code S2. Trees presented in Figure 5 are available separately: (A) https://achtman-lab.github.io/GrapeTree/MSTree_holder.html?tree=https://raw.githubusercontent.com/zheminzhou/PEPPAN_data/master/Strep.content.json; (B) https://achtman-lab.github.io/GrapeTree/MSTree_holder.html?tree=https://raw.githubusercontent.com/zheminzhou/PEPPAN_data/master/Strep.CGAV.json.

## Acknowledgements

This project was supported by the Wellcome Trust (202792/Z/16/Z) and EnteroBase development was funded by the BBSRC (BB/L020319/1). We gratefully acknowledge helps on the interpretation of *Streptococcus* taxonomy by Mogens Kilian, Daniel Jaén Luchoro and Francisco Salva Serra, and critical comments on the text from Nina Luhmann.

## Author Contributions

Z.Z. developed the pipelines, analyzed data and prepared the figures. M.A., J.C., and Z.Z. interpreted the results and wrote the manuscript.

