## Supplemental Text for "Accurate reconstruction of bacterial pan- and core- genomes with PEPPAN"

### Supplemental Text 1. PEPPAN: Phylogeny Enhanced Prediction of a PAN-genome

Here we describe the detailed steps of the PEPPAN pipeline in Supplemental Fig. S1.

#### 1.1 Identifying representative gene sequences

##### 1.1.1 Iterative clustering of genes

The default input for PEPPAN is a set of pre-computed gene annotations in GFF3 format (Ensembl Release 98 2019). PEPPAN then extracts all annotated CDS features from that input set. PEPPAN can also include other annotated features such as ncRNAs in its analysis if they are specified using the '—feature' option. PEPPAN can also use externally curated gene sets as reference gene sequences for its re-annotation step, such as those from a wgMLST scheme.

PEPPAN runs Linclust (Steinegger and Soding 2018) iteratively on the entire set of input genes to identify representative gene clusters for the input gene set, starting with clusters of 100% nucleotide sequence identity, and iteratively lowering the identity threshold in ten 1% steps to 90% sequence identity. In each round, sequences with less than 80% coverage across the aligned region are excluded from the cluster. After each iteration, only the longest sequence is retained for each cluster, and these are then used as input to the next clustering round. The output from the final step consists of a set of one representative gene sequence for each 90% sequence homology cluster, which is then used for downstream analyses.

##### 1.1.2 Minimum spanning trees from all-against-all comparisons of representative genes

BLASTn (Altschul *et al.* 1990) and DIAMOND (Buchfink *et al.* 2015) are used to compute all-against-all comparisons of all representative gene sequences, and scores for the quality of pairwise alignments of gene sequences. PEPPAN then calculates a collection of discrete minimum spanning trees (MSTs) from scores in which ≥40% of the codons (in the shorter gene) were aligned, and which shared ≥40% identity across the alignment.

##### 1.1.3 Genetic distances between genomes

PEPPAN extracts a set of the most common genes across all the genomes to calculate a proxy for the average genetic distances between genomes. That proxy is used in section 1.3.2 to evaluate pairwise genetic distances of individual genes. This goal requires excluding paralogs. To this end, all genes are removed from MSTs in which multiple sequences in the cluster originated from a common genome, leaving only single copy genes from different genomes. As a result, the greater the number of remaining single-copy genes per MST, the most likely they are to constitute part of the core genome. PEPPAN identifies the MSTs with the greatest number of genes, up to a maximum of 1000 MSTs, and uses that set as a proxy for the core genome. for an estimate of average genetic distance between genomes.

Pair-wise genetic distances of individual genes from each MST are calculated based on their membership within the representative gene clusters identified in section 1.1.1: (1) for two genes belonging to the same 90% identity cluster, the genetic distance equals one minus the largest clustering threshold that was needed to cluster them; (2) for two genes from different 90% identity clusters, the genetic distance equals the value of the most distant edge in the MST connecting these two clusters (section 1.1.2). Pairwise genetic distances of individual genes are summarized over all MSTs and a log-normal distribution is fitted for each pair of genomes, yielding an estimate of average genetic distance ($\bar{D}$) and standard deviation ($\sigma$) for each pair of genomes.

#### 1.2 Identifying all gene candidates in all genomes

In order to identify genes independently of individual genome annotations, each representative gene sequence is aligned to all genome sequences using BLASTn and DIAMOND. DIAMOND alignments are then back-translated into nucleotide alignments. Alignment scores from both tools are re-calculated with a scoring scheme of +3 and -1 for each matched and mismatched site and -6 and -1 for gap-opening and extension penalties. When a representative gene is aligned to the same region of a genome by both tools, only the highest-scoring alignment is kept. Otherwise both alignments are kept. By default, PEPPAN considers all alignments as putative gene candidates if they span at least 50% of the length of a representative gene with 50% nucleotide sequence identity. This identity threshold is much lower than the average nucleotide identity (68-85%) for 144 genera between genomes from the same genus (Barco *et al.* 2020), and thus allows the identification of most of the gene candidates from an entire genus. All gene candidates for each representative gene sequence are retained that pass these filtering rules. Each putative gene candidate is scored by comparing it with the representative gene as well as according to the gene annotations in the original GFF3 file. PEPPAN calculates this score *s* as:

$$s = (l * i) * \sqrt{l * (r_q * r_r)^{1/2}}$$

where $l$ is the length of the open reading frame of the putative gene candidate, $i$ is the fraction of identity between the gene candidate and the representative gene, $r_q$ is the aligned length of the gene candidate relative to the representative gene and $r_r$ is the length of the portion of the gene candidate that overlaps an annotated gene in the original GFF3 annotation divided by the total length of the gene candidate. If a gene candidate does not overlap an annotated gene by at least 0.1 of its length, $r_r$ is set to 0.1. These scores are saved for subsequent use during the filtering of conflicting gene candidates in section 1.3.4.

#### 1.3 Identifying clusters of orthologous genes

##### 1.3.1 Gene candidate alignment and phylogeny estimation

Pseudo-sequences that are easy to handle are generated for each gene candidate in order to facilitate accurate multi-sequence alignments. These consist of a copy of the corresponding representative gene sequence except that the polymorphic sites are replaced by the nucleotide variants from the gene candidates, and the sites that are missing in the alignments are replaced by "-". This procedure results in pseudo-sequences whose lengths are identical to that of the reference allele and yield the desired multiple alignment without further processing. RapidNJ (Simonsen *et al.* 2011) is then used to build a gene tree from the multiple alignment for each set of gene candidates (Supplemental Fig. S1F). These alignments and trees are used in the next section for identification of paralogous genes by comparing them with the average genetic distances between the genomes.

##### 1.3.2 Phylogeny-based paralog splitting

Each branch in a gene tree separates the genes into two sub-clusters. Genes are scored as orthologs when the branch between them is compatible with a descent from a common ancestor (Supplemental Fig. S1E-H). This decision is reached by comparing the branch lengths separating two sub-clusters of genes with the average genetic distance between their genomes of origin.

Consider the simple scenario a sub-cluster of two gene candidates, *i* and *j*,. PEPPAN calculates the paralogous score for these two gene candidates as:

$$p_{i,j} = \frac{d_{i,j}}{e^{\bar{D}_{i,j}+3*\sigma_{i,j}}}$$

where $d_{i,j}$ is the genetic distance between the two genes, and $\bar{D}_{i,j}$ and $\sigma_{i,j}$ are the distance and variance for the log-normal distribution of the genetic distance between the two genomes carrying these genes (section 1.1.3), except when both gene candidates are co-present in a common genome in which case it is arbitrarily set to $\bar{D}$=0.005 and $\sigma$=2. A value of $p_{i,j}$>1 indicates that the genes represent a pair of paralogs that most likely diverged prior to the common ancestor of their genomes, because the cumulative sequence differences between the genes are greater than 3$\sigma$ of the average genetic distance of their genomes (equivalent to a p-value of < 0.0028). These genes are therefore classified as paralogs. Otherwise they are putative orthologs.

This calculation can be extended to two sub-clusters of genes (*M* and *N*) by calculating a weighted average of the paralogous scores for all genes in the two groups as:

$$\bar{p} = \frac{\sum_{i \epsilon M} \sum_{j \epsilon N} p_{i,j} * e^{-\bar{D}_{i,j}}}{\sum_{i \epsilon M} \sum_{j \epsilon N} e^{-\bar{D}_{i,j}}}$$

The *p* scores are weighted by the inverse of the genetic distances of the respective genomes, because paralogous genes are more easily identified by comparing closely related genomes than more divergent ones. Each gene tree is split iteratively, starting at the branch with the greatest $\bar{p}$ value if $\bar{p}$ >1 (Supplemental Fig. S1H). Branches in each of the sub-trees are scored and subjected to further splits until all branches have a value of $\bar{p}$ ≤1. Each of the resulting sub-trees thus represents a putative ortholog group.

##### 1.3.3 Resolve conflicting gene candidates

The locations of multiple putative orthologs can overlap in some genomes according to genomic annotations. Such conflicts can come from two sources, which each needs a distinct approach for its resolution.

(1) Inconsistent gene predictions for a corresponding genomic region across multiple genomes. In order to resolve such conflicts, a summed information score $\hat{s}$ is calculated for every ortholog group as:

$$\hat{s} = \sum_{g \in G} s_g$$

where $s_g$ is the information score for every gene candidate *g* in the ortholog group *G* that was calculated in section 1.2. PEPPAN retains the ortholog group with the greatest summed information score, and discards other groups.

(2) Some orthologous genes that are assigned to distinct homology groups by hierarchical clustering (section 1.1.1) may be linked in a common gene MST (section 1.1.2) even though their sequence identities are <90% or their alignment coverage <80%. Gene candidates from such linked representative genes may result in overlapping regions in some genomes. To identify this type of conflict, PEPPAN compares the information scores of pairs of overlapping ortholog groups, and merges the pairs when more than 1/3 of the mappings of the gene candidates in each ortholog group overlap, and their representative genes are from the same gene MST (section 1.1.2).

##### 1.3.4 Use synteny to identify and remove recent duplicates

Paralogous genes with ≥99% identity are likely to have recently arisen by gene duplications, and cannot be detected by phylogeny-based paralog analysis. However, the neighboring genes will differ between duplicated gene candidates in distinct regions of the genome. In order to test for such paralogs, all gene candidates from each genome are ordered by their genomic coordinates, and PEPPAN extracts up to three neighboring genes upstream and downstream of each gene candidate in a putative orthologs group. A minimum spanning tree is calculated based on the number of unique neighbors for each pair of gene candidates, and the final set of orthologous genes is arbitrarily restricted to those clusters of gene candidates with branch lengths of no more than 4 different neighboring genes.

#### 1.4 Pseudogene identification

Gene candidates are selected based on aligning each representative gene to all genomes (section 1.2), and might include pseudogenes because open reading frames (ORFs) are not considered in the alignment. The option '—pseudogene' controls the minimum acceptable length of the CDS in a gene candidate relative to the representative gene, with a default value of 80% (Lerat and Ochman 2004). The integrity of each gene candidate is filtered by this cutoff separately according to four criteria (Supplemental Fig. S1C): (1) ≥80% coverage of the representative gene; (2) the open reading frame contains ≥80% of codons in the representative gene, and is not broken by a frameshift or nonsense mutation; (3) a start codon is present ≤60 bp upstream of the 5' end of the alignment; (4) a stop codon is present at or downstream of the 3' end. Candidate genes that pass all four criteria are scored as coding genes, and all others as pseudogenes.

### Supplemental Text 2. SimPan: A pipeline for Simulating the evolution of Pan-genomes

The infinite many gene (IMG) model (Baumdicker *et al.* 2012) is arguably the simplest existing neutral mathematical model for the evolution of a pan-genome. In this model, accessory genes are inserted and deleted at fixed frequencies at nodes in a phylogenetic tree. The IMG model is the basis for several existing pan-genome simulators (Dalquen *et al.* 2012; Ferres *et al.* 2019), including the workflow in PanX (Ding *et al.* 2018). However, the IMG model does not match the U-shaped gene frequency distributions observed in multiple real bacterial pan-genomes, and which is thought to result from decreased gene insertion/deletion frequencies over long-term evolution (Collins and Higgs 2012; Lobkovsky *et al.* 2013). Alternative theoretical models have been proposed that account for the U-shaped distribution by a recent expansion of population size (Haegeman and Weitz 2012), as a consequence of long-term purifying selection (Lobkovsky *et al.* 2013; Zhou *et al.* 2014), or as the result of a special gene class that tends to be lost soon after it is acquired (Collins and Higgs 2012; Croucher *et al.* 2014). However, we were not aware of an existing pipeline for simulation of a pan-genome according to a U-shaped gene frequency distribution. Furthermore, none of the theoretical models account for homoplastic gene insertions/deletions, nor for large insertion/deletion events that span multiple consecutive genes.

SimPan was developed to simulate pan-genomes in a more realistic way by accounting for these issues. (1) SimPan infers homoplastic gene insertions/deletions by using SimBac (Brown *et al.* 2016) that includes homologous recombination events during the generation of local trees. (2) SimPan reshapes each local tree in a time-dependent manner, and reduces the long-term stability of gene insertions/deletions by penalizing the length of basal branches. (3) SimPan allows insertions/deletions of multiple consecutive genes.

#### 2.1 Simulation of two gene classes

SimPan first generates two gene classes according to input parameters (Supplemental Table S1): backbone genes (nBackbone: 4000) which are present in the ancestor of the population, and mobile genes (nMobile: 20,000) which are sources of horizontal genetic transfers. A pool of paralogous sources of recombination which resulted from historical gene duplications and are now only distantly related to each other by 60% identity (idenParalog) is created by choosing a random set of 5 percent of the backbone genes (pBackbone) and 40% (pMobile) of the mobile genes. The phylogenies of these paralogous genes are simulated using SimBac (Brown *et al.* 2016). Furthermore, SimPan assigns the sizes of the coding regions from a geometric distribution with an average length of 900 bp for each gene (geneLen). The sizes for 5'-intergenic and 3'-intergenic regions surrounding each coding region are assigned by drawing two additional numbers from a second geometric distribution, with an average length of 50 bp for each gene (igrLen). The total size of the sequences in the simulation equals the total sizes of backbone and mobile genes (nBackbone + nMobile), multiplied by the total number of genomes (genomeNum: 20).

#### 2.2 Simulation of local trees and INDEL trees

SimBac (Brown *et al.* 2016) simulates a clonal phylogeny of genomes, and then creates a local tree for every backbone and mobile gene to reflect the effects of random homologous recombination (rec: 0.05, recLen: 1000). SimPan reshapes each local tree into an INDEL tree for later use to simulate gene insertion/deletions by resizing the branch lengths in the trees according to their distances to the tips. This reshaping simulates an exponential increasing frequency of gene insertion or deletions along the branches from root to tips in the tree, which is consistent with observations from multiple real bacterial pan-genomes (Lobkovsky *et al.* 2013; Zhou *et al.* 2014) and results in a U-shaped gene frequency distribution as observed in natural genomes (Collins and Higgs 2012; Lobkovsky *et al.* 2013).

Given a reshaping factor $\lambda$=100, the new length ($l'$) of each branch in the INDEL tree is calculated as:

$$l' = \lambda * (e^{-\lambda * s} - e^{-\lambda * (s+l)})$$

where $l$ is the length of the branch in the local tree and $s$ is its distance to the tips.

#### 2.3 Simulation of insertion and deletion events

Each deletion or insertion event is simulated by choosing a random INDEL tree and a random branch in that tree. In the case of a deletion, the genome descending from that chosen branch is subjected to the deletion of a random block of backbone genes (default average block length (deletionBlock): 3 genes), which is propagated to all descendent genomes. Further random branches in random INDEL trees are subjected to deletion events until the core genome contains less than a user-defined number of genes (nCore: 3500). Subsequently random insertions of consecutive mobile genes (default average block length (insertBlock): 10) are performed on random branches of random INDEL trees until the average genome exceeds a user-defined maximum number of genes (aveSize: 4,500).

#### 2.4 Simulation of gene and genomic sequences

Coding and intergenic regions of the genes are simulated according to the M1 and HKY models, respectively, with the program INDELible (Fletcher and Yang 2009) using the local trees initially generated by SimBac. Additional short insertions and deletions (shortIndelLen: 10 bp) are also simulated with INDELible at a rate of 0.01 per mutation (shortIndelRate: 0.01). INDELible does not simulate start and stop codons. Therefore, the first and last codons of each coding region are replaced by a random bacterial start and stop codon according to the empirical distributions of start (ATG: 83%; GTG: 14%; TTG: 3%) and stop (TAA: 63%; TGA: 29%; TGA: 8%) codons in *E. coli* K-12 (Blattner *et al.* 1997). All gene sequences are then concatenated into genomic sequences, and stored as files in FASTA format for the generation of annotated genomes in GFF (Ensembl Release 98 2019) format.

### Supplemental Text 3. Using PEPPAN for whole-genome MLST schemes

We have used PEPPAN to create whole genome multi-locus sequence typing (wgMLST) schemes for multiple genera in EnteroBase (http://enterobase.warwick.ac.uk). These schemes have been used to identify genomes of *Salmonella* which contain a rare accessory gene (Chen *et al.* 2019) and to select subsets of conserved core genome loci for cgMLST genotyping schemes (Alikhan *et al.* 2018; Zhou *et al.* 2020). Here, we describe the manual curation steps of PEPPAN-generated pan-genomes that are needed to establish a reliable wgMLST scheme.

#### 3.1 Selection of a reliable set of reference genomes

PEPPAN re-annotates all genomes to ensure the consistency of gene annotations. By default, input genes are iteratively clustered and the longest sequence within each cluster is chosen as representative for that orhtolog cluster (section 1.1.1). The representative genes determine the start and end coordinates of the final pan-genes. This approach works well on input genomes that have been automatically annotated without manual curations. However, genome sets that are used to build wgMLST schemes often contain reference genomes whose annotations have been curated by experts and that include gene names that are widely accepted by the community (Blattner *et al.* 1997; Sebaihia *et al.* 2006). In other cases, annotations have been checked experimentally (Kröger *et al.* 2012). These manual annotations are likely to be more reliable than those predicted automatically. PEPPAN therefore allows users to incorporate such prior knowledge in the pan-genome, by defining a flexible priority order for input genomes using the “-P” (Priority) parameter. For example, we assigned each of the representative *Streptococcus* genomes into one of five priority levels before constructing the pan-genome (Supplemental Table S2), based on a combination of its “RefSeq category” (Tatusova *et al.* 2014) and assembly status. The five priority levels were: 5: a manually selected ‘gold standard’ reference genome with high-quality RefSeq annotation; 4: a complete representative genome; 3: a draft representative genome; 2: other complete genomes; 1: other draft genomes. When genomes of different priority levels are clustered, genes from genomes with the greatest priority are preferably selected as representatives of the gene cluster. In the case of multiple genomes with identically low priorities, the longest gene will be selected. For our *Streptococcus* dataset, these priority levels resulted in the representative genes for 753 of the 754 relaxed core genes having originated from genomes in priority level 1 (Supplemental Table S3).

#### 3.2 Identification of genes that are too similar

Each pan-gene identified by PEPPAN is assigned to a group of orthologs that has been screened for paralogs according to phylogeny- and synteny-based approaches. However, this approach is too complicated for assigning sequences from newly assembled genomes to loci and alleles by an MLST nomenclature server. Instead, the EnteroBase nomenclature server calls MLST loci and alleles in a novel genome by aligning a pre-defined set of representative sequences to that genome (Zhou *et al.* 2020). Due to sequence diversity and horizontal gene transfer, novel genomes may include sequence variants that align to multiple similar pan-genes, resulting in some cases to gene assignments to multiple representative gene sequences. The existence of such problematical representative pan-genes can be identified by the MLSTdb function in the EnteroBase EToKi package (https://github.com/zheminzhou/EToKi). EToKi MLSTdb runs an all-against-all comparison on all representative gene sequences generated by PEPPAN, and removes sets of genes with ≥70% identity at either the nucleotide or amino acid level. All wgMLST schemes in EnteroBase were subjected to this final cleaning stage before being made publicly available.

#### Supplemental Text 4. Pan-genomes of *Streptococcus* ANI95% clusters

The genomic analyses presented here identified 29 ANI95% clusters of *Streptococcus* which contained at least 10 genomes. A median frequency of 5.4% of their CDSs (range 3.2-12.9%) were scored as pseudogenes by PEPPAN, and the average numbers of pseudogenes per genome ranged from 220 (12.9% of all CDS) in the primary yoghurt fermentation bacterium (Bolotin *et al.* 2004; Hols *et al.* 2005; Goh *et al.* 2011), *S. thermophilus* (MG_21), down to 58 (3.2%) in a pathogen responsible for bovine mastitis) (Ward *et al.* 2009; Hossain *et al.* 2015), *S. uberis* (MG_66).

The pan- and core-genome sizes also varied dramatically between the 29 ANI95% clusters (Supplemental Table S5). For example, 321 genomes of *S. suis* (MG_11) yielded a pan-genome of 9,947 pan-genes with 989 core genes whereas 441 genomes of *S. pyogenes* (MG_1) defined a pan-genome of only 4,246 pan-genes with 1,249 core-genes. This may reflect sampling bias or differences in population size or demographics of individual species, but we have not pursued this issue.

To calculate the growth rates of the pan-genomes for each of the 29 ANI 95% clusters, we used PEPPAN_parser to fit a power law curve to rarefaction curves of the estimated size of the pan-genomes with increasing numbers of genomes (Tettelin *et al.* 2008) (Supplemental Table S5). The power-law parameter, α, gives an estimate of the growth trend of the pan-genome. A lower α suggests a faster growth rate of the pan-genome, whereas a higher value suggests a slower growth rate. In particular, the size of pan-genome is considered to be finite (closed pan-genome) when $\alpha > 1$ and infinite (open pan-genome) when $\alpha \leq 1$. We infer that the pan-genome of the *Streptococcus* genus is open, as are the pan-genomes of 28 ANI95% clusters within *Streptococcus*. However, the pan-genome of *S. sobrinus* (MG_41) is closed, with an average α value of 1.04. *S. sobrinus* also had the smallest pan-genome of all *Streptococcus* species clusters, which is consistent with a closed pan-genome. Yet *S. sobrinus* also contains an average of 198 pseudogenes per genome (10.4% of CDSs). One explanation of this small, closed pan-genome might have been extreme sample bias and a small population size (Park *et al.* 2019) because most *S. sobrinus* genomes in this study were isolated from a single country (Brazil). However, instead of extremely low genetic diversity, the diversity of *S. sobrinus* genomes was greater than that of several other *Streptococcus* species (Supplemental Table S5), and a phylogeny based on their core SNPs revealed high genetic diversity in *S. sobrinus* (Achtman and Zhou 2019). The *S. sobrinus* genomes are of relatively low quality (Supplemental Table S2A). Low quality genomes can also reduce the size of the pan-genome and increase the number of pseudogenes. Alternatively, the reduced pan-genome size and the high number of pseudogenes in *S.sobrinus* may be related to its association with dental caries (Bowen *et al.* 2018).
