## Supplemental Figure S1 for "Accurate reconstruction of bacterial pan- and core- genomes with PEPPAN"

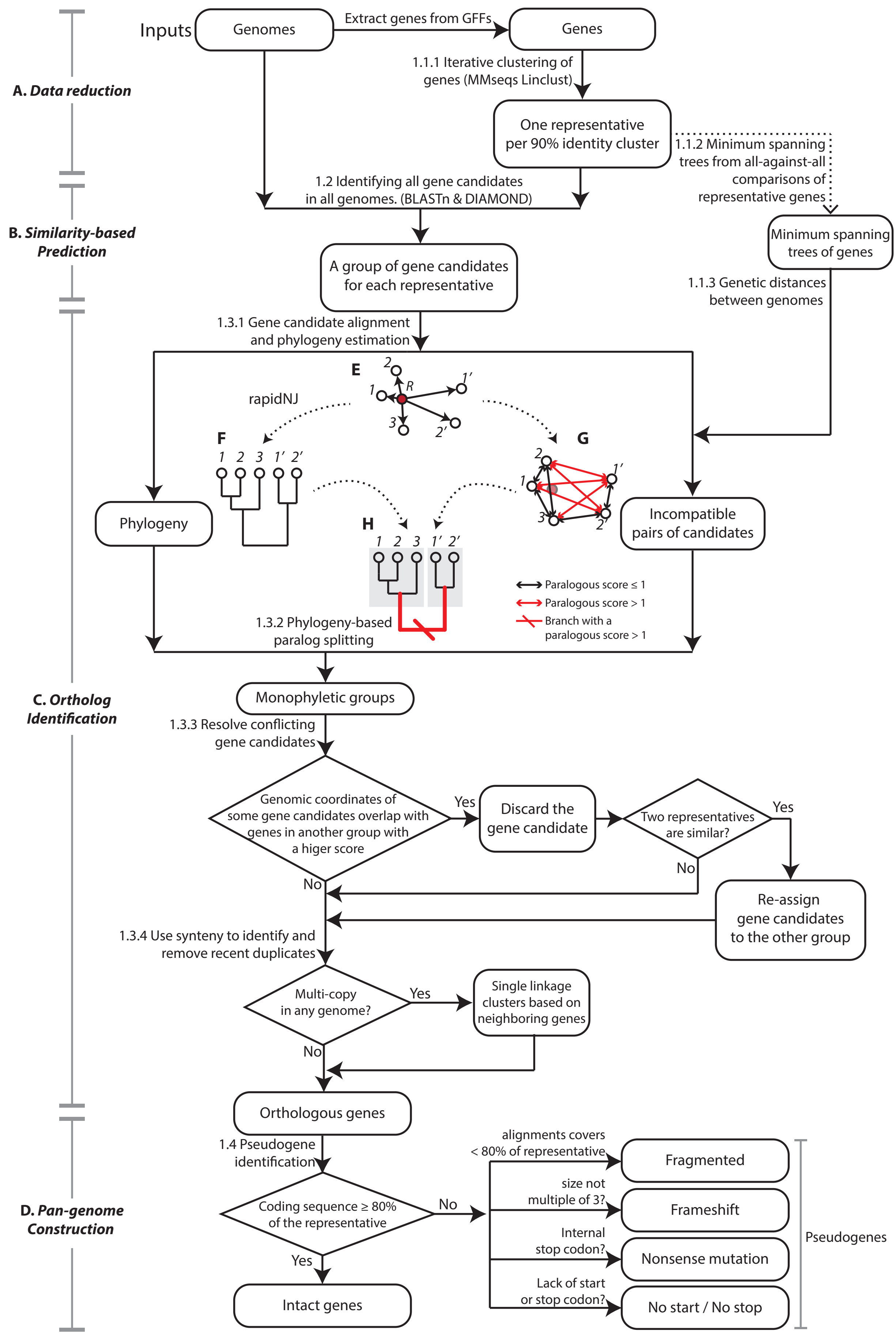


**Supplemental Figure S1. The detailed workflow of PEPPAN algorithm.** (A-D) The four successive groups of operations as shown in Figure 1A. The operations of 1.1 to 1.4 correlate with sub-headings in Supplemental Text 1. (E-H) Cartoon of an example for the phylogeny-based paralog splitting. (E) A homologous group of five gene candidates (circles) derived from alignments of a representative gene (R; red dot) on three genomes (1-3). (F) RapidNJ tree of the homologous group. (G) Compatible (black) and incompatible (red) pairs of gene candidates by their paralogous scores. (H) Split of gene tree on the branch with the greatest paralogous score.
