## Supplemental Figure S2 for "Accurate reconstruction of bacterial pan- and core- genomes with PEPPAN"

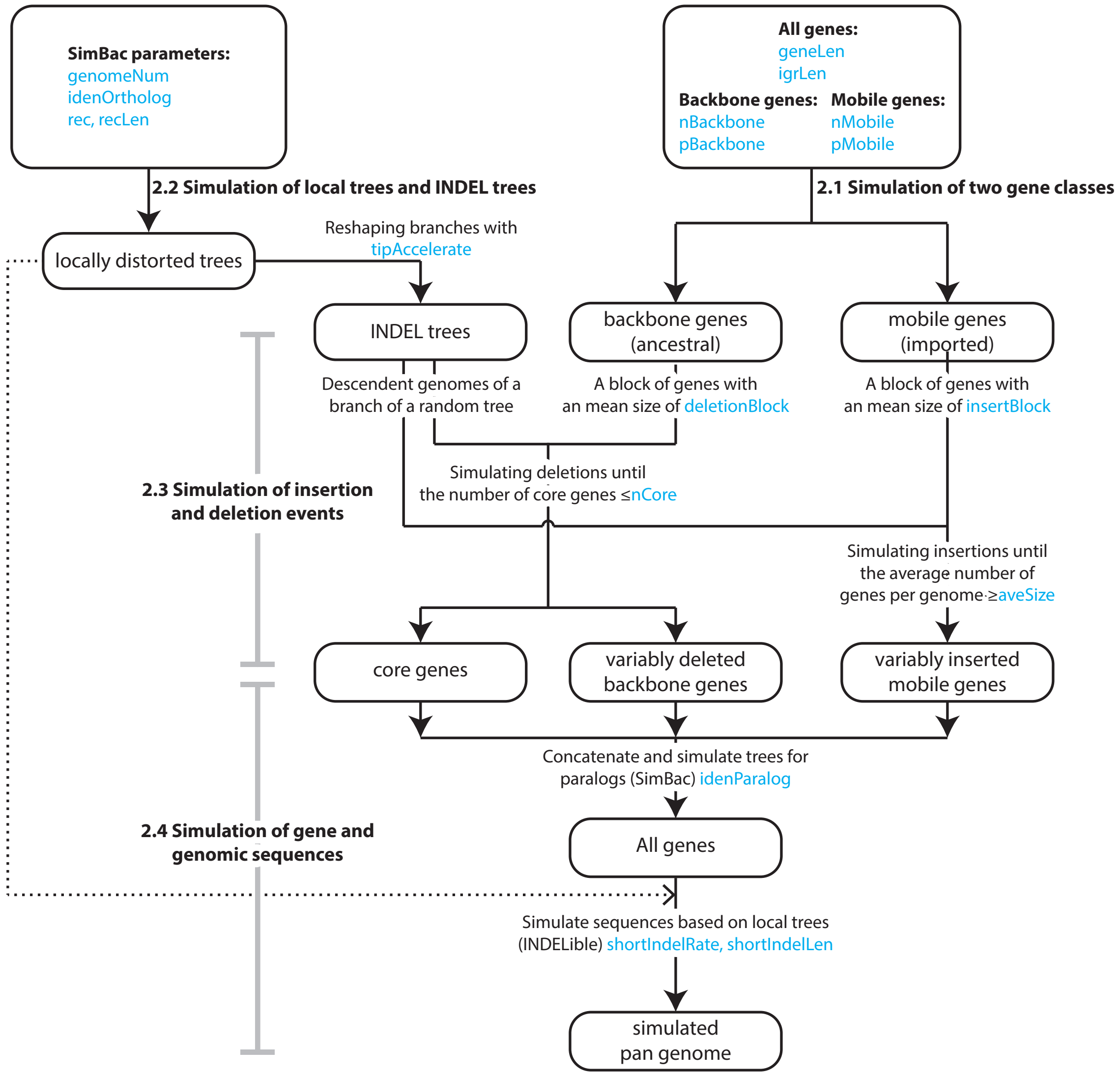


**Supplemental Figure S2. The detailed workflow of SimPan.** The operations initiated with 2.x and in bold fonts are sub-titles that corresponds to sub-titles in Supplemental Text 2. The words in blue are the names of the parameters to be specified by the users (Supplemental Table S1).
