## Supplemental Figure S3 for "Accurate reconstruction of bacterial pan- and core- genomes with PEPPAN"

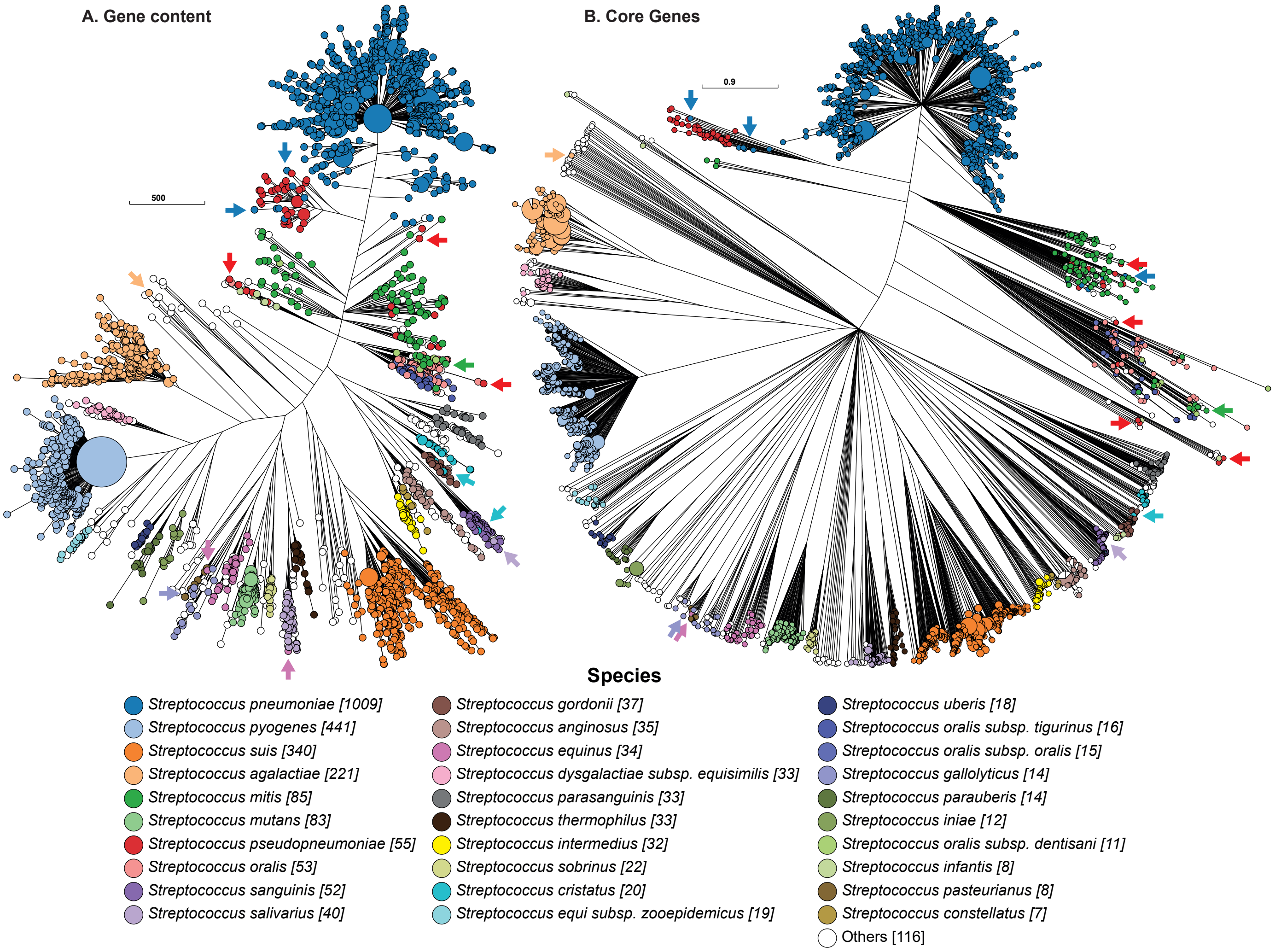


**Supplemental Figure S3. Mapping of Species label for each *Streptococcus* genome onto phylogenies generated by accessory gene contents (A) and by allelic variations in core genes (B).** The trees in A and B are the same as in Fig. 5, except that the nodes are color-coded by Species labels deposited in GenBank, as shown in the Key. The sizes of the nodes are proportional to the numbers of genomes they encompass. Some species labels that are inconsistent with both phylogenies are highlighted with colored arrows.
