## Supplemental Table S1 for "Accurate reconstruction of bacterial pan- and core- genomes with PEPPAN"

Supplemental Table S1. Available options in SimPan

| Option | Description | Default |
|---|---|---|
| genomeNum | number of genomes in population | 20 |
| idenOrtholog | average nucleotide identities of orthologous genes | 0.98 |
| rec | expected proportion of homoplastic sites between pairs of genomes | 0.05 |
| recLen | expected length (bp) of a recombinant event | 1000 |
| geneLen | mean, minimum, and maximum gene length (bp) | 900,150,6000 |
| igrLen | mean, minimum and maximum length (bp) of intergenic regions | 50,0,300 |
| nBackbone | number of backbone genes present in the common ancestor | 4000 |
| pBackbone | proportion of paralogs in the backbone genes | 0.05 |
| nMobile | number of mobile genes | 20000 |
| pMobile | proportion of paralogs in the mobile genes | 0.4 |
| tipAccelerate | gradient by which to accelerate rate of recent gene indel events | 100 |
| deletionBlock | mean, minimum and maximum number of genes in a deletion event | 3,0,30 |
| insertBlock | mean, minimum and maximum number of genes in an insertion event | 10,0,100 |
| nCore | number of strict core genes | 3500 |
| aveSize | average number of genes per genome | 4500 |
| idenParalog | average nucleotide identities of paralogous genes | 0.6 |
| shortIndelRate | average frequency of indel events relative to mutation rates | 0.01 |
| shortIndelLen | average length (bp) of short indel events within a gene | 10 |
