## Supplemental Table S5 for "Accurate reconstruction of bacterial pan- and core- genomes with PEPPAN"

Supplemental Table S5. *Streptococcus* pan-genomes from 29 ANI 95% clusters with ≥10 genomes.

| Group | Species | No. of Genomes | mean ANI% | No. of CDSs | No. of Genes | % pseudo | No. of Core genes | No. of core CDSs | No. of pan genes | α |
|---|---|---|---|---|---|---|---|---|---|---|
| **MG_4** | *S. agalactiae* | 224 | 98.86 | 1,895 | 1,994 | 5.0 | 1,037 | 433 | 6,361 | 0.80 |
| **MG_69** | *S. anginosus* | 57 | 95.94 | 1,730 | 1,832 | 5.6 | 1,184 | 979 | 5,393 | 0.79 |
| **MG_34** | *S. constellatus* | 16 | 97.96 | 1,674 | 1,785 | 6.2 | 1,348 | 1,117 | 3,418 | 0.67 |
| **MG_37** | *S. cristatus* | 12 | 95.62 | 1,800 | 1,892 | 4.9 | 1,460 | 1,364 | 3,375 | 0.65 |
| **MG_2** | *S. dysgalactiae* | 41 | 98.17 | 1,865 | 1,959 | 4.8 | 1,396 | 1,124 | 4,620 | 0.75 |
| **MG_114** | *S. equi* | 23 | 97.37 | 1,755 | 1,846 | 4.9 | 1,398 | 1,201 | 3,467 | 0.73 |
| **MG_8** | *S. equinus* | 23 | 97.72 | 1,649 | 1,718 | 4.0 | 1,286 | 975 | 3,410 | 0.62 |
| **MG_23** | *S. equinus* | 11 | 96.65 | 1,723 | 1,806 | 4.6 | 1,396 | 1,257 | 3,088 | 0.64 |
| **MG_29** | *S. gallolyticus* | 36 | 96.54 | 1,914 | 2,048 | 6.5 | 1,165 | 836 | 6,206 | 0.76 |
| **MG_32** | *S. gordonii* | 42 | 96.17 | 1,940 | 2,026 | 4.3 | 1,534 | 1,239 | 4,051 | 0.75 |
| **MG_101** | *S. iniae* | 12 | 99.63 | 1,836 | 1,928 | 4.7 | 1,551 | 1,400 | 2,854 | 0.81 |
| **MG_150** | *S. intermedius* | 30 | 98.37 | 1,726 | 1,820 | 5.2 | 1,412 | 1,151 | 3,643 | 0.81 |
| **MG_33** | *S. mitis* | 20 | 95.36 | 1,776 | 1,900 | 6.5 | 1,229 | 929 | 4,335 | 0.74 |
| **MG_43** | *S. mitis* | 14 | 95.76 | 1,696 | 1,773 | 4.3 | 1,352 | 1,139 | 3,034 | 0.80 |
| **MG_156** | *S. mutans* | 84 | 98.84 | 1,744 | 1,837 | 5.0 | 1,206 | 914 | 3,385 | 0.89 |
| **MG_18** | *S. oralis subsp. oralis* | 54 | 95.19 | 1,774 | 1,849 | 4.1 | 1,323 | 1085 | 4,805 | 0.71 |
| **MG_143** | *S. oralis subsp. tigurinus* | 10 | 95.91 | 1,752 | 1,815 | 3.5 | 1,464 | 1,393 | 2,799 | 0.71 |
| **MG_13** | *S. parasanguinis* | 33 | 95.38 | 1,864 | 1,951 | 4.5 | 1,343 | 1,105 | 4,267 | 0.82 |
| **MG_12** | *S. parasanguinis* | 10 | 96.92 | 1,863 | 1,949 | 4.4 | 1,448 | 1,274 | 3,132 | 0.78 |
| **MG_55** | *S. parauberis* | 14 | 99.08 | 1,882 | 1,980 | 4.9 | 1,520 | 1,311 | 3,749 | 0.60 |
| **MG_27** | *S. pneumoniae* | 997 | 98.50 | 1,857 | 1,992 | 6.8 | 840 | 195 | 6,457 | 0.84 |
| **MG_75** | *S. pseudopneumoniae* | 44 | 97.31 | 1,877 | 2,020 | 7.1 | 1,498 | 1,211 | 3,223 | 0.91 |
| **MG_1** | *S. pyogenes* | 441 | 98.78 | 1,590 | 1,656 | 4.0 | 1,249 | 761 | 4,246 | 0.85 |
| **MG_7** | *S. salivarius* | 53 | 95.78 | 1,880 | 1,968 | 4.5 | 1,323 | 1,083 | 4,657 | 0.81 |
| **MG_16** | *S. sanguinis* | 47 | 95.41 | 2,135 | 2,212 | 3.5 | 1,641 | 1,460 | 4,549 | 0.87 |
| **MG_41** | *S. sobrinus* | 21 | 98.82 | 1,702 | 1,900 | 10.4 | 1,529 | 614 | 2,577 | 1.04 |
| **MG_11** | *S. suis* | 321 | 95.51 | 1,952 | 2,061 | 5.3 | 989 | 648 | 9,947 | 0.82 |
| **MG_21** | *S. thermophilus* | 33 | 98.80 | 1,491 | 1,711 | 12.9 | 1,273 | 956 | 3,028 | 0.73 |
| **MG_66** | *S. uberis* | 17 | 99.07 | 1,746 | 1,804 | 3.2 | 1,463 | 1,391 | 3,132 | 0.59 |
