## Supplemental Table S6 for "Accurate reconstruction of bacterial pan- and core- genomes with PEPPAN"

Supplemental Table S6. Comparison of *Salmonella* pan-genome estimates by a precursor of PEPPAN on 537 genomes in 2015 (Zhou et al. 2020) with Roary or PEPPAN on a more representative set of 926 genomes (Alikhan et al. 2018).

| | | 2015 set (537 genomes) | 2018 set (926 genomes) | | |
|---|---|---|---|---|---|
| | Criteria (Percent present) | PEPPAN precursor (all genes) | Roary (genes ≥90 bp) | PEPPAN (genes ≥90 bp) | PEPPAN (all genes) |
| **No. pan genes** | | 26,643 | 84,041 | 52,325 | 52,337 |
| Core | 99-100% | 3295 | 1596 | 3114 | 3121 |
| Soft core | 95-100% | 3637 | 2085 | 3275 | 3282 |
| Shell | 15-95% | 1676 | 3570 | 2507 | 2509 |
| Cloud | 0-15% | 21,330 | 78,386 | 46,543 | 46,546 |

NOTE: Data on Roary analyses are from Park and Andam, 2020 (Park and Andam 2020), which also served as the source for the definitions of core, soft core, shell and cloud gene sets.
