## Supplemental Code S1 for "Accurate reconstruction of bacterial pan- and core- genomes with PEPPAN": index.rst.txt

.. PEPPA documentation master file, created by
   sphinx-quickstart on Wed Apr 22 11:18:44 2020.
   You can adapt this file completely to your liking, but it should at least
   contain the root `toctree` directive.

Welcome to PEPPA's documentation!
=================================

.. toctree::
   :maxdepth: 2
   :caption: Contents:

    installation <usage/installation.rst>
    quickstart <usage/quickstart.rst>
    parameters <usage/parameters.rst>
    inputs <usage/inputs.rst>
    outputs <usage/outputs.rst>

About PEPPA
-----------
PEPPA (Phylogeny Enhanced Pipeline for PAn-genome) is a pipeline that can construct a pan-genome from thousands of genetically diversified bacterial genomes. 
PEPPA implements a combination of tree- and synteny-based approaches to identify and exclude paralogous genes, 
as well as similarity-based gene predictions that support consistent annotations of genes and pseudogenes in individual genomes. 

Citation
--------
If you use GrapeTree please cite the pre-print in BioRxiv:

Z Zhou, M Achtman (2020) "Accurate reconstruction of the pan- and core- genomes of bacteria with PEPPA"
bioRxiv, doi: [https://doi.org/10.1101/2020.01.03.894154](https://doi.org/10.1101/2020.01.03.894154)
    

Indices and tables
==================

* :ref:`genindex`
* :ref:`modindex`
* :ref:`search`
