## Supplemental Code S1 for "Accurate reconstruction of bacterial pan- and core- genomes with PEPPAN": documentation_options.js

```
var DOCUMENTATION_OPTIONS = {
    URL_ROOT: document.getElementById("documentation_options").getAttribute('data-url_root'),
    VERSION: '1.0',
    LANGUAGE: 'None',
    COLLAPSE_INDEX: false,
    BUILDER: 'html',
    FILE_SUFFIX: '.html',
    LINK_SUFFIX: '.html',
    HAS_SOURCE: true,
    SOURCELINK_SUFFIX: '.txt',
    NAVIGATION_WITH_KEYS: false
};
```
