## Supplemental Code S1 for "Accurate reconstruction of bacterial pan- and core- genomes with PEPPAN": jquery-3.4.1.js

```
/*!
 * jQuery JavaScript Library v3.4.1
 * https://jquery.com/
 *
 * Includes Sizzle.js
 * https://sizzlejs.com/
 *
 * Copyright JS Foundation and other contributors
 * Released under the MIT license
 * https://jquery.org/license
 *
 * Date: 2019-05-01T21:04Z
 */
( function( global, factory ) {

	"use strict";

	if ( typeof module === "object" && typeof module.exports === "object" ) {

		// For CommonJS and CommonJS-like environments where a proper `window`
		// is present, execute the factory and get jQuery.
		// For environments that do not have a `window` with a `document`
		// (such as Node.js), expose a factory as module.exports.
		// This accentuates the need for the creation of a real `window`.
		// e.g. var jQuery = require("jquery")(window);
		// See ticket #14549 for more info.
		module.exports = global.document ?
			factory( global, true ) :
			function( w ) {
				if ( !w.document ) {
					throw new Error( "jQuery requires a window with a document" );
				}
				return factory( w );
			};
	} else {
		factory( global );
	}

// Pass this if window is not defined yet
} )( typeof window !== "undefined" ? window : this, function( window, noGlobal ) {

// Edge <= 12 - 13+, Firefox <=18 - 45+, IE 10 - 11, Safari 5.1 - 9+, iOS 6 - 9.1
// throw exceptions when non-strict code (e.g., ASP.NET 4.5) accesses strict mode
// arguments.callee.caller (trac-13335). But as of jQuery 3.0 (2016), strict mode should be common
// enough that all such attempts are guarded in a try block.
"use strict";

var arr = [];

var document = window.document;

var getProto = Object.getPrototypeOf;

var slice = arr.slice;

var concat = arr.concat;

var push = arr.push;

var indexOf = arr.indexOf;

var class2type = {};

var toString = class2type.toString;

var hasOwn = class2type.hasOwnProperty;

var fnToString = hasOwn.toString;

var ObjectFunctionString = fnToString.call( Object );

var support = {};

var isFunction = function isFunction( obj ) {

      // Support: Chrome <=57, Firefox <=52
      // In some browsers, typeof returns "function" for HTML <object> elements
      // (i.e., `typeof document.createElement( "object" ) === "function"`).
      // We don't want to classify *any* DOM node as a function.
      return typeof obj === "function" && typeof obj.nodeType !== "number";
  };


var isWindow = function isWindow( obj ) {
		return obj != null && obj === obj.window;
	};


	var preservedScriptAttributes = {
		type: true,
		src: true,
		nonce: true,
		noModule: true
	};

	function DOMEval( code, node, doc ) {
		doc = doc || document;

		var i, val,
			script = doc.createElement( "script" );

		script.text = code;
		if ( node ) {
			for ( i in preservedScriptAttributes ) {

				// Support: Firefox 64+, Edge 18+
				// Some browsers don't support the "nonce" property on scripts.
				// On the other hand, just using `getAttribute` is not enough as
				// the `nonce` attribute is reset to an empty string whenever it
				// becomes browsing-context connected.
				// See https://github.com/whatwg/html/issues/2369
				// See https://html.spec.whatwg.org/#nonce-attributes
				// The `node.getAttribute` check was added for the sake of
				// `jQuery.globalEval` so that it can fake a nonce-containing node
				// via an object.
				val = node[ i ] || node.getAttribute && node.getAttribute( i );
				if ( val ) {
					script.setAttribute( i, val );
				}
			}
		}
		doc.head.appendChild( script ).parentNode.removeChild( script );
	}


function toType( obj ) {
	if ( obj == null ) {
		return obj + "";
	}

	// Support: Android <=2.3 only (functionish RegExp)
	return typeof obj === "object" || typeof obj === "function" ?
		class2type[ toString.call( obj ) ] || "object" :
		typeof obj;
}
/* global Symbol */
// Defining this global in .eslintrc.json would create a danger of using the global
// unguarded in another place, it seems safer to define global only for this module


var
	version = "3.4.1",

	// Define a local copy of jQuery
	jQuery = function( selector, context ) {

		// The jQuery object is actually just the init constructor 'enhanced'
		// Need init if jQuery is called (just allow error to be thrown if not included)
		return new jQuery.fn.init( selector, context );
	},

	// Support: Android <=4.0 only
	// Make sure we trim BOM and NBSP
	rtrim = /^[\s\uFEFF\xA0]+|[\s\uFEFF\xA0]+$/g;

jQuery.fn = jQuery.prototype = {

	// The current version of jQuery being used
	jquery: version,

	constructor: jQuery,

	// The default length of a jQuery object is 0
	length: 0,

	toArray: function() {
		return slice.call( this );
	},

	// Get the Nth element in the matched element set OR
	// Get the whole matched element set as a clean array
	get: function( num ) {

		// Return all the elements in a clean array
		if ( num == null ) {
			return slice.call( this );
		}

		// Return just the one element from the set
		return num < 0 ? this[ num + this.length ] : this[ num ];
	},

	// Take an array of elements and push it onto the stack
	// (returning the new matched element set)
	pushStack: function( elems ) {

		// Build a new jQuery matched element set
		var ret = jQuery.merge( this.constructor(), elems );

		// Add the old object onto the stack (as a reference)
		ret.prevObject = this;

		// Return the newly-formed element set
		return ret;
	},

	// Execute a callback for every element in the matched set.
	each: function( callback ) {
		return jQuery.each( this, callback );
	},

	map: function( callback ) {
		return this.pushStack( jQuery.map( this, function( elem, i ) {
			return callback.call( elem, i, elem );
		} ) );
	},

	slice: function() {
		return this.pushStack( slice.apply( this, arguments ) );
	},

	first: function() {
		return this.eq( 0 );
	},

	last: function() {
		return this.eq( -1 );
	},

	eq: function( i ) {
		var len = this.length,
			j = +i + ( i < 0 ? len : 0 );
		return this.pushStack( j >= 0 && j < len ? [ this[ j ] ] : [] );
	},

	end: function() {
		return this.prevObject || this.constructor();
	},

	// For internal use only.
	// Behaves like an Array's method, not like a jQuery method.
	push: push,
	sort: arr.sort,
	splice: arr.splice
};

jQuery.extend = jQuery.fn.extend = function() {
	var options, name, src, copy, copyIsArray, clone,
		target = arguments[ 0 ] || {},
		i = 1,
		length = arguments.length,
		deep = false;

	// Handle a deep copy situation
	if ( typeof target === "boolean" ) {
		deep = target;

		// Skip the boolean and the target
		target = arguments[ i ] || {};
		i++;
	}

	// Handle case when target is a string or something (possible in deep copy)
	if ( typeof target !== "object" && !isFunction( target ) ) {
		target = {};
	}

	// Extend jQuery itself if only one argument is passed
	if ( i === length ) {
		target = this;
		i--;
	}

	for ( ; i < length; i++ ) {

		// Only deal with non-null/undefined values
		if ( ( options = arguments[ i ] ) != null ) {

			// Extend the base object
			for ( name in options ) {
				copy = options[ name ];

				// Prevent Object.prototype pollution
				// Prevent never-ending loop
				if ( name === "__proto__" || target === copy ) {
					continue;
				}

				// Recurse if we're merging plain objects or arrays
				if ( deep && copy && ( jQuery.isPlainObject( copy ) ||
					( copyIsArray = Array.isArray( copy ) ) ) ) {
					src = target[ name ];

					// Ensure proper type for the source value
					if ( copyIsArray && !Array.isArray( src ) ) {
						clone = [];
					} else if ( !copyIsArray && !jQuery.isPlainObject( src ) ) {
						clone = {};
					} else {
						clone = src;
					}
					copyIsArray = false;

					// Never move original objects, clone them
					target[ name ] = jQuery.extend( deep, clone, copy );

				// Don't bring in undefined values
				} else if ( copy !== undefined ) {
					target[ name ] = copy;
				}
			}
		}
	}

	// Return the modified object
	return target;
};

jQuery.extend( {

	// Unique for each copy of jQuery on the page
	expando: "jQuery" + ( version + Math.random() ).replace( /\D/g, "" ),

	// Assume jQuery is ready without the ready module
	isReady: true,

	error: function( msg ) {
		throw new Error( msg );
	},

	noop: function() {},

	isPlainObject: function( obj ) {
		var proto, Ctor;

		// Detect obvious negatives
		// Use toString instead of jQuery.type to catch host objects
		if ( !obj || toString.call( obj ) !== "[object Object]" ) {
			return false;
		}

		proto = getProto( obj );

		// Objects with no prototype (e.g., `Object.create( null )`) are plain
		if ( !proto ) {
			return true;
		}

		// Objects with prototype are plain iff they were constructed by a global Object function
		Ctor = hasOwn.call( proto, "constructor" ) && proto.constructor;
		return typeof Ctor === "function" && fnToString.call( Ctor ) === ObjectFunctionString;
	},

	isEmptyObject: function( obj ) {
		var name;

		for ( name in obj ) {
			return false;
		}
		return true;
	},

	// Evaluates a script in a global context
	globalEval: function( code, options ) {
		DOMEval( code, { nonce: options && options.nonce } );
	},

	each: function( obj, callback ) {
		var length, i = 0;

		if ( isArrayLike( obj ) ) {
			length = obj.length;
			for ( ; i < length; i++ ) {
				if ( callback.call( obj[ i ], i, obj[ i ] ) === false ) {
					break;
				}
			}
		} else {
			for ( i in obj ) {
				if ( callback.call( obj[ i ], i, obj[ i ] ) === false ) {
					break;
				}
			}
		}

		return obj;
	},

	// Support: Android <=4.0 only
	trim: function( text ) {
		return text == null ?
			"" :
			( text + "" ).replace( rtrim, "" );
	},

	// results is for internal usage only
	makeArray: function( arr, results ) {
		var ret = results || [];

		if ( arr != null ) {
			if ( isArrayLike( Object( arr ) ) ) {
				jQuery.merge( ret,
					typeof arr === "string" ?
					[ arr ] : arr
				);
			} else {
				push.call( ret, arr );
			}
		}

		return ret;
	},

	inArray: function( elem, arr, i ) {
		return arr == null ? -1 : indexOf.call( arr, elem, i );
	},

	// Support: Android <=4.0 only, PhantomJS 1 only
	// push.apply(_, arraylike) throws on ancient WebKit
	merge: function( first, second ) {
		var len = +second.length,
			j = 0,
			i = first.length;

		for ( ; j < len; j++ ) {
			first[ i++ ] = second[ j ];
		}

		first.length = i;

		return first;
	},

	grep: function( elems, callback, invert ) {
		var callbackInverse,
			matches = [],
			i = 0,
			length = elems.length,
			callbackExpect = !invert;

		// Go through the array, only saving the items
		// that pass the validator function
		for ( ; i < length; i++ ) {
			callbackInverse = !callback( elems[ i ], i );
			if ( callbackInverse !== callbackExpect ) {
				matches.push( elems[ i ] );
			}
		}

		return matches;
	},

	// arg is for internal usage only
	map: function( elems, callback, arg ) {
		var length, value,
			i = 0,
			ret = [];

		// Go through the array, translating each of the items to their new values
		if ( isArrayLike( elems ) ) {
			length = elems.length;
			for ( ; i < length; i++ ) {
				value = callback( elems[ i ], i, arg );

				if ( value != null ) {
					ret.push( value );
				}
			}

		// Go through every key on the object,
		} else {
			for ( i in elems ) {
				value = callback( elems[ i ], i, arg );

				if ( value != null ) {
					ret.push( value );
				}
			}
		}

		// Flatten any nested arrays
		return concat.apply( [], ret );
	},

	// A global GUID counter for objects
	guid: 1,

	// jQuery.support is not used in Core but other projects attach their
	// properties to it so it needs to exist.
	support: support
} );

if ( typeof Symbol === "function" ) {
	jQuery.fn[ Symbol.iterator ] = arr[ Symbol.iterator ];
}

// Populate the class2type map
jQuery.each( "Boolean Number String Function Array Date RegExp Object Error Symbol".split( " " ),
function( i, name ) {
	class2type[ "[object " + name + "]" ] = name.toLowerCase();
} );

function isArrayLike( obj ) {

	// Support: real iOS 8.2 only (not reproducible in simulator)
	// `in` check used to prevent JIT error (gh-2145)
	// hasOwn isn't used here due to false negatives
	// regarding Nodelist length in IE
	var length = !!obj && "length" in obj && obj.length,
		type = toType( obj );

	if ( isFunction( obj ) || isWindow( obj ) ) {
		return false;
	}

	return type === "array" || length === 0 ||
		typeof length === "number" && length > 0 && ( length - 1 ) in obj;
}
var Sizzle =
/*!
 * Sizzle CSS Selector Engine v2.3.4
 * https://sizzlejs.com/
 *
 * Copyright JS Foundation and other contributors
 * Released under the MIT license
 * https://js.foundation/
 *
 * Date: 2019-04-08
 */
(function( window ) {

var i,
	support,
	Expr,
	getText,
	isXML,
	tokenize,
	compile,
	select,
	outermostContext,
	sortInput,
	hasDuplicate,

	// Local document vars
	setDocument,
	document,
	docElem,
	documentIsHTML,
	rbuggyQSA,
	rbuggyMatches,
	matches,
	contains,

	// Instance-specific data
	expando = "sizzle" + 1 * new Date(),
	preferredDoc = window.document,
	dirruns = 0,
	done = 0,
	classCache = createCache(),
	tokenCache = createCache(),
	compilerCache = createCache(),
	nonnativeSelectorCache = createCache(),
	sortOrder = function( a, b ) {
		if ( a === b ) {
			hasDuplicate = true;
		}
		return 0;
	},

	// Instance methods
	hasOwn = ({}).hasOwnProperty,
	arr = [],
	pop = arr.pop,
	push_native = arr.push,
	push = arr.push,
	slice = arr.slice,
	// Use a stripped-down indexOf as it's faster than native
	// https://jsperf.com/thor-indexof-vs-for/5
	indexOf = function( list, elem ) {
		var i = 0,
			len = list.length;
		for ( ; i < len; i++ ) {
			if ( list[i] === elem ) {
				return i;
			}
		}
		return -1;
	},

	booleans = "checked|selected|async|autofocus|autoplay|controls|defer|disabled|hidden|ismap|loop|multiple|open|readonly|required|scoped",

	// Regular expressions

	// http://www.w3.org/TR/css3-selectors/#whitespace
	whitespace = "[\\x20\\t\\r\\n\\f]",

	// http://www.w3.org/TR/CSS21/syndata.html#value-def-identifier
	identifier = "(?:\\\\.|[\\w-]|[^\0-\\xa0])+",

	// Attribute selectors: http://www.w3.org/TR/selectors/#attribute-selectors
	attributes = "\\[" + whitespace + "*(" + identifier + ")(?:" + whitespace +
		// Operator (capture 2)
		"*([*^$|!~]?=)" + whitespace +
		// "Attribute values must be CSS identifiers [capture 5] or strings [capture 3 or capture 4]"
		"*(?:'((?:\\\\.|[^\\\\'])*)'|\"((?:\\\\.|[^\\\\\"])*)\"|(" + identifier + "))|)" + whitespace +
		"*\\]",

	pseudos = ":(" + identifier + ")(?:\\((" +
		// To reduce the number of selectors needing tokenize in the preFilter, prefer arguments:
		// 1. quoted (capture 3; capture 4 or capture 5)
		"('((?:\\\\.|[^\\\\'])*)'|\"((?:\\\\.|[^\\\\\"])*)\")|" +
		// 2. simple (capture 6)
		"((?:\\\\.|[^\\\\()[\\]]|" + attributes + ")*)|" +
		// 3. anything else (capture 2)
		".*" +
		")\\)|)",

	// Leading and non-escaped trailing whitespace, capturing some non-whitespace characters preceding the latter
	rwhitespace = new RegExp( whitespace + "+", "g" ),
	rtrim = new RegExp( "^" + whitespace + "+|((?:^|[^\\\\])(?:\\\\.)*)" + whitespace + "+$", "g" ),

	rcomma = new RegExp( "^" + whitespace + "*," + whitespace + "*" ),
	rcombinators = new RegExp( "^" + whitespace + "*([>+~]|" + whitespace + ")" + whitespace + "*" ),
	rdescend = new RegExp( whitespace + "|>" ),

	rpseudo = new RegExp( pseudos ),
	ridentifier = new RegExp( "^" + identifier + "$" ),

	matchExpr = {
		"ID": new RegExp( "^#(" + identifier + ")" ),
		"CLASS": new RegExp( "^\\.(" + identifier + ")" ),
		"TAG": new RegExp( "^(" + identifier + "|[*])" ),
		"ATTR": new RegExp( "^" + attributes ),
		"PSEUDO": new RegExp( "^" + pseudos ),
		"CHILD": new RegExp( "^:(only|first|last|nth|nth-last)-(child|of-type)(?:\\(" + whitespace +
			"*(even|odd|(([+-]|)(\\d*)n|)" + whitespace + "*(?:([+-]|)" + whitespace +
			"*(\\d+)|))" + whitespace + "*\\)|)", "i" ),
		"bool": new RegExp( "^(?:" + booleans + ")$", "i" ),
		// For use in libraries implementing .is()
		// We use this for POS matching in `select`
		"needsContext": new RegExp( "^" + whitespace + "*[>+~]|:(even|odd|eq|gt|lt|nth|first|last)(?:\\(" +
			whitespace + "*((?:-\\d)?\\d*)" + whitespace + "*\\)|)(?=[^-]|$)", "i" )
	},

	rhtml = /HTML$/i,
	rinputs = /^(?:input|select|textarea|button)$/i,
	rheader = /^h\d$/i,

	rnative = /^[^{]+\{\s*\[native \w/,

	// Easily-parseable/retrievable ID or TAG or CLASS selectors
	rquickExpr = /^(?:#([\w-]+)|(\w+)|\.([\w-]+))$/,

	rsibling = /[+~]/,

	// CSS escapes
	// http://www.w3.org/TR/CSS21/syndata.html#escaped-characters
	runescape = new RegExp( "\\\\([\\da-f]{1,6}" + whitespace + "?|(" + whitespace + ")|.)", "ig" ),
	funescape = function( _, escaped, escapedWhitespace ) {
		var high = "0x" + escaped - 0x10000;
		// NaN means non-codepoint
		// Support: Firefox<24
		// Workaround erroneous numeric interpretation of +"0x"
		return high !== high || escapedWhitespace ?
			escaped :
			high < 0 ?
				// BMP codepoint
				String.fromCharCode( high + 0x10000 ) :
				// Supplemental Plane codepoint (surrogate pair)
				String.fromCharCode( high >> 10 | 0xD800, high & 0x3FF | 0xDC00 );
	},

	// CSS string/identifier serialization
	// https://drafts.csswg.org/cssom/#common-serializing-idioms
	rcssescape = /([\0-\x1f\x7f]|^-?\d)|^-$|[^\0-\x1f\x7f-\uFFFF\w-]/g,
	fcssescape = function( ch, asCodePoint ) {
		if ( asCodePoint ) {

			// U+0000 NULL becomes U+FFFD REPLACEMENT CHARACTER
			if ( ch === "\0" ) {
				return "\uFFFD";
			}

			// Control characters and (dependent upon position) numbers get escaped as code points
			return ch.slice( 0, -1 ) + "\\" + ch.charCodeAt( ch.length - 1 ).toString( 16 ) + " ";
		}

		// Other potentially-special ASCII characters get backslash-escaped
		return "\\" + ch;
	},

	// Used for iframes
	// See setDocument()
	// Removing the function wrapper causes a "Permission Denied"
	// error in IE
	unloadHandler = function() {
		setDocument();
	},

	inDisabledFieldset = addCombinator(
		function( elem ) {
			return elem.disabled === true && elem.nodeName.toLowerCase() === "fieldset";
		},
		{ dir: "parentNode", next: "legend" }
	);

// Optimize for push.apply( _, NodeList )
try {
	push.apply(
		(arr = slice.call( preferredDoc.childNodes )),
		preferredDoc.childNodes
	);
	// Support: Android<4.0
	// Detect silently failing push.apply
	arr[ preferredDoc.childNodes.length ].nodeType;
} catch ( e ) {
	push = { apply: arr.length ?

		// Leverage slice if possible
		function( target, els ) {
			push_native.apply( target, slice.call(els) );
		} :

		// Support: IE<9
		// Otherwise append directly
		function( target, els ) {
			var j = target.length,
				i = 0;
			// Can't trust NodeList.length
			while ( (target[j++] = els[i++]) ) {}
			target.length = j - 1;
		}
	};
}

function Sizzle( selector, context, results, seed ) {
	var m, i, elem, nid, match, groups, newSelector,
		newContext = context && context.ownerDocument,

		// nodeType defaults to 9, since context defaults to document
		nodeType = context ? context.nodeType : 9;

	results = results || [];

	// Return early from calls with invalid selector or context
	if ( typeof selector !== "string" || !selector ||
		nodeType !== 1 && nodeType !== 9 && nodeType !== 11 ) {

		return results;
	}

	// Try to shortcut find operations (as opposed to filters) in HTML documents
	if ( !seed ) {

		if ( ( context ? context.ownerDocument || context : preferredDoc ) !== document ) {
			setDocument( context );
		}
		context = context || document;

		if ( documentIsHTML ) {

			// If the selector is sufficiently simple, try using a "get*By*" DOM method
			// (excepting DocumentFragment context, where the methods don't exist)
			if ( nodeType !== 11 && (match = rquickExpr.exec( selector )) ) {

				// ID selector
				if ( (m = match[1]) ) {

					// Document context
					if ( nodeType === 9 ) {
						if ( (elem = context.getElementById( m )) ) {

							// Support: IE, Opera, Webkit
							// TODO: identify versions
							// getElementById can match elements by name instead of ID
							if ( elem.id === m ) {
								results.push( elem );
								return results;
							}
						} else {
							return results;
						}

					// Element context
					} else {

						// Support: IE, Opera, Webkit
						// TODO: identify versions
						// getElementById can match elements by name instead of ID
						if ( newContext && (elem = newContext.getElementById( m )) &&
							contains( context, elem ) &&
							elem.id === m ) {

							results.push( elem );
							return results;
						}
					}

				// Type selector
				} else if ( match[2] ) {
					push.apply( results, context.getElementsByTagName( selector ) );
					return results;

				// Class selector
				} else if ( (m = match[3]) && support.getElementsByClassName &&
					context.getElementsByClassName ) {

					push.apply( results, context.getElementsByClassName( m ) );
					return results;
				}
			}

			// Take advantage of querySelectorAll
			if ( support.qsa &&
				!nonnativeSelectorCache[ selector + " " ] &&
				(!rbuggyQSA || !rbuggyQSA.test( selector )) &&

				// Support: IE 8 only
				// Exclude object elements
				(nodeType !== 1 || context.nodeName.toLowerCase() !== "object") ) {

				newSelector = selector;
				newContext = context;

				// qSA considers elements outside a scoping root when evaluating child or
				// descendant combinators, which is not what we want.
				// In such cases, we work around the behavior by prefixing every selector in the
				// list with an ID selector referencing the scope context.
				// Thanks to Andrew Dupont for this technique.
				if ( nodeType === 1 && rdescend.test( selector ) ) {

					// Capture the context ID, setting it first if necessary
					if ( (nid = context.getAttribute( "id" )) ) {
						nid = nid.replace( rcssescape, fcssescape );
					} else {
						context.setAttribute( "id", (nid = expando) );
					}

					// Prefix every selector in the list
					groups = tokenize( selector );
					i = groups.length;
					while ( i-- ) {
						groups[i] = "#" + nid + " " + toSelector( groups[i] );
					}
					newSelector = groups.join( "," );

					// Expand context for sibling selectors
					newContext = rsibling.test( selector ) && testContext( context.parentNode ) ||
						context;
				}

				try {
					push.apply( results,
						newContext.querySelectorAll( newSelector )
					);
					return results;
				} catch ( qsaError ) {
					nonnativeSelectorCache( selector, true );
				} finally {
					if ( nid === expando ) {
						context.removeAttribute( "id" );
					}
				}
			}
		}
	}

	// All others
	return select( selector.replace( rtrim, "$1" ), context, results, seed );
}

/**
 * Create key-value caches of limited size
 * @returns {function(string, object)} Returns the Object data after storing it on itself with
 *	property name the (space-suffixed) string and (if the cache is larger than Expr.cacheLength)
 *	deleting the oldest entry
 */
function createCache() {
	var keys = [];

	function cache( key, value ) {
		// Use (key + " ") to avoid collision with native prototype properties (see Issue #157)
		if ( keys.push( key + " " ) > Expr.cacheLength ) {
			// Only keep the most recent entries
			delete cache[ keys.shift() ];
		}
		return (cache[ key + " " ] = value);
	}
	return cache;
}

/**
 * Mark a function for special use by Sizzle
 * @param {Function} fn The function to mark
 */
function markFunction( fn ) {
	fn[ expando ] = true;
	return fn;
}

/**
 * Support testing using an element
 * @param {Function} fn Passed the created element and returns a boolean result
 */
function assert( fn ) {
	var el = document.createElement("fieldset");

	try {
		return !!fn( el );
	} catch (e) {
		return false;
	} finally {
		// Remove from its parent by default
		if ( el.parentNode ) {
			el.parentNode.removeChild( el );
		}
		// release memory in IE
		el = null;
	}
}

/**
 * Adds the same handler for all of the specified attrs
 * @param {String} attrs Pipe-separated list of attributes
 * @param {Function} handler The method that will be applied
 */
function addHandle( attrs, handler ) {
	var arr = attrs.split("|"),
		i = arr.length;

	while ( i-- ) {
		Expr.attrHandle[ arr[i] ] = handler;
	}
}

/**
 * Checks document order of two siblings
 * @param {Element} a
 * @param {Element} b
 * @returns {Number} Returns less than 0 if a precedes b, greater than 0 if a follows b
 */
function siblingCheck( a, b ) {
	var cur = b && a,
		diff = cur && a.nodeType === 1 && b.nodeType === 1 &&
			a.sourceIndex - b.sourceIndex;

	// Use IE sourceIndex if available on both nodes
	if ( diff ) {
		return diff;
	}

	// Check if b follows a
	if ( cur ) {
		while ( (cur = cur.nextSibling) ) {
			if ( cur === b ) {
				return -1;
			}
		}
	}

	return a ? 1 : -1;
}

/**
 * Returns a function to use in pseudos for input types
 * @param {String} type
 */
function createInputPseudo( type ) {
	return function( elem ) {
		var name = elem.nodeName.toLowerCase();
		return name === "input" && elem.type === type;
	};
}

/**
 * Returns a function to use in pseudos for buttons
 * @param {String} type
 */
function createButtonPseudo( type ) {
	return function( elem ) {
		var name = elem.nodeName.toLowerCase();
		return (name === "input" || name === "button") && elem.type === type;
	};
}

/**
 * Returns a function to use in pseudos for :enabled/:disabled
 * @param {Boolean} disabled true for :disabled; false for :enabled
 */
function createDisabledPseudo( disabled ) {

	// Known :disabled false positives: fieldset[disabled] > legend:nth-of-type(n+2) :can-disable
	return function( elem ) {

		// Only certain elements can match :enabled or :disabled
		// https://html.spec.whatwg.org/multipage/scripting.html#selector-enabled
		// https://html.spec.whatwg.org/multipage/scripting.html#selector-disabled
		if ( "form" in elem ) {

			// Check for inherited disabledness on relevant non-disabled elements:
			// * listed form-associated elements in a disabled fieldset
			//   https://html.spec.whatwg.org/multipage/forms.html#category-listed
			//   https://html.spec.whatwg.org/multipage/forms.html#concept-fe-disabled
			// * option elements in a disabled optgroup
			//   https://html.spec.whatwg.org/multipage/forms.html#concept-option-disabled
			// All such elements have a "form" property.
			if ( elem.parentNode && elem.disabled === false ) {

				// Option elements defer to a parent optgroup if present
				if ( "label" in elem ) {
					if ( "label" in elem.parentNode ) {
						return elem.parentNode.disabled === disabled;
					} else {
						return elem.disabled === disabled;
					}
				}

				// Support: IE 6 - 11
				// Use the isDisabled shortcut property to check for disabled fieldset ancestors
				return elem.isDisabled === disabled ||

					// Where there is no isDisabled, check manually
					/* jshint -W018 */
					elem.isDisabled !== !disabled &&
						inDisabledFieldset( elem ) === disabled;
			}

			return elem.disabled === disabled;

		// Try to winnow out elements that can't be disabled before trusting the disabled property.
		// Some victims get caught in our net (label, legend, menu, track), but it shouldn't
		// even exist on them, let alone have a boolean value.
		} else if ( "label" in elem ) {
			return elem.disabled === disabled;
		}

		// Remaining elements are neither :enabled nor :disabled
		return false;
	};
}

/**
 * Returns a function to use in pseudos for positionals
 * @param {Function} fn
 */
function createPositionalPseudo( fn ) {
	return markFunction(function( argument ) {
		argument = +argument;
		return markFunction(function( seed, matches ) {
			var j,
				matchIndexes = fn( [], seed.length, argument ),
				i = matchIndexes.length;

			// Match elements found at the specified indexes
			while ( i-- ) {
				if ( seed[ (j = matchIndexes[i]) ] ) {
					seed[j] = !(matches[j] = seed[j]);
				}
			}
		});
	});
}

/**
 * Checks a node for validity as a Sizzle context
 * @param {Element|Object=} context
 * @returns {Element|Object|Boolean} The input node if acceptable, otherwise a falsy value
 */
function testContext( context ) {
	return context && typeof context.getElementsByTagName !== "undefined" && context;
}

// Expose support vars for convenience
support = Sizzle.support = {};

/**
 * Detects XML nodes
 * @param {Element|Object} elem An element or a document
 * @returns {Boolean} True iff elem is a non-HTML XML node
 */
isXML = Sizzle.isXML = function( elem ) {
	var namespace = elem.namespaceURI,
		docElem = (elem.ownerDocument || elem).documentElement;

	// Support: IE <=8
	// Assume HTML when documentElement doesn't yet exist, such as inside loading iframes
	// https://bugs.jquery.com/ticket/4833
	return !rhtml.test( namespace || docElem && docElem.nodeName || "HTML" );
};

/**
 * Sets document-related variables once based on the current document
 * @param {Element|Object} [doc] An element or document object to use to set the document
 * @returns {Object} Returns the current document
 */
setDocument = Sizzle.setDocument = function( node ) {
	var hasCompare, subWindow,
		doc = node ? node.ownerDocument || node : preferredDoc;

	// Return early if doc is invalid or already selected
	if ( doc === document || doc.nodeType !== 9 || !doc.documentElement ) {
		return document;
	}

	// Update global variables
	document = doc;
	docElem = document.documentElement;
	documentIsHTML = !isXML( document );

	// Support: IE 9-11, Edge
	// Accessing iframe documents after unload throws "permission denied" errors (jQuery #13936)
	if ( preferredDoc !== document &&
		(subWindow = document.defaultView) && subWindow.top !== subWindow ) {

		// Support: IE 11, Edge
		if ( subWindow.addEventListener ) {
			subWindow.addEventListener( "unload", unloadHandler, false );

		// Support: IE 9 - 10 only
		} else if ( subWindow.attachEvent ) {
			subWindow.attachEvent( "onunload", unloadHandler );
		}
	}

	/* Attributes
	---------------------------------------------------------------------- */

	// Support: IE<8
	// Verify that getAttribute really returns attributes and not properties
	// (excepting IE8 booleans)
	support.attributes = assert(function( el ) {
		el.className = "i";
		return !el.getAttribute("className");
	});

	/* getElement(s)By*
	---------------------------------------------------------------------- */

	// Check if getElementsByTagName("*") returns only elements
	support.getElementsByTagName = assert(function( el ) {
		el.appendChild( document.createComment("") );
		return !el.getElementsByTagName("*").length;
	});

	// Support: IE<9
	support.getElementsByClassName = rnative.test( document.getElementsByClassName );

	// Support: IE<10
	// Check if getElementById returns elements by name
	// The broken getElementById methods don't pick up programmatically-set names,
	// so use a roundabout getElementsByName test
	support.getById = assert(function( el ) {
		docElem.appendChild( el ).id = expando;
		return !document.getElementsByName || !document.getElementsByName( expando ).length;
	});

	// ID filter and find
	if ( support.getById ) {
		Expr.filter["ID"] = function( id ) {
			var attrId = id.replace( runescape, funescape );
			return function( elem ) {
				return elem.getAttribute("id") === attrId;
			};
		};
		Expr.find["ID"] = function( id, context ) {
			if ( typeof context.getElementById !== "undefined" && documentIsHTML ) {
				var elem = context.getElementById( id );
				return elem ? [ elem ] : [];
			}
		};
	} else {
		Expr.filter["ID"] =  function( id ) {
			var attrId = id.replace( runescape, funescape );
			return function( elem ) {
				var node = typeof elem.getAttributeNode !== "undefined" &&
					elem.getAttributeNode("id");
				return node && node.value === attrId;
			};
		};

		// Support: IE 6 - 7 only
		// getElementById is not reliable as a find shortcut
		Expr.find["ID"] = function( id, context ) {
			if ( typeof context.getElementById !== "undefined" && documentIsHTML ) {
				var node, i, elems,
					elem = context.getElementById( id );

				if ( elem ) {

					// Verify the id attribute
					node = elem.getAttributeNode("id");
					if ( node && node.value === id ) {
						return [ elem ];
					}

					// Fall back on getElementsByName
					elems = context.getElementsByName( id );
					i = 0;
					while ( (elem = elems[i++]) ) {
						node = elem.getAttributeNode("id");
						if ( node && node.value === id ) {
							return [ elem ];
						}
					}
				}

				return [];
			}
		};
	}

	// Tag
	Expr.find["TAG"] = support.getElementsByTagName ?
		function( tag, context ) {
			if ( typeof context.getElementsByTagName !== "undefined" ) {
				return context.getElementsByTagName( tag );

			// DocumentFragment nodes don't have gEBTN
			} else if ( support.qsa ) {
				return context.querySelectorAll( tag );
			}
		} :

		function( tag, context ) {
			var elem,
				tmp = [],
				i = 0,
				// By happy coincidence, a (broken) gEBTN appears on DocumentFragment nodes too
				results = context.getElementsByTagName( tag );

			// Filter out possible comments
			if ( tag === "*" ) {
				while ( (elem = results[i++]) ) {
					if ( elem.nodeType === 1 ) {
						tmp.push( elem );
					}
				}

				return tmp;
			}
			return results;
		};

	// Class
	Expr.find["CLASS"] = support.getElementsByClassName && function( className, context ) {
		if ( typeof context.getElementsByClassName !== "undefined" && documentIsHTML ) {
			return context.getElementsByClassName( className );
		}
	};

	/* QSA/matchesSelector
	---------------------------------------------------------------------- */

	// QSA and matchesSelector support

	// matchesSelector(:active) reports false when true (IE9/Opera 11.5)
	rbuggyMatches = [];

	// qSa(:focus) reports false when true (Chrome 21)
	// We allow this because of a bug in IE8/9 that throws an error
	// whenever `document.activeElement` is accessed on an iframe
	// So, we allow :focus to pass through QSA all the time to avoid the IE error
	// See https://bugs.jquery.com/ticket/13378
	rbuggyQSA = [];

	if ( (support.qsa = rnative.test( document.querySelectorAll )) ) {
		// Build QSA regex
		// Regex strategy adopted from Diego Perini
		assert(function( el ) {
			// Select is set to empty string on purpose
			// This is to test IE's treatment of not explicitly
			// setting a boolean content attribute,
			// since its presence should be enough
			// https://bugs.jquery.com/ticket/12359
			docElem.appendChild( el ).innerHTML = "<a id='" + expando + "'></a>" +
				"<select id='" + expando + "-\r\\' msallowcapture=''>" +
				"<option selected=''></option></select>";

			// Support: IE8, Opera 11-12.16
			// Nothing should be selected when empty strings follow ^= or $= or *=
			// The test attribute must be unknown in Opera but "safe" for WinRT
			// https://msdn.microsoft.com/en-us/library/ie/hh465388.aspx#attribute_section
			if ( el.querySelectorAll("[msallowcapture^='']").length ) {
				rbuggyQSA.push( "[*^$]=" + whitespace + "*(?:''|\"\")" );
			}

			// Support: IE8
			// Boolean attributes and "value" are not treated correctly
			if ( !el.querySelectorAll("[selected]").length ) {
				rbuggyQSA.push( "\\[" + whitespace + "*(?:value|" + booleans + ")" );
			}

			// Support: Chrome<29, Android<4.4, Safari<7.0+, iOS<7.0+, PhantomJS<1.9.8+
			if ( !el.querySelectorAll( "[id~=" + expando + "-]" ).length ) {
				rbuggyQSA.push("~=");
			}

			// Webkit/Opera - :checked should return selected option elements
			// http://www.w3.org/TR/2011/REC-css3-selectors-20110929/#checked
			// IE8 throws error here and will not see later tests
			if ( !el.querySelectorAll(":checked").length ) {
				rbuggyQSA.push(":checked");
			}

			// Support: Safari 8+, iOS 8+
			// https://bugs.webkit.org/show_bug.cgi?id=136851
			// In-page `selector#id sibling-combinator selector` fails
			if ( !el.querySelectorAll( "a#" + expando + "+*" ).length ) {
				rbuggyQSA.push(".#.+[+~]");
			}
		});

		assert(function( el ) {
			el.innerHTML = "<a href='' disabled='disabled'></a>" +
				"<select disabled='disabled'><option/></select>";

			// Support: Windows 8 Native Apps
			// The type and name attributes are restricted during .innerHTML assignment
			var input = document.createElement("input");
			input.setAttribute( "type", "hidden" );
			el.appendChild( input ).setAttribute( "name", "D" );

			// Support: IE8
			// Enforce case-sensitivity of name attribute
			if ( el.querySelectorAll("[name=d]").length ) {
				rbuggyQSA.push( "name" + whitespace + "*[*^$|!~]?=" );
			}

			// FF 3.5 - :enabled/:disabled and hidden elements (hidden elements are still enabled)
			// IE8 throws error here and will not see later tests
			if ( el.querySelectorAll(":enabled").length !== 2 ) {
				rbuggyQSA.push( ":enabled", ":disabled" );
			}

			// Support: IE9-11+
			// IE's :disabled selector does not pick up the children of disabled fieldsets
			docElem.appendChild( el ).disabled = true;
			if ( el.querySelectorAll(":disabled").length !== 2 ) {
				rbuggyQSA.push( ":enabled", ":disabled" );
			}

			// Opera 10-11 does not throw on post-comma invalid pseudos
			el.querySelectorAll("*,:x");
			rbuggyQSA.push(",.*:");
		});
	}

	if ( (support.matchesSelector = rnative.test( (matches = docElem.matches ||
		docElem.webkitMatchesSelector ||
		docElem.mozMatchesSelector ||
		docElem.oMatchesSelector ||
		docElem.msMatchesSelector) )) ) {

		assert(function( el ) {
			// Check to see if it's possible to do matchesSelector
			// on a disconnected node (IE 9)
			support.disconnectedMatch = matches.call( el, "*" );

			// This should fail with an exception
			// Gecko does not error, returns false instead
			matches.call( el, "[s!='']:x" );
			rbuggyMatches.push( "!=", pseudos );
		});
	}

	rbuggyQSA = rbuggyQSA.length && new RegExp( rbuggyQSA.join("|") );
	rbuggyMatches = rbuggyMatches.length && new RegExp( rbuggyMatches.join("|") );

	/* Contains
	---------------------------------------------------------------------- */
	hasCompare = rnative.test( docElem.compareDocumentPosition );

	// Element contains another
	// Purposefully self-exclusive
	// As in, an element does not contain itself
	contains = hasCompare || rnative.test( docElem.contains ) ?
		function( a, b ) {
			var adown = a.nodeType === 9 ? a.documentElement : a,
				bup = b && b.parentNode;
			return a === bup || !!( bup && bup.nodeType === 1 && (
				adown.contains ?
					adown.contains( bup ) :
					a.compareDocumentPosition && a.compareDocumentPosition( bup ) & 16
			));
		} :
		function( a, b ) {
			if ( b ) {
				while ( (b = b.parentNode) ) {
					if ( b === a ) {
						return true;
					}
				}
			}
			return false;
		};

	/* Sorting
	---------------------------------------------------------------------- */

	// Document order sorting
	sortOrder = hasCompare ?
	function( a, b ) {

		// Flag for duplicate removal
		if ( a === b ) {
			hasDuplicate = true;
			return 0;
		}

		// Sort on method existence if only one input has compareDocumentPosition
		var compare = !a.compareDocumentPosition - !b.compareDocumentPosition;
		if ( compare ) {
			return compare;
		}

		// Calculate position if both inputs belong to the same document
		compare = ( a.ownerDocument || a ) === ( b.ownerDocument || b ) ?
			a.compareDocumentPosition( b ) :

			// Otherwise we know they are disconnected
			1;

		// Disconnected nodes
		if ( compare & 1 ||
			(!support.sortDetached && b.compareDocumentPosition( a ) === compare) ) {

			// Choose the first element that is related to our preferred document
			if ( a === document || a.ownerDocument === preferredDoc && contains(preferredDoc, a) ) {
				return -1;
			}
			if ( b === document || b.ownerDocument === preferredDoc && contains(preferredDoc, b) ) {
				return 1;
			}

			// Maintain original order
			return sortInput ?
				( indexOf( sortInput, a ) - indexOf( sortInput, b ) ) :
				0;
		}

		return compare & 4 ? -1 : 1;
	} :
	function( a, b ) {
		// Exit early if the nodes are identical
		if ( a === b ) {
			hasDuplicate = true;
			return 0;
		}

		var cur,
			i = 0,
			aup = a.parentNode,
			bup = b.parentNode,
			ap = [ a ],
			bp = [ b ];

		// Parentless nodes are either documents or disconnected
		if ( !aup || !bup ) {
			return a === document ? -1 :
				b === document ? 1 :
				aup ? -1 :
				bup ? 1 :
				sortInput ?
				( indexOf( sortInput, a ) - indexOf( sortInput, b ) ) :
				0;

		// If the nodes are siblings, we can do a quick check
		} else if ( aup === bup ) {
			return siblingCheck( a, b );
		}

		// Otherwise we need full lists of their ancestors for comparison
		cur = a;
		while ( (cur = cur.parentNode) ) {
			ap.unshift( cur );
		}
		cur = b;
		while ( (cur = cur.parentNode) ) {
			bp.unshift( cur );
		}

		// Walk down the tree looking for a discrepancy
		while ( ap[i] === bp[i] ) {
			i++;
		}

		return i ?
			// Do a sibling check if the nodes have a common ancestor
			siblingCheck( ap[i], bp[i] ) :

			// Otherwise nodes in our document sort first
			ap[i] === preferredDoc ? -1 :
			bp[i] === preferredDoc ? 1 :
			0;
	};

	return document;
};

Sizzle.matches = function( expr, elements ) {
	return Sizzle( expr, null, null, elements );
};

Sizzle.matchesSelector = function( elem, expr ) {
	// Set document vars if needed
	if ( ( elem.ownerDocument || elem ) !== document ) {
		setDocument( elem );
	}

	if ( support.matchesSelector && documentIsHTML &&
		!nonnativeSelectorCache[ expr + " " ] &&
		( !rbuggyMatches || !rbuggyMatches.test( expr ) ) &&
		( !rbuggyQSA     || !rbuggyQSA.test( expr ) ) ) {

		try {
			var ret = matches.call( elem, expr );

			// IE 9's matchesSelector returns false on disconnected nodes
			if ( ret || support.disconnectedMatch ||
					// As well, disconnected nodes are said to be in a document
					// fragment in IE 9
					elem.document && elem.document.nodeType !== 11 ) {
				return ret;
			}
		} catch (e) {
			nonnativeSelectorCache( expr, true );
		}
	}

	return Sizzle( expr, document, null, [ elem ] ).length > 0;
};

Sizzle.contains = function( context, elem ) {
	// Set document vars if needed
	if ( ( context.ownerDocument || context ) !== document ) {
		setDocument( context );
	}
	return contains( context, elem );
};

Sizzle.attr = function( elem, name ) {
	// Set document vars if needed
	if ( ( elem.ownerDocument || elem ) !== document ) {
		setDocument( elem );
	}

	var fn = Expr.attrHandle[ name.toLowerCase() ],
		// Don't get fooled by Object.prototype properties (jQuery #13807)
		val = fn && hasOwn.call( Expr.attrHandle, name.toLowerCase() ) ?
			fn( elem, name, !documentIsHTML ) :
			undefined;

	return val !== undefined ?
		val :
		support.attributes || !documentIsHTML ?
			elem.getAttribute( name ) :
			(val = elem.getAttributeNode(name)) && val.specified ?
				val.value :
				null;
};

Sizzle.escape = function( sel ) {
	return (sel + "").replace( rcssescape, fcssescape );
};

Sizzle.error = function( msg ) {
	throw new Error( "Syntax error, unrecognized expression: " + msg );
};

/**
 * Document sorting and removing duplicates
 * @param {ArrayLike} results
 */
Sizzle.uniqueSort = function( results ) {
	var elem,
		duplicates = [],
		j = 0,
		i = 0;

	// Unless we *know* we can detect duplicates, assume their presence
	hasDuplicate = !support.detectDuplicates;
	sortInput = !support.sortStable && results.slice( 0 );
	results.sort( sortOrder );

	if ( hasDuplicate ) {
		while ( (elem = results[i++]) ) {
			if ( elem === results[ i ] ) {
				j = duplicates.push( i );
			}
		}
		while ( j-- ) {
			results.splice( duplicates[ j ], 1 );
		}
	}

	// Clear input after sorting to release objects
	// See https://github.com/jquery/sizzle/pull/225
	sortInput = null;

	return results;
};

/**
 * Utility function for retrieving the text value of an array of DOM nodes
 * @param {Array|Element} elem
 */
getText = Sizzle.getText = function( elem ) {
	var node,
		ret = "",
		i = 0,
		nodeType = elem.nodeType;

	if ( !nodeType ) {
		// If no nodeType, this is expected to be an array
		while ( (node = elem[i++]) ) {
			// Do not traverse comment nodes
			ret += getText( node );
		}
	} else if ( nodeType === 1 || nodeType === 9 || nodeType === 11 ) {
		// Use textContent for elements
		// innerText usage removed for consistency of new lines (jQuery #11153)
		if ( typeof elem.textContent === "string" ) {
			return elem.textContent;
		} else {
			// Traverse its children
			for ( elem = elem.firstChild; elem; elem = elem.nextSibling ) {
				ret += getText( elem );
			}
		}
	} else if ( nodeType === 3 || nodeType === 4 ) {
		return elem.nodeValue;
	}
	// Do not include comment or processing instruction nodes

	return ret;
};

Expr = Sizzle.selectors = {

	// Can be adjusted by the user
	cacheLength: 50,

	createPseudo: markFunction,

	match: matchExpr,

	attrHandle: {},

	find: {},

	relative: {
		">": { dir: "parentNode", first: true },
		" ": { dir: "parentNode" },
		"+": { dir: "previousSibling", first: true },
		"~": { dir: "previousSibling" }
	},

	preFilter: {
		"ATTR": function( match ) {
			match[1] = match[1].replace( runescape, funescape );

			// Move the given value to match[3] whether quoted or unquoted
			match[3] = ( match[3] || match[4] || match[5] || "" ).replace( runescape, funescape );

			if ( match[2] === "~=" ) {
				match[3] = " " + match[3] + " ";
			}

			return match.slice( 0, 4 );
		},

		"CHILD": function( match ) {
			/* matches from matchExpr["CHILD"]
				1 type (only|nth|...)
				2 what (child|of-type)
				3 argument (even|odd|\d*|\d*n([+-]\d+)?|...)
				4 xn-component of xn+y argument ([+-]?\d*n|)
				5 sign of xn-component
				6 x of xn-component
				7 sign of y-component
				8 y of y-component
			*/
			match[1] = match[1].toLowerCase();

			if ( match[1].slice( 0, 3 ) === "nth" ) {
				// nth-* requires argument
				if ( !match[3] ) {
					Sizzle.error( match[0] );
				}

				// numeric x and y parameters for Expr.filter.CHILD
				// remember that false/true cast respectively to 0/1
				match[4] = +( match[4] ? match[5] + (match[6] || 1) : 2 * ( match[3] === "even" || match[3] === "odd" ) );
				match[5] = +( ( match[7] + match[8] ) || match[3] === "odd" );

			// other types prohibit arguments
			} else if ( match[3] ) {
				Sizzle.error( match[0] );
			}

			return match;
		},

		"PSEUDO": function( match ) {
			var excess,
				unquoted = !match[6] && match[2];

			if ( matchExpr["CHILD"].test( match[0] ) ) {
				return null;
			}

			// Accept quoted arguments as-is
			if ( match[3] ) {
				match[2] = match[4] || match[5] || "";

			// Strip excess characters from unquoted arguments
			} else if ( unquoted && rpseudo.test( unquoted ) &&
				// Get excess from tokenize (recursively)
				(excess = tokenize( unquoted, true )) &&
				// advance to the next closing parenthesis
				(excess = unquoted.indexOf( ")", unquoted.length - excess ) - unquoted.length) ) {

				// excess is a negative index
				match[0] = match[0].slice( 0, excess );
				match[2] = unquoted.slice( 0, excess );
			}

			// Return only captures needed by the pseudo filter method (type and argument)
			return match.slice( 0, 3 );
		}
	},

	filter: {

		"TAG": function( nodeNameSelector ) {
			var nodeName = nodeNameSelector.replace( runescape, funescape ).toLowerCase();
			return nodeNameSelector === "*" ?
				function() { return true; } :
				function( elem ) {
					return elem.nodeName && elem.nodeName.toLowerCase() === nodeName;
				};
		},

		"CLASS": function( className ) {
			var pattern = classCache[ className + " " ];

			return pattern ||
				(pattern = new RegExp( "(^|" + whitespace + ")" + className + "(" + whitespace + "|$)" )) &&
				classCache( className, function( elem ) {
					return pattern.test( typeof elem.className === "string" && elem.className || typeof elem.getAttribute !== "undefined" && elem.getAttribute("class") || "" );
				});
		},

		"ATTR": function( name, operator, check ) {
			return function( elem ) {
				var result = Sizzle.attr( elem, name );

				if ( result == null ) {
					return operator === "!=";
				}
				if ( !operator ) {
					return true;
				}

				result += "";

				return operator === "=" ? result === check :
					operator === "!=" ? result !== check :
					operator === "^=" ? check && result.indexOf( check ) === 0 :
					operator === "*=" ? check && result.indexOf( check ) > -1 :
					operator === "$=" ? check && result.slice( -check.length ) === check :
					operator === "~=" ? ( " " + result.replace( rwhitespace, " " ) + " " ).indexOf( check ) > -1 :
					operator === "|=" ? result === check || result.slice( 0, check.length + 1 ) === check + "-" :
					false;
			};
		},

		"CHILD": function( type, what, argument, first, last ) {
			var simple = type.slice( 0, 3 ) !== "nth",
				forward = type.slice( -4 ) !== "last",
				ofType = what === "of-type";

			return first === 1 && last === 0 ?

				// Shortcut for :nth-*(n)
				function( elem ) {
					return !!elem.parentNode;
				} :

				function( elem, context, xml ) {
					var cache, uniqueCache, outerCache, node, nodeIndex, start,
						dir = simple !== forward ? "nextSibling" : "previousSibling",
						parent = elem.parentNode,
						name = ofType && elem.nodeName.toLowerCase(),
						useCache = !xml && !ofType,
						diff = false;

					if ( parent ) {

						// :(first|last|only)-(child|of-type)
						if ( simple ) {
							while ( dir ) {
								node = elem;
								while ( (node = node[ dir ]) ) {
									if ( ofType ?
										node.nodeName.toLowerCase() === name :
										node.nodeType === 1 ) {

										return false;
									}
								}
								// Reverse direction for :only-* (if we haven't yet done so)
								start = dir = type === "only" && !start && "nextSibling";
							}
							return true;
						}

						start = [ forward ? parent.firstChild : parent.lastChild ];

						// non-xml :nth-child(...) stores cache data on `parent`
						if ( forward && useCache ) {

							// Seek `elem` from a previously-cached index

							// ...in a gzip-friendly way
							node = parent;
							outerCache = node[ expando ] || (node[ expando ] = {});

							// Support: IE <9 only
							// Defend against cloned attroperties (jQuery gh-1709)
							uniqueCache = outerCache[ node.uniqueID ] ||
								(outerCache[ node.uniqueID ] = {});

							cache = uniqueCache[ type ] || [];
							nodeIndex = cache[ 0 ] === dirruns && cache[ 1 ];
							diff = nodeIndex && cache[ 2 ];
							node = nodeIndex && parent.childNodes[ nodeIndex ];

							while ( (node = ++nodeIndex && node && node[ dir ] ||

								// Fallback to seeking `elem` from the start
								(diff = nodeIndex = 0) || start.pop()) ) {

								// When found, cache indexes on `parent` and break
								if ( node.nodeType === 1 && ++diff && node === elem ) {
									uniqueCache[ type ] = [ dirruns, nodeIndex, diff ];
									break;
								}
							}

						} else {
							// Use previously-cached element index if available
							if ( useCache ) {
								// ...in a gzip-friendly way
								node = elem;
								outerCache = node[ expando ] || (node[ expando ] = {});

								// Support: IE <9 only
								// Defend against cloned attroperties (jQuery gh-1709)
								uniqueCache = outerCache[ node.uniqueID ] ||
									(outerCache[ node.uniqueID ] = {});

								cache = uniqueCache[ type ] || [];
								nodeIndex = cache[ 0 ] === dirruns && cache[ 1 ];
								diff = nodeIndex;
							}

							// xml :nth-child(...)
							// or :nth-last-child(...) or :nth(-last)?-of-type(...)
							if ( diff === false ) {
								// Use the same loop as above to seek `elem` from the start
								while ( (node = ++nodeIndex && node && node[ dir ] ||
									(diff = nodeIndex = 0) || start.pop()) ) {

									if ( ( ofType ?
										node.nodeName.toLowerCase() === name :
										node.nodeType === 1 ) &&
										++diff ) {

										// Cache the index of each encountered element
										if ( useCache ) {
											outerCache = node[ expando ] || (node[ expando ] = {});

											// Support: IE <9 only
											// Defend against cloned attroperties (jQuery gh-1709)
											uniqueCache = outerCache[ node.uniqueID ] ||
												(outerCache[ node.uniqueID ] = {});

											uniqueCache[ type ] = [ dirruns, diff ];
										}

										if ( node === elem ) {
											break;
										}
									}
								}
							}
						}

						// Incorporate the offset, then check against cycle size
						diff -= last;
						return diff === first || ( diff % first === 0 && diff / first >= 0 );
					}
				};
		},

		"PSEUDO": function( pseudo, argument ) {
			// pseudo-class names are case-insensitive
			// http://www.w3.org/TR/selectors/#pseudo-classes
			// Prioritize by case sensitivity in case custom pseudos are added with uppercase letters
			// Remember that setFilters inherits from pseudos
			var args,
				fn = Expr.pseudos[ pseudo ] || Expr.setFilters[ pseudo.toLowerCase() ] ||
					Sizzle.error( "unsupported pseudo: " + pseudo );

			// The user may use createPseudo to indicate that
			// arguments are needed to create the filter function
			// just as Sizzle does
			if ( fn[ expando ] ) {
				return fn( argument );
			}

			// But maintain support for old signatures
			if ( fn.length > 1 ) {
				args = [ pseudo, pseudo, "", argument ];
				return Expr.setFilters.hasOwnProperty( pseudo.toLowerCase() ) ?
					markFunction(function( seed, matches ) {
						var idx,
							matched = fn( seed, argument ),
							i = matched.length;
						while ( i-- ) {
							idx = indexOf( seed, matched[i] );
							seed[ idx ] = !( matches[ idx ] = matched[i] );
						}
					}) :
					function( elem ) {
						return fn( elem, 0, args );
					};
			}

			return fn;
		}
	},

	pseudos: {
		// Potentially complex pseudos
		"not": markFunction(function( selector ) {
			// Trim the selector passed to compile
			// to avoid treating leading and trailing
			// spaces as combinators
			var input = [],
				results = [],
				matcher = compile( selector.replace( rtrim, "$1" ) );

			return matcher[ expando ] ?
				markFunction(function( seed, matches, context, xml ) {
					var elem,
						unmatched = matcher( seed, null, xml, [] ),
						i = seed.length;

					// Match elements unmatched by `matcher`
					while ( i-- ) {
						if ( (elem = unmatched[i]) ) {
							seed[i] = !(matches[i] = elem);
						}
					}
				}) :
				function( elem, context, xml ) {
					input[0] = elem;
					matcher( input, null, xml, results );
					// Don't keep the element (issue #299)
					input[0] = null;
					return !results.pop();
				};
		}),

		"has": markFunction(function( selector ) {
			return function( elem ) {
				return Sizzle( selector, elem ).length > 0;
			};
		}),

		"contains": markFunction(function( text ) {
			text = text.replace( runescape, funescape );
			return function( elem ) {
				return ( elem.textContent || getText( elem ) ).indexOf( text ) > -1;
			};
		}),

		// "Whether an element is represented by a :lang() selector
		// is based solely on the element's language value
		// being equal to the identifier C,
		// or beginning with the identifier C immediately followed by "-".
		// The matching of C against the element's language value is performed case-insensitively.
		// The identifier C does not have to be a valid language name."
		// http://www.w3.org/TR/selectors/#lang-pseudo
		"lang": markFunction( function( lang ) {
			// lang value must be a valid identifier
			if ( !ridentifier.test(lang || "") ) {
				Sizzle.error( "unsupported lang: " + lang );
			}
			lang = lang.replace( runescape, funescape ).toLowerCase();
			return function( elem ) {
				var elemLang;
				do {
					if ( (elemLang = documentIsHTML ?
						elem.lang :
						elem.getAttribute("xml:lang") || elem.getAttribute("lang")) ) {

						elemLang = elemLang.toLowerCase();
						return elemLang === lang || elemLang.indexOf( lang + "-" ) === 0;
					}
				} while ( (elem = elem.parentNode) && elem.nodeType === 1 );
				return false;
			};
		}),

		// Miscellaneous
		"target": function( elem ) {
			var hash = window.location && window.location.hash;
			return hash && hash.slice( 1 ) === elem.id;
		},

		"root": function( elem ) {
			return elem === docElem;
		},

		"focus": function( elem ) {
			return elem === document.activeElement && (!document.hasFocus || document.hasFocus()) && !!(elem.type || elem.href || ~elem.tabIndex);
		},

		// Boolean properties
		"enabled": createDisabledPseudo( false ),
		"disabled": createDisabledPseudo( true ),

		"checked": function( elem ) {
			// In CSS3, :checked should return both checked and selected elements
			// http://www.w3.org/TR/2011/REC-css3-selectors-20110929/#checked
			var nodeName = elem.nodeName.toLowerCase();
			return (nodeName === "input" && !!elem.checked) || (nodeName === "option" && !!elem.selected);
		},

		"selected": function( elem ) {
			// Accessing this property makes selected-by-default
			// options in Safari work properly
			if ( elem.parentNode ) {
				elem.parentNode.selectedIndex;
			}

			return elem.selected === true;
		},

		// Contents
		"empty": function( elem ) {
			// http://www.w3.org/TR/selectors/#empty-pseudo
			// :empty is negated by element (1) or content nodes (text: 3; cdata: 4; entity ref: 5),
			//   but not by others (comment: 8; processing instruction: 7; etc.)
			// nodeType < 6 works because attributes (2) do not appear as children
			for ( elem = elem.firstChild; elem; elem = elem.nextSibling ) {
				if ( elem.nodeType < 6 ) {
					return false;
				}
			}
			return true;
		},

		"parent": function( elem ) {
			return !Expr.pseudos["empty"]( elem );
		},

		// Element/input types
		"header": function( elem ) {
			return rheader.test( elem.nodeName );
		},

		"input": function( elem ) {
			return rinputs.test( elem.nodeName );
		},

		"button": function( elem ) {
			var name = elem.nodeName.toLowerCase();
			return name === "input" && elem.type === "button" || name === "button";
		},

		"text": function( elem ) {
			var attr;
			return elem.nodeName.toLowerCase() === "input" &&
				elem.type === "text" &&

				// Support: IE<8
				// New HTML5 attribute values (e.g., "search") appear with elem.type === "text"
				( (attr = elem.getAttribute("type")) == null || attr.toLowerCase() === "text" );
		},

		// Position-in-collection
		"first": createPositionalPseudo(function() {
			return [ 0 ];
		}),

		"last": createPositionalPseudo(function( matchIndexes, length ) {
			return [ length - 1 ];
		}),

		"eq": createPositionalPseudo(function( matchIndexes, length, argument ) {
			return [ argument < 0 ? argument + length : argument ];
		}),

		"even": createPositionalPseudo(function( matchIndexes, length ) {
			var i = 0;
			for ( ; i < length; i += 2 ) {
				matchIndexes.push( i );
			}
			return matchIndexes;
		}),

		"odd": createPositionalPseudo(function( matchIndexes, length ) {
			var i = 1;
			for ( ; i < length; i += 2 ) {
				matchIndexes.push( i );
			}
			return matchIndexes;
		}),

		"lt": createPositionalPseudo(function( matchIndexes, length, argument ) {
			var i = argument < 0 ?
				argument + length :
				argument > length ?
					length :
					argument;
			for ( ; --i >= 0; ) {
				matchIndexes.push( i );
			}
			return matchIndexes;
		}),

		"gt": createPositionalPseudo(function( matchIndexes, length, argument ) {
			var i = argument < 0 ? argument + length : argument;
			for ( ; ++i < length; ) {
				matchIndexes.push( i );
			}
			return matchIndexes;
		})
	}
};

Expr.pseudos["nth"] = Expr.pseudos["eq"];

// Add button/input type pseudos
for ( i in { radio: true, checkbox: true, file: true, password: true, image: true } ) {
	Expr.pseudos[ i ] = createInputPseudo( i );
}
for ( i in { submit: true, reset: true } ) {
	Expr.pseudos[ i ] = createButtonPseudo( i );
}

// Easy API for creating new setFilters
function setFilters() {}
setFilters.prototype = Expr.filters = Expr.pseudos;
Expr.setFilters = new setFilters();

tokenize = Sizzle.tokenize = function( selector, parseOnly ) {
	var matched, match, tokens, type,
		soFar, groups, preFilters,
		cached = tokenCache[ selector + " " ];

	if ( cached ) {
		return parseOnly ? 0 : cached.slice( 0 );
	}

	soFar = selector;
	groups = [];
	preFilters = Expr.preFilter;

	while ( soFar ) {

		// Comma and first run
		if ( !matched || (match = rcomma.exec( soFar )) ) {
			if ( match ) {
				// Don't consume trailing commas as valid
				soFar = soFar.slice( match[0].length ) || soFar;
			}
			groups.push( (tokens = []) );
		}

		matched = false;

		// Combinators
		if ( (match = rcombinators.exec( soFar )) ) {
			matched = match.shift();
			tokens.push({
				value: matched,
				// Cast descendant combinators to space
				type: match[0].replace( rtrim, " " )
			});
			soFar = soFar.slice( matched.length );
		}

		// Filters
		for ( type in Expr.filter ) {
			if ( (match = matchExpr[ type ].exec( soFar )) && (!preFilters[ type ] ||
				(match = preFilters[ type ]( match ))) ) {
				matched = match.shift();
				tokens.push({
					value: matched,
					type: type,
					matches: match
				});
				soFar = soFar.slice( matched.length );
			}
		}

		if ( !matched ) {
			break;
		}
	}

	// Return the length of the invalid excess
	// if we're just parsing
	// Otherwise, throw an error or return tokens
	return parseOnly ?
		soFar.length :
		soFar ?
			Sizzle.error( selector ) :
			// Cache the tokens
			tokenCache( selector, groups ).slice( 0 );
};

function toSelector( tokens ) {
	var i = 0,
		len = tokens.length,
		selector = "";
	for ( ; i < len; i++ ) {
		selector += tokens[i].value;
	}
	return selector;
}

function addCombinator( matcher, combinator, base ) {
	var dir = combinator.dir,
		skip = combinator.next,
		key = skip || dir,
		checkNonElements = base && key === "parentNode",
		doneName = done++;

	return combinator.first ?
		// Check against closest ancestor/preceding element
		function( elem, context, xml ) {
			while ( (elem = elem[ dir ]) ) {
				if ( elem.nodeType === 1 || checkNonElements ) {
					return matcher( elem, context, xml );
				}
			}
			return false;
		} :

		// Check against all ancestor/preceding elements
		function( elem, context, xml ) {
			var oldCache, uniqueCache, outerCache,
				newCache = [ dirruns, doneName ];

			// We can't set arbitrary data on XML nodes, so they don't benefit from combinator caching
			if ( xml ) {
				while ( (elem = elem[ dir ]) ) {
					if ( elem.nodeType === 1 || checkNonElements ) {
						if ( matcher( elem, context, xml ) ) {
							return true;
						}
					}
				}
			} else {
				while ( (elem = elem[ dir ]) ) {
					if ( elem.nodeType === 1 || checkNonElements ) {
						outerCache = elem[ expando ] || (elem[ expando ] = {});

						// Support: IE <9 only
						// Defend against cloned attroperties (jQuery gh-1709)
						uniqueCache = outerCache[ elem.uniqueID ] || (outerCache[ elem.uniqueID ] = {});

						if ( skip && skip === elem.nodeName.toLowerCase() ) {
							elem = elem[ dir ] || elem;
						} else if ( (oldCache = uniqueCache[ key ]) &&
							oldCache[ 0 ] === dirruns && oldCache[ 1 ] === doneName ) {

							// Assign to newCache so results back-propagate to previous elements
							return (newCache[ 2 ] = oldCache[ 2 ]);
						} else {
							// Reuse newcache so results back-propagate to previous elements
							uniqueCache[ key ] = newCache;

							// A match means we're done; a fail means we have to keep checking
							if ( (newCache[ 2 ] = matcher( elem, context, xml )) ) {
								return true;
							}
						}
					}
				}
			}
			return false;
		};
}

function elementMatcher( matchers ) {
	return matchers.length > 1 ?
		function( elem, context, xml ) {
			var i = matchers.length;
			while ( i-- ) {
				if ( !matchers[i]( elem, context, xml ) ) {
					return false;
				}
			}
			return true;
		} :
		matchers[0];
}

function multipleContexts( selector, contexts, results ) {
	var i = 0,
		len = contexts.length;
	for ( ; i < len; i++ ) {
		Sizzle( selector, contexts[i], results );
	}
	return results;
}

function condense( unmatched, map, filter, context, xml ) {
	var elem,
		newUnmatched = [],
		i = 0,
		len = unmatched.length,
		mapped = map != null;

	for ( ; i < len; i++ ) {
		if ( (elem = unmatched[i]) ) {
			if ( !filter || filter( elem, context, xml ) ) {
				newUnmatched.push( elem );
				if ( mapped ) {
					map.push( i );
				}
			}
		}
	}

	return newUnmatched;
}

function setMatcher( preFilter, selector, matcher, postFilter, postFinder, postSelector ) {
	if ( postFilter && !postFilter[ expando ] ) {
		postFilter = setMatcher( postFilter );
	}
	if ( postFinder && !postFinder[ expando ] ) {
		postFinder = setMatcher( postFinder, postSelector );
	}
	return markFunction(function( seed, results, context, xml ) {
		var temp, i, elem,
			preMap = [],
			postMap = [],
			preexisting = results.length,

			// Get initial elements from seed or context
			elems = seed || multipleContexts( selector || "*", context.nodeType ? [ context ] : context, [] ),

			// Prefilter to get matcher input, preserving a map for seed-results synchronization
			matcherIn = preFilter && ( seed || !selector ) ?
				condense( elems, preMap, preFilter, context, xml ) :
				elems,

			matcherOut = matcher ?
				// If we have a postFinder, or filtered seed, or non-seed postFilter or preexisting results,
				postFinder || ( seed ? preFilter : preexisting || postFilter ) ?

					// ...intermediate processing is necessary
					[] :

					// ...otherwise use results directly
					results :
				matcherIn;

		// Find primary matches
		if ( matcher ) {
			matcher( matcherIn, matcherOut, context, xml );
		}

		// Apply postFilter
		if ( postFilter ) {
			temp = condense( matcherOut, postMap );
			postFilter( temp, [], context, xml );

			// Un-match failing elements by moving them back to matcherIn
			i = temp.length;
			while ( i-- ) {
				if ( (elem = temp[i]) ) {
					matcherOut[ postMap[i] ] = !(matcherIn[ postMap[i] ] = elem);
				}
			}
		}

		if ( seed ) {
			if ( postFinder || preFilter ) {
				if ( postFinder ) {
					// Get the final matcherOut by condensing this intermediate into postFinder contexts
					temp = [];
					i = matcherOut.length;
					while ( i-- ) {
						if ( (elem = matcherOut[i]) ) {
							// Restore matcherIn since elem is not yet a final match
							temp.push( (matcherIn[i] = elem) );
						}
					}
					postFinder( null, (matcherOut = []), temp, xml );
				}

				// Move matched elements from seed to results to keep them synchronized
				i = matcherOut.length;
				while ( i-- ) {
					if ( (elem = matcherOut[i]) &&
						(temp = postFinder ? indexOf( seed, elem ) : preMap[i]) > -1 ) {

						seed[temp] = !(results[temp] = elem);
					}
				}
			}

		// Add elements to results, through postFinder if defined
		} else {
			matcherOut = condense(
				matcherOut === results ?
					matcherOut.splice( preexisting, matcherOut.length ) :
					matcherOut
			);
			if ( postFinder ) {
				postFinder( null, results, matcherOut, xml );
			} else {
				push.apply( results, matcherOut );
			}
		}
	});
}

function matcherFromTokens( tokens ) {
	var checkContext, matcher, j,
		len = tokens.length,
		leadingRelative = Expr.relative[ tokens[0].type ],
		implicitRelative = leadingRelative || Expr.relative[" "],
		i = leadingRelative ? 1 : 0,

		// The foundational matcher ensures that elements are reachable from top-level context(s)
		matchContext = addCombinator( function( elem ) {
			return elem === checkContext;
		}, implicitRelative, true ),
		matchAnyContext = addCombinator( function( elem ) {
			return indexOf( checkContext, elem ) > -1;
		}, implicitRelative, true ),
		matchers = [ function( elem, context, xml ) {
			var ret = ( !leadingRelative && ( xml || context !== outermostContext ) ) || (
				(checkContext = context).nodeType ?
					matchContext( elem, context, xml ) :
					matchAnyContext( elem, context, xml ) );
			// Avoid hanging onto element (issue #299)
			checkContext = null;
			return ret;
		} ];

	for ( ; i < len; i++ ) {
		if ( (matcher = Expr.relative[ tokens[i].type ]) ) {
			matchers = [ addCombinator(elementMatcher( matchers ), matcher) ];
		} else {
			matcher = Expr.filter[ tokens[i].type ].apply( null, tokens[i].matches );

			// Return special upon seeing a positional matcher
			if ( matcher[ expando ] ) {
				// Find the next relative operator (if any) for proper handling
				j = ++i;
				for ( ; j < len; j++ ) {
					if ( Expr.relative[ tokens[j].type ] ) {
						break;
					}
				}
				return setMatcher(
					i > 1 && elementMatcher( matchers ),
					i > 1 && toSelector(
						// If the preceding token was a descendant combinator, insert an implicit any-element `*`
						tokens.slice( 0, i - 1 ).concat({ value: tokens[ i - 2 ].type === " " ? "*" : "" })
					).replace( rtrim, "$1" ),
					matcher,
					i < j && matcherFromTokens( tokens.slice( i, j ) ),
					j < len && matcherFromTokens( (tokens = tokens.slice( j )) ),
					j < len && toSelector( tokens )
				);
			}
			matchers.push( matcher );
		}
	}

	return elementMatcher( matchers );
}

function matcherFromGroupMatchers( elementMatchers, setMatchers ) {
	var bySet = setMatchers.length > 0,
		byElement = elementMatchers.length > 0,
		superMatcher = function( seed, context, xml, results, outermost ) {
			var elem, j, matcher,
				matchedCount = 0,
				i = "0",
				unmatched = seed && [],
				setMatched = [],
				contextBackup = outermostContext,
				// We must always have either seed elements or outermost context
				elems = seed || byElement && Expr.find["TAG"]( "*", outermost ),
				// Use integer dirruns iff this is the outermost matcher
				dirrunsUnique = (dirruns += contextBackup == null ? 1 : Math.random() || 0.1),
				len = elems.length;

			if ( outermost ) {
				outermostContext = context === document || context || outermost;
			}

			// Add elements passing elementMatchers directly to results
			// Support: IE<9, Safari
			// Tolerate NodeList properties (IE: "length"; Safari: <number>) matching elements by id
			for ( ; i !== len && (elem = elems[i]) != null; i++ ) {
				if ( byElement && elem ) {
					j = 0;
					if ( !context && elem.ownerDocument !== document ) {
						setDocument( elem );
						xml = !documentIsHTML;
					}
					while ( (matcher = elementMatchers[j++]) ) {
						if ( matcher( elem, context || document, xml) ) {
							results.push( elem );
							break;
						}
					}
					if ( outermost ) {
						dirruns = dirrunsUnique;
					}
				}

				// Track unmatched elements for set filters
				if ( bySet ) {
					// They will have gone through all possible matchers
					if ( (elem = !matcher && elem) ) {
						matchedCount--;
					}

					// Lengthen the array for every element, matched or not
					if ( seed ) {
						unmatched.push( elem );
					}
				}
			}

			// `i` is now the count of elements visited above, and adding it to `matchedCount`
			// makes the latter nonnegative.
			matchedCount += i;

			// Apply set filters to unmatched elements
			// NOTE: This can be skipped if there are no unmatched elements (i.e., `matchedCount`
			// equals `i`), unless we didn't visit _any_ elements in the above loop because we have
			// no element matchers and no seed.
			// Incrementing an initially-string "0" `i` allows `i` to remain a string only in that
			// case, which will result in a "00" `matchedCount` that differs from `i` but is also
			// numerically zero.
			if ( bySet && i !== matchedCount ) {
				j = 0;
				while ( (matcher = setMatchers[j++]) ) {
					matcher( unmatched, setMatched, context, xml );
				}

				if ( seed ) {
					// Reintegrate element matches to eliminate the need for sorting
					if ( matchedCount > 0 ) {
						while ( i-- ) {
							if ( !(unmatched[i] || setMatched[i]) ) {
								setMatched[i] = pop.call( results );
							}
						}
					}

					// Discard index placeholder values to get only actual matches
					setMatched = condense( setMatched );
				}

				// Add matches to results
				push.apply( results, setMatched );

				// Seedless set matches succeeding multiple successful matchers stipulate sorting
				if ( outermost && !seed && setMatched.length > 0 &&
					( matchedCount + setMatchers.length ) > 1 ) {

					Sizzle.uniqueSort( results );
				}
			}

			// Override manipulation of globals by nested matchers
			if ( outermost ) {
				dirruns = dirrunsUnique;
				outermostContext = contextBackup;
			}

			return unmatched;
		};

	return bySet ?
		markFunction( superMatcher ) :
		superMatcher;
}

compile = Sizzle.compile = function( selector, match /* Internal Use Only */ ) {
	var i,
		setMatchers = [],
		elementMatchers = [],
		cached = compilerCache[ selector + " " ];

	if ( !cached ) {
		// Generate a function of recursive functions that can be used to check each element
		if ( !match ) {
			match = tokenize( selector );
		}
		i = match.length;
		while ( i-- ) {
			cached = matcherFromTokens( match[i] );
			if ( cached[ expando ] ) {
				setMatchers.push( cached );
			} else {
				elementMatchers.push( cached );
			}
		}

		// Cache the compiled function
		cached = compilerCache( selector, matcherFromGroupMatchers( elementMatchers, setMatchers ) );

		// Save selector and tokenization
		cached.selector = selector;
	}
	return cached;
};

/**
 * A low-level selection function that works with Sizzle's compiled
 *  selector functions
 * @param {String|Function} selector A selector or a pre-compiled
 *  selector function built with Sizzle.compile
 * @param {Element} context
 * @param {Array} [results]
 * @param {Array} [seed] A set of elements to match against
 */
select = Sizzle.select = function( selector, context, results, seed ) {
	var i, tokens, token, type, find,
		compiled = typeof selector === "function" && selector,
		match = !seed && tokenize( (selector = compiled.selector || selector) );

	results = results || [];

	// Try to minimize operations if there is only one selector in the list and no seed
	// (the latter of which guarantees us context)
	if ( match.length === 1 ) {

		// Reduce context if the leading compound selector is an ID
		tokens = match[0] = match[0].slice( 0 );
		if ( tokens.length > 2 && (token = tokens[0]).type === "ID" &&
				context.nodeType === 9 && documentIsHTML && Expr.relative[ tokens[1].type ] ) {

			context = ( Expr.find["ID"]( token.matches[0].replace(runescape, funescape), context ) || [] )[0];
			if ( !context ) {
				return results;

			// Precompiled matchers will still verify ancestry, so step up a level
			} else if ( compiled ) {
				context = context.parentNode;
			}

			selector = selector.slice( tokens.shift().value.length );
		}

		// Fetch a seed set for right-to-left matching
		i = matchExpr["needsContext"].test( selector ) ? 0 : tokens.length;
		while ( i-- ) {
			token = tokens[i];

			// Abort if we hit a combinator
			if ( Expr.relative[ (type = token.type) ] ) {
				break;
			}
			if ( (find = Expr.find[ type ]) ) {
				// Search, expanding context for leading sibling combinators
				if ( (seed = find(
					token.matches[0].replace( runescape, funescape ),
					rsibling.test( tokens[0].type ) && testContext( context.parentNode ) || context
				)) ) {

					// If seed is empty or no tokens remain, we can return early
					tokens.splice( i, 1 );
					selector = seed.length && toSelector( tokens );
					if ( !selector ) {
						push.apply( results, seed );
						return results;
					}

					break;
				}
			}
		}
	}

	// Compile and execute a filtering function if one is not provided
	// Provide `match` to avoid retokenization if we modified the selector above
	( compiled || compile( selector, match ) )(
		seed,
		context,
		!documentIsHTML,
		results,
		!context || rsibling.test( selector ) && testContext( context.parentNode ) || context
	);
	return results;
};

// One-time assignments

// Sort stability
support.sortStable = expando.split("").sort( sortOrder ).join("") === expando;

// Support: Chrome 14-35+
// Always assume duplicates if they aren't passed to the comparison function
support.detectDuplicates = !!hasDuplicate;

// Initialize against the default document
setDocument();

// Support: Webkit<537.32 - Safari 6.0.3/Chrome 25 (fixed in Chrome 27)
// Detached nodes confoundingly follow *each other*
support.sortDetached = assert(function( el ) {
	// Should return 1, but returns 4 (following)
	return el.compareDocumentPosition( document.createElement("fieldset") ) & 1;
});

// Support: IE<8
// Prevent attribute/property "interpolation"
// https://msdn.microsoft.com/en-us/library/ms536429%28VS.85%29.aspx
if ( !assert(function( el ) {
	el.innerHTML = "<a href='#'></a>";
	return el.firstChild.getAttribute("href") === "#" ;
}) ) {
	addHandle( "type|href|height|width", function( elem, name, isXML ) {
		if ( !isXML ) {
			return elem.getAttribute( name, name.toLowerCase() === "type" ? 1 : 2 );
		}
	});
}

// Support: IE<9
// Use defaultValue in place of getAttribute("value")
if ( !support.attributes || !assert(function( el ) {
	el.innerHTML = "<input/>";
	el.firstChild.setAttribute( "value", "" );
	return el.firstChild.getAttribute( "value" ) === "";
}) ) {
	addHandle( "value", function( elem, name, isXML ) {
		if ( !isXML && elem.nodeName.toLowerCase() === "input" ) {
			return elem.defaultValue;
		}
	});
}

// Support: IE<9
// Use getAttributeNode to fetch booleans when getAttribute lies
if ( !assert(function( el ) {
	return el.getAttribute("disabled") == null;
}) ) {
	addHandle( booleans, function( elem, name, isXML ) {
		var val;
		if ( !isXML ) {
			return elem[ name ] === true ? name.toLowerCase() :
					(val = elem.getAttributeNode( name )) && val.specified ?
					val.value :
				null;
		}
	});
}

return Sizzle;

})( window );


jQuery.find = Sizzle;
jQuery.expr = Sizzle.selectors;

// Deprecated
jQuery.expr[ ":" ] = jQuery.expr.pseudos;
jQuery.uniqueSort = jQuery.unique = Sizzle.uniqueSort;
jQuery.text = Sizzle.getText;
jQuery.isXMLDoc = Sizzle.isXML;
jQuery.contains = Sizzle.contains;
jQuery.escapeSelector = Sizzle.escape;


var dir = function( elem, dir, until ) {
	var matched = [],
		truncate = until !== undefined;

	while ( ( elem = elem[ dir ] ) && elem.nodeType !== 9 ) {
		if ( elem.nodeType === 1 ) {
			if ( truncate && jQuery( elem ).is( until ) ) {
				break;
			}
			matched.push( elem );
		}
	}
	return matched;
};


var siblings = function( n, elem ) {
	var matched = [];

	for ( ; n; n = n.nextSibling ) {
		if ( n.nodeType === 1 && n !== elem ) {
			matched.push( n );
		}
	}

	return matched;
};


var rneedsContext = jQuery.expr.match.needsContext;


function nodeName( elem, name ) {

  return elem.nodeName && elem.nodeName.toLowerCase() === name.toLowerCase();

};
var rsingleTag = ( /^<([a-z][^\/\0>:\x20\t\r\n\f]*)[\x20\t\r\n\f]*\/?>(?:<\/\1>|)$/i );


// Implement the identical functionality for filter and not
function winnow( elements, qualifier, not ) {
	if ( isFunction( qualifier ) ) {
		return jQuery.grep( elements, function( elem, i ) {
			return !!qualifier.call( elem, i, elem ) !== not;
		} );
	}

	// Single element
	if ( qualifier.nodeType ) {
		return jQuery.grep( elements, function( elem ) {
			return ( elem === qualifier ) !== not;
		} );
	}

	// Arraylike of elements (jQuery, arguments, Array)
	if ( typeof qualifier !== "string" ) {
		return jQuery.grep( elements, function( elem ) {
			return ( indexOf.call( qualifier, elem ) > -1 ) !== not;
		} );
	}

	// Filtered directly for both simple and complex selectors
	return jQuery.filter( qualifier, elements, not );
}

jQuery.filter = function( expr, elems, not ) {
	var elem = elems[ 0 ];

	if ( not ) {
		expr = ":not(" + expr + ")";
	}

	if ( elems.length === 1 && elem.nodeType === 1 ) {
		return jQuery.find.matchesSelector( elem, expr ) ? [ elem ] : [];
	}

	return jQuery.find.matches( expr, jQuery.grep( elems, function( elem ) {
		return elem.nodeType === 1;
	} ) );
};

jQuery.fn.extend( {
	find: function( selector ) {
		var i, ret,
			len = this.length,
			self = this;

		if ( typeof selector !== "string" ) {
			return this.pushStack( jQuery( selector ).filter( function() {
				for ( i = 0; i < len; i++ ) {
					if ( jQuery.contains( self[ i ], this ) ) {
						return true;
					}
				}
			} ) );
		}

		ret = this.pushStack( [] );

		for ( i = 0; i < len; i++ ) {
			jQuery.find( selector, self[ i ], ret );
		}

		return len > 1 ? jQuery.uniqueSort( ret ) : ret;
	},
	filter: function( selector ) {
		return this.pushStack( winnow( this, selector || [], false ) );
	},
	not: function( selector ) {
		return this.pushStack( winnow( this, selector || [], true ) );
	},
	is: function( selector ) {
		return !!winnow(
			this,

			// If this is a positional/relative selector, check membership in the returned set
			// so $("p:first").is("p:last") won't return true for a doc with two "p".
			typeof selector === "string" && rneedsContext.test( selector ) ?
				jQuery( selector ) :
				selector || [],
			false
		).length;
	}
} );


// Initialize a jQuery object


// A central reference to the root jQuery(document)
var rootjQuery,

	// A simple way to check for HTML strings
	// Prioritize #id over <tag> to avoid XSS via location.hash (#9521)
	// Strict HTML recognition (#11290: must start with <)
	// Shortcut simple #id case for speed
	rquickExpr = /^(?:\s*(<[\w\W]+>)[^>]*|#([\w-]+))$/,

	init = jQuery.fn.init = function( selector, context, root ) {
		var match, elem;

		// HANDLE: $(""), $(null), $(undefined), $(false)
		if ( !selector ) {
			return this;
		}

		// Method init() accepts an alternate rootjQuery
		// so migrate can support jQuery.sub (gh-2101)
		root = root || rootjQuery;

		// Handle HTML strings
		if ( typeof selector === "string" ) {
			if ( selector[ 0 ] === "<" &&
				selector[ selector.length - 1 ] === ">" &&
				selector.length >= 3 ) {

				// Assume that strings that start and end with <> are HTML and skip the regex check
				match = [ null, selector, null ];

			} else {
				match = rquickExpr.exec( selector );
			}

			// Match html or make sure no context is specified for #id
			if ( match && ( match[ 1 ] || !context ) ) {

				// HANDLE: $(html) -> $(array)
				if ( match[ 1 ] ) {
					context = context instanceof jQuery ? context[ 0 ] : context;

					// Option to run scripts is true for back-compat
					// Intentionally let the error be thrown if parseHTML is not present
					jQuery.merge( this, jQuery.parseHTML(
						match[ 1 ],
						context && context.nodeType ? context.ownerDocument || context : document,
						true
					) );

					// HANDLE: $(html, props)
					if ( rsingleTag.test( match[ 1 ] ) && jQuery.isPlainObject( context ) ) {
						for ( match in context ) {

							// Properties of context are called as methods if possible
							if ( isFunction( this[ match ] ) ) {
								this[ match ]( context[ match ] );

							// ...and otherwise set as attributes
							} else {
								this.attr( match, context[ match ] );
							}
						}
					}

					return this;

				// HANDLE: $(#id)
				} else {
					elem = document.getElementById( match[ 2 ] );

					if ( elem ) {

						// Inject the element directly into the jQuery object
						this[ 0 ] = elem;
						this.length = 1;
					}
					return this;
				}

			// HANDLE: $(expr, $(...))
			} else if ( !context || context.jquery ) {
				return ( context || root ).find( selector );

			// HANDLE: $(expr, context)
			// (which is just equivalent to: $(context).find(expr)
			} else {
				return this.constructor( context ).find( selector );
			}

		// HANDLE: $(DOMElement)
		} else if ( selector.nodeType ) {
			this[ 0 ] = selector;
			this.length = 1;
			return this;

		// HANDLE: $(function)
		// Shortcut for document ready
		} else if ( isFunction( selector ) ) {
			return root.ready !== undefined ?
				root.ready( selector ) :

				// Execute immediately if ready is not present
				selector( jQuery );
		}

		return jQuery.makeArray( selector, this );
	};

// Give the init function the jQuery prototype for later instantiation
init.prototype = jQuery.fn;

// Initialize central reference
rootjQuery = jQuery( document );


var rparentsprev = /^(?:parents|prev(?:Until|All))/,

	// Methods guaranteed to produce a unique set when starting from a unique set
	guaranteedUnique = {
		children: true,
		contents: true,
		next: true,
		prev: true
	};

jQuery.fn.extend( {
	has: function( target ) {
		var targets = jQuery( target, this ),
			l = targets.length;

		return this.filter( function() {
			var i = 0;
			for ( ; i < l; i++ ) {
				if ( jQuery.contains( this, targets[ i ] ) ) {
					return true;
				}
			}
		} );
	},

	closest: function( selectors, context ) {
		var cur,
			i = 0,
			l = this.length,
			matched = [],
			targets = typeof selectors !== "string" && jQuery( selectors );

		// Positional selectors never match, since there's no _selection_ context
		if ( !rneedsContext.test( selectors ) ) {
			for ( ; i < l; i++ ) {
				for ( cur = this[ i ]; cur && cur !== context; cur = cur.parentNode ) {

					// Always skip document fragments
					if ( cur.nodeType < 11 && ( targets ?
						targets.index( cur ) > -1 :

						// Don't pass non-elements to Sizzle
						cur.nodeType === 1 &&
							jQuery.find.matchesSelector( cur, selectors ) ) ) {

						matched.push( cur );
						break;
					}
				}
			}
		}

		return this.pushStack( matched.length > 1 ? jQuery.uniqueSort( matched ) : matched );
	},

	// Determine the position of an element within the set
	index: function( elem ) {

		// No argument, return index in parent
		if ( !elem ) {
			return ( this[ 0 ] && this[ 0 ].parentNode ) ? this.first().prevAll().length : -1;
		}

		// Index in selector
		if ( typeof elem === "string" ) {
			return indexOf.call( jQuery( elem ), this[ 0 ] );
		}

		// Locate the position of the desired element
		return indexOf.call( this,

			// If it receives a jQuery object, the first element is used
			elem.jquery ? elem[ 0 ] : elem
		);
	},

	add: function( selector, context ) {
		return this.pushStack(
			jQuery.uniqueSort(
				jQuery.merge( this.get(), jQuery( selector, context ) )
			)
		);
	},

	addBack: function( selector ) {
		return this.add( selector == null ?
			this.prevObject : this.prevObject.filter( selector )
		);
	}
} );

function sibling( cur, dir ) {
	while ( ( cur = cur[ dir ] ) && cur.nodeType !== 1 ) {}
	return cur;
}

jQuery.each( {
	parent: function( elem ) {
		var parent = elem.parentNode;
		return parent && parent.nodeType !== 11 ? parent : null;
	},
	parents: function( elem ) {
		return dir( elem, "parentNode" );
	},
	parentsUntil: function( elem, i, until ) {
		return dir( elem, "parentNode", until );
	},
	next: function( elem ) {
		return sibling( elem, "nextSibling" );
	},
	prev: function( elem ) {
		return sibling( elem, "previousSibling" );
	},
	nextAll: function( elem ) {
		return dir( elem, "nextSibling" );
	},
	prevAll: function( elem ) {
		return dir( elem, "previousSibling" );
	},
	nextUntil: function( elem, i, until ) {
		return dir( elem, "nextSibling", until );
	},
	prevUntil: function( elem, i, until ) {
		return dir( elem, "previousSibling", until );
	},
	siblings: function( elem ) {
		return siblings( ( elem.parentNode || {} ).firstChild, elem );
	},
	children: function( elem ) {
		return siblings( elem.firstChild );
	},
	contents: function( elem ) {
		if ( typeof elem.contentDocument !== "undefined" ) {
			return elem.contentDocument;
		}

		// Support: IE 9 - 11 only, iOS 7 only, Android Browser <=4.3 only
		// Treat the template element as a regular one in browsers that
		// don't support it.
		if ( nodeName( elem, "template" ) ) {
			elem = elem.content || elem;
		}

		return jQuery.merge( [], elem.childNodes );
	}
}, function( name, fn ) {
	jQuery.fn[ name ] = function( until, selector ) {
		var matched = jQuery.map( this, fn, until );

		if ( name.slice( -5 ) !== "Until" ) {
			selector = until;
		}

		if ( selector && typeof selector === "string" ) {
			matched = jQuery.filter( selector, matched );
		}

		if ( this.length > 1 ) {

			// Remove duplicates
			if ( !guaranteedUnique[ name ] ) {
				jQuery.uniqueSort( matched );
			}

			// Reverse order for parents* and prev-derivatives
			if ( rparentsprev.test( name ) ) {
				matched.reverse();
			}
		}

		return this.pushStack( matched );
	};
} );
var rnothtmlwhite = ( /[^\x20\t\r\n\f]+/g );


// Convert String-formatted options into Object-formatted ones
function createOptions( options ) {
	var object = {};
	jQuery.each( options.match( rnothtmlwhite ) || [], function( _, flag ) {
		object[ flag ] = true;
	} );
	return object;
}

/*
 * Create a callback list using the following parameters:
 *
 *	options: an optional list of space-separated options that will change how
 *			the callback list behaves or a more traditional option object
 *
 * By default a callback list will act like an event callback list and can be
 * "fired" multiple times.
 *
 * Possible options:
 *
 *	once:			will ensure the callback list can only be fired once (like a Deferred)
 *
 *	memory:			will keep track of previous values and will call any callback added
 *					after the list has been fired right away with the latest "memorized"
 *					values (like a Deferred)
 *
 *	unique:			will ensure a callback can only be added once (no duplicate in the list)
 *
 *	stopOnFalse:	interrupt callings when a callback returns false
 *
 */
jQuery.Callbacks = function( options ) {

	// Convert options from String-formatted to Object-formatted if needed
	// (we check in cache first)
	options = typeof options === "string" ?
		createOptions( options ) :
		jQuery.extend( {}, options );

	var // Flag to know if list is currently firing
		firing,

		// Last fire value for non-forgettable lists
		memory,

		// Flag to know if list was already fired
		fired,

		// Flag to prevent firing
		locked,

		// Actual callback list
		list = [],

		// Queue of execution data for repeatable lists
		queue = [],

		// Index of currently firing callback (modified by add/remove as needed)
		firingIndex = -1,

		// Fire callbacks
		fire = function() {

			// Enforce single-firing
			locked = locked || options.once;

			// Execute callbacks for all pending executions,
			// respecting firingIndex overrides and runtime changes
			fired = firing = true;
			for ( ; queue.length; firingIndex = -1 ) {
				memory = queue.shift();
				while ( ++firingIndex < list.length ) {

					// Run callback and check for early termination
					if ( list[ firingIndex ].apply( memory[ 0 ], memory[ 1 ] ) === false &&
						options.stopOnFalse ) {

						// Jump to end and forget the data so .add doesn't re-fire
						firingIndex = list.length;
						memory = false;
					}
				}
			}

			// Forget the data if we're done with it
			if ( !options.memory ) {
				memory = false;
			}

			firing = false;

			// Clean up if we're done firing for good
			if ( locked ) {

				// Keep an empty list if we have data for future add calls
				if ( memory ) {
					list = [];

				// Otherwise, this object is spent
				} else {
					list = "";
				}
			}
		},

		// Actual Callbacks object
		self = {

			// Add a callback or a collection of callbacks to the list
			add: function() {
				if ( list ) {

					// If we have memory from a past run, we should fire after adding
					if ( memory && !firing ) {
						firingIndex = list.length - 1;
						queue.push( memory );
					}

					( function add( args ) {
						jQuery.each( args, function( _, arg ) {
							if ( isFunction( arg ) ) {
								if ( !options.unique || !self.has( arg ) ) {
									list.push( arg );
								}
							} else if ( arg && arg.length && toType( arg ) !== "string" ) {

								// Inspect recursively
								add( arg );
							}
						} );
					} )( arguments );

					if ( memory && !firing ) {
						fire();
					}
				}
				return this;
			},

			// Remove a callback from the list
			remove: function() {
				jQuery.each( arguments, function( _, arg ) {
					var index;
					while ( ( index = jQuery.inArray( arg, list, index ) ) > -1 ) {
						list.splice( index, 1 );

						// Handle firing indexes
						if ( index <= firingIndex ) {
							firingIndex--;
						}
					}
				} );
				return this;
			},

			// Check if a given callback is in the list.
			// If no argument is given, return whether or not list has callbacks attached.
			has: function( fn ) {
				return fn ?
					jQuery.inArray( fn, list ) > -1 :
					list.length > 0;
			},

			// Remove all callbacks from the list
			empty: function() {
				if ( list ) {
					list = [];
				}
				return this;
			},

			// Disable .fire and .add
			// Abort any current/pending executions
			// Clear all callbacks and values
			disable: function() {
				locked = queue = [];
				list = memory = "";
				return this;
			},
			disabled: function() {
				return !list;
			},

			// Disable .fire
			// Also disable .add unless we have memory (since it would have no effect)
			// Abort any pending executions
			lock: function() {
				locked = queue = [];
				if ( !memory && !firing ) {
					list = memory = "";
				}
				return this;
			},
			locked: function() {
				return !!locked;
			},

			// Call all callbacks with the given context and arguments
			fireWith: function( context, args ) {
				if ( !locked ) {
					args = args || [];
					args = [ context, args.slice ? args.slice() : args ];
					queue.push( args );
					if ( !firing ) {
						fire();
					}
				}
				return this;
			},

			// Call all the callbacks with the given arguments
			fire: function() {
				self.fireWith( this, arguments );
				return this;
			},

			// To know if the callbacks have already been called at least once
			fired: function() {
				return !!fired;
			}
		};

	return self;
};


function Identity( v ) {
	return v;
}
function Thrower( ex ) {
	throw ex;
}

function adoptValue( value, resolve, reject, noValue ) {
	var method;

	try {

		// Check for promise aspect first to privilege synchronous behavior
		if ( value && isFunction( ( method = value.promise ) ) ) {
			method.call( value ).done( resolve ).fail( reject );

		// Other thenables
		} else if ( value && isFunction( ( method = value.then ) ) ) {
			method.call( value, resolve, reject );

		// Other non-thenables
		} else {

			// Control `resolve` arguments by letting Array#slice cast boolean `noValue` to integer:
			// * false: [ value ].slice( 0 ) => resolve( value )
			// * true: [ value ].slice( 1 ) => resolve()
			resolve.apply( undefined, [ value ].slice( noValue ) );
		}

	// For Promises/A+, convert exceptions into rejections
	// Since jQuery.when doesn't unwrap thenables, we can skip the extra checks appearing in
	// Deferred#then to conditionally suppress rejection.
	} catch ( value ) {

		// Support: Android 4.0 only
		// Strict mode functions invoked without .call/.apply get global-object context
		reject.apply( undefined, [ value ] );
	}
}

jQuery.extend( {

	Deferred: function( func ) {
		var tuples = [

				// action, add listener, callbacks,
				// ... .then handlers, argument index, [final state]
				[ "notify", "progress", jQuery.Callbacks( "memory" ),
					jQuery.Callbacks( "memory" ), 2 ],
				[ "resolve", "done", jQuery.Callbacks( "once memory" ),
					jQuery.Callbacks( "once memory" ), 0, "resolved" ],
				[ "reject", "fail", jQuery.Callbacks( "once memory" ),
					jQuery.Callbacks( "once memory" ), 1, "rejected" ]
			],
			state = "pending",
			promise = {
				state: function() {
					return state;
				},
				always: function() {
					deferred.done( arguments ).fail( arguments );
					return this;
				},
				"catch": function( fn ) {
					return promise.then( null, fn );
				},

				// Keep pipe for back-compat
				pipe: function( /* fnDone, fnFail, fnProgress */ ) {
					var fns = arguments;

					return jQuery.Deferred( function( newDefer ) {
						jQuery.each( tuples, function( i, tuple ) {

							// Map tuples (progress, done, fail) to arguments (done, fail, progress)
							var fn = isFunction( fns[ tuple[ 4 ] ] ) && fns[ tuple[ 4 ] ];

							// deferred.progress(function() { bind to newDefer or newDefer.notify })
							// deferred.done(function() { bind to newDefer or newDefer.resolve })
							// deferred.fail(function() { bind to newDefer or newDefer.reject })
							deferred[ tuple[ 1 ] ]( function() {
								var returned = fn && fn.apply( this, arguments );
								if ( returned && isFunction( returned.promise ) ) {
									returned.promise()
										.progress( newDefer.notify )
										.done( newDefer.resolve )
										.fail( newDefer.reject );
								} else {
									newDefer[ tuple[ 0 ] + "With" ](
										this,
										fn ? [ returned ] : arguments
									);
								}
							} );
						} );
						fns = null;
					} ).promise();
				},
				then: function( onFulfilled, onRejected, onProgress ) {
					var maxDepth = 0;
					function resolve( depth, deferred, handler, special ) {
						return function() {
							var that = this,
								args = arguments,
								mightThrow = function() {
									var returned, then;

									// Support: Promises/A+ section 2.3.3.3.3
									// https://promisesaplus.com/#point-59
									// Ignore double-resolution attempts
									if ( depth < maxDepth ) {
										return;
									}

									returned = handler.apply( that, args );

									// Support: Promises/A+ section 2.3.1
									// https://promisesaplus.com/#point-48
									if ( returned === deferred.promise() ) {
										throw new TypeError( "Thenable self-resolution" );
									}

									// Support: Promises/A+ sections 2.3.3.1, 3.5
									// https://promisesaplus.com/#point-54
									// https://promisesaplus.com/#point-75
									// Retrieve `then` only once
									then = returned &&

										// Support: Promises/A+ section 2.3.4
										// https://promisesaplus.com/#point-64
										// Only check objects and functions for thenability
										( typeof returned === "object" ||
											typeof returned === "function" ) &&
										returned.then;

									// Handle a returned thenable
									if ( isFunction( then ) ) {

										// Special processors (notify) just wait for resolution
										if ( special ) {
											then.call(
												returned,
												resolve( maxDepth, deferred, Identity, special ),
												resolve( maxDepth, deferred, Thrower, special )
											);

										// Normal processors (resolve) also hook into progress
										} else {

											// ...and disregard older resolution values
											maxDepth++;

											then.call(
												returned,
												resolve( maxDepth, deferred, Identity, special ),
												resolve( maxDepth, deferred, Thrower, special ),
												resolve( maxDepth, deferred, Identity,
													deferred.notifyWith )
											);
										}

									// Handle all other returned values
									} else {

										// Only substitute handlers pass on context
										// and multiple values (non-spec behavior)
										if ( handler !== Identity ) {
											that = undefined;
											args = [ returned ];
										}

										// Process the value(s)
										// Default process is resolve
										( special || deferred.resolveWith )( that, args );
									}
								},

								// Only normal processors (resolve) catch and reject exceptions
								process = special ?
									mightThrow :
									function() {
										try {
											mightThrow();
										} catch ( e ) {

											if ( jQuery.Deferred.exceptionHook ) {
												jQuery.Deferred.exceptionHook( e,
													process.stackTrace );
											}

											// Support: Promises/A+ section 2.3.3.3.4.1
											// https://promisesaplus.com/#point-61
											// Ignore post-resolution exceptions
											if ( depth + 1 >= maxDepth ) {

												// Only substitute handlers pass on context
												// and multiple values (non-spec behavior)
												if ( handler !== Thrower ) {
													that = undefined;
													args = [ e ];
												}

												deferred.rejectWith( that, args );
											}
										}
									};

							// Support: Promises/A+ section 2.3.3.3.1
							// https://promisesaplus.com/#point-57
							// Re-resolve promises immediately to dodge false rejection from
							// subsequent errors
							if ( depth ) {
								process();
							} else {

								// Call an optional hook to record the stack, in case of exception
								// since it's otherwise lost when execution goes async
								if ( jQuery.Deferred.getStackHook ) {
									process.stackTrace = jQuery.Deferred.getStackHook();
								}
								window.setTimeout( process );
							}
						};
					}

					return jQuery.Deferred( function( newDefer ) {

						// progress_handlers.add( ... )
						tuples[ 0 ][ 3 ].add(
							resolve(
								0,
								newDefer,
								isFunction( onProgress ) ?
									onProgress :
									Identity,
								newDefer.notifyWith
							)
						);

						// fulfilled_handlers.add( ... )
						tuples[ 1 ][ 3 ].add(
							resolve(
								0,
								newDefer,
								isFunction( onFulfilled ) ?
									onFulfilled :
									Identity
							)
						);

						// rejected_handlers.add( ... )
						tuples[ 2 ][ 3 ].add(
							resolve(
								0,
								newDefer,
								isFunction( onRejected ) ?
									onRejected :
									Thrower
							)
						);
					} ).promise();
				},

				// Get a promise for this deferred
				// If obj is provided, the promise aspect is added to the object
				promise: function( obj ) {
					return obj != null ? jQuery.extend( obj, promise ) : promise;
				}
			},
			deferred = {};

		// Add list-specific methods
		jQuery.each( tuples, function( i, tuple ) {
			var list = tuple[ 2 ],
				stateString = tuple[ 5 ];

			// promise.progress = list.add
			// promise.done = list.add
			// promise.fail = list.add
			promise[ tuple[ 1 ] ] = list.add;

			// Handle state
			if ( stateString ) {
				list.add(
					function() {

						// state = "resolved" (i.e., fulfilled)
						// state = "rejected"
						state = stateString;
					},

					// rejected_callbacks.disable
					// fulfilled_callbacks.disable
					tuples[ 3 - i ][ 2 ].disable,

					// rejected_handlers.disable
					// fulfilled_handlers.disable
					tuples[ 3 - i ][ 3 ].disable,

					// progress_callbacks.lock
					tuples[ 0 ][ 2 ].lock,

					// progress_handlers.lock
					tuples[ 0 ][ 3 ].lock
				);
			}

			// progress_handlers.fire
			// fulfilled_handlers.fire
			// rejected_handlers.fire
			list.add( tuple[ 3 ].fire );

			// deferred.notify = function() { deferred.notifyWith(...) }
			// deferred.resolve = function() { deferred.resolveWith(...) }
			// deferred.reject = function() { deferred.rejectWith(...) }
			deferred[ tuple[ 0 ] ] = function() {
				deferred[ tuple[ 0 ] + "With" ]( this === deferred ? undefined : this, arguments );
				return this;
			};

			// deferred.notifyWith = list.fireWith
			// deferred.resolveWith = list.fireWith
			// deferred.rejectWith = list.fireWith
			deferred[ tuple[ 0 ] + "With" ] = list.fireWith;
		} );

		// Make the deferred a promise
		promise.promise( deferred );

		// Call given func if any
		if ( func ) {
			func.call( deferred, deferred );
		}

		// All done!
		return deferred;
	},

	// Deferred helper
	when: function( singleValue ) {
		var

			// count of uncompleted subordinates
			remaining = arguments.length,

			// count of unprocessed arguments
			i = remaining,

			// subordinate fulfillment data
			resolveContexts = Array( i ),
			resolveValues = slice.call( arguments ),

			// the master Deferred
			master = jQuery.Deferred(),

			// subordinate callback factory
			updateFunc = function( i ) {
				return function( value ) {
					resolveContexts[ i ] = this;
					resolveValues[ i ] = arguments.length > 1 ? slice.call( arguments ) : value;
					if ( !( --remaining ) ) {
						master.resolveWith( resolveContexts, resolveValues );
					}
				};
			};

		// Single- and empty arguments are adopted like Promise.resolve
		if ( remaining <= 1 ) {
			adoptValue( singleValue, master.done( updateFunc( i ) ).resolve, master.reject,
				!remaining );

			// Use .then() to unwrap secondary thenables (cf. gh-3000)
			if ( master.state() === "pending" ||
				isFunction( resolveValues[ i ] && resolveValues[ i ].then ) ) {

				return master.then();
			}
		}

		// Multiple arguments are aggregated like Promise.all array elements
		while ( i-- ) {
			adoptValue( resolveValues[ i ], updateFunc( i ), master.reject );
		}

		return master.promise();
	}
} );


// These usually indicate a programmer mistake during development,
// warn about them ASAP rather than swallowing them by default.
var rerrorNames = /^(Eval|Internal|Range|Reference|Syntax|Type|URI)Error$/;

jQuery.Deferred.exceptionHook = function( error, stack ) {

	// Support: IE 8 - 9 only
	// Console exists when dev tools are open, which can happen at any time
	if ( window.console && window.console.warn && error && rerrorNames.test( error.name ) ) {
		window.console.warn( "jQuery.Deferred exception: " + error.message, error.stack, stack );
	}
};


jQuery.readyException = function( error ) {
	window.setTimeout( function() {
		throw error;
	} );
};


// The deferred used on DOM ready
var readyList = jQuery.Deferred();

jQuery.fn.ready = function( fn ) {

	readyList
		.then( fn )

		// Wrap jQuery.readyException in a function so that the lookup
		// happens at the time of error handling instead of callback
		// registration.
		.catch( function( error ) {
			jQuery.readyException( error );
		} );

	return this;
};

jQuery.extend( {

	// Is the DOM ready to be used? Set to true once it occurs.
	isReady: false,

	// A counter to track how many items to wait for before
	// the ready event fires. See #6781
	readyWait: 1,

	// Handle when the DOM is ready
	ready: function( wait ) {

		// Abort if there are pending holds or we're already ready
		if ( wait === true ? --jQuery.readyWait : jQuery.isReady ) {
			return;
		}

		// Remember that the DOM is ready
		jQuery.isReady = true;

		// If a normal DOM Ready event fired, decrement, and wait if need be
		if ( wait !== true && --jQuery.readyWait > 0 ) {
			return;
		}

		// If there are functions bound, to execute
		readyList.resolveWith( document, [ jQuery ] );
	}
} );

jQuery.ready.then = readyList.then;

// The ready event handler and self cleanup method
function completed() {
	document.removeEventListener( "DOMContentLoaded", completed );
	window.removeEventListener( "load", completed );
	jQuery.ready();
}

// Catch cases where $(document).ready() is called
// after the browser event has already occurred.
// Support: IE <=9 - 10 only
// Older IE sometimes signals "interactive" too soon
if ( document.readyState === "complete" ||
	( document.readyState !== "loading" && !document.documentElement.doScroll ) ) {

	// Handle it asynchronously to allow scripts the opportunity to delay ready
	window.setTimeout( jQuery.ready );

} else {

	// Use the handy event callback
	document.addEventListener( "DOMContentLoaded", completed );

	// A fallback to window.onload, that will always work
	window.addEventListener( "load", completed );
}


// Multifunctional method to get and set values of a collection
// The value/s can optionally be executed if it's a function
var access = function( elems, fn, key, value, chainable, emptyGet, raw ) {
	var i = 0,
		len = elems.length,
		bulk = key == null;

	// Sets many values
	if ( toType( key ) === "object" ) {
		chainable = true;
		for ( i in key ) {
			access( elems, fn, i, key[ i ], true, emptyGet, raw );
		}

	// Sets one value
	} else if ( value !== undefined ) {
		chainable = true;

		if ( !isFunction( value ) ) {
			raw = true;
		}

		if ( bulk ) {

			// Bulk operations run against the entire set
			if ( raw ) {
				fn.call( elems, value );
				fn = null;

			// ...except when executing function values
			} else {
				bulk = fn;
				fn = function( elem, key, value ) {
					return bulk.call( jQuery( elem ), value );
				};
			}
		}

		if ( fn ) {
			for ( ; i < len; i++ ) {
				fn(
					elems[ i ], key, raw ?
					value :
					value.call( elems[ i ], i, fn( elems[ i ], key ) )
				);
			}
		}
	}

	if ( chainable ) {
		return elems;
	}

	// Gets
	if ( bulk ) {
		return fn.call( elems );
	}

	return len ? fn( elems[ 0 ], key ) : emptyGet;
};


// Matches dashed string for camelizing
var rmsPrefix = /^-ms-/,
	rdashAlpha = /-([a-z])/g;

// Used by camelCase as callback to replace()
function fcamelCase( all, letter ) {
	return letter.toUpperCase();
}

// Convert dashed to camelCase; used by the css and data modules
// Support: IE <=9 - 11, Edge 12 - 15
// Microsoft forgot to hump their vendor prefix (#9572)
function camelCase( string ) {
	return string.replace( rmsPrefix, "ms-" ).replace( rdashAlpha, fcamelCase );
}
var acceptData = function( owner ) {

	// Accepts only:
	//  - Node
	//    - Node.ELEMENT_NODE
	//    - Node.DOCUMENT_NODE
	//  - Object
	//    - Any
	return owner.nodeType === 1 || owner.nodeType === 9 || !( +owner.nodeType );
};


function Data() {
	this.expando = jQuery.expando + Data.uid++;
}

Data.uid = 1;

Data.prototype = {

	cache: function( owner ) {

		// Check if the owner object already has a cache
		var value = owner[ this.expando ];

		// If not, create one
		if ( !value ) {
			value = {};

			// We can accept data for non-element nodes in modern browsers,
			// but we should not, see #8335.
			// Always return an empty object.
			if ( acceptData( owner ) ) {

				// If it is a node unlikely to be stringify-ed or looped over
				// use plain assignment
				if ( owner.nodeType ) {
					owner[ this.expando ] = value;

				// Otherwise secure it in a non-enumerable property
				// configurable must be true to allow the property to be
				// deleted when data is removed
				} else {
					Object.defineProperty( owner, this.expando, {
						value: value,
						configurable: true
					} );
				}
			}
		}

		return value;
	},
	set: function( owner, data, value ) {
		var prop,
			cache = this.cache( owner );

		// Handle: [ owner, key, value ] args
		// Always use camelCase key (gh-2257)
		if ( typeof data === "string" ) {
			cache[ camelCase( data ) ] = value;

		// Handle: [ owner, { properties } ] args
		} else {

			// Copy the properties one-by-one to the cache object
			for ( prop in data ) {
				cache[ camelCase( prop ) ] = data[ prop ];
			}
		}
		return cache;
	},
	get: function( owner, key ) {
		return key === undefined ?
			this.cache( owner ) :

			// Always use camelCase key (gh-2257)
			owner[ this.expando ] && owner[ this.expando ][ camelCase( key ) ];
	},
	access: function( owner, key, value ) {

		// In cases where either:
		//
		//   1. No key was specified
		//   2. A string key was specified, but no value provided
		//
		// Take the "read" path and allow the get method to determine
		// which value to return, respectively either:
		//
		//   1. The entire cache object
		//   2. The data stored at the key
		//
		if ( key === undefined ||
				( ( key && typeof key === "string" ) && value === undefined ) ) {

			return this.get( owner, key );
		}

		// When the key is not a string, or both a key and value
		// are specified, set or extend (existing objects) with either:
		//
		//   1. An object of properties
		//   2. A key and value
		//
		this.set( owner, key, value );

		// Since the "set" path can have two possible entry points
		// return the expected data based on which path was taken[*]
		return value !== undefined ? value : key;
	},
	remove: function( owner, key ) {
		var i,
			cache = owner[ this.expando ];

		if ( cache === undefined ) {
			return;
		}

		if ( key !== undefined ) {

			// Support array or space separated string of keys
			if ( Array.isArray( key ) ) {

				// If key is an array of keys...
				// We always set camelCase keys, so remove that.
				key = key.map( camelCase );
			} else {
				key = camelCase( key );

				// If a key with the spaces exists, use it.
				// Otherwise, create an array by matching non-whitespace
				key = key in cache ?
					[ key ] :
					( key.match( rnothtmlwhite ) || [] );
			}

			i = key.length;

			while ( i-- ) {
				delete cache[ key[ i ] ];
			}
		}

		// Remove the expando if there's no more data
		if ( key === undefined || jQuery.isEmptyObject( cache ) ) {

			// Support: Chrome <=35 - 45
			// Webkit & Blink performance suffers when deleting properties
			// from DOM nodes, so set to undefined instead
			// https://bugs.chromium.org/p/chromium/issues/detail?id=378607 (bug restricted)
			if ( owner.nodeType ) {
				owner[ this.expando ] = undefined;
			} else {
				delete owner[ this.expando ];
			}
		}
	},
	hasData: function( owner ) {
		var cache = owner[ this.expando ];
		return cache !== undefined && !jQuery.isEmptyObject( cache );
	}
};
var dataPriv = new Data();

var dataUser = new Data();


//	Implementation Summary
//
//	1. Enforce API surface and semantic compatibility with 1.9.x branch
//	2. Improve the module's maintainability by reducing the storage
//		paths to a single mechanism.
//	3. Use the same single mechanism to support "private" and "user" data.
//	4. _Never_ expose "private" data to user code (TODO: Drop _data, _removeData)
//	5. Avoid exposing implementation details on user objects (eg. expando properties)
//	6. Provide a clear path for implementation upgrade to WeakMap in 2014

var rbrace = /^(?:\{[\w\W]*\}|\[[\w\W]*\])$/,
	rmultiDash = /[A-Z]/g;

function getData( data ) {
	if ( data === "true" ) {
		return true;
	}

	if ( data === "false" ) {
		return false;
	}

	if ( data === "null" ) {
		return null;
	}

	// Only convert to a number if it doesn't change the string
	if ( data === +data + "" ) {
		return +data;
	}

	if ( rbrace.test( data ) ) {
		return JSON.parse( data );
	}

	return data;
}

function dataAttr( elem, key, data ) {
	var name;

	// If nothing was found internally, try to fetch any
	// data from the HTML5 data-* attribute
	if ( data === undefined && elem.nodeType === 1 ) {
		name = "data-" + key.replace( rmultiDash, "-$&" ).toLowerCase();
		data = elem.getAttribute( name );

		if ( typeof data === "string" ) {
			try {
				data = getData( data );
			} catch ( e ) {}

			// Make sure we set the data so it isn't changed later
			dataUser.set( elem, key, data );
		} else {
			data = undefined;
		}
	}
	return data;
}

jQuery.extend( {
	hasData: function( elem ) {
		return dataUser.hasData( elem ) || dataPriv.hasData( elem );
	},

	data: function( elem, name, data ) {
		return dataUser.access( elem, name, data );
	},

	removeData: function( elem, name ) {
		dataUser.remove( elem, name );
	},

	// TODO: Now that all calls to _data and _removeData have been replaced
	// with direct calls to dataPriv methods, these can be deprecated.
	_data: function( elem, name, data ) {
		return dataPriv.access( elem, name, data );
	},

	_removeData: function( elem, name ) {
		dataPriv.remove( elem, name );
	}
} );

jQuery.fn.extend( {
	data: function( key, value ) {
		var i, name, data,
			elem = this[ 0 ],
			attrs = elem && elem.attributes;

		// Gets all values
		if ( key === undefined ) {
			if ( this.length ) {
				data = dataUser.get( elem );

				if ( elem.nodeType === 1 && !dataPriv.get( elem, "hasDataAttrs" ) ) {
					i = attrs.length;
					while ( i-- ) {

						// Support: IE 11 only
						// The attrs elements can be null (#14894)
						if ( attrs[ i ] ) {
							name = attrs[ i ].name;
							if ( name.indexOf( "data-" ) === 0 ) {
								name = camelCase( name.slice( 5 ) );
								dataAttr( elem, name, data[ name ] );
							}
						}
					}
					dataPriv.set( elem, "hasDataAttrs", true );
				}
			}

			return data;
		}

		// Sets multiple values
		if ( typeof key === "object" ) {
			return this.each( function() {
				dataUser.set( this, key );
			} );
		}

		return access( this, function( value ) {
			var data;

			// The calling jQuery object (element matches) is not empty
			// (and therefore has an element appears at this[ 0 ]) and the
			// `value` parameter was not undefined. An empty jQuery object
			// will result in `undefined` for elem = this[ 0 ] which will
			// throw an exception if an attempt to read a data cache is made.
			if ( elem && value === undefined ) {

				// Attempt to get data from the cache
				// The key will always be camelCased in Data
				data = dataUser.get( elem, key );
				if ( data !== undefined ) {
					return data;
				}

				// Attempt to "discover" the data in
				// HTML5 custom data-* attrs
				data = dataAttr( elem, key );
				if ( data !== undefined ) {
					return data;
				}

				// We tried really hard, but the data doesn't exist.
				return;
			}

			// Set the data...
			this.each( function() {

				// We always store the camelCased key
				dataUser.set( this, key, value );
			} );
		}, null, value, arguments.length > 1, null, true );
	},

	removeData: function( key ) {
		return this.each( function() {
			dataUser.remove( this, key );
		} );
	}
} );


jQuery.extend( {
	queue: function( elem, type, data ) {
		var queue;

		if ( elem ) {
			type = ( type || "fx" ) + "queue";
			queue = dataPriv.get( elem, type );

			// Speed up dequeue by getting out quickly if this is just a lookup
			if ( data ) {
				if ( !queue || Array.isArray( data ) ) {
					queue = dataPriv.access( elem, type, jQuery.makeArray( data ) );
				} else {
					queue.push( data );
				}
			}
			return queue || [];
		}
	},

	dequeue: function( elem, type ) {
		type = type || "fx";

		var queue = jQuery.queue( elem, type ),
			startLength = queue.length,
			fn = queue.shift(),
			hooks = jQuery._queueHooks( elem, type ),
			next = function() {
				jQuery.dequeue( elem, type );
			};

		// If the fx queue is dequeued, always remove the progress sentinel
		if ( fn === "inprogress" ) {
			fn = queue.shift();
			startLength--;
		}

		if ( fn ) {

			// Add a progress sentinel to prevent the fx queue from being
			// automatically dequeued
			if ( type === "fx" ) {
				queue.unshift( "inprogress" );
			}

			// Clear up the last queue stop function
			delete hooks.stop;
			fn.call( elem, next, hooks );
		}

		if ( !startLength && hooks ) {
			hooks.empty.fire();
		}
	},

	// Not public - generate a queueHooks object, or return the current one
	_queueHooks: function( elem, type ) {
		var key = type + "queueHooks";
		return dataPriv.get( elem, key ) || dataPriv.access( elem, key, {
			empty: jQuery.Callbacks( "once memory" ).add( function() {
				dataPriv.remove( elem, [ type + "queue", key ] );
			} )
		} );
	}
} );

jQuery.fn.extend( {
	queue: function( type, data ) {
		var setter = 2;

		if ( typeof type !== "string" ) {
			data = type;
			type = "fx";
			setter--;
		}

		if ( arguments.length < setter ) {
			return jQuery.queue( this[ 0 ], type );
		}

		return data === undefined ?
			this :
			this.each( function() {
				var queue = jQuery.queue( this, type, data );

				// Ensure a hooks for this queue
				jQuery._queueHooks( this, type );

				if ( type === "fx" && queue[ 0 ] !== "inprogress" ) {
					jQuery.dequeue( this, type );
				}
			} );
	},
	dequeue: function( type ) {
		return this.each( function() {
			jQuery.dequeue( this, type );
		} );
	},
	clearQueue: function( type ) {
		return this.queue( type || "fx", [] );
	},

	// Get a promise resolved when queues of a certain type
	// are emptied (fx is the type by default)
	promise: function( type, obj ) {
		var tmp,
			count = 1,
			defer = jQuery.Deferred(),
			elements = this,
			i = this.length,
			resolve = function() {
				if ( !( --count ) ) {
					defer.resolveWith( elements, [ elements ] );
				}
			};

		if ( typeof type !== "string" ) {
			obj = type;
			type = undefined;
		}
		type = type || "fx";

		while ( i-- ) {
			tmp = dataPriv.get( elements[ i ], type + "queueHooks" );
			if ( tmp && tmp.empty ) {
				count++;
				tmp.empty.add( resolve );
			}
		}
		resolve();
		return defer.promise( obj );
	}
} );
var pnum = ( /[+-]?(?:\d*\.|)\d+(?:[eE][+-]?\d+|)/ ).source;

var rcssNum = new RegExp( "^(?:([+-])=|)(" + pnum + ")([a-z%]*)$", "i" );


var cssExpand = [ "Top", "Right", "Bottom", "Left" ];

var documentElement = document.documentElement;


	var isAttached = function( elem ) {
			return jQuery.contains( elem.ownerDocument, elem );
		},
		composed = { composed: true };

	// Support: IE 9 - 11+, Edge 12 - 18+, iOS 10.0 - 10.2 only
	// Check attachment across shadow DOM boundaries when possible (gh-3504)
	// Support: iOS 10.0-10.2 only
	// Early iOS 10 versions support `attachShadow` but not `getRootNode`,
	// leading to errors. We need to check for `getRootNode`.
	if ( documentElement.getRootNode ) {
		isAttached = function( elem ) {
			return jQuery.contains( elem.ownerDocument, elem ) ||
				elem.getRootNode( composed ) === elem.ownerDocument;
		};
	}
var isHiddenWithinTree = function( elem, el ) {

		// isHiddenWithinTree might be called from jQuery#filter function;
		// in that case, element will be second argument
		elem = el || elem;

		// Inline style trumps all
		return elem.style.display === "none" ||
			elem.style.display === "" &&

			// Otherwise, check computed style
			// Support: Firefox <=43 - 45
			// Disconnected elements can have computed display: none, so first confirm that elem is
			// in the document.
			isAttached( elem ) &&

			jQuery.css( elem, "display" ) === "none";
	};

var swap = function( elem, options, callback, args ) {
	var ret, name,
		old = {};

	// Remember the old values, and insert the new ones
	for ( name in options ) {
		old[ name ] = elem.style[ name ];
		elem.style[ name ] = options[ name ];
	}

	ret = callback.apply( elem, args || [] );

	// Revert the old values
	for ( name in options ) {
		elem.style[ name ] = old[ name ];
	}

	return ret;
};


function adjustCSS( elem, prop, valueParts, tween ) {
	var adjusted, scale,
		maxIterations = 20,
		currentValue = tween ?
			function() {
				return tween.cur();
			} :
			function() {
				return jQuery.css( elem, prop, "" );
			},
		initial = currentValue(),
		unit = valueParts && valueParts[ 3 ] || ( jQuery.cssNumber[ prop ] ? "" : "px" ),

		// Starting value computation is required for potential unit mismatches
		initialInUnit = elem.nodeType &&
			( jQuery.cssNumber[ prop ] || unit !== "px" && +initial ) &&
			rcssNum.exec( jQuery.css( elem, prop ) );

	if ( initialInUnit && initialInUnit[ 3 ] !== unit ) {

		// Support: Firefox <=54
		// Halve the iteration target value to prevent interference from CSS upper bounds (gh-2144)
		initial = initial / 2;

		// Trust units reported by jQuery.css
		unit = unit || initialInUnit[ 3 ];

		// Iteratively approximate from a nonzero starting point
		initialInUnit = +initial || 1;

		while ( maxIterations-- ) {

			// Evaluate and update our best guess (doubling guesses that zero out).
			// Finish if the scale equals or crosses 1 (making the old*new product non-positive).
			jQuery.style( elem, prop, initialInUnit + unit );
			if ( ( 1 - scale ) * ( 1 - ( scale = currentValue() / initial || 0.5 ) ) <= 0 ) {
				maxIterations = 0;
			}
			initialInUnit = initialInUnit / scale;

		}

		initialInUnit = initialInUnit * 2;
		jQuery.style( elem, prop, initialInUnit + unit );

		// Make sure we update the tween properties later on
		valueParts = valueParts || [];
	}

	if ( valueParts ) {
		initialInUnit = +initialInUnit || +initial || 0;

		// Apply relative offset (+=/-=) if specified
		adjusted = valueParts[ 1 ] ?
			initialInUnit + ( valueParts[ 1 ] + 1 ) * valueParts[ 2 ] :
			+valueParts[ 2 ];
		if ( tween ) {
			tween.unit = unit;
			tween.start = initialInUnit;
			tween.end = adjusted;
		}
	}
	return adjusted;
}


var defaultDisplayMap = {};

function getDefaultDisplay( elem ) {
	var temp,
		doc = elem.ownerDocument,
		nodeName = elem.nodeName,
		display = defaultDisplayMap[ nodeName ];

	if ( display ) {
		return display;
	}

	temp = doc.body.appendChild( doc.createElement( nodeName ) );
	display = jQuery.css( temp, "display" );

	temp.parentNode.removeChild( temp );

	if ( display === "none" ) {
		display = "block";
	}
	defaultDisplayMap[ nodeName ] = display;

	return display;
}

function showHide( elements, show ) {
	var display, elem,
		values = [],
		index = 0,
		length = elements.length;

	// Determine new display value for elements that need to change
	for ( ; index < length; index++ ) {
		elem = elements[ index ];
		if ( !elem.style ) {
			continue;
		}

		display = elem.style.display;
		if ( show ) {

			// Since we force visibility upon cascade-hidden elements, an immediate (and slow)
			// check is required in this first loop unless we have a nonempty display value (either
			// inline or about-to-be-restored)
			if ( display === "none" ) {
				values[ index ] = dataPriv.get( elem, "display" ) || null;
				if ( !values[ index ] ) {
					elem.style.display = "";
				}
			}
			if ( elem.style.display === "" && isHiddenWithinTree( elem ) ) {
				values[ index ] = getDefaultDisplay( elem );
			}
		} else {
			if ( display !== "none" ) {
				values[ index ] = "none";

				// Remember what we're overwriting
				dataPriv.set( elem, "display", display );
			}
		}
	}

	// Set the display of the elements in a second loop to avoid constant reflow
	for ( index = 0; index < length; index++ ) {
		if ( values[ index ] != null ) {
			elements[ index ].style.display = values[ index ];
		}
	}

	return elements;
}

jQuery.fn.extend( {
	show: function() {
		return showHide( this, true );
	},
	hide: function() {
		return showHide( this );
	},
	toggle: function( state ) {
		if ( typeof state === "boolean" ) {
			return state ? this.show() : this.hide();
		}

		return this.each( function() {
			if ( isHiddenWithinTree( this ) ) {
				jQuery( this ).show();
			} else {
				jQuery( this ).hide();
			}
		} );
	}
} );
var rcheckableType = ( /^(?:checkbox|radio)$/i );

var rtagName = ( /<([a-z][^\/\0>\x20\t\r\n\f]*)/i );

var rscriptType = ( /^$|^module$|\/(?:java|ecma)script/i );


// We have to close these tags to support XHTML (#13200)
var wrapMap = {

	// Support: IE <=9 only
	option: [ 1, "<select multiple='multiple'>", "</select>" ],

	// XHTML parsers do not magically insert elements in the
	// same way that tag soup parsers do. So we cannot shorten
	// this by omitting <tbody> or other required elements.
	thead: [ 1, "<table>", "</table>" ],
	col: [ 2, "<table><colgroup>", "</colgroup></table>" ],
	tr: [ 2, "<table><tbody>", "</tbody></table>" ],
	td: [ 3, "<table><tbody><tr>", "</tr></tbody></table>" ],

	_default: [ 0, "", "" ]
};

// Support: IE <=9 only
wrapMap.optgroup = wrapMap.option;

wrapMap.tbody = wrapMap.tfoot = wrapMap.colgroup = wrapMap.caption = wrapMap.thead;
wrapMap.th = wrapMap.td;


function getAll( context, tag ) {

	// Support: IE <=9 - 11 only
	// Use typeof to avoid zero-argument method invocation on host objects (#15151)
	var ret;

	if ( typeof context.getElementsByTagName !== "undefined" ) {
		ret = context.getElementsByTagName( tag || "*" );

	} else if ( typeof context.querySelectorAll !== "undefined" ) {
		ret = context.querySelectorAll( tag || "*" );

	} else {
		ret = [];
	}

	if ( tag === undefined || tag && nodeName( context, tag ) ) {
		return jQuery.merge( [ context ], ret );
	}

	return ret;
}


// Mark scripts as having already been evaluated
function setGlobalEval( elems, refElements ) {
	var i = 0,
		l = elems.length;

	for ( ; i < l; i++ ) {
		dataPriv.set(
			elems[ i ],
			"globalEval",
			!refElements || dataPriv.get( refElements[ i ], "globalEval" )
		);
	}
}


var rhtml = /<|&#?\w+;/;

function buildFragment( elems, context, scripts, selection, ignored ) {
	var elem, tmp, tag, wrap, attached, j,
		fragment = context.createDocumentFragment(),
		nodes = [],
		i = 0,
		l = elems.length;

	for ( ; i < l; i++ ) {
		elem = elems[ i ];

		if ( elem || elem === 0 ) {

			// Add nodes directly
			if ( toType( elem ) === "object" ) {

				// Support: Android <=4.0 only, PhantomJS 1 only
				// push.apply(_, arraylike) throws on ancient WebKit
				jQuery.merge( nodes, elem.nodeType ? [ elem ] : elem );

			// Convert non-html into a text node
			} else if ( !rhtml.test( elem ) ) {
				nodes.push( context.createTextNode( elem ) );

			// Convert html into DOM nodes
			} else {
				tmp = tmp || fragment.appendChild( context.createElement( "div" ) );

				// Deserialize a standard representation
				tag = ( rtagName.exec( elem ) || [ "", "" ] )[ 1 ].toLowerCase();
				wrap = wrapMap[ tag ] || wrapMap._default;
				tmp.innerHTML = wrap[ 1 ] + jQuery.htmlPrefilter( elem ) + wrap[ 2 ];

				// Descend through wrappers to the right content
				j = wrap[ 0 ];
				while ( j-- ) {
					tmp = tmp.lastChild;
				}

				// Support: Android <=4.0 only, PhantomJS 1 only
				// push.apply(_, arraylike) throws on ancient WebKit
				jQuery.merge( nodes, tmp.childNodes );

				// Remember the top-level container
				tmp = fragment.firstChild;

				// Ensure the created nodes are orphaned (#12392)
				tmp.textContent = "";
			}
		}
	}

	// Remove wrapper from fragment
	fragment.textContent = "";

	i = 0;
	while ( ( elem = nodes[ i++ ] ) ) {

		// Skip elements already in the context collection (trac-4087)
		if ( selection && jQuery.inArray( elem, selection ) > -1 ) {
			if ( ignored ) {
				ignored.push( elem );
			}
			continue;
		}

		attached = isAttached( elem );

		// Append to fragment
		tmp = getAll( fragment.appendChild( elem ), "script" );

		// Preserve script evaluation history
		if ( attached ) {
			setGlobalEval( tmp );
		}

		// Capture executables
		if ( scripts ) {
			j = 0;
			while ( ( elem = tmp[ j++ ] ) ) {
				if ( rscriptType.test( elem.type || "" ) ) {
					scripts.push( elem );
				}
			}
		}
	}

	return fragment;
}


( function() {
	var fragment = document.createDocumentFragment(),
		div = fragment.appendChild( document.createElement( "div" ) ),
		input = document.createElement( "input" );

	// Support: Android 4.0 - 4.3 only
	// Check state lost if the name is set (#11217)
	// Support: Windows Web Apps (WWA)
	// `name` and `type` must use .setAttribute for WWA (#14901)
	input.setAttribute( "type", "radio" );
	input.setAttribute( "checked", "checked" );
	input.setAttribute( "name", "t" );

	div.appendChild( input );

	// Support: Android <=4.1 only
	// Older WebKit doesn't clone checked state correctly in fragments
	support.checkClone = div.cloneNode( true ).cloneNode( true ).lastChild.checked;

	// Support: IE <=11 only
	// Make sure textarea (and checkbox) defaultValue is properly cloned
	div.innerHTML = "<textarea>x</textarea>";
	support.noCloneChecked = !!div.cloneNode( true ).lastChild.defaultValue;
} )();


var
	rkeyEvent = /^key/,
	rmouseEvent = /^(?:mouse|pointer|contextmenu|drag|drop)|click/,
	rtypenamespace = /^([^.]*)(?:\.(.+)|)/;

function returnTrue() {
	return true;
}

function returnFalse() {
	return false;
}

// Support: IE <=9 - 11+
// focus() and blur() are asynchronous, except when they are no-op.
// So expect focus to be synchronous when the element is already active,
// and blur to be synchronous when the element is not already active.
// (focus and blur are always synchronous in other supported browsers,
// this just defines when we can count on it).
function expectSync( elem, type ) {
	return ( elem === safeActiveElement() ) === ( type === "focus" );
}

// Support: IE <=9 only
// Accessing document.activeElement can throw unexpectedly
// https://bugs.jquery.com/ticket/13393
function safeActiveElement() {
	try {
		return document.activeElement;
	} catch ( err ) { }
}

function on( elem, types, selector, data, fn, one ) {
	var origFn, type;

	// Types can be a map of types/handlers
	if ( typeof types === "object" ) {

		// ( types-Object, selector, data )
		if ( typeof selector !== "string" ) {

			// ( types-Object, data )
			data = data || selector;
			selector = undefined;
		}
		for ( type in types ) {
			on( elem, type, selector, data, types[ type ], one );
		}
		return elem;
	}

	if ( data == null && fn == null ) {

		// ( types, fn )
		fn = selector;
		data = selector = undefined;
	} else if ( fn == null ) {
		if ( typeof selector === "string" ) {

			// ( types, selector, fn )
			fn = data;
			data = undefined;
		} else {

			// ( types, data, fn )
			fn = data;
			data = selector;
			selector = undefined;
		}
	}
	if ( fn === false ) {
		fn = returnFalse;
	} else if ( !fn ) {
		return elem;
	}

	if ( one === 1 ) {
		origFn = fn;
		fn = function( event ) {

			// Can use an empty set, since event contains the info
			jQuery().off( event );
			return origFn.apply( this, arguments );
		};

		// Use same guid so caller can remove using origFn
		fn.guid = origFn.guid || ( origFn.guid = jQuery.guid++ );
	}
	return elem.each( function() {
		jQuery.event.add( this, types, fn, data, selector );
	} );
}

/*
 * Helper functions for managing events -- not part of the public interface.
 * Props to Dean Edwards' addEvent library for many of the ideas.
 */
jQuery.event = {

	global: {},

	add: function( elem, types, handler, data, selector ) {

		var handleObjIn, eventHandle, tmp,
			events, t, handleObj,
			special, handlers, type, namespaces, origType,
			elemData = dataPriv.get( elem );

		// Don't attach events to noData or text/comment nodes (but allow plain objects)
		if ( !elemData ) {
			return;
		}

		// Caller can pass in an object of custom data in lieu of the handler
		if ( handler.handler ) {
			handleObjIn = handler;
			handler = handleObjIn.handler;
			selector = handleObjIn.selector;
		}

		// Ensure that invalid selectors throw exceptions at attach time
		// Evaluate against documentElement in case elem is a non-element node (e.g., document)
		if ( selector ) {
			jQuery.find.matchesSelector( documentElement, selector );
		}

		// Make sure that the handler has a unique ID, used to find/remove it later
		if ( !handler.guid ) {
			handler.guid = jQuery.guid++;
		}

		// Init the element's event structure and main handler, if this is the first
		if ( !( events = elemData.events ) ) {
			events = elemData.events = {};
		}
		if ( !( eventHandle = elemData.handle ) ) {
			eventHandle = elemData.handle = function( e ) {

				// Discard the second event of a jQuery.event.trigger() and
				// when an event is called after a page has unloaded
				return typeof jQuery !== "undefined" && jQuery.event.triggered !== e.type ?
					jQuery.event.dispatch.apply( elem, arguments ) : undefined;
			};
		}

		// Handle multiple events separated by a space
		types = ( types || "" ).match( rnothtmlwhite ) || [ "" ];
		t = types.length;
		while ( t-- ) {
			tmp = rtypenamespace.exec( types[ t ] ) || [];
			type = origType = tmp[ 1 ];
			namespaces = ( tmp[ 2 ] || "" ).split( "." ).sort();

			// There *must* be a type, no attaching namespace-only handlers
			if ( !type ) {
				continue;
			}

			// If event changes its type, use the special event handlers for the changed type
			special = jQuery.event.special[ type ] || {};

			// If selector defined, determine special event api type, otherwise given type
			type = ( selector ? special.delegateType : special.bindType ) || type;

			// Update special based on newly reset type
			special = jQuery.event.special[ type ] || {};

			// handleObj is passed to all event handlers
			handleObj = jQuery.extend( {
				type: type,
				origType: origType,
				data: data,
				handler: handler,
				guid: handler.guid,
				selector: selector,
				needsContext: selector && jQuery.expr.match.needsContext.test( selector ),
				namespace: namespaces.join( "." )
			}, handleObjIn );

			// Init the event handler queue if we're the first
			if ( !( handlers = events[ type ] ) ) {
				handlers = events[ type ] = [];
				handlers.delegateCount = 0;

				// Only use addEventListener if the special events handler returns false
				if ( !special.setup ||
					special.setup.call( elem, data, namespaces, eventHandle ) === false ) {

					if ( elem.addEventListener ) {
						elem.addEventListener( type, eventHandle );
					}
				}
			}

			if ( special.add ) {
				special.add.call( elem, handleObj );

				if ( !handleObj.handler.guid ) {
					handleObj.handler.guid = handler.guid;
				}
			}

			// Add to the element's handler list, delegates in front
			if ( selector ) {
				handlers.splice( handlers.delegateCount++, 0, handleObj );
			} else {
				handlers.push( handleObj );
			}

			// Keep track of which events have ever been used, for event optimization
			jQuery.event.global[ type ] = true;
		}

	},

	// Detach an event or set of events from an element
	remove: function( elem, types, handler, selector, mappedTypes ) {

		var j, origCount, tmp,
			events, t, handleObj,
			special, handlers, type, namespaces, origType,
			elemData = dataPriv.hasData( elem ) && dataPriv.get( elem );

		if ( !elemData || !( events = elemData.events ) ) {
			return;
		}

		// Once for each type.namespace in types; type may be omitted
		types = ( types || "" ).match( rnothtmlwhite ) || [ "" ];
		t = types.length;
		while ( t-- ) {
			tmp = rtypenamespace.exec( types[ t ] ) || [];
			type = origType = tmp[ 1 ];
			namespaces = ( tmp[ 2 ] || "" ).split( "." ).sort();

			// Unbind all events (on this namespace, if provided) for the element
			if ( !type ) {
				for ( type in events ) {
					jQuery.event.remove( elem, type + types[ t ], handler, selector, true );
				}
				continue;
			}

			special = jQuery.event.special[ type ] || {};
			type = ( selector ? special.delegateType : special.bindType ) || type;
			handlers = events[ type ] || [];
			tmp = tmp[ 2 ] &&
				new RegExp( "(^|\\.)" + namespaces.join( "\\.(?:.*\\.|)" ) + "(\\.|$)" );

			// Remove matching events
			origCount = j = handlers.length;
			while ( j-- ) {
				handleObj = handlers[ j ];

				if ( ( mappedTypes || origType === handleObj.origType ) &&
					( !handler || handler.guid === handleObj.guid ) &&
					( !tmp || tmp.test( handleObj.namespace ) ) &&
					( !selector || selector === handleObj.selector ||
						selector === "**" && handleObj.selector ) ) {
					handlers.splice( j, 1 );

					if ( handleObj.selector ) {
						handlers.delegateCount--;
					}
					if ( special.remove ) {
						special.remove.call( elem, handleObj );
					}
				}
			}

			// Remove generic event handler if we removed something and no more handlers exist
			// (avoids potential for endless recursion during removal of special event handlers)
			if ( origCount && !handlers.length ) {
				if ( !special.teardown ||
					special.teardown.call( elem, namespaces, elemData.handle ) === false ) {

					jQuery.removeEvent( elem, type, elemData.handle );
				}

				delete events[ type ];
			}
		}

		// Remove data and the expando if it's no longer used
		if ( jQuery.isEmptyObject( events ) ) {
			dataPriv.remove( elem, "handle events" );
		}
	},

	dispatch: function( nativeEvent ) {

		// Make a writable jQuery.Event from the native event object
		var event = jQuery.event.fix( nativeEvent );

		var i, j, ret, matched, handleObj, handlerQueue,
			args = new Array( arguments.length ),
			handlers = ( dataPriv.get( this, "events" ) || {} )[ event.type ] || [],
			special = jQuery.event.special[ event.type ] || {};

		// Use the fix-ed jQuery.Event rather than the (read-only) native event
		args[ 0 ] = event;

		for ( i = 1; i < arguments.length; i++ ) {
			args[ i ] = arguments[ i ];
		}

		event.delegateTarget = this;

		// Call the preDispatch hook for the mapped type, and let it bail if desired
		if ( special.preDispatch && special.preDispatch.call( this, event ) === false ) {
			return;
		}

		// Determine handlers
		handlerQueue = jQuery.event.handlers.call( this, event, handlers );

		// Run delegates first; they may want to stop propagation beneath us
		i = 0;
		while ( ( matched = handlerQueue[ i++ ] ) && !event.isPropagationStopped() ) {
			event.currentTarget = matched.elem;

			j = 0;
			while ( ( handleObj = matched.handlers[ j++ ] ) &&
				!event.isImmediatePropagationStopped() ) {

				// If the event is namespaced, then each handler is only invoked if it is
				// specially universal or its namespaces are a superset of the event's.
				if ( !event.rnamespace || handleObj.namespace === false ||
					event.rnamespace.test( handleObj.namespace ) ) {

					event.handleObj = handleObj;
					event.data = handleObj.data;

					ret = ( ( jQuery.event.special[ handleObj.origType ] || {} ).handle ||
						handleObj.handler ).apply( matched.elem, args );

					if ( ret !== undefined ) {
						if ( ( event.result = ret ) === false ) {
							event.preventDefault();
							event.stopPropagation();
						}
					}
				}
			}
		}

		// Call the postDispatch hook for the mapped type
		if ( special.postDispatch ) {
			special.postDispatch.call( this, event );
		}

		return event.result;
	},

	handlers: function( event, handlers ) {
		var i, handleObj, sel, matchedHandlers, matchedSelectors,
			handlerQueue = [],
			delegateCount = handlers.delegateCount,
			cur = event.target;

		// Find delegate handlers
		if ( delegateCount &&

			// Support: IE <=9
			// Black-hole SVG <use> instance trees (trac-13180)
			cur.nodeType &&

			// Support: Firefox <=42
			// Suppress spec-violating clicks indicating a non-primary pointer button (trac-3861)
			// https://www.w3.org/TR/DOM-Level-3-Events/#event-type-click
			// Support: IE 11 only
			// ...but not arrow key "clicks" of radio inputs, which can have `button` -1 (gh-2343)
			!( event.type === "click" && event.button >= 1 ) ) {

			for ( ; cur !== this; cur = cur.parentNode || this ) {

				// Don't check non-elements (#13208)
				// Don't process clicks on disabled elements (#6911, #8165, #11382, #11764)
				if ( cur.nodeType === 1 && !( event.type === "click" && cur.disabled === true ) ) {
					matchedHandlers = [];
					matchedSelectors = {};
					for ( i = 0; i < delegateCount; i++ ) {
						handleObj = handlers[ i ];

						// Don't conflict with Object.prototype properties (#13203)
						sel = handleObj.selector + " ";

						if ( matchedSelectors[ sel ] === undefined ) {
							matchedSelectors[ sel ] = handleObj.needsContext ?
								jQuery( sel, this ).index( cur ) > -1 :
								jQuery.find( sel, this, null, [ cur ] ).length;
						}
						if ( matchedSelectors[ sel ] ) {
							matchedHandlers.push( handleObj );
						}
					}
					if ( matchedHandlers.length ) {
						handlerQueue.push( { elem: cur, handlers: matchedHandlers } );
					}
				}
			}
		}

		// Add the remaining (directly-bound) handlers
		cur = this;
		if ( delegateCount < handlers.length ) {
			handlerQueue.push( { elem: cur, handlers: handlers.slice( delegateCount ) } );
		}

		return handlerQueue;
	},

	addProp: function( name, hook ) {
		Object.defineProperty( jQuery.Event.prototype, name, {
			enumerable: true,
			configurable: true,

			get: isFunction( hook ) ?
				function() {
					if ( this.originalEvent ) {
							return hook( this.originalEvent );
					}
				} :
				function() {
					if ( this.originalEvent ) {
							return this.originalEvent[ name ];
					}
				},

			set: function( value ) {
				Object.defineProperty( this, name, {
					enumerable: true,
					configurable: true,
					writable: true,
					value: value
				} );
			}
		} );
	},

	fix: function( originalEvent ) {
		return originalEvent[ jQuery.expando ] ?
			originalEvent :
			new jQuery.Event( originalEvent );
	},

	special: {
		load: {

			// Prevent triggered image.load events from bubbling to window.load
			noBubble: true
		},
		click: {

			// Utilize native event to ensure correct state for checkable inputs
			setup: function( data ) {

				// For mutual compressibility with _default, replace `this` access with a local var.
				// `|| data` is dead code meant only to preserve the variable through minification.
				var el = this || data;

				// Claim the first handler
				if ( rcheckableType.test( el.type ) &&
					el.click && nodeName( el, "input" ) ) {

					// dataPriv.set( el, "click", ... )
					leverageNative( el, "click", returnTrue );
				}

				// Return false to allow normal processing in the caller
				return false;
			},
			trigger: function( data ) {

				// For mutual compressibility with _default, replace `this` access with a local var.
				// `|| data` is dead code meant only to preserve the variable through minification.
				var el = this || data;

				// Force setup before triggering a click
				if ( rcheckableType.test( el.type ) &&
					el.click && nodeName( el, "input" ) ) {

					leverageNative( el, "click" );
				}

				// Return non-false to allow normal event-path propagation
				return true;
			},

			// For cross-browser consistency, suppress native .click() on links
			// Also prevent it if we're currently inside a leveraged native-event stack
			_default: function( event ) {
				var target = event.target;
				return rcheckableType.test( target.type ) &&
					target.click && nodeName( target, "input" ) &&
					dataPriv.get( target, "click" ) ||
					nodeName( target, "a" );
			}
		},

		beforeunload: {
			postDispatch: function( event ) {

				// Support: Firefox 20+
				// Firefox doesn't alert if the returnValue field is not set.
				if ( event.result !== undefined && event.originalEvent ) {
					event.originalEvent.returnValue = event.result;
				}
			}
		}
	}
};

// Ensure the presence of an event listener that handles manually-triggered
// synthetic events by interrupting progress until reinvoked in response to
// *native* events that it fires directly, ensuring that state changes have
// already occurred before other listeners are invoked.
function leverageNative( el, type, expectSync ) {

	// Missing expectSync indicates a trigger call, which must force setup through jQuery.event.add
	if ( !expectSync ) {
		if ( dataPriv.get( el, type ) === undefined ) {
			jQuery.event.add( el, type, returnTrue );
		}
		return;
	}

	// Register the controller as a special universal handler for all event namespaces
	dataPriv.set( el, type, false );
	jQuery.event.add( el, type, {
		namespace: false,
		handler: function( event ) {
			var notAsync, result,
				saved = dataPriv.get( this, type );

			if ( ( event.isTrigger & 1 ) && this[ type ] ) {

				// Interrupt processing of the outer synthetic .trigger()ed event
				// Saved data should be false in such cases, but might be a leftover capture object
				// from an async native handler (gh-4350)
				if ( !saved.length ) {

					// Store arguments for use when handling the inner native event
					// There will always be at least one argument (an event object), so this array
					// will not be confused with a leftover capture object.
					saved = slice.call( arguments );
					dataPriv.set( this, type, saved );

					// Trigger the native event and capture its result
					// Support: IE <=9 - 11+
					// focus() and blur() are asynchronous
					notAsync = expectSync( this, type );
					this[ type ]();
					result = dataPriv.get( this, type );
					if ( saved !== result || notAsync ) {
						dataPriv.set( this, type, false );
					} else {
						result = {};
					}
					if ( saved !== result ) {

						// Cancel the outer synthetic event
						event.stopImmediatePropagation();
						event.preventDefault();
						return result.value;
					}

				// If this is an inner synthetic event for an event with a bubbling surrogate
				// (focus or blur), assume that the surrogate already propagated from triggering the
				// native event and prevent that from happening again here.
				// This technically gets the ordering wrong w.r.t. to `.trigger()` (in which the
				// bubbling surrogate propagates *after* the non-bubbling base), but that seems
				// less bad than duplication.
				} else if ( ( jQuery.event.special[ type ] || {} ).delegateType ) {
					event.stopPropagation();
				}

			// If this is a native event triggered above, everything is now in order
			// Fire an inner synthetic event with the original arguments
			} else if ( saved.length ) {

				// ...and capture the result
				dataPriv.set( this, type, {
					value: jQuery.event.trigger(

						// Support: IE <=9 - 11+
						// Extend with the prototype to reset the above stopImmediatePropagation()
						jQuery.extend( saved[ 0 ], jQuery.Event.prototype ),
						saved.slice( 1 ),
						this
					)
				} );

				// Abort handling of the native event
				event.stopImmediatePropagation();
			}
		}
	} );
}

jQuery.removeEvent = function( elem, type, handle ) {

	// This "if" is needed for plain objects
	if ( elem.removeEventListener ) {
		elem.removeEventListener( type, handle );
	}
};

jQuery.Event = function( src, props ) {

	// Allow instantiation without the 'new' keyword
	if ( !( this instanceof jQuery.Event ) ) {
		return new jQuery.Event( src, props );
	}

	// Event object
	if ( src && src.type ) {
		this.originalEvent = src;
		this.type = src.type;

		// Events bubbling up the document may have been marked as prevented
		// by a handler lower down the tree; reflect the correct value.
		this.isDefaultPrevented = src.defaultPrevented ||
				src.defaultPrevented === undefined &&

				// Support: Android <=2.3 only
				src.returnValue === false ?
			returnTrue :
			returnFalse;

		// Create target properties
		// Support: Safari <=6 - 7 only
		// Target should not be a text node (#504, #13143)
		this.target = ( src.target && src.target.nodeType === 3 ) ?
			src.target.parentNode :
			src.target;

		this.currentTarget = src.currentTarget;
		this.relatedTarget = src.relatedTarget;

	// Event type
	} else {
		this.type = src;
	}

	// Put explicitly provided properties onto the event object
	if ( props ) {
		jQuery.extend( this, props );
	}

	// Create a timestamp if incoming event doesn't have one
	this.timeStamp = src && src.timeStamp || Date.now();

	// Mark it as fixed
	this[ jQuery.expando ] = true;
};

// jQuery.Event is based on DOM3 Events as specified by the ECMAScript Language Binding
// https://www.w3.org/TR/2003/WD-DOM-Level-3-Events-20030331/ecma-script-binding.html
jQuery.Event.prototype = {
	constructor: jQuery.Event,
	isDefaultPrevented: returnFalse,
	isPropagationStopped: returnFalse,
	isImmediatePropagationStopped: returnFalse,
	isSimulated: false,

	preventDefault: function() {
		var e = this.originalEvent;

		this.isDefaultPrevented = returnTrue;

		if ( e && !this.isSimulated ) {
			e.preventDefault();
		}
	},
	stopPropagation: function() {
		var e = this.originalEvent;

		this.isPropagationStopped = returnTrue;

		if ( e && !this.isSimulated ) {
			e.stopPropagation();
		}
	},
	stopImmediatePropagation: function() {
		var e = this.originalEvent;

		this.isImmediatePropagationStopped = returnTrue;

		if ( e && !this.isSimulated ) {
			e.stopImmediatePropagation();
		}

		this.stopPropagation();
	}
};

// Includes all common event props including KeyEvent and MouseEvent specific props
jQuery.each( {
	altKey: true,
	bubbles: true,
	cancelable: true,
	changedTouches: true,
	ctrlKey: true,
	detail: true,
	eventPhase: true,
	metaKey: true,
	pageX: true,
	pageY: true,
	shiftKey: true,
	view: true,
	"char": true,
	code: true,
	charCode: true,
	key: true,
	keyCode: true,
	button: true,
	buttons: true,
	clientX: true,
	clientY: true,
	offsetX: true,
	offsetY: true,
	pointerId: true,
	pointerType: true,
	screenX: true,
	screenY: true,
	targetTouches: true,
	toElement: true,
	touches: true,

	which: function( event ) {
		var button = event.button;

		// Add which for key events
		if ( event.which == null && rkeyEvent.test( event.type ) ) {
			return event.charCode != null ? event.charCode : event.keyCode;
		}

		// Add which for click: 1 === left; 2 === middle; 3 === right
		if ( !event.which && button !== undefined && rmouseEvent.test( event.type ) ) {
			if ( button & 1 ) {
				return 1;
			}

			if ( button & 2 ) {
				return 3;
			}

			if ( button & 4 ) {
				return 2;
			}

			return 0;
		}

		return event.which;
	}
}, jQuery.event.addProp );

jQuery.each( { focus: "focusin", blur: "focusout" }, function( type, delegateType ) {
	jQuery.event.special[ type ] = {

		// Utilize native event if possible so blur/focus sequence is correct
		setup: function() {

			// Claim the first handler
			// dataPriv.set( this, "focus", ... )
			// dataPriv.set( this, "blur", ... )
			leverageNative( this, type, expectSync );

			// Return false to allow normal processing in the caller
			return false;
		},
		trigger: function() {

			// Force setup before trigger
			leverageNative( this, type );

			// Return non-false to allow normal event-path propagation
			return true;
		},

		delegateType: delegateType
	};
} );

// Create mouseenter/leave events using mouseover/out and event-time checks
// so that event delegation works in jQuery.
// Do the same for pointerenter/pointerleave and pointerover/pointerout
//
// Support: Safari 7 only
// Safari sends mouseenter too often; see:
// https://bugs.chromium.org/p/chromium/issues/detail?id=470258
// for the description of the bug (it existed in older Chrome versions as well).
jQuery.each( {
	mouseenter: "mouseover",
	mouseleave: "mouseout",
	pointerenter: "pointerover",
	pointerleave: "pointerout"
}, function( orig, fix ) {
	jQuery.event.special[ orig ] = {
		delegateType: fix,
		bindType: fix,

		handle: function( event ) {
			var ret,
				target = this,
				related = event.relatedTarget,
				handleObj = event.handleObj;

			// For mouseenter/leave call the handler if related is outside the target.
			// NB: No relatedTarget if the mouse left/entered the browser window
			if ( !related || ( related !== target && !jQuery.contains( target, related ) ) ) {
				event.type = handleObj.origType;
				ret = handleObj.handler.apply( this, arguments );
				event.type = fix;
			}
			return ret;
		}
	};
} );

jQuery.fn.extend( {

	on: function( types, selector, data, fn ) {
		return on( this, types, selector, data, fn );
	},
	one: function( types, selector, data, fn ) {
		return on( this, types, selector, data, fn, 1 );
	},
	off: function( types, selector, fn ) {
		var handleObj, type;
		if ( types && types.preventDefault && types.handleObj ) {

			// ( event )  dispatched jQuery.Event
			handleObj = types.handleObj;
			jQuery( types.delegateTarget ).off(
				handleObj.namespace ?
					handleObj.origType + "." + handleObj.namespace :
					handleObj.origType,
				handleObj.selector,
				handleObj.handler
			);
			return this;
		}
		if ( typeof types === "object" ) {

			// ( types-object [, selector] )
			for ( type in types ) {
				this.off( type, selector, types[ type ] );
			}
			return this;
		}
		if ( selector === false || typeof selector === "function" ) {

			// ( types [, fn] )
			fn = selector;
			selector = undefined;
		}
		if ( fn === false ) {
			fn = returnFalse;
		}
		return this.each( function() {
			jQuery.event.remove( this, types, fn, selector );
		} );
	}
} );


var

	/* eslint-disable max-len */

	// See https://github.com/eslint/eslint/issues/3229
	rxhtmlTag = /<(?!area|br|col|embed|hr|img|input|link|meta|param)(([a-z][^\/\0>\x20\t\r\n\f]*)[^>]*)\/>/gi,

	/* eslint-enable */

	// Support: IE <=10 - 11, Edge 12 - 13 only
	// In IE/Edge using regex groups here causes severe slowdowns.
	// See https://connect.microsoft.com/IE/feedback/details/1736512/
	rnoInnerhtml = /<script|<style|<link/i,

	// checked="checked" or checked
	rchecked = /checked\s*(?:[^=]|=\s*.checked.)/i,
	rcleanScript = /^\s*<!(?:\[CDATA\[|--)|(?:\]\]|--)>\s*$/g;

// Prefer a tbody over its parent table for containing new rows
function manipulationTarget( elem, content ) {
	if ( nodeName( elem, "table" ) &&
		nodeName( content.nodeType !== 11 ? content : content.firstChild, "tr" ) ) {

		return jQuery( elem ).children( "tbody" )[ 0 ] || elem;
	}

	return elem;
}

// Replace/restore the type attribute of script elements for safe DOM manipulation
function disableScript( elem ) {
	elem.type = ( elem.getAttribute( "type" ) !== null ) + "/" + elem.type;
	return elem;
}
function restoreScript( elem ) {
	if ( ( elem.type || "" ).slice( 0, 5 ) === "true/" ) {
		elem.type = elem.type.slice( 5 );
	} else {
		elem.removeAttribute( "type" );
	}

	return elem;
}

function cloneCopyEvent( src, dest ) {
	var i, l, type, pdataOld, pdataCur, udataOld, udataCur, events;

	if ( dest.nodeType !== 1 ) {
		return;
	}

	// 1. Copy private data: events, handlers, etc.
	if ( dataPriv.hasData( src ) ) {
		pdataOld = dataPriv.access( src );
		pdataCur = dataPriv.set( dest, pdataOld );
		events = pdataOld.events;

		if ( events ) {
			delete pdataCur.handle;
			pdataCur.events = {};

			for ( type in events ) {
				for ( i = 0, l = events[ type ].length; i < l; i++ ) {
					jQuery.event.add( dest, type, events[ type ][ i ] );
				}
			}
		}
	}

	// 2. Copy user data
	if ( dataUser.hasData( src ) ) {
		udataOld = dataUser.access( src );
		udataCur = jQuery.extend( {}, udataOld );

		dataUser.set( dest, udataCur );
	}
}

// Fix IE bugs, see support tests
function fixInput( src, dest ) {
	var nodeName = dest.nodeName.toLowerCase();

	// Fails to persist the checked state of a cloned checkbox or radio button.
	if ( nodeName === "input" && rcheckableType.test( src.type ) ) {
		dest.checked = src.checked;

	// Fails to return the selected option to the default selected state when cloning options
	} else if ( nodeName === "input" || nodeName === "textarea" ) {
		dest.defaultValue = src.defaultValue;
	}
}

function domManip( collection, args, callback, ignored ) {

	// Flatten any nested arrays
	args = concat.apply( [], args );

	var fragment, first, scripts, hasScripts, node, doc,
		i = 0,
		l = collection.length,
		iNoClone = l - 1,
		value = args[ 0 ],
		valueIsFunction = isFunction( value );

	// We can't cloneNode fragments that contain checked, in WebKit
	if ( valueIsFunction ||
			( l > 1 && typeof value === "string" &&
				!support.checkClone && rchecked.test( value ) ) ) {
		return collection.each( function( index ) {
			var self = collection.eq( index );
			if ( valueIsFunction ) {
				args[ 0 ] = value.call( this, index, self.html() );
			}
			domManip( self, args, callback, ignored );
		} );
	}

	if ( l ) {
		fragment = buildFragment( args, collection[ 0 ].ownerDocument, false, collection, ignored );
		first = fragment.firstChild;

		if ( fragment.childNodes.length === 1 ) {
			fragment = first;
		}

		// Require either new content or an interest in ignored elements to invoke the callback
		if ( first || ignored ) {
			scripts = jQuery.map( getAll( fragment, "script" ), disableScript );
			hasScripts = scripts.length;

			// Use the original fragment for the last item
			// instead of the first because it can end up
			// being emptied incorrectly in certain situations (#8070).
			for ( ; i < l; i++ ) {
				node = fragment;

				if ( i !== iNoClone ) {
					node = jQuery.clone( node, true, true );

					// Keep references to cloned scripts for later restoration
					if ( hasScripts ) {

						// Support: Android <=4.0 only, PhantomJS 1 only
						// push.apply(_, arraylike) throws on ancient WebKit
						jQuery.merge( scripts, getAll( node, "script" ) );
					}
				}

				callback.call( collection[ i ], node, i );
			}

			if ( hasScripts ) {
				doc = scripts[ scripts.length - 1 ].ownerDocument;

				// Reenable scripts
				jQuery.map( scripts, restoreScript );

				// Evaluate executable scripts on first document insertion
				for ( i = 0; i < hasScripts; i++ ) {
					node = scripts[ i ];
					if ( rscriptType.test( node.type || "" ) &&
						!dataPriv.access( node, "globalEval" ) &&
						jQuery.contains( doc, node ) ) {

						if ( node.src && ( node.type || "" ).toLowerCase()  !== "module" ) {

							// Optional AJAX dependency, but won't run scripts if not present
							if ( jQuery._evalUrl && !node.noModule ) {
								jQuery._evalUrl( node.src, {
									nonce: node.nonce || node.getAttribute( "nonce" )
								} );
							}
						} else {
							DOMEval( node.textContent.replace( rcleanScript, "" ), node, doc );
						}
					}
				}
			}
		}
	}

	return collection;
}

function remove( elem, selector, keepData ) {
	var node,
		nodes = selector ? jQuery.filter( selector, elem ) : elem,
		i = 0;

	for ( ; ( node = nodes[ i ] ) != null; i++ ) {
		if ( !keepData && node.nodeType === 1 ) {
			jQuery.cleanData( getAll( node ) );
		}

		if ( node.parentNode ) {
			if ( keepData && isAttached( node ) ) {
				setGlobalEval( getAll( node, "script" ) );
			}
			node.parentNode.removeChild( node );
		}
	}

	return elem;
}

jQuery.extend( {
	htmlPrefilter: function( html ) {
		return html.replace( rxhtmlTag, "<$1></$2>" );
	},

	clone: function( elem, dataAndEvents, deepDataAndEvents ) {
		var i, l, srcElements, destElements,
			clone = elem.cloneNode( true ),
			inPage = isAttached( elem );

		// Fix IE cloning issues
		if ( !support.noCloneChecked && ( elem.nodeType === 1 || elem.nodeType === 11 ) &&
				!jQuery.isXMLDoc( elem ) ) {

			// We eschew Sizzle here for performance reasons: https://jsperf.com/getall-vs-sizzle/2
			destElements = getAll( clone );
			srcElements = getAll( elem );

			for ( i = 0, l = srcElements.length; i < l; i++ ) {
				fixInput( srcElements[ i ], destElements[ i ] );
			}
		}

		// Copy the events from the original to the clone
		if ( dataAndEvents ) {
			if ( deepDataAndEvents ) {
				srcElements = srcElements || getAll( elem );
				destElements = destElements || getAll( clone );

				for ( i = 0, l = srcElements.length; i < l; i++ ) {
					cloneCopyEvent( srcElements[ i ], destElements[ i ] );
				}
			} else {
				cloneCopyEvent( elem, clone );
			}
		}

		// Preserve script evaluation history
		destElements = getAll( clone, "script" );
		if ( destElements.length > 0 ) {
			setGlobalEval( destElements, !inPage && getAll( elem, "script" ) );
		}

		// Return the cloned set
		return clone;
	},

	cleanData: function( elems ) {
		var data, elem, type,
			special = jQuery.event.special,
			i = 0;

		for ( ; ( elem = elems[ i ] ) !== undefined; i++ ) {
			if ( acceptData( elem ) ) {
				if ( ( data = elem[ dataPriv.expando ] ) ) {
					if ( data.events ) {
						for ( type in data.events ) {
							if ( special[ type ] ) {
								jQuery.event.remove( elem, type );

							// This is a shortcut to avoid jQuery.event.remove's overhead
							} else {
								jQuery.removeEvent( elem, type, data.handle );
							}
						}
					}

					// Support: Chrome <=35 - 45+
					// Assign undefined instead of using delete, see Data#remove
					elem[ dataPriv.expando ] = undefined;
				}
				if ( elem[ dataUser.expando ] ) {

					// Support: Chrome <=35 - 45+
					// Assign undefined instead of using delete, see Data#remove
					elem[ dataUser.expando ] = undefined;
				}
			}
		}
	}
} );

jQuery.fn.extend( {
	detach: function( selector ) {
		return remove( this, selector, true );
	},

	remove: function( selector ) {
		return remove( this, selector );
	},

	text: function( value ) {
		return access( this, function( value ) {
			return value === undefined ?
				jQuery.text( this ) :
				this.empty().each( function() {
					if ( this.nodeType === 1 || this.nodeType === 11 || this.nodeType === 9 ) {
						this.textContent = value;
					}
				} );
		}, null, value, arguments.length );
	},

	append: function() {
		return domManip( this, arguments, function( elem ) {
			if ( this.nodeType === 1 || this.nodeType === 11 || this.nodeType === 9 ) {
				var target = manipulationTarget( this, elem );
				target.appendChild( elem );
			}
		} );
	},

	prepend: function() {
		return domManip( this, arguments, function( elem ) {
			if ( this.nodeType === 1 || this.nodeType === 11 || this.nodeType === 9 ) {
				var target = manipulationTarget( this, elem );
				target.insertBefore( elem, target.firstChild );
			}
		} );
	},

	before: function() {
		return domManip( this, arguments, function( elem ) {
			if ( this.parentNode ) {
				this.parentNode.insertBefore( elem, this );
			}
		} );
	},

	after: function() {
		return domManip( this, arguments, function( elem ) {
			if ( this.parentNode ) {
				this.parentNode.insertBefore( elem, this.nextSibling );
			}
		} );
	},

	empty: function() {
		var elem,
			i = 0;

		for ( ; ( elem = this[ i ] ) != null; i++ ) {
			if ( elem.nodeType === 1 ) {

				// Prevent memory leaks
				jQuery.cleanData( getAll( elem, false ) );

				// Remove any remaining nodes
				elem.textContent = "";
			}
		}

		return this;
	},

	clone: function( dataAndEvents, deepDataAndEvents ) {
		dataAndEvents = dataAndEvents == null ? false : dataAndEvents;
		deepDataAndEvents = deepDataAndEvents == null ? dataAndEvents : deepDataAndEvents;

		return this.map( function() {
			return jQuery.clone( this, dataAndEvents, deepDataAndEvents );
		} );
	},

	html: function( value ) {
		return access( this, function( value ) {
			var elem = this[ 0 ] || {},
				i = 0,
				l = this.length;

			if ( value === undefined && elem.nodeType === 1 ) {
				return elem.innerHTML;
			}

			// See if we can take a shortcut and just use innerHTML
			if ( typeof value === "string" && !rnoInnerhtml.test( value ) &&
				!wrapMap[ ( rtagName.exec( value ) || [ "", "" ] )[ 1 ].toLowerCase() ] ) {

				value = jQuery.htmlPrefilter( value );

				try {
					for ( ; i < l; i++ ) {
						elem = this[ i ] || {};

						// Remove element nodes and prevent memory leaks
						if ( elem.nodeType === 1 ) {
							jQuery.cleanData( getAll( elem, false ) );
							elem.innerHTML = value;
						}
					}

					elem = 0;

				// If using innerHTML throws an exception, use the fallback method
				} catch ( e ) {}
			}

			if ( elem ) {
				this.empty().append( value );
			}
		}, null, value, arguments.length );
	},

	replaceWith: function() {
		var ignored = [];

		// Make the changes, replacing each non-ignored context element with the new content
		return domManip( this, arguments, function( elem ) {
			var parent = this.parentNode;

			if ( jQuery.inArray( this, ignored ) < 0 ) {
				jQuery.cleanData( getAll( this ) );
				if ( parent ) {
					parent.replaceChild( elem, this );
				}
			}

		// Force callback invocation
		}, ignored );
	}
} );

jQuery.each( {
	appendTo: "append",
	prependTo: "prepend",
	insertBefore: "before",
	insertAfter: "after",
	replaceAll: "replaceWith"
}, function( name, original ) {
	jQuery.fn[ name ] = function( selector ) {
		var elems,
			ret = [],
			insert = jQuery( selector ),
			last = insert.length - 1,
			i = 0;

		for ( ; i <= last; i++ ) {
			elems = i === last ? this : this.clone( true );
			jQuery( insert[ i ] )[ original ]( elems );

			// Support: Android <=4.0 only, PhantomJS 1 only
			// .get() because push.apply(_, arraylike) throws on ancient WebKit
			push.apply( ret, elems.get() );
		}

		return this.pushStack( ret );
	};
} );
var rnumnonpx = new RegExp( "^(" + pnum + ")(?!px)[a-z%]+$", "i" );

var getStyles = function( elem ) {

		// Support: IE <=11 only, Firefox <=30 (#15098, #14150)
		// IE throws on elements created in popups
		// FF meanwhile throws on frame elements through "defaultView.getComputedStyle"
		var view = elem.ownerDocument.defaultView;

		if ( !view || !view.opener ) {
			view = window;
		}

		return view.getComputedStyle( elem );
	};

var rboxStyle = new RegExp( cssExpand.join( "|" ), "i" );


( function() {

	// Executing both pixelPosition & boxSizingReliable tests require only one layout
	// so they're executed at the same time to save the second computation.
	function computeStyleTests() {

		// This is a singleton, we need to execute it only once
		if ( !div ) {
			return;
		}

		container.style.cssText = "position:absolute;left:-11111px;width:60px;" +
			"margin-top:1px;padding:0;border:0";
		div.style.cssText =
			"position:relative;display:block;box-sizing:border-box;overflow:scroll;" +
			"margin:auto;border:1px;padding:1px;" +
			"width:60%;top:1%";
		documentElement.appendChild( container ).appendChild( div );

		var divStyle = window.getComputedStyle( div );
		pixelPositionVal = divStyle.top !== "1%";

		// Support: Android 4.0 - 4.3 only, Firefox <=3 - 44
		reliableMarginLeftVal = roundPixelMeasures( divStyle.marginLeft ) === 12;

		// Support: Android 4.0 - 4.3 only, Safari <=9.1 - 10.1, iOS <=7.0 - 9.3
		// Some styles come back with percentage values, even though they shouldn't
		div.style.right = "60%";
		pixelBoxStylesVal = roundPixelMeasures( divStyle.right ) === 36;

		// Support: IE 9 - 11 only
		// Detect misreporting of content dimensions for box-sizing:border-box elements
		boxSizingReliableVal = roundPixelMeasures( divStyle.width ) === 36;

		// Support: IE 9 only
		// Detect overflow:scroll screwiness (gh-3699)
		// Support: Chrome <=64
		// Don't get tricked when zoom affects offsetWidth (gh-4029)
		div.style.position = "absolute";
		scrollboxSizeVal = roundPixelMeasures( div.offsetWidth / 3 ) === 12;

		documentElement.removeChild( container );

		// Nullify the div so it wouldn't be stored in the memory and
		// it will also be a sign that checks already performed
		div = null;
	}

	function roundPixelMeasures( measure ) {
		return Math.round( parseFloat( measure ) );
	}

	var pixelPositionVal, boxSizingReliableVal, scrollboxSizeVal, pixelBoxStylesVal,
		reliableMarginLeftVal,
		container = document.createElement( "div" ),
		div = document.createElement( "div" );

	// Finish early in limited (non-browser) environments
	if ( !div.style ) {
		return;
	}

	// Support: IE <=9 - 11 only
	// Style of cloned element affects source element cloned (#8908)
	div.style.backgroundClip = "content-box";
	div.cloneNode( true ).style.backgroundClip = "";
	support.clearCloneStyle = div.style.backgroundClip === "content-box";

	jQuery.extend( support, {
		boxSizingReliable: function() {
			computeStyleTests();
			return boxSizingReliableVal;
		},
		pixelBoxStyles: function() {
			computeStyleTests();
			return pixelBoxStylesVal;
		},
		pixelPosition: function() {
			computeStyleTests();
			return pixelPositionVal;
		},
		reliableMarginLeft: function() {
			computeStyleTests();
			return reliableMarginLeftVal;
		},
		scrollboxSize: function() {
			computeStyleTests();
			return scrollboxSizeVal;
		}
	} );
} )();


function curCSS( elem, name, computed ) {
	var width, minWidth, maxWidth, ret,

		// Support: Firefox 51+
		// Retrieving style before computed somehow
		// fixes an issue with getting wrong values
		// on detached elements
		style = elem.style;

	computed = computed || getStyles( elem );

	// getPropertyValue is needed for:
	//   .css('filter') (IE 9 only, #12537)
	//   .css('--customProperty) (#3144)
	if ( computed ) {
		ret = computed.getPropertyValue( name ) || computed[ name ];

		if ( ret === "" && !isAttached( elem ) ) {
			ret = jQuery.style( elem, name );
		}

		// A tribute to the "awesome hack by Dean Edwards"
		// Android Browser returns percentage for some values,
		// but width seems to be reliably pixels.
		// This is against the CSSOM draft spec:
		// https://drafts.csswg.org/cssom/#resolved-values
		if ( !support.pixelBoxStyles() && rnumnonpx.test( ret ) && rboxStyle.test( name ) ) {

			// Remember the original values
			width = style.width;
			minWidth = style.minWidth;
			maxWidth = style.maxWidth;

			// Put in the new values to get a computed value out
			style.minWidth = style.maxWidth = style.width = ret;
			ret = computed.width;

			// Revert the changed values
			style.width = width;
			style.minWidth = minWidth;
			style.maxWidth = maxWidth;
		}
	}

	return ret !== undefined ?

		// Support: IE <=9 - 11 only
		// IE returns zIndex value as an integer.
		ret + "" :
		ret;
}


function addGetHookIf( conditionFn, hookFn ) {

	// Define the hook, we'll check on the first run if it's really needed.
	return {
		get: function() {
			if ( conditionFn() ) {

				// Hook not needed (or it's not possible to use it due
				// to missing dependency), remove it.
				delete this.get;
				return;
			}

			// Hook needed; redefine it so that the support test is not executed again.
			return ( this.get = hookFn ).apply( this, arguments );
		}
	};
}


var cssPrefixes = [ "Webkit", "Moz", "ms" ],
	emptyStyle = document.createElement( "div" ).style,
	vendorProps = {};

// Return a vendor-prefixed property or undefined
function vendorPropName( name ) {

	// Check for vendor prefixed names
	var capName = name[ 0 ].toUpperCase() + name.slice( 1 ),
		i = cssPrefixes.length;

	while ( i-- ) {
		name = cssPrefixes[ i ] + capName;
		if ( name in emptyStyle ) {
			return name;
		}
	}
}

// Return a potentially-mapped jQuery.cssProps or vendor prefixed property
function finalPropName( name ) {
	var final = jQuery.cssProps[ name ] || vendorProps[ name ];

	if ( final ) {
		return final;
	}
	if ( name in emptyStyle ) {
		return name;
	}
	return vendorProps[ name ] = vendorPropName( name ) || name;
}


var

	// Swappable if display is none or starts with table
	// except "table", "table-cell", or "table-caption"
	// See here for display values: https://developer.mozilla.org/en-US/docs/CSS/display
	rdisplayswap = /^(none|table(?!-c[ea]).+)/,
	rcustomProp = /^--/,
	cssShow = { position: "absolute", visibility: "hidden", display: "block" },
	cssNormalTransform = {
		letterSpacing: "0",
		fontWeight: "400"
	};

function setPositiveNumber( elem, value, subtract ) {

	// Any relative (+/-) values have already been
	// normalized at this point
	var matches = rcssNum.exec( value );
	return matches ?

		// Guard against undefined "subtract", e.g., when used as in cssHooks
		Math.max( 0, matches[ 2 ] - ( subtract || 0 ) ) + ( matches[ 3 ] || "px" ) :
		value;
}

function boxModelAdjustment( elem, dimension, box, isBorderBox, styles, computedVal ) {
	var i = dimension === "width" ? 1 : 0,
		extra = 0,
		delta = 0;

	// Adjustment may not be necessary
	if ( box === ( isBorderBox ? "border" : "content" ) ) {
		return 0;
	}

	for ( ; i < 4; i += 2 ) {

		// Both box models exclude margin
		if ( box === "margin" ) {
			delta += jQuery.css( elem, box + cssExpand[ i ], true, styles );
		}

		// If we get here with a content-box, we're seeking "padding" or "border" or "margin"
		if ( !isBorderBox ) {

			// Add padding
			delta += jQuery.css( elem, "padding" + cssExpand[ i ], true, styles );

			// For "border" or "margin", add border
			if ( box !== "padding" ) {
				delta += jQuery.css( elem, "border" + cssExpand[ i ] + "Width", true, styles );

			// But still keep track of it otherwise
			} else {
				extra += jQuery.css( elem, "border" + cssExpand[ i ] + "Width", true, styles );
			}

		// If we get here with a border-box (content + padding + border), we're seeking "content" or
		// "padding" or "margin"
		} else {

			// For "content", subtract padding
			if ( box === "content" ) {
				delta -= jQuery.css( elem, "padding" + cssExpand[ i ], true, styles );
			}

			// For "content" or "padding", subtract border
			if ( box !== "margin" ) {
				delta -= jQuery.css( elem, "border" + cssExpand[ i ] + "Width", true, styles );
			}
		}
	}

	// Account for positive content-box scroll gutter when requested by providing computedVal
	if ( !isBorderBox && computedVal >= 0 ) {

		// offsetWidth/offsetHeight is a rounded sum of content, padding, scroll gutter, and border
		// Assuming integer scroll gutter, subtract the rest and round down
		delta += Math.max( 0, Math.ceil(
			elem[ "offset" + dimension[ 0 ].toUpperCase() + dimension.slice( 1 ) ] -
			computedVal -
			delta -
			extra -
			0.5

		// If offsetWidth/offsetHeight is unknown, then we can't determine content-box scroll gutter
		// Use an explicit zero to avoid NaN (gh-3964)
		) ) || 0;
	}

	return delta;
}

function getWidthOrHeight( elem, dimension, extra ) {

	// Start with computed style
	var styles = getStyles( elem ),

		// To avoid forcing a reflow, only fetch boxSizing if we need it (gh-4322).
		// Fake content-box until we know it's needed to know the true value.
		boxSizingNeeded = !support.boxSizingReliable() || extra,
		isBorderBox = boxSizingNeeded &&
			jQuery.css( elem, "boxSizing", false, styles ) === "border-box",
		valueIsBorderBox = isBorderBox,

		val = curCSS( elem, dimension, styles ),
		offsetProp = "offset" + dimension[ 0 ].toUpperCase() + dimension.slice( 1 );

	// Support: Firefox <=54
	// Return a confounding non-pixel value or feign ignorance, as appropriate.
	if ( rnumnonpx.test( val ) ) {
		if ( !extra ) {
			return val;
		}
		val = "auto";
	}


	// Fall back to offsetWidth/offsetHeight when value is "auto"
	// This happens for inline elements with no explicit setting (gh-3571)
	// Support: Android <=4.1 - 4.3 only
	// Also use offsetWidth/offsetHeight for misreported inline dimensions (gh-3602)
	// Support: IE 9-11 only
	// Also use offsetWidth/offsetHeight for when box sizing is unreliable
	// We use getClientRects() to check for hidden/disconnected.
	// In those cases, the computed value can be trusted to be border-box
	if ( ( !support.boxSizingReliable() && isBorderBox ||
		val === "auto" ||
		!parseFloat( val ) && jQuery.css( elem, "display", false, styles ) === "inline" ) &&
		elem.getClientRects().length ) {

		isBorderBox = jQuery.css( elem, "boxSizing", false, styles ) === "border-box";

		// Where available, offsetWidth/offsetHeight approximate border box dimensions.
		// Where not available (e.g., SVG), assume unreliable box-sizing and interpret the
		// retrieved value as a content box dimension.
		valueIsBorderBox = offsetProp in elem;
		if ( valueIsBorderBox ) {
			val = elem[ offsetProp ];
		}
	}

	// Normalize "" and auto
	val = parseFloat( val ) || 0;

	// Adjust for the element's box model
	return ( val +
		boxModelAdjustment(
			elem,
			dimension,
			extra || ( isBorderBox ? "border" : "content" ),
			valueIsBorderBox,
			styles,

			// Provide the current computed size to request scroll gutter calculation (gh-3589)
			val
		)
	) + "px";
}

jQuery.extend( {

	// Add in style property hooks for overriding the default
	// behavior of getting and setting a style property
	cssHooks: {
		opacity: {
			get: function( elem, computed ) {
				if ( computed ) {

					// We should always get a number back from opacity
					var ret = curCSS( elem, "opacity" );
					return ret === "" ? "1" : ret;
				}
			}
		}
	},

	// Don't automatically add "px" to these possibly-unitless properties
	cssNumber: {
		"animationIterationCount": true,
		"columnCount": true,
		"fillOpacity": true,
		"flexGrow": true,
		"flexShrink": true,
		"fontWeight": true,
		"gridArea": true,
		"gridColumn": true,
		"gridColumnEnd": true,
		"gridColumnStart": true,
		"gridRow": true,
		"gridRowEnd": true,
		"gridRowStart": true,
		"lineHeight": true,
		"opacity": true,
		"order": true,
		"orphans": true,
		"widows": true,
		"zIndex": true,
		"zoom": true
	},

	// Add in properties whose names you wish to fix before
	// setting or getting the value
	cssProps: {},

	// Get and set the style property on a DOM Node
	style: function( elem, name, value, extra ) {

		// Don't set styles on text and comment nodes
		if ( !elem || elem.nodeType === 3 || elem.nodeType === 8 || !elem.style ) {
			return;
		}

		// Make sure that we're working with the right name
		var ret, type, hooks,
			origName = camelCase( name ),
			isCustomProp = rcustomProp.test( name ),
			style = elem.style;

		// Make sure that we're working with the right name. We don't
		// want to query the value if it is a CSS custom property
		// since they are user-defined.
		if ( !isCustomProp ) {
			name = finalPropName( origName );
		}

		// Gets hook for the prefixed version, then unprefixed version
		hooks = jQuery.cssHooks[ name ] || jQuery.cssHooks[ origName ];

		// Check if we're setting a value
		if ( value !== undefined ) {
			type = typeof value;

			// Convert "+=" or "-=" to relative numbers (#7345)
			if ( type === "string" && ( ret = rcssNum.exec( value ) ) && ret[ 1 ] ) {
				value = adjustCSS( elem, name, ret );

				// Fixes bug #9237
				type = "number";
			}

			// Make sure that null and NaN values aren't set (#7116)
			if ( value == null || value !== value ) {
				return;
			}

			// If a number was passed in, add the unit (except for certain CSS properties)
			// The isCustomProp check can be removed in jQuery 4.0 when we only auto-append
			// "px" to a few hardcoded values.
			if ( type === "number" && !isCustomProp ) {
				value += ret && ret[ 3 ] || ( jQuery.cssNumber[ origName ] ? "" : "px" );
			}

			// background-* props affect original clone's values
			if ( !support.clearCloneStyle && value === "" && name.indexOf( "background" ) === 0 ) {
				style[ name ] = "inherit";
			}

			// If a hook was provided, use that value, otherwise just set the specified value
			if ( !hooks || !( "set" in hooks ) ||
				( value = hooks.set( elem, value, extra ) ) !== undefined ) {

				if ( isCustomProp ) {
					style.setProperty( name, value );
				} else {
					style[ name ] = value;
				}
			}

		} else {

			// If a hook was provided get the non-computed value from there
			if ( hooks && "get" in hooks &&
				( ret = hooks.get( elem, false, extra ) ) !== undefined ) {

				return ret;
			}

			// Otherwise just get the value from the style object
			return style[ name ];
		}
	},

	css: function( elem, name, extra, styles ) {
		var val, num, hooks,
			origName = camelCase( name ),
			isCustomProp = rcustomProp.test( name );

		// Make sure that we're working with the right name. We don't
		// want to modify the value if it is a CSS custom property
		// since they are user-defined.
		if ( !isCustomProp ) {
			name = finalPropName( origName );
		}

		// Try prefixed name followed by the unprefixed name
		hooks = jQuery.cssHooks[ name ] || jQuery.cssHooks[ origName ];

		// If a hook was provided get the computed value from there
		if ( hooks && "get" in hooks ) {
			val = hooks.get( elem, true, extra );
		}

		// Otherwise, if a way to get the computed value exists, use that
		if ( val === undefined ) {
			val = curCSS( elem, name, styles );
		}

		// Convert "normal" to computed value
		if ( val === "normal" && name in cssNormalTransform ) {
			val = cssNormalTransform[ name ];
		}

		// Make numeric if forced or a qualifier was provided and val looks numeric
		if ( extra === "" || extra ) {
			num = parseFloat( val );
			return extra === true || isFinite( num ) ? num || 0 : val;
		}

		return val;
	}
} );

jQuery.each( [ "height", "width" ], function( i, dimension ) {
	jQuery.cssHooks[ dimension ] = {
		get: function( elem, computed, extra ) {
			if ( computed ) {

				// Certain elements can have dimension info if we invisibly show them
				// but it must have a current display style that would benefit
				return rdisplayswap.test( jQuery.css( elem, "display" ) ) &&

					// Support: Safari 8+
					// Table columns in Safari have non-zero offsetWidth & zero
					// getBoundingClientRect().width unless display is changed.
					// Support: IE <=11 only
					// Running getBoundingClientRect on a disconnected node
					// in IE throws an error.
					( !elem.getClientRects().length || !elem.getBoundingClientRect().width ) ?
						swap( elem, cssShow, function() {
							return getWidthOrHeight( elem, dimension, extra );
						} ) :
						getWidthOrHeight( elem, dimension, extra );
			}
		},

		set: function( elem, value, extra ) {
			var matches,
				styles = getStyles( elem ),

				// Only read styles.position if the test has a chance to fail
				// to avoid forcing a reflow.
				scrollboxSizeBuggy = !support.scrollboxSize() &&
					styles.position === "absolute",

				// To avoid forcing a reflow, only fetch boxSizing if we need it (gh-3991)
				boxSizingNeeded = scrollboxSizeBuggy || extra,
				isBorderBox = boxSizingNeeded &&
					jQuery.css( elem, "boxSizing", false, styles ) === "border-box",
				subtract = extra ?
					boxModelAdjustment(
						elem,
						dimension,
						extra,
						isBorderBox,
						styles
					) :
					0;

			// Account for unreliable border-box dimensions by comparing offset* to computed and
			// faking a content-box to get border and padding (gh-3699)
			if ( isBorderBox && scrollboxSizeBuggy ) {
				subtract -= Math.ceil(
					elem[ "offset" + dimension[ 0 ].toUpperCase() + dimension.slice( 1 ) ] -
					parseFloat( styles[ dimension ] ) -
					boxModelAdjustment( elem, dimension, "border", false, styles ) -
					0.5
				);
			}

			// Convert to pixels if value adjustment is needed
			if ( subtract && ( matches = rcssNum.exec( value ) ) &&
				( matches[ 3 ] || "px" ) !== "px" ) {

				elem.style[ dimension ] = value;
				value = jQuery.css( elem, dimension );
			}

			return setPositiveNumber( elem, value, subtract );
		}
	};
} );

jQuery.cssHooks.marginLeft = addGetHookIf( support.reliableMarginLeft,
	function( elem, computed ) {
		if ( computed ) {
			return ( parseFloat( curCSS( elem, "marginLeft" ) ) ||
				elem.getBoundingClientRect().left -
					swap( elem, { marginLeft: 0 }, function() {
						return elem.getBoundingClientRect().left;
					} )
				) + "px";
		}
	}
);

// These hooks are used by animate to expand properties
jQuery.each( {
	margin: "",
	padding: "",
	border: "Width"
}, function( prefix, suffix ) {
	jQuery.cssHooks[ prefix + suffix ] = {
		expand: function( value ) {
			var i = 0,
				expanded = {},

				// Assumes a single number if not a string
				parts = typeof value === "string" ? value.split( " " ) : [ value ];

			for ( ; i < 4; i++ ) {
				expanded[ prefix + cssExpand[ i ] + suffix ] =
					parts[ i ] || parts[ i - 2 ] || parts[ 0 ];
			}

			return expanded;
		}
	};

	if ( prefix !== "margin" ) {
		jQuery.cssHooks[ prefix + suffix ].set = setPositiveNumber;
	}
} );

jQuery.fn.extend( {
	css: function( name, value ) {
		return access( this, function( elem, name, value ) {
			var styles, len,
				map = {},
				i = 0;

			if ( Array.isArray( name ) ) {
				styles = getStyles( elem );
				len = name.length;

				for ( ; i < len; i++ ) {
					map[ name[ i ] ] = jQuery.css( elem, name[ i ], false, styles );
				}

				return map;
			}

			return value !== undefined ?
				jQuery.style( elem, name, value ) :
				jQuery.css( elem, name );
		}, name, value, arguments.length > 1 );
	}
} );


function Tween( elem, options, prop, end, easing ) {
	return new Tween.prototype.init( elem, options, prop, end, easing );
}
jQuery.Tween = Tween;

Tween.prototype = {
	constructor: Tween,
	init: function( elem, options, prop, end, easing, unit ) {
		this.elem = elem;
		this.prop = prop;
		this.easing = easing || jQuery.easing._default;
		this.options = options;
		this.start = this.now = this.cur();
		this.end = end;
		this.unit = unit || ( jQuery.cssNumber[ prop ] ? "" : "px" );
	},
	cur: function() {
		var hooks = Tween.propHooks[ this.prop ];

		return hooks && hooks.get ?
			hooks.get( this ) :
			Tween.propHooks._default.get( this );
	},
	run: function( percent ) {
		var eased,
			hooks = Tween.propHooks[ this.prop ];

		if ( this.options.duration ) {
			this.pos = eased = jQuery.easing[ this.easing ](
				percent, this.options.duration * percent, 0, 1, this.options.duration
			);
		} else {
			this.pos = eased = percent;
		}
		this.now = ( this.end - this.start ) * eased + this.start;

		if ( this.options.step ) {
			this.options.step.call( this.elem, this.now, this );
		}

		if ( hooks && hooks.set ) {
			hooks.set( this );
		} else {
			Tween.propHooks._default.set( this );
		}
		return this;
	}
};

Tween.prototype.init.prototype = Tween.prototype;

Tween.propHooks = {
	_default: {
		get: function( tween ) {
			var result;

			// Use a property on the element directly when it is not a DOM element,
			// or when there is no matching style property that exists.
			if ( tween.elem.nodeType !== 1 ||
				tween.elem[ tween.prop ] != null && tween.elem.style[ tween.prop ] == null ) {
				return tween.elem[ tween.prop ];
			}

			// Passing an empty string as a 3rd parameter to .css will automatically
			// attempt a parseFloat and fallback to a string if the parse fails.
			// Simple values such as "10px" are parsed to Float;
			// complex values such as "rotate(1rad)" are returned as-is.
			result = jQuery.css( tween.elem, tween.prop, "" );

			// Empty strings, null, undefined and "auto" are converted to 0.
			return !result || result === "auto" ? 0 : result;
		},
		set: function( tween ) {

			// Use step hook for back compat.
			// Use cssHook if its there.
			// Use .style if available and use plain properties where available.
			if ( jQuery.fx.step[ tween.prop ] ) {
				jQuery.fx.step[ tween.prop ]( tween );
			} else if ( tween.elem.nodeType === 1 && (
					jQuery.cssHooks[ tween.prop ] ||
					tween.elem.style[ finalPropName( tween.prop ) ] != null ) ) {
				jQuery.style( tween.elem, tween.prop, tween.now + tween.unit );
			} else {
				tween.elem[ tween.prop ] = tween.now;
			}
		}
	}
};

// Support: IE <=9 only
// Panic based approach to setting things on disconnected nodes
Tween.propHooks.scrollTop = Tween.propHooks.scrollLeft = {
	set: function( tween ) {
		if ( tween.elem.nodeType && tween.elem.parentNode ) {
			tween.elem[ tween.prop ] = tween.now;
		}
	}
};

jQuery.easing = {
	linear: function( p ) {
		return p;
	},
	swing: function( p ) {
		return 0.5 - Math.cos( p * Math.PI ) / 2;
	},
	_default: "swing"
};

jQuery.fx = Tween.prototype.init;

// Back compat <1.8 extension point
jQuery.fx.step = {};


var
	fxNow, inProgress,
	rfxtypes = /^(?:toggle|show|hide)$/,
	rrun = /queueHooks$/;

function schedule() {
	if ( inProgress ) {
		if ( document.hidden === false && window.requestAnimationFrame ) {
			window.requestAnimationFrame( schedule );
		} else {
			window.setTimeout( schedule, jQuery.fx.interval );
		}

		jQuery.fx.tick();
	}
}

// Animations created synchronously will run synchronously
function createFxNow() {
	window.setTimeout( function() {
		fxNow = undefined;
	} );
	return ( fxNow = Date.now() );
}

// Generate parameters to create a standard animation
function genFx( type, includeWidth ) {
	var which,
		i = 0,
		attrs = { height: type };

	// If we include width, step value is 1 to do all cssExpand values,
	// otherwise step value is 2 to skip over Left and Right
	includeWidth = includeWidth ? 1 : 0;
	for ( ; i < 4; i += 2 - includeWidth ) {
		which = cssExpand[ i ];
		attrs[ "margin" + which ] = attrs[ "padding" + which ] = type;
	}

	if ( includeWidth ) {
		attrs.opacity = attrs.width = type;
	}

	return attrs;
}

function createTween( value, prop, animation ) {
	var tween,
		collection = ( Animation.tweeners[ prop ] || [] ).concat( Animation.tweeners[ "*" ] ),
		index = 0,
		length = collection.length;
	for ( ; index < length; index++ ) {
		if ( ( tween = collection[ index ].call( animation, prop, value ) ) ) {

			// We're done with this property
			return tween;
		}
	}
}

function defaultPrefilter( elem, props, opts ) {
	var prop, value, toggle, hooks, oldfire, propTween, restoreDisplay, display,
		isBox = "width" in props || "height" in props,
		anim = this,
		orig = {},
		style = elem.style,
		hidden = elem.nodeType && isHiddenWithinTree( elem ),
		dataShow = dataPriv.get( elem, "fxshow" );

	// Queue-skipping animations hijack the fx hooks
	if ( !opts.queue ) {
		hooks = jQuery._queueHooks( elem, "fx" );
		if ( hooks.unqueued == null ) {
			hooks.unqueued = 0;
			oldfire = hooks.empty.fire;
			hooks.empty.fire = function() {
				if ( !hooks.unqueued ) {
					oldfire();
				}
			};
		}
		hooks.unqueued++;

		anim.always( function() {

			// Ensure the complete handler is called before this completes
			anim.always( function() {
				hooks.unqueued--;
				if ( !jQuery.queue( elem, "fx" ).length ) {
					hooks.empty.fire();
				}
			} );
		} );
	}

	// Detect show/hide animations
	for ( prop in props ) {
		value = props[ prop ];
		if ( rfxtypes.test( value ) ) {
			delete props[ prop ];
			toggle = toggle || value === "toggle";
			if ( value === ( hidden ? "hide" : "show" ) ) {

				// Pretend to be hidden if this is a "show" and
				// there is still data from a stopped show/hide
				if ( value === "show" && dataShow && dataShow[ prop ] !== undefined ) {
					hidden = true;

				// Ignore all other no-op show/hide data
				} else {
					continue;
				}
			}
			orig[ prop ] = dataShow && dataShow[ prop ] || jQuery.style( elem, prop );
		}
	}

	// Bail out if this is a no-op like .hide().hide()
	propTween = !jQuery.isEmptyObject( props );
	if ( !propTween && jQuery.isEmptyObject( orig ) ) {
		return;
	}

	// Restrict "overflow" and "display" styles during box animations
	if ( isBox && elem.nodeType === 1 ) {

		// Support: IE <=9 - 11, Edge 12 - 15
		// Record all 3 overflow attributes because IE does not infer the shorthand
		// from identically-valued overflowX and overflowY and Edge just mirrors
		// the overflowX value there.
		opts.overflow = [ style.overflow, style.overflowX, style.overflowY ];

		// Identify a display type, preferring old show/hide data over the CSS cascade
		restoreDisplay = dataShow && dataShow.display;
		if ( restoreDisplay == null ) {
			restoreDisplay = dataPriv.get( elem, "display" );
		}
		display = jQuery.css( elem, "display" );
		if ( display === "none" ) {
			if ( restoreDisplay ) {
				display = restoreDisplay;
			} else {

				// Get nonempty value(s) by temporarily forcing visibility
				showHide( [ elem ], true );
				restoreDisplay = elem.style.display || restoreDisplay;
				display = jQuery.css( elem, "display" );
				showHide( [ elem ] );
			}
		}

		// Animate inline elements as inline-block
		if ( display === "inline" || display === "inline-block" && restoreDisplay != null ) {
			if ( jQuery.css( elem, "float" ) === "none" ) {

				// Restore the original display value at the end of pure show/hide animations
				if ( !propTween ) {
					anim.done( function() {
						style.display = restoreDisplay;
					} );
					if ( restoreDisplay == null ) {
						display = style.display;
						restoreDisplay = display === "none" ? "" : display;
					}
				}
				style.display = "inline-block";
			}
		}
	}

	if ( opts.overflow ) {
		style.overflow = "hidden";
		anim.always( function() {
			style.overflow = opts.overflow[ 0 ];
			style.overflowX = opts.overflow[ 1 ];
			style.overflowY = opts.overflow[ 2 ];
		} );
	}

	// Implement show/hide animations
	propTween = false;
	for ( prop in orig ) {

		// General show/hide setup for this element animation
		if ( !propTween ) {
			if ( dataShow ) {
				if ( "hidden" in dataShow ) {
					hidden = dataShow.hidden;
				}
			} else {
				dataShow = dataPriv.access( elem, "fxshow", { display: restoreDisplay } );
			}

			// Store hidden/visible for toggle so `.stop().toggle()` "reverses"
			if ( toggle ) {
				dataShow.hidden = !hidden;
			}

			// Show elements before animating them
			if ( hidden ) {
				showHide( [ elem ], true );
			}

			/* eslint-disable no-loop-func */

			anim.done( function() {

			/* eslint-enable no-loop-func */

				// The final step of a "hide" animation is actually hiding the element
				if ( !hidden ) {
					showHide( [ elem ] );
				}
				dataPriv.remove( elem, "fxshow" );
				for ( prop in orig ) {
					jQuery.style( elem, prop, orig[ prop ] );
				}
			} );
		}

		// Per-property setup
		propTween = createTween( hidden ? dataShow[ prop ] : 0, prop, anim );
		if ( !( prop in dataShow ) ) {
			dataShow[ prop ] = propTween.start;
			if ( hidden ) {
				propTween.end = propTween.start;
				propTween.start = 0;
			}
		}
	}
}

function propFilter( props, specialEasing ) {
	var index, name, easing, value, hooks;

	// camelCase, specialEasing and expand cssHook pass
	for ( index in props ) {
		name = camelCase( index );
		easing = specialEasing[ name ];
		value = props[ index ];
		if ( Array.isArray( value ) ) {
			easing = value[ 1 ];
			value = props[ index ] = value[ 0 ];
		}

		if ( index !== name ) {
			props[ name ] = value;
			delete props[ index ];
		}

		hooks = jQuery.cssHooks[ name ];
		if ( hooks && "expand" in hooks ) {
			value = hooks.expand( value );
			delete props[ name ];

			// Not quite $.extend, this won't overwrite existing keys.
			// Reusing 'index' because we have the correct "name"
			for ( index in value ) {
				if ( !( index in props ) ) {
					props[ index ] = value[ index ];
					specialEasing[ index ] = easing;
				}
			}
		} else {
			specialEasing[ name ] = easing;
		}
	}
}

function Animation( elem, properties, options ) {
	var result,
		stopped,
		index = 0,
		length = Animation.prefilters.length,
		deferred = jQuery.Deferred().always( function() {

			// Don't match elem in the :animated selector
			delete tick.elem;
		} ),
		tick = function() {
			if ( stopped ) {
				return false;
			}
			var currentTime = fxNow || createFxNow(),
				remaining = Math.max( 0, animation.startTime + animation.duration - currentTime ),

				// Support: Android 2.3 only
				// Archaic crash bug won't allow us to use `1 - ( 0.5 || 0 )` (#12497)
				temp = remaining / animation.duration || 0,
				percent = 1 - temp,
				index = 0,
				length = animation.tweens.length;

			for ( ; index < length; index++ ) {
				animation.tweens[ index ].run( percent );
			}

			deferred.notifyWith( elem, [ animation, percent, remaining ] );

			// If there's more to do, yield
			if ( percent < 1 && length ) {
				return remaining;
			}

			// If this was an empty animation, synthesize a final progress notification
			if ( !length ) {
				deferred.notifyWith( elem, [ animation, 1, 0 ] );
			}

			// Resolve the animation and report its conclusion
			deferred.resolveWith( elem, [ animation ] );
			return false;
		},
		animation = deferred.promise( {
			elem: elem,
			props: jQuery.extend( {}, properties ),
			opts: jQuery.extend( true, {
				specialEasing: {},
				easing: jQuery.easing._default
			}, options ),
			originalProperties: properties,
			originalOptions: options,
			startTime: fxNow || createFxNow(),
			duration: options.duration,
			tweens: [],
			createTween: function( prop, end ) {
				var tween = jQuery.Tween( elem, animation.opts, prop, end,
						animation.opts.specialEasing[ prop ] || animation.opts.easing );
				animation.tweens.push( tween );
				return tween;
			},
			stop: function( gotoEnd ) {
				var index = 0,

					// If we are going to the end, we want to run all the tweens
					// otherwise we skip this part
					length = gotoEnd ? animation.tweens.length : 0;
				if ( stopped ) {
					return this;
				}
				stopped = true;
				for ( ; index < length; index++ ) {
					animation.tweens[ index ].run( 1 );
				}

				// Resolve when we played the last frame; otherwise, reject
				if ( gotoEnd ) {
					deferred.notifyWith( elem, [ animation, 1, 0 ] );
					deferred.resolveWith( elem, [ animation, gotoEnd ] );
				} else {
					deferred.rejectWith( elem, [ animation, gotoEnd ] );
				}
				return this;
			}
		} ),
		props = animation.props;

	propFilter( props, animation.opts.specialEasing );

	for ( ; index < length; index++ ) {
		result = Animation.prefilters[ index ].call( animation, elem, props, animation.opts );
		if ( result ) {
			if ( isFunction( result.stop ) ) {
				jQuery._queueHooks( animation.elem, animation.opts.queue ).stop =
					result.stop.bind( result );
			}
			return result;
		}
	}

	jQuery.map( props, createTween, animation );

	if ( isFunction( animation.opts.start ) ) {
		animation.opts.start.call( elem, animation );
	}

	// Attach callbacks from options
	animation
		.progress( animation.opts.progress )
		.done( animation.opts.done, animation.opts.complete )
		.fail( animation.opts.fail )
		.always( animation.opts.always );

	jQuery.fx.timer(
		jQuery.extend( tick, {
			elem: elem,
			anim: animation,
			queue: animation.opts.queue
		} )
	);

	return animation;
}

jQuery.Animation = jQuery.extend( Animation, {

	tweeners: {
		"*": [ function( prop, value ) {
			var tween = this.createTween( prop, value );
			adjustCSS( tween.elem, prop, rcssNum.exec( value ), tween );
			return tween;
		} ]
	},

	tweener: function( props, callback ) {
		if ( isFunction( props ) ) {
			callback = props;
			props = [ "*" ];
		} else {
			props = props.match( rnothtmlwhite );
		}

		var prop,
			index = 0,
			length = props.length;

		for ( ; index < length; index++ ) {
			prop = props[ index ];
			Animation.tweeners[ prop ] = Animation.tweeners[ prop ] || [];
			Animation.tweeners[ prop ].unshift( callback );
		}
	},

	prefilters: [ defaultPrefilter ],

	prefilter: function( callback, prepend ) {
		if ( prepend ) {
			Animation.prefilters.unshift( callback );
		} else {
			Animation.prefilters.push( callback );
		}
	}
} );

jQuery.speed = function( speed, easing, fn ) {
	var opt = speed && typeof speed === "object" ? jQuery.extend( {}, speed ) : {
		complete: fn || !fn && easing ||
			isFunction( speed ) && speed,
		duration: speed,
		easing: fn && easing || easing && !isFunction( easing ) && easing
	};

	// Go to the end state if fx are off
	if ( jQuery.fx.off ) {
		opt.duration = 0;

	} else {
		if ( typeof opt.duration !== "number" ) {
			if ( opt.duration in jQuery.fx.speeds ) {
				opt.duration = jQuery.fx.speeds[ opt.duration ];

			} else {
				opt.duration = jQuery.fx.speeds._default;
			}
		}
	}

	// Normalize opt.queue - true/undefined/null -> "fx"
	if ( opt.queue == null || opt.queue === true ) {
		opt.queue = "fx";
	}

	// Queueing
	opt.old = opt.complete;

	opt.complete = function() {
		if ( isFunction( opt.old ) ) {
			opt.old.call( this );
		}

		if ( opt.queue ) {
			jQuery.dequeue( this, opt.queue );
		}
	};

	return opt;
};

jQuery.fn.extend( {
	fadeTo: function( speed, to, easing, callback ) {

		// Show any hidden elements after setting opacity to 0
		return this.filter( isHiddenWithinTree ).css( "opacity", 0 ).show()

			// Animate to the value specified
			.end().animate( { opacity: to }, speed, easing, callback );
	},
	animate: function( prop, speed, easing, callback ) {
		var empty = jQuery.isEmptyObject( prop ),
			optall = jQuery.speed( speed, easing, callback ),
			doAnimation = function() {

				// Operate on a copy of prop so per-property easing won't be lost
				var anim = Animation( this, jQuery.extend( {}, prop ), optall );

				// Empty animations, or finishing resolves immediately
				if ( empty || dataPriv.get( this, "finish" ) ) {
					anim.stop( true );
				}
			};
			doAnimation.finish = doAnimation;

		return empty || optall.queue === false ?
			this.each( doAnimation ) :
			this.queue( optall.queue, doAnimation );
	},
	stop: function( type, clearQueue, gotoEnd ) {
		var stopQueue = function( hooks ) {
			var stop = hooks.stop;
			delete hooks.stop;
			stop( gotoEnd );
		};

		if ( typeof type !== "string" ) {
			gotoEnd = clearQueue;
			clearQueue = type;
			type = undefined;
		}
		if ( clearQueue && type !== false ) {
			this.queue( type || "fx", [] );
		}

		return this.each( function() {
			var dequeue = true,
				index = type != null && type + "queueHooks",
				timers = jQuery.timers,
				data = dataPriv.get( this );

			if ( index ) {
				if ( data[ index ] && data[ index ].stop ) {
					stopQueue( data[ index ] );
				}
			} else {
				for ( index in data ) {
					if ( data[ index ] && data[ index ].stop && rrun.test( index ) ) {
						stopQueue( data[ index ] );
					}
				}
			}

			for ( index = timers.length; index--; ) {
				if ( timers[ index ].elem === this &&
					( type == null || timers[ index ].queue === type ) ) {

					timers[ index ].anim.stop( gotoEnd );
					dequeue = false;
					timers.splice( index, 1 );
				}
			}

			// Start the next in the queue if the last step wasn't forced.
			// Timers currently will call their complete callbacks, which
			// will dequeue but only if they were gotoEnd.
			if ( dequeue || !gotoEnd ) {
				jQuery.dequeue( this, type );
			}
		} );
	},
	finish: function( type ) {
		if ( type !== false ) {
			type = type || "fx";
		}
		return this.each( function() {
			var index,
				data = dataPriv.get( this ),
				queue = data[ type + "queue" ],
				hooks = data[ type + "queueHooks" ],
				timers = jQuery.timers,
				length = queue ? queue.length : 0;

			// Enable finishing flag on private data
			data.finish = true;

			// Empty the queue first
			jQuery.queue( this, type, [] );

			if ( hooks && hooks.stop ) {
				hooks.stop.call( this, true );
			}

			// Look for any active animations, and finish them
			for ( index = timers.length; index--; ) {
				if ( timers[ index ].elem === this && timers[ index ].queue === type ) {
					timers[ index ].anim.stop( true );
					timers.splice( index, 1 );
				}
			}

			// Look for any animations in the old queue and finish them
			for ( index = 0; index < length; index++ ) {
				if ( queue[ index ] && queue[ index ].finish ) {
					queue[ index ].finish.call( this );
				}
			}

			// Turn off finishing flag
			delete data.finish;
		} );
	}
} );

jQuery.each( [ "toggle", "show", "hide" ], function( i, name ) {
	var cssFn = jQuery.fn[ name ];
	jQuery.fn[ name ] = function( speed, easing, callback ) {
		return speed == null || typeof speed === "boolean" ?
			cssFn.apply( this, arguments ) :
			this.animate( genFx( name, true ), speed, easing, callback );
	};
} );

// Generate shortcuts for custom animations
jQuery.each( {
	slideDown: genFx( "show" ),
	slideUp: genFx( "hide" ),
	slideToggle: genFx( "toggle" ),
	fadeIn: { opacity: "show" },
	fadeOut: { opacity: "hide" },
	fadeToggle: { opacity: "toggle" }
}, function( name, props ) {
	jQuery.fn[ name ] = function( speed, easing, callback ) {
		return this.animate( props, speed, easing, callback );
	};
} );

jQuery.timers = [];
jQuery.fx.tick = function() {
	var timer,
		i = 0,
		timers = jQuery.timers;

	fxNow = Date.now();

	for ( ; i < timers.length; i++ ) {
		timer = timers[ i ];

		// Run the timer and safely remove it when done (allowing for external removal)
		if ( !timer() && timers[ i ] === timer ) {
			timers.splice( i--, 1 );
		}
	}

	if ( !timers.length ) {
		jQuery.fx.stop();
	}
	fxNow = undefined;
};

jQuery.fx.timer = function( timer ) {
	jQuery.timers.push( timer );
	jQuery.fx.start();
};

jQuery.fx.interval = 13;
jQuery.fx.start = function() {
	if ( inProgress ) {
		return;
	}

	inProgress = true;
	schedule();
};

jQuery.fx.stop = function() {
	inProgress = null;
};

jQuery.fx.speeds = {
	slow: 600,
	fast: 200,

	// Default speed
	_default: 400
};


// Based off of the plugin by Clint Helfers, with permission.
// https://web.archive.org/web/20100324014747/http://blindsignals.com/index.php/2009/07/jquery-delay/
jQuery.fn.delay = function( time, type ) {
	time = jQuery.fx ? jQuery.fx.speeds[ time ] || time : time;
	type = type || "fx";

	return this.queue( type, function( next, hooks ) {
		var timeout = window.setTimeout( next, time );
		hooks.stop = function() {
			window.clearTimeout( timeout );
		};
	} );
};


( function() {
	var input = document.createElement( "input" ),
		select = document.createElement( "select" ),
		opt = select.appendChild( document.createElement( "option" ) );

	input.type = "checkbox";

	// Support: Android <=4.3 only
	// Default value for a checkbox should be "on"
	support.checkOn = input.value !== "";

	// Support: IE <=11 only
	// Must access selectedIndex to make default options select
	support.optSelected = opt.selected;

	// Support: IE <=11 only
	// An input loses its value after becoming a radio
	input = document.createElement( "input" );
	input.value = "t";
	input.type = "radio";
	support.radioValue = input.value === "t";
} )();


var boolHook,
	attrHandle = jQuery.expr.attrHandle;

jQuery.fn.extend( {
	attr: function( name, value ) {
		return access( this, jQuery.attr, name, value, arguments.length > 1 );
	},

	removeAttr: function( name ) {
		return this.each( function() {
			jQuery.removeAttr( this, name );
		} );
	}
} );

jQuery.extend( {
	attr: function( elem, name, value ) {
		var ret, hooks,
			nType = elem.nodeType;

		// Don't get/set attributes on text, comment and attribute nodes
		if ( nType === 3 || nType === 8 || nType === 2 ) {
			return;
		}

		// Fallback to prop when attributes are not supported
		if ( typeof elem.getAttribute === "undefined" ) {
			return jQuery.prop( elem, name, value );
		}

		// Attribute hooks are determined by the lowercase version
		// Grab necessary hook if one is defined
		if ( nType !== 1 || !jQuery.isXMLDoc( elem ) ) {
			hooks = jQuery.attrHooks[ name.toLowerCase() ] ||
				( jQuery.expr.match.bool.test( name ) ? boolHook : undefined );
		}

		if ( value !== undefined ) {
			if ( value === null ) {
				jQuery.removeAttr( elem, name );
				return;
			}

			if ( hooks && "set" in hooks &&
				( ret = hooks.set( elem, value, name ) ) !== undefined ) {
				return ret;
			}

			elem.setAttribute( name, value + "" );
			return value;
		}

		if ( hooks && "get" in hooks && ( ret = hooks.get( elem, name ) ) !== null ) {
			return ret;
		}

		ret = jQuery.find.attr( elem, name );

		// Non-existent attributes return null, we normalize to undefined
		return ret == null ? undefined : ret;
	},

	attrHooks: {
		type: {
			set: function( elem, value ) {
				if ( !support.radioValue && value === "radio" &&
					nodeName( elem, "input" ) ) {
					var val = elem.value;
					elem.setAttribute( "type", value );
					if ( val ) {
						elem.value = val;
					}
					return value;
				}
			}
		}
	},

	removeAttr: function( elem, value ) {
		var name,
			i = 0,

			// Attribute names can contain non-HTML whitespace characters
			// https://html.spec.whatwg.org/multipage/syntax.html#attributes-2
			attrNames = value && value.match( rnothtmlwhite );

		if ( attrNames && elem.nodeType === 1 ) {
			while ( ( name = attrNames[ i++ ] ) ) {
				elem.removeAttribute( name );
			}
		}
	}
} );

// Hooks for boolean attributes
boolHook = {
	set: function( elem, value, name ) {
		if ( value === false ) {

			// Remove boolean attributes when set to false
			jQuery.removeAttr( elem, name );
		} else {
			elem.setAttribute( name, name );
		}
		return name;
	}
};

jQuery.each( jQuery.expr.match.bool.source.match( /\w+/g ), function( i, name ) {
	var getter = attrHandle[ name ] || jQuery.find.attr;

	attrHandle[ name ] = function( elem, name, isXML ) {
		var ret, handle,
			lowercaseName = name.toLowerCase();

		if ( !isXML ) {

			// Avoid an infinite loop by temporarily removing this function from the getter
			handle = attrHandle[ lowercaseName ];
			attrHandle[ lowercaseName ] = ret;
			ret = getter( elem, name, isXML ) != null ?
				lowercaseName :
				null;
			attrHandle[ lowercaseName ] = handle;
		}
		return ret;
	};
} );


var rfocusable = /^(?:input|select|textarea|button)$/i,
	rclickable = /^(?:a|area)$/i;

jQuery.fn.extend( {
	prop: function( name, value ) {
		return access( this, jQuery.prop, name, value, arguments.length > 1 );
	},

	removeProp: function( name ) {
		return this.each( function() {
			delete this[ jQuery.propFix[ name ] || name ];
		} );
	}
} );

jQuery.extend( {
	prop: function( elem, name, value ) {
		var ret, hooks,
			nType = elem.nodeType;

		// Don't get/set properties on text, comment and attribute nodes
		if ( nType === 3 || nType === 8 || nType === 2 ) {
			return;
		}

		if ( nType !== 1 || !jQuery.isXMLDoc( elem ) ) {

			// Fix name and attach hooks
			name = jQuery.propFix[ name ] || name;
			hooks = jQuery.propHooks[ name ];
		}

		if ( value !== undefined ) {
			if ( hooks && "set" in hooks &&
				( ret = hooks.set( elem, value, name ) ) !== undefined ) {
				return ret;
			}

			return ( elem[ name ] = value );
		}

		if ( hooks && "get" in hooks && ( ret = hooks.get( elem, name ) ) !== null ) {
			return ret;
		}

		return elem[ name ];
	},

	propHooks: {
		tabIndex: {
			get: function( elem ) {

				// Support: IE <=9 - 11 only
				// elem.tabIndex doesn't always return the
				// correct value when it hasn't been explicitly set
				// https://web.archive.org/web/20141116233347/http://fluidproject.org/blog/2008/01/09/getting-setting-and-removing-tabindex-values-with-javascript/
				// Use proper attribute retrieval(#12072)
				var tabindex = jQuery.find.attr( elem, "tabindex" );

				if ( tabindex ) {
					return parseInt( tabindex, 10 );
				}

				if (
					rfocusable.test( elem.nodeName ) ||
					rclickable.test( elem.nodeName ) &&
					elem.href
				) {
					return 0;
				}

				return -1;
			}
		}
	},

	propFix: {
		"for": "htmlFor",
		"class": "className"
	}
} );

// Support: IE <=11 only
// Accessing the selectedIndex property
// forces the browser to respect setting selected
// on the option
// The getter ensures a default option is selected
// when in an optgroup
// eslint rule "no-unused-expressions" is disabled for this code
// since it considers such accessions noop
if ( !support.optSelected ) {
	jQuery.propHooks.selected = {
		get: function( elem ) {

			/* eslint no-unused-expressions: "off" */

			var parent = elem.parentNode;
			if ( parent && parent.parentNode ) {
				parent.parentNode.selectedIndex;
			}
			return null;
		},
		set: function( elem ) {

			/* eslint no-unused-expressions: "off" */

			var parent = elem.parentNode;
			if ( parent ) {
				parent.selectedIndex;

				if ( parent.parentNode ) {
					parent.parentNode.selectedIndex;
				}
			}
		}
	};
}

jQuery.each( [
	"tabIndex",
	"readOnly",
	"maxLength",
	"cellSpacing",
	"cellPadding",
	"rowSpan",
	"colSpan",
	"useMap",
	"frameBorder",
	"contentEditable"
], function() {
	jQuery.propFix[ this.toLowerCase() ] = this;
} );


	// Strip and collapse whitespace according to HTML spec
	// https://infra.spec.whatwg.org/#strip-and-collapse-ascii-whitespace
	function stripAndCollapse( value ) {
		var tokens = value.match( rnothtmlwhite ) || [];
		return tokens.join( " " );
	}


function getClass( elem ) {
	return elem.getAttribute && elem.getAttribute( "class" ) || "";
}

function classesToArray( value ) {
	if ( Array.isArray( value ) ) {
		return value;
	}
	if ( typeof value === "string" ) {
		return value.match( rnothtmlwhite ) || [];
	}
	return [];
}

jQuery.fn.extend( {
	addClass: function( value ) {
		var classes, elem, cur, curValue, clazz, j, finalValue,
			i = 0;

		if ( isFunction( value ) ) {
			return this.each( function( j ) {
				jQuery( this ).addClass( value.call( this, j, getClass( this ) ) );
			} );
		}

		classes = classesToArray( value );

		if ( classes.length ) {
			while ( ( elem = this[ i++ ] ) ) {
				curValue = getClass( elem );
				cur = elem.nodeType === 1 && ( " " + stripAndCollapse( curValue ) + " " );

				if ( cur ) {
					j = 0;
					while ( ( clazz = classes[ j++ ] ) ) {
						if ( cur.indexOf( " " + clazz + " " ) < 0 ) {
							cur += clazz + " ";
						}
					}

					// Only assign if different to avoid unneeded rendering.
					finalValue = stripAndCollapse( cur );
					if ( curValue !== finalValue ) {
						elem.setAttribute( "class", finalValue );
					}
				}
			}
		}

		return this;
	},

	removeClass: function( value ) {
		var classes, elem, cur, curValue, clazz, j, finalValue,
			i = 0;

		if ( isFunction( value ) ) {
			return this.each( function( j ) {
				jQuery( this ).removeClass( value.call( this, j, getClass( this ) ) );
			} );
		}

		if ( !arguments.length ) {
			return this.attr( "class", "" );
		}

		classes = classesToArray( value );

		if ( classes.length ) {
			while ( ( elem = this[ i++ ] ) ) {
				curValue = getClass( elem );

				// This expression is here for better compressibility (see addClass)
				cur = elem.nodeType === 1 && ( " " + stripAndCollapse( curValue ) + " " );

				if ( cur ) {
					j = 0;
					while ( ( clazz = classes[ j++ ] ) ) {

						// Remove *all* instances
						while ( cur.indexOf( " " + clazz + " " ) > -1 ) {
							cur = cur.replace( " " + clazz + " ", " " );
						}
					}

					// Only assign if different to avoid unneeded rendering.
					finalValue = stripAndCollapse( cur );
					if ( curValue !== finalValue ) {
						elem.setAttribute( "class", finalValue );
					}
				}
			}
		}

		return this;
	},

	toggleClass: function( value, stateVal ) {
		var type = typeof value,
			isValidValue = type === "string" || Array.isArray( value );

		if ( typeof stateVal === "boolean" && isValidValue ) {
			return stateVal ? this.addClass( value ) : this.removeClass( value );
		}

		if ( isFunction( value ) ) {
			return this.each( function( i ) {
				jQuery( this ).toggleClass(
					value.call( this, i, getClass( this ), stateVal ),
					stateVal
				);
			} );
		}

		return this.each( function() {
			var className, i, self, classNames;

			if ( isValidValue ) {

				// Toggle individual class names
				i = 0;
				self = jQuery( this );
				classNames = classesToArray( value );

				while ( ( className = classNames[ i++ ] ) ) {

					// Check each className given, space separated list
					if ( self.hasClass( className ) ) {
						self.removeClass( className );
					} else {
						self.addClass( className );
					}
				}

			// Toggle whole class name
			} else if ( value === undefined || type === "boolean" ) {
				className = getClass( this );
				if ( className ) {

					// Store className if set
					dataPriv.set( this, "__className__", className );
				}

				// If the element has a class name or if we're passed `false`,
				// then remove the whole classname (if there was one, the above saved it).
				// Otherwise bring back whatever was previously saved (if anything),
				// falling back to the empty string if nothing was stored.
				if ( this.setAttribute ) {
					this.setAttribute( "class",
						className || value === false ?
						"" :
						dataPriv.get( this, "__className__" ) || ""
					);
				}
			}
		} );
	},

	hasClass: function( selector ) {
		var className, elem,
			i = 0;

		className = " " + selector + " ";
		while ( ( elem = this[ i++ ] ) ) {
			if ( elem.nodeType === 1 &&
				( " " + stripAndCollapse( getClass( elem ) ) + " " ).indexOf( className ) > -1 ) {
					return true;
			}
		}

		return false;
	}
} );


var rreturn = /\r/g;

jQuery.fn.extend( {
	val: function( value ) {
		var hooks, ret, valueIsFunction,
			elem = this[ 0 ];

		if ( !arguments.length ) {
			if ( elem ) {
				hooks = jQuery.valHooks[ elem.type ] ||
					jQuery.valHooks[ elem.nodeName.toLowerCase() ];

				if ( hooks &&
					"get" in hooks &&
					( ret = hooks.get( elem, "value" ) ) !== undefined
				) {
					return ret;
				}

				ret = elem.value;

				// Handle most common string cases
				if ( typeof ret === "string" ) {
					return ret.replace( rreturn, "" );
				}

				// Handle cases where value is null/undef or number
				return ret == null ? "" : ret;
			}

			return;
		}

		valueIsFunction = isFunction( value );

		return this.each( function( i ) {
			var val;

			if ( this.nodeType !== 1 ) {
				return;
			}

			if ( valueIsFunction ) {
				val = value.call( this, i, jQuery( this ).val() );
			} else {
				val = value;
			}

			// Treat null/undefined as ""; convert numbers to string
			if ( val == null ) {
				val = "";

			} else if ( typeof val === "number" ) {
				val += "";

			} else if ( Array.isArray( val ) ) {
				val = jQuery.map( val, function( value ) {
					return value == null ? "" : value + "";
				} );
			}

			hooks = jQuery.valHooks[ this.type ] || jQuery.valHooks[ this.nodeName.toLowerCase() ];

			// If set returns undefined, fall back to normal setting
			if ( !hooks || !( "set" in hooks ) || hooks.set( this, val, "value" ) === undefined ) {
				this.value = val;
			}
		} );
	}
} );

jQuery.extend( {
	valHooks: {
		option: {
			get: function( elem ) {

				var val = jQuery.find.attr( elem, "value" );
				return val != null ?
					val :

					// Support: IE <=10 - 11 only
					// option.text throws exceptions (#14686, #14858)
					// Strip and collapse whitespace
					// https://html.spec.whatwg.org/#strip-and-collapse-whitespace
					stripAndCollapse( jQuery.text( elem ) );
			}
		},
		select: {
			get: function( elem ) {
				var value, option, i,
					options = elem.options,
					index = elem.selectedIndex,
					one = elem.type === "select-one",
					values = one ? null : [],
					max = one ? index + 1 : options.length;

				if ( index < 0 ) {
					i = max;

				} else {
					i = one ? index : 0;
				}

				// Loop through all the selected options
				for ( ; i < max; i++ ) {
					option = options[ i ];

					// Support: IE <=9 only
					// IE8-9 doesn't update selected after form reset (#2551)
					if ( ( option.selected || i === index ) &&

							// Don't return options that are disabled or in a disabled optgroup
							!option.disabled &&
							( !option.parentNode.disabled ||
								!nodeName( option.parentNode, "optgroup" ) ) ) {

						// Get the specific value for the option
						value = jQuery( option ).val();

						// We don't need an array for one selects
						if ( one ) {
							return value;
						}

						// Multi-Selects return an array
						values.push( value );
					}
				}

				return values;
			},

			set: function( elem, value ) {
				var optionSet, option,
					options = elem.options,
					values = jQuery.makeArray( value ),
					i = options.length;

				while ( i-- ) {
					option = options[ i ];

					/* eslint-disable no-cond-assign */

					if ( option.selected =
						jQuery.inArray( jQuery.valHooks.option.get( option ), values ) > -1
					) {
						optionSet = true;
					}

					/* eslint-enable no-cond-assign */
				}

				// Force browsers to behave consistently when non-matching value is set
				if ( !optionSet ) {
					elem.selectedIndex = -1;
				}
				return values;
			}
		}
	}
} );

// Radios and checkboxes getter/setter
jQuery.each( [ "radio", "checkbox" ], function() {
	jQuery.valHooks[ this ] = {
		set: function( elem, value ) {
			if ( Array.isArray( value ) ) {
				return ( elem.checked = jQuery.inArray( jQuery( elem ).val(), value ) > -1 );
			}
		}
	};
	if ( !support.checkOn ) {
		jQuery.valHooks[ this ].get = function( elem ) {
			return elem.getAttribute( "value" ) === null ? "on" : elem.value;
		};
	}
} );


// Return jQuery for attributes-only inclusion


support.focusin = "onfocusin" in window;


var rfocusMorph = /^(?:focusinfocus|focusoutblur)$/,
	stopPropagationCallback = function( e ) {
		e.stopPropagation();
	};

jQuery.extend( jQuery.event, {

	trigger: function( event, data, elem, onlyHandlers ) {

		var i, cur, tmp, bubbleType, ontype, handle, special, lastElement,
			eventPath = [ elem || document ],
			type = hasOwn.call( event, "type" ) ? event.type : event,
			namespaces = hasOwn.call( event, "namespace" ) ? event.namespace.split( "." ) : [];

		cur = lastElement = tmp = elem = elem || document;

		// Don't do events on text and comment nodes
		if ( elem.nodeType === 3 || elem.nodeType === 8 ) {
			return;
		}

		// focus/blur morphs to focusin/out; ensure we're not firing them right now
		if ( rfocusMorph.test( type + jQuery.event.triggered ) ) {
			return;
		}

		if ( type.indexOf( "." ) > -1 ) {

			// Namespaced trigger; create a regexp to match event type in handle()
			namespaces = type.split( "." );
			type = namespaces.shift();
			namespaces.sort();
		}
		ontype = type.indexOf( ":" ) < 0 && "on" + type;

		// Caller can pass in a jQuery.Event object, Object, or just an event type string
		event = event[ jQuery.expando ] ?
			event :
			new jQuery.Event( type, typeof event === "object" && event );

		// Trigger bitmask: & 1 for native handlers; & 2 for jQuery (always true)
		event.isTrigger = onlyHandlers ? 2 : 3;
		event.namespace = namespaces.join( "." );
		event.rnamespace = event.namespace ?
			new RegExp( "(^|\\.)" + namespaces.join( "\\.(?:.*\\.|)" ) + "(\\.|$)" ) :
			null;

		// Clean up the event in case it is being reused
		event.result = undefined;
		if ( !event.target ) {
			event.target = elem;
		}

		// Clone any incoming data and prepend the event, creating the handler arg list
		data = data == null ?
			[ event ] :
			jQuery.makeArray( data, [ event ] );

		// Allow special events to draw outside the lines
		special = jQuery.event.special[ type ] || {};
		if ( !onlyHandlers && special.trigger && special.trigger.apply( elem, data ) === false ) {
			return;
		}

		// Determine event propagation path in advance, per W3C events spec (#9951)
		// Bubble up to document, then to window; watch for a global ownerDocument var (#9724)
		if ( !onlyHandlers && !special.noBubble && !isWindow( elem ) ) {

			bubbleType = special.delegateType || type;
			if ( !rfocusMorph.test( bubbleType + type ) ) {
				cur = cur.parentNode;
			}
			for ( ; cur; cur = cur.parentNode ) {
				eventPath.push( cur );
				tmp = cur;
			}

			// Only add window if we got to document (e.g., not plain obj or detached DOM)
			if ( tmp === ( elem.ownerDocument || document ) ) {
				eventPath.push( tmp.defaultView || tmp.parentWindow || window );
			}
		}

		// Fire handlers on the event path
		i = 0;
		while ( ( cur = eventPath[ i++ ] ) && !event.isPropagationStopped() ) {
			lastElement = cur;
			event.type = i > 1 ?
				bubbleType :
				special.bindType || type;

			// jQuery handler
			handle = ( dataPriv.get( cur, "events" ) || {} )[ event.type ] &&
				dataPriv.get( cur, "handle" );
			if ( handle ) {
				handle.apply( cur, data );
			}

			// Native handler
			handle = ontype && cur[ ontype ];
			if ( handle && handle.apply && acceptData( cur ) ) {
				event.result = handle.apply( cur, data );
				if ( event.result === false ) {
					event.preventDefault();
				}
			}
		}
		event.type = type;

		// If nobody prevented the default action, do it now
		if ( !onlyHandlers && !event.isDefaultPrevented() ) {

			if ( ( !special._default ||
				special._default.apply( eventPath.pop(), data ) === false ) &&
				acceptData( elem ) ) {

				// Call a native DOM method on the target with the same name as the event.
				// Don't do default actions on window, that's where global variables be (#6170)
				if ( ontype && isFunction( elem[ type ] ) && !isWindow( elem ) ) {

					// Don't re-trigger an onFOO event when we call its FOO() method
					tmp = elem[ ontype ];

					if ( tmp ) {
						elem[ ontype ] = null;
					}

					// Prevent re-triggering of the same event, since we already bubbled it above
					jQuery.event.triggered = type;

					if ( event.isPropagationStopped() ) {
						lastElement.addEventListener( type, stopPropagationCallback );
					}

					elem[ type ]();

					if ( event.isPropagationStopped() ) {
						lastElement.removeEventListener( type, stopPropagationCallback );
					}

					jQuery.event.triggered = undefined;

					if ( tmp ) {
						elem[ ontype ] = tmp;
					}
				}
			}
		}

		return event.result;
	},

	// Piggyback on a donor event to simulate a different one
	// Used only for `focus(in | out)` events
	simulate: function( type, elem, event ) {
		var e = jQuery.extend(
			new jQuery.Event(),
			event,
			{
				type: type,
				isSimulated: true
			}
		);

		jQuery.event.trigger( e, null, elem );
	}

} );

jQuery.fn.extend( {

	trigger: function( type, data ) {
		return this.each( function() {
			jQuery.event.trigger( type, data, this );
		} );
	},
	triggerHandler: function( type, data ) {
		var elem = this[ 0 ];
		if ( elem ) {
			return jQuery.event.trigger( type, data, elem, true );
		}
	}
} );


// Support: Firefox <=44
// Firefox doesn't have focus(in | out) events
// Related ticket - https://bugzilla.mozilla.org/show_bug.cgi?id=687787
//
// Support: Chrome <=48 - 49, Safari <=9.0 - 9.1
// focus(in | out) events fire after focus & blur events,
// which is spec violation - http://www.w3.org/TR/DOM-Level-3-Events/#events-focusevent-event-order
// Related ticket - https://bugs.chromium.org/p/chromium/issues/detail?id=449857
if ( !support.focusin ) {
	jQuery.each( { focus: "focusin", blur: "focusout" }, function( orig, fix ) {

		// Attach a single capturing handler on the document while someone wants focusin/focusout
		var handler = function( event ) {
			jQuery.event.simulate( fix, event.target, jQuery.event.fix( event ) );
		};

		jQuery.event.special[ fix ] = {
			setup: function() {
				var doc = this.ownerDocument || this,
					attaches = dataPriv.access( doc, fix );

				if ( !attaches ) {
					doc.addEventListener( orig, handler, true );
				}
				dataPriv.access( doc, fix, ( attaches || 0 ) + 1 );
			},
			teardown: function() {
				var doc = this.ownerDocument || this,
					attaches = dataPriv.access( doc, fix ) - 1;

				if ( !attaches ) {
					doc.removeEventListener( orig, handler, true );
					dataPriv.remove( doc, fix );

				} else {
					dataPriv.access( doc, fix, attaches );
				}
			}
		};
	} );
}
var location = window.location;

var nonce = Date.now();

var rquery = ( /\?/ );


// Cross-browser xml parsing
jQuery.parseXML = function( data ) {
	var xml;
	if ( !data || typeof data !== "string" ) {
		return null;
	}

	// Support: IE 9 - 11 only
	// IE throws on parseFromString with invalid input.
	try {
		xml = ( new window.DOMParser() ).parseFromString( data, "text/xml" );
	} catch ( e ) {
		xml = undefined;
	}

	if ( !xml || xml.getElementsByTagName( "parsererror" ).length ) {
		jQuery.error( "Invalid XML: " + data );
	}
	return xml;
};


var
	rbracket = /\[\]$/,
	rCRLF = /\r?\n/g,
	rsubmitterTypes = /^(?:submit|button|image|reset|file)$/i,
	rsubmittable = /^(?:input|select|textarea|keygen)/i;

function buildParams( prefix, obj, traditional, add ) {
	var name;

	if ( Array.isArray( obj ) ) {

		// Serialize array item.
		jQuery.each( obj, function( i, v ) {
			if ( traditional || rbracket.test( prefix ) ) {

				// Treat each array item as a scalar.
				add( prefix, v );

			} else {

				// Item is non-scalar (array or object), encode its numeric index.
				buildParams(
					prefix + "[" + ( typeof v === "object" && v != null ? i : "" ) + "]",
					v,
					traditional,
					add
				);
			}
		} );

	} else if ( !traditional && toType( obj ) === "object" ) {

		// Serialize object item.
		for ( name in obj ) {
			buildParams( prefix + "[" + name + "]", obj[ name ], traditional, add );
		}

	} else {

		// Serialize scalar item.
		add( prefix, obj );
	}
}

// Serialize an array of form elements or a set of
// key/values into a query string
jQuery.param = function( a, traditional ) {
	var prefix,
		s = [],
		add = function( key, valueOrFunction ) {

			// If value is a function, invoke it and use its return value
			var value = isFunction( valueOrFunction ) ?
				valueOrFunction() :
				valueOrFunction;

			s[ s.length ] = encodeURIComponent( key ) + "=" +
				encodeURIComponent( value == null ? "" : value );
		};

	if ( a == null ) {
		return "";
	}

	// If an array was passed in, assume that it is an array of form elements.
	if ( Array.isArray( a ) || ( a.jquery && !jQuery.isPlainObject( a ) ) ) {

		// Serialize the form elements
		jQuery.each( a, function() {
			add( this.name, this.value );
		} );

	} else {

		// If traditional, encode the "old" way (the way 1.3.2 or older
		// did it), otherwise encode params recursively.
		for ( prefix in a ) {
			buildParams( prefix, a[ prefix ], traditional, add );
		}
	}

	// Return the resulting serialization
	return s.join( "&" );
};

jQuery.fn.extend( {
	serialize: function() {
		return jQuery.param( this.serializeArray() );
	},
	serializeArray: function() {
		return this.map( function() {

			// Can add propHook for "elements" to filter or add form elements
			var elements = jQuery.prop( this, "elements" );
			return elements ? jQuery.makeArray( elements ) : this;
		} )
		.filter( function() {
			var type = this.type;

			// Use .is( ":disabled" ) so that fieldset[disabled] works
			return this.name && !jQuery( this ).is( ":disabled" ) &&
				rsubmittable.test( this.nodeName ) && !rsubmitterTypes.test( type ) &&
				( this.checked || !rcheckableType.test( type ) );
		} )
		.map( function( i, elem ) {
			var val = jQuery( this ).val();

			if ( val == null ) {
				return null;
			}

			if ( Array.isArray( val ) ) {
				return jQuery.map( val, function( val ) {
					return { name: elem.name, value: val.replace( rCRLF, "\r\n" ) };
				} );
			}

			return { name: elem.name, value: val.replace( rCRLF, "\r\n" ) };
		} ).get();
	}
} );


var
	r20 = /%20/g,
	rhash = /#.*$/,
	rantiCache = /([?&])_=[^&]*/,
	rheaders = /^(.*?):[ \t]*([^\r\n]*)$/mg,

	// #7653, #8125, #8152: local protocol detection
	rlocalProtocol = /^(?:about|app|app-storage|.+-extension|file|res|widget):$/,
	rnoContent = /^(?:GET|HEAD)$/,
	rprotocol = /^\/\//,

	/* Prefilters
	 * 1) They are useful to introduce custom dataTypes (see ajax/jsonp.js for an example)
	 * 2) These are called:
	 *    - BEFORE asking for a transport
	 *    - AFTER param serialization (s.data is a string if s.processData is true)
	 * 3) key is the dataType
	 * 4) the catchall symbol "*" can be used
	 * 5) execution will start with transport dataType and THEN continue down to "*" if needed
	 */
	prefilters = {},

	/* Transports bindings
	 * 1) key is the dataType
	 * 2) the catchall symbol "*" can be used
	 * 3) selection will start with transport dataType and THEN go to "*" if needed
	 */
	transports = {},

	// Avoid comment-prolog char sequence (#10098); must appease lint and evade compression
	allTypes = "*/".concat( "*" ),

	// Anchor tag for parsing the document origin
	originAnchor = document.createElement( "a" );
	originAnchor.href = location.href;

// Base "constructor" for jQuery.ajaxPrefilter and jQuery.ajaxTransport
function addToPrefiltersOrTransports( structure ) {

	// dataTypeExpression is optional and defaults to "*"
	return function( dataTypeExpression, func ) {

		if ( typeof dataTypeExpression !== "string" ) {
			func = dataTypeExpression;
			dataTypeExpression = "*";
		}

		var dataType,
			i = 0,
			dataTypes = dataTypeExpression.toLowerCase().match( rnothtmlwhite ) || [];

		if ( isFunction( func ) ) {

			// For each dataType in the dataTypeExpression
			while ( ( dataType = dataTypes[ i++ ] ) ) {

				// Prepend if requested
				if ( dataType[ 0 ] === "+" ) {
					dataType = dataType.slice( 1 ) || "*";
					( structure[ dataType ] = structure[ dataType ] || [] ).unshift( func );

				// Otherwise append
				} else {
					( structure[ dataType ] = structure[ dataType ] || [] ).push( func );
				}
			}
		}
	};
}

// Base inspection function for prefilters and transports
function inspectPrefiltersOrTransports( structure, options, originalOptions, jqXHR ) {

	var inspected = {},
		seekingTransport = ( structure === transports );

	function inspect( dataType ) {
		var selected;
		inspected[ dataType ] = true;
		jQuery.each( structure[ dataType ] || [], function( _, prefilterOrFactory ) {
			var dataTypeOrTransport = prefilterOrFactory( options, originalOptions, jqXHR );
			if ( typeof dataTypeOrTransport === "string" &&
				!seekingTransport && !inspected[ dataTypeOrTransport ] ) {

				options.dataTypes.unshift( dataTypeOrTransport );
				inspect( dataTypeOrTransport );
				return false;
			} else if ( seekingTransport ) {
				return !( selected = dataTypeOrTransport );
			}
		} );
		return selected;
	}

	return inspect( options.dataTypes[ 0 ] ) || !inspected[ "*" ] && inspect( "*" );
}

// A special extend for ajax options
// that takes "flat" options (not to be deep extended)
// Fixes #9887
function ajaxExtend( target, src ) {
	var key, deep,
		flatOptions = jQuery.ajaxSettings.flatOptions || {};

	for ( key in src ) {
		if ( src[ key ] !== undefined ) {
			( flatOptions[ key ] ? target : ( deep || ( deep = {} ) ) )[ key ] = src[ key ];
		}
	}
	if ( deep ) {
		jQuery.extend( true, target, deep );
	}

	return target;
}

/* Handles responses to an ajax request:
 * - finds the right dataType (mediates between content-type and expected dataType)
 * - returns the corresponding response
 */
function ajaxHandleResponses( s, jqXHR, responses ) {

	var ct, type, finalDataType, firstDataType,
		contents = s.contents,
		dataTypes = s.dataTypes;

	// Remove auto dataType and get content-type in the process
	while ( dataTypes[ 0 ] === "*" ) {
		dataTypes.shift();
		if ( ct === undefined ) {
			ct = s.mimeType || jqXHR.getResponseHeader( "Content-Type" );
		}
	}

	// Check if we're dealing with a known content-type
	if ( ct ) {
		for ( type in contents ) {
			if ( contents[ type ] && contents[ type ].test( ct ) ) {
				dataTypes.unshift( type );
				break;
			}
		}
	}

	// Check to see if we have a response for the expected dataType
	if ( dataTypes[ 0 ] in responses ) {
		finalDataType = dataTypes[ 0 ];
	} else {

		// Try convertible dataTypes
		for ( type in responses ) {
			if ( !dataTypes[ 0 ] || s.converters[ type + " " + dataTypes[ 0 ] ] ) {
				finalDataType = type;
				break;
			}
			if ( !firstDataType ) {
				firstDataType = type;
			}
		}

		// Or just use first one
		finalDataType = finalDataType || firstDataType;
	}

	// If we found a dataType
	// We add the dataType to the list if needed
	// and return the corresponding response
	if ( finalDataType ) {
		if ( finalDataType !== dataTypes[ 0 ] ) {
			dataTypes.unshift( finalDataType );
		}
		return responses[ finalDataType ];
	}
}

/* Chain conversions given the request and the original response
 * Also sets the responseXXX fields on the jqXHR instance
 */
function ajaxConvert( s, response, jqXHR, isSuccess ) {
	var conv2, current, conv, tmp, prev,
		converters = {},

		// Work with a copy of dataTypes in case we need to modify it for conversion
		dataTypes = s.dataTypes.slice();

	// Create converters map with lowercased keys
	if ( dataTypes[ 1 ] ) {
		for ( conv in s.converters ) {
			converters[ conv.toLowerCase() ] = s.converters[ conv ];
		}
	}

	current = dataTypes.shift();

	// Convert to each sequential dataType
	while ( current ) {

		if ( s.responseFields[ current ] ) {
			jqXHR[ s.responseFields[ current ] ] = response;
		}

		// Apply the dataFilter if provided
		if ( !prev && isSuccess && s.dataFilter ) {
			response = s.dataFilter( response, s.dataType );
		}

		prev = current;
		current = dataTypes.shift();

		if ( current ) {

			// There's only work to do if current dataType is non-auto
			if ( current === "*" ) {

				current = prev;

			// Convert response if prev dataType is non-auto and differs from current
			} else if ( prev !== "*" && prev !== current ) {

				// Seek a direct converter
				conv = converters[ prev + " " + current ] || converters[ "* " + current ];

				// If none found, seek a pair
				if ( !conv ) {
					for ( conv2 in converters ) {

						// If conv2 outputs current
						tmp = conv2.split( " " );
						if ( tmp[ 1 ] === current ) {

							// If prev can be converted to accepted input
							conv = converters[ prev + " " + tmp[ 0 ] ] ||
								converters[ "* " + tmp[ 0 ] ];
							if ( conv ) {

								// Condense equivalence converters
								if ( conv === true ) {
									conv = converters[ conv2 ];

								// Otherwise, insert the intermediate dataType
								} else if ( converters[ conv2 ] !== true ) {
									current = tmp[ 0 ];
									dataTypes.unshift( tmp[ 1 ] );
								}
								break;
							}
						}
					}
				}

				// Apply converter (if not an equivalence)
				if ( conv !== true ) {

					// Unless errors are allowed to bubble, catch and return them
					if ( conv && s.throws ) {
						response = conv( response );
					} else {
						try {
							response = conv( response );
						} catch ( e ) {
							return {
								state: "parsererror",
								error: conv ? e : "No conversion from " + prev + " to " + current
							};
						}
					}
				}
			}
		}
	}

	return { state: "success", data: response };
}

jQuery.extend( {

	// Counter for holding the number of active queries
	active: 0,

	// Last-Modified header cache for next request
	lastModified: {},
	etag: {},

	ajaxSettings: {
		url: location.href,
		type: "GET",
		isLocal: rlocalProtocol.test( location.protocol ),
		global: true,
		processData: true,
		async: true,
		contentType: "application/x-www-form-urlencoded; charset=UTF-8",

		/*
		timeout: 0,
		data: null,
		dataType: null,
		username: null,
		password: null,
		cache: null,
		throws: false,
		traditional: false,
		headers: {},
		*/

		accepts: {
			"*": allTypes,
			text: "text/plain",
			html: "text/html",
			xml: "application/xml, text/xml",
			json: "application/json, text/javascript"
		},

		contents: {
			xml: /\bxml\b/,
			html: /\bhtml/,
			json: /\bjson\b/
		},

		responseFields: {
			xml: "responseXML",
			text: "responseText",
			json: "responseJSON"
		},

		// Data converters
		// Keys separate source (or catchall "*") and destination types with a single space
		converters: {

			// Convert anything to text
			"* text": String,

			// Text to html (true = no transformation)
			"text html": true,

			// Evaluate text as a json expression
			"text json": JSON.parse,

			// Parse text as xml
			"text xml": jQuery.parseXML
		},

		// For options that shouldn't be deep extended:
		// you can add your own custom options here if
		// and when you create one that shouldn't be
		// deep extended (see ajaxExtend)
		flatOptions: {
			url: true,
			context: true
		}
	},

	// Creates a full fledged settings object into target
	// with both ajaxSettings and settings fields.
	// If target is omitted, writes into ajaxSettings.
	ajaxSetup: function( target, settings ) {
		return settings ?

			// Building a settings object
			ajaxExtend( ajaxExtend( target, jQuery.ajaxSettings ), settings ) :

			// Extending ajaxSettings
			ajaxExtend( jQuery.ajaxSettings, target );
	},

	ajaxPrefilter: addToPrefiltersOrTransports( prefilters ),
	ajaxTransport: addToPrefiltersOrTransports( transports ),

	// Main method
	ajax: function( url, options ) {

		// If url is an object, simulate pre-1.5 signature
		if ( typeof url === "object" ) {
			options = url;
			url = undefined;
		}

		// Force options to be an object
		options = options || {};

		var transport,

			// URL without anti-cache param
			cacheURL,

			// Response headers
			responseHeadersString,
			responseHeaders,

			// timeout handle
			timeoutTimer,

			// Url cleanup var
			urlAnchor,

			// Request state (becomes false upon send and true upon completion)
			completed,

			// To know if global events are to be dispatched
			fireGlobals,

			// Loop variable
			i,

			// uncached part of the url
			uncached,

			// Create the final options object
			s = jQuery.ajaxSetup( {}, options ),

			// Callbacks context
			callbackContext = s.context || s,

			// Context for global events is callbackContext if it is a DOM node or jQuery collection
			globalEventContext = s.context &&
				( callbackContext.nodeType || callbackContext.jquery ) ?
					jQuery( callbackContext ) :
					jQuery.event,

			// Deferreds
			deferred = jQuery.Deferred(),
			completeDeferred = jQuery.Callbacks( "once memory" ),

			// Status-dependent callbacks
			statusCode = s.statusCode || {},

			// Headers (they are sent all at once)
			requestHeaders = {},
			requestHeadersNames = {},

			// Default abort message
			strAbort = "canceled",

			// Fake xhr
			jqXHR = {
				readyState: 0,

				// Builds headers hashtable if needed
				getResponseHeader: function( key ) {
					var match;
					if ( completed ) {
						if ( !responseHeaders ) {
							responseHeaders = {};
							while ( ( match = rheaders.exec( responseHeadersString ) ) ) {
								responseHeaders[ match[ 1 ].toLowerCase() + " " ] =
									( responseHeaders[ match[ 1 ].toLowerCase() + " " ] || [] )
										.concat( match[ 2 ] );
							}
						}
						match = responseHeaders[ key.toLowerCase() + " " ];
					}
					return match == null ? null : match.join( ", " );
				},

				// Raw string
				getAllResponseHeaders: function() {
					return completed ? responseHeadersString : null;
				},

				// Caches the header
				setRequestHeader: function( name, value ) {
					if ( completed == null ) {
						name = requestHeadersNames[ name.toLowerCase() ] =
							requestHeadersNames[ name.toLowerCase() ] || name;
						requestHeaders[ name ] = value;
					}
					return this;
				},

				// Overrides response content-type header
				overrideMimeType: function( type ) {
					if ( completed == null ) {
						s.mimeType = type;
					}
					return this;
				},

				// Status-dependent callbacks
				statusCode: function( map ) {
					var code;
					if ( map ) {
						if ( completed ) {

							// Execute the appropriate callbacks
							jqXHR.always( map[ jqXHR.status ] );
						} else {

							// Lazy-add the new callbacks in a way that preserves old ones
							for ( code in map ) {
								statusCode[ code ] = [ statusCode[ code ], map[ code ] ];
							}
						}
					}
					return this;
				},

				// Cancel the request
				abort: function( statusText ) {
					var finalText = statusText || strAbort;
					if ( transport ) {
						transport.abort( finalText );
					}
					done( 0, finalText );
					return this;
				}
			};

		// Attach deferreds
		deferred.promise( jqXHR );

		// Add protocol if not provided (prefilters might expect it)
		// Handle falsy url in the settings object (#10093: consistency with old signature)
		// We also use the url parameter if available
		s.url = ( ( url || s.url || location.href ) + "" )
			.replace( rprotocol, location.protocol + "//" );

		// Alias method option to type as per ticket #12004
		s.type = options.method || options.type || s.method || s.type;

		// Extract dataTypes list
		s.dataTypes = ( s.dataType || "*" ).toLowerCase().match( rnothtmlwhite ) || [ "" ];

		// A cross-domain request is in order when the origin doesn't match the current origin.
		if ( s.crossDomain == null ) {
			urlAnchor = document.createElement( "a" );

			// Support: IE <=8 - 11, Edge 12 - 15
			// IE throws exception on accessing the href property if url is malformed,
			// e.g. http://example.com:80x/
			try {
				urlAnchor.href = s.url;

				// Support: IE <=8 - 11 only
				// Anchor's host property isn't correctly set when s.url is relative
				urlAnchor.href = urlAnchor.href;
				s.crossDomain = originAnchor.protocol + "//" + originAnchor.host !==
					urlAnchor.protocol + "//" + urlAnchor.host;
			} catch ( e ) {

				// If there is an error parsing the URL, assume it is crossDomain,
				// it can be rejected by the transport if it is invalid
				s.crossDomain = true;
			}
		}

		// Convert data if not already a string
		if ( s.data && s.processData && typeof s.data !== "string" ) {
			s.data = jQuery.param( s.data, s.traditional );
		}

		// Apply prefilters
		inspectPrefiltersOrTransports( prefilters, s, options, jqXHR );

		// If request was aborted inside a prefilter, stop there
		if ( completed ) {
			return jqXHR;
		}

		// We can fire global events as of now if asked to
		// Don't fire events if jQuery.event is undefined in an AMD-usage scenario (#15118)
		fireGlobals = jQuery.event && s.global;

		// Watch for a new set of requests
		if ( fireGlobals && jQuery.active++ === 0 ) {
			jQuery.event.trigger( "ajaxStart" );
		}

		// Uppercase the type
		s.type = s.type.toUpperCase();

		// Determine if request has content
		s.hasContent = !rnoContent.test( s.type );

		// Save the URL in case we're toying with the If-Modified-Since
		// and/or If-None-Match header later on
		// Remove hash to simplify url manipulation
		cacheURL = s.url.replace( rhash, "" );

		// More options handling for requests with no content
		if ( !s.hasContent ) {

			// Remember the hash so we can put it back
			uncached = s.url.slice( cacheURL.length );

			// If data is available and should be processed, append data to url
			if ( s.data && ( s.processData || typeof s.data === "string" ) ) {
				cacheURL += ( rquery.test( cacheURL ) ? "&" : "?" ) + s.data;

				// #9682: remove data so that it's not used in an eventual retry
				delete s.data;
			}

			// Add or update anti-cache param if needed
			if ( s.cache === false ) {
				cacheURL = cacheURL.replace( rantiCache, "$1" );
				uncached = ( rquery.test( cacheURL ) ? "&" : "?" ) + "_=" + ( nonce++ ) + uncached;
			}

			// Put hash and anti-cache on the URL that will be requested (gh-1732)
			s.url = cacheURL + uncached;

		// Change '%20' to '+' if this is encoded form body content (gh-2658)
		} else if ( s.data && s.processData &&
			( s.contentType || "" ).indexOf( "application/x-www-form-urlencoded" ) === 0 ) {
			s.data = s.data.replace( r20, "+" );
		}

		// Set the If-Modified-Since and/or If-None-Match header, if in ifModified mode.
		if ( s.ifModified ) {
			if ( jQuery.lastModified[ cacheURL ] ) {
				jqXHR.setRequestHeader( "If-Modified-Since", jQuery.lastModified[ cacheURL ] );
			}
			if ( jQuery.etag[ cacheURL ] ) {
				jqXHR.setRequestHeader( "If-None-Match", jQuery.etag[ cacheURL ] );
			}
		}

		// Set the correct header, if data is being sent
		if ( s.data && s.hasContent && s.contentType !== false || options.contentType ) {
			jqXHR.setRequestHeader( "Content-Type", s.contentType );
		}

		// Set the Accepts header for the server, depending on the dataType
		jqXHR.setRequestHeader(
			"Accept",
			s.dataTypes[ 0 ] && s.accepts[ s.dataTypes[ 0 ] ] ?
				s.accepts[ s.dataTypes[ 0 ] ] +
					( s.dataTypes[ 0 ] !== "*" ? ", " + allTypes + "; q=0.01" : "" ) :
				s.accepts[ "*" ]
		);

		// Check for headers option
		for ( i in s.headers ) {
			jqXHR.setRequestHeader( i, s.headers[ i ] );
		}

		// Allow custom headers/mimetypes and early abort
		if ( s.beforeSend &&
			( s.beforeSend.call( callbackContext, jqXHR, s ) === false || completed ) ) {

			// Abort if not done already and return
			return jqXHR.abort();
		}

		// Aborting is no longer a cancellation
		strAbort = "abort";

		// Install callbacks on deferreds
		completeDeferred.add( s.complete );
		jqXHR.done( s.success );
		jqXHR.fail( s.error );

		// Get transport
		transport = inspectPrefiltersOrTransports( transports, s, options, jqXHR );

		// If no transport, we auto-abort
		if ( !transport ) {
			done( -1, "No Transport" );
		} else {
			jqXHR.readyState = 1;

			// Send global event
			if ( fireGlobals ) {
				globalEventContext.trigger( "ajaxSend", [ jqXHR, s ] );
			}

			// If request was aborted inside ajaxSend, stop there
			if ( completed ) {
				return jqXHR;
			}

			// Timeout
			if ( s.async && s.timeout > 0 ) {
				timeoutTimer = window.setTimeout( function() {
					jqXHR.abort( "timeout" );
				}, s.timeout );
			}

			try {
				completed = false;
				transport.send( requestHeaders, done );
			} catch ( e ) {

				// Rethrow post-completion exceptions
				if ( completed ) {
					throw e;
				}

				// Propagate others as results
				done( -1, e );
			}
		}

		// Callback for when everything is done
		function done( status, nativeStatusText, responses, headers ) {
			var isSuccess, success, error, response, modified,
				statusText = nativeStatusText;

			// Ignore repeat invocations
			if ( completed ) {
				return;
			}

			completed = true;

			// Clear timeout if it exists
			if ( timeoutTimer ) {
				window.clearTimeout( timeoutTimer );
			}

			// Dereference transport for early garbage collection
			// (no matter how long the jqXHR object will be used)
			transport = undefined;

			// Cache response headers
			responseHeadersString = headers || "";

			// Set readyState
			jqXHR.readyState = status > 0 ? 4 : 0;

			// Determine if successful
			isSuccess = status >= 200 && status < 300 || status === 304;

			// Get response data
			if ( responses ) {
				response = ajaxHandleResponses( s, jqXHR, responses );
			}

			// Convert no matter what (that way responseXXX fields are always set)
			response = ajaxConvert( s, response, jqXHR, isSuccess );

			// If successful, handle type chaining
			if ( isSuccess ) {

				// Set the If-Modified-Since and/or If-None-Match header, if in ifModified mode.
				if ( s.ifModified ) {
					modified = jqXHR.getResponseHeader( "Last-Modified" );
					if ( modified ) {
						jQuery.lastModified[ cacheURL ] = modified;
					}
					modified = jqXHR.getResponseHeader( "etag" );
					if ( modified ) {
						jQuery.etag[ cacheURL ] = modified;
					}
				}

				// if no content
				if ( status === 204 || s.type === "HEAD" ) {
					statusText = "nocontent";

				// if not modified
				} else if ( status === 304 ) {
					statusText = "notmodified";

				// If we have data, let's convert it
				} else {
					statusText = response.state;
					success = response.data;
					error = response.error;
					isSuccess = !error;
				}
			} else {

				// Extract error from statusText and normalize for non-aborts
				error = statusText;
				if ( status || !statusText ) {
					statusText = "error";
					if ( status < 0 ) {
						status = 0;
					}
				}
			}

			// Set data for the fake xhr object
			jqXHR.status = status;
			jqXHR.statusText = ( nativeStatusText || statusText ) + "";

			// Success/Error
			if ( isSuccess ) {
				deferred.resolveWith( callbackContext, [ success, statusText, jqXHR ] );
			} else {
				deferred.rejectWith( callbackContext, [ jqXHR, statusText, error ] );
			}

			// Status-dependent callbacks
			jqXHR.statusCode( statusCode );
			statusCode = undefined;

			if ( fireGlobals ) {
				globalEventContext.trigger( isSuccess ? "ajaxSuccess" : "ajaxError",
					[ jqXHR, s, isSuccess ? success : error ] );
			}

			// Complete
			completeDeferred.fireWith( callbackContext, [ jqXHR, statusText ] );

			if ( fireGlobals ) {
				globalEventContext.trigger( "ajaxComplete", [ jqXHR, s ] );

				// Handle the global AJAX counter
				if ( !( --jQuery.active ) ) {
					jQuery.event.trigger( "ajaxStop" );
				}
			}
		}

		return jqXHR;
	},

	getJSON: function( url, data, callback ) {
		return jQuery.get( url, data, callback, "json" );
	},

	getScript: function( url, callback ) {
		return jQuery.get( url, undefined, callback, "script" );
	}
} );

jQuery.each( [ "get", "post" ], function( i, method ) {
	jQuery[ method ] = function( url, data, callback, type ) {

		// Shift arguments if data argument was omitted
		if ( isFunction( data ) ) {
			type = type || callback;
			callback = data;
			data = undefined;
		}

		// The url can be an options object (which then must have .url)
		return jQuery.ajax( jQuery.extend( {
			url: url,
			type: method,
			dataType: type,
			data: data,
			success: callback
		}, jQuery.isPlainObject( url ) && url ) );
	};
} );


jQuery._evalUrl = function( url, options ) {
	return jQuery.ajax( {
		url: url,

		// Make this explicit, since user can override this through ajaxSetup (#11264)
		type: "GET",
		dataType: "script",
		cache: true,
		async: false,
		global: false,

		// Only evaluate the response if it is successful (gh-4126)
		// dataFilter is not invoked for failure responses, so using it instead
		// of the default converter is kludgy but it works.
		converters: {
			"text script": function() {}
		},
		dataFilter: function( response ) {
			jQuery.globalEval( response, options );
		}
	} );
};


jQuery.fn.extend( {
	wrapAll: function( html ) {
		var wrap;

		if ( this[ 0 ] ) {
			if ( isFunction( html ) ) {
				html = html.call( this[ 0 ] );
			}

			// The elements to wrap the target around
			wrap = jQuery( html, this[ 0 ].ownerDocument ).eq( 0 ).clone( true );

			if ( this[ 0 ].parentNode ) {
				wrap.insertBefore( this[ 0 ] );
			}

			wrap.map( function() {
				var elem = this;

				while ( elem.firstElementChild ) {
					elem = elem.firstElementChild;
				}

				return elem;
			} ).append( this );
		}

		return this;
	},

	wrapInner: function( html ) {
		if ( isFunction( html ) ) {
			return this.each( function( i ) {
				jQuery( this ).wrapInner( html.call( this, i ) );
			} );
		}

		return this.each( function() {
			var self = jQuery( this ),
				contents = self.contents();

			if ( contents.length ) {
				contents.wrapAll( html );

			} else {
				self.append( html );
			}
		} );
	},

	wrap: function( html ) {
		var htmlIsFunction = isFunction( html );

		return this.each( function( i ) {
			jQuery( this ).wrapAll( htmlIsFunction ? html.call( this, i ) : html );
		} );
	},

	unwrap: function( selector ) {
		this.parent( selector ).not( "body" ).each( function() {
			jQuery( this ).replaceWith( this.childNodes );
		} );
		return this;
	}
} );


jQuery.expr.pseudos.hidden = function( elem ) {
	return !jQuery.expr.pseudos.visible( elem );
};
jQuery.expr.pseudos.visible = function( elem ) {
	return !!( elem.offsetWidth || elem.offsetHeight || elem.getClientRects().length );
};


jQuery.ajaxSettings.xhr = function() {
	try {
		return new window.XMLHttpRequest();
	} catch ( e ) {}
};

var xhrSuccessStatus = {

		// File protocol always yields status code 0, assume 200
		0: 200,

		// Support: IE <=9 only
		// #1450: sometimes IE returns 1223 when it should be 204
		1223: 204
	},
	xhrSupported = jQuery.ajaxSettings.xhr();

support.cors = !!xhrSupported && ( "withCredentials" in xhrSupported );
support.ajax = xhrSupported = !!xhrSupported;

jQuery.ajaxTransport( function( options ) {
	var callback, errorCallback;

	// Cross domain only allowed if supported through XMLHttpRequest
	if ( support.cors || xhrSupported && !options.crossDomain ) {
		return {
			send: function( headers, complete ) {
				var i,
					xhr = options.xhr();

				xhr.open(
					options.type,
					options.url,
					options.async,
					options.username,
					options.password
				);

				// Apply custom fields if provided
				if ( options.xhrFields ) {
					for ( i in options.xhrFields ) {
						xhr[ i ] = options.xhrFields[ i ];
					}
				}

				// Override mime type if needed
				if ( options.mimeType && xhr.overrideMimeType ) {
					xhr.overrideMimeType( options.mimeType );
				}

				// X-Requested-With header
				// For cross-domain requests, seeing as conditions for a preflight are
				// akin to a jigsaw puzzle, we simply never set it to be sure.
				// (it can always be set on a per-request basis or even using ajaxSetup)
				// For same-domain requests, won't change header if already provided.
				if ( !options.crossDomain && !headers[ "X-Requested-With" ] ) {
					headers[ "X-Requested-With" ] = "XMLHttpRequest";
				}

				// Set headers
				for ( i in headers ) {
					xhr.setRequestHeader( i, headers[ i ] );
				}

				// Callback
				callback = function( type ) {
					return function() {
						if ( callback ) {
							callback = errorCallback = xhr.onload =
								xhr.onerror = xhr.onabort = xhr.ontimeout =
									xhr.onreadystatechange = null;

							if ( type === "abort" ) {
								xhr.abort();
							} else if ( type === "error" ) {

								// Support: IE <=9 only
								// On a manual native abort, IE9 throws
								// errors on any property access that is not readyState
								if ( typeof xhr.status !== "number" ) {
									complete( 0, "error" );
								} else {
									complete(

										// File: protocol always yields status 0; see #8605, #14207
										xhr.status,
										xhr.statusText
									);
								}
							} else {
								complete(
									xhrSuccessStatus[ xhr.status ] || xhr.status,
									xhr.statusText,

									// Support: IE <=9 only
									// IE9 has no XHR2 but throws on binary (trac-11426)
									// For XHR2 non-text, let the caller handle it (gh-2498)
									( xhr.responseType || "text" ) !== "text"  ||
									typeof xhr.responseText !== "string" ?
										{ binary: xhr.response } :
										{ text: xhr.responseText },
									xhr.getAllResponseHeaders()
								);
							}
						}
					};
				};

				// Listen to events
				xhr.onload = callback();
				errorCallback = xhr.onerror = xhr.ontimeout = callback( "error" );

				// Support: IE 9 only
				// Use onreadystatechange to replace onabort
				// to handle uncaught aborts
				if ( xhr.onabort !== undefined ) {
					xhr.onabort = errorCallback;
				} else {
					xhr.onreadystatechange = function() {

						// Check readyState before timeout as it changes
						if ( xhr.readyState === 4 ) {

							// Allow onerror to be called first,
							// but that will not handle a native abort
							// Also, save errorCallback to a variable
							// as xhr.onerror cannot be accessed
							window.setTimeout( function() {
								if ( callback ) {
									errorCallback();
								}
							} );
						}
					};
				}

				// Create the abort callback
				callback = callback( "abort" );

				try {

					// Do send the request (this may raise an exception)
					xhr.send( options.hasContent && options.data || null );
				} catch ( e ) {

					// #14683: Only rethrow if this hasn't been notified as an error yet
					if ( callback ) {
						throw e;
					}
				}
			},

			abort: function() {
				if ( callback ) {
					callback();
				}
			}
		};
	}
} );


// Prevent auto-execution of scripts when no explicit dataType was provided (See gh-2432)
jQuery.ajaxPrefilter( function( s ) {
	if ( s.crossDomain ) {
		s.contents.script = false;
	}
} );

// Install script dataType
jQuery.ajaxSetup( {
	accepts: {
		script: "text/javascript, application/javascript, " +
			"application/ecmascript, application/x-ecmascript"
	},
	contents: {
		script: /\b(?:java|ecma)script\b/
	},
	converters: {
		"text script": function( text ) {
			jQuery.globalEval( text );
			return text;
		}
	}
} );

// Handle cache's special case and crossDomain
jQuery.ajaxPrefilter( "script", function( s ) {
	if ( s.cache === undefined ) {
		s.cache = false;
	}
	if ( s.crossDomain ) {
		s.type = "GET";
	}
} );

// Bind script tag hack transport
jQuery.ajaxTransport( "script", function( s ) {

	// This transport only deals with cross domain or forced-by-attrs requests
	if ( s.crossDomain || s.scriptAttrs ) {
		var script, callback;
		return {
			send: function( _, complete ) {
				script = jQuery( "<script>" )
					.attr( s.scriptAttrs || {} )
					.prop( { charset: s.scriptCharset, src: s.url } )
					.on( "load error", callback = function( evt ) {
						script.remove();
						callback = null;
						if ( evt ) {
							complete( evt.type === "error" ? 404 : 200, evt.type );
						}
					} );

				// Use native DOM manipulation to avoid our domManip AJAX trickery
				document.head.appendChild( script[ 0 ] );
			},
			abort: function() {
				if ( callback ) {
					callback();
				}
			}
		};
	}
} );


var oldCallbacks = [],
	rjsonp = /(=)\?(?=&|$)|\?\?/;

// Default jsonp settings
jQuery.ajaxSetup( {
	jsonp: "callback",
	jsonpCallback: function() {
		var callback = oldCallbacks.pop() || ( jQuery.expando + "_" + ( nonce++ ) );
		this[ callback ] = true;
		return callback;
	}
} );

// Detect, normalize options and install callbacks for jsonp requests
jQuery.ajaxPrefilter( "json jsonp", function( s, originalSettings, jqXHR ) {

	var callbackName, overwritten, responseContainer,
		jsonProp = s.jsonp !== false && ( rjsonp.test( s.url ) ?
			"url" :
			typeof s.data === "string" &&
				( s.contentType || "" )
					.indexOf( "application/x-www-form-urlencoded" ) === 0 &&
				rjsonp.test( s.data ) && "data"
		);

	// Handle iff the expected data type is "jsonp" or we have a parameter to set
	if ( jsonProp || s.dataTypes[ 0 ] === "jsonp" ) {

		// Get callback name, remembering preexisting value associated with it
		callbackName = s.jsonpCallback = isFunction( s.jsonpCallback ) ?
			s.jsonpCallback() :
			s.jsonpCallback;

		// Insert callback into url or form data
		if ( jsonProp ) {
			s[ jsonProp ] = s[ jsonProp ].replace( rjsonp, "$1" + callbackName );
		} else if ( s.jsonp !== false ) {
			s.url += ( rquery.test( s.url ) ? "&" : "?" ) + s.jsonp + "=" + callbackName;
		}

		// Use data converter to retrieve json after script execution
		s.converters[ "script json" ] = function() {
			if ( !responseContainer ) {
				jQuery.error( callbackName + " was not called" );
			}
			return responseContainer[ 0 ];
		};

		// Force json dataType
		s.dataTypes[ 0 ] = "json";

		// Install callback
		overwritten = window[ callbackName ];
		window[ callbackName ] = function() {
			responseContainer = arguments;
		};

		// Clean-up function (fires after converters)
		jqXHR.always( function() {

			// If previous value didn't exist - remove it
			if ( overwritten === undefined ) {
				jQuery( window ).removeProp( callbackName );

			// Otherwise restore preexisting value
			} else {
				window[ callbackName ] = overwritten;
			}

			// Save back as free
			if ( s[ callbackName ] ) {

				// Make sure that re-using the options doesn't screw things around
				s.jsonpCallback = originalSettings.jsonpCallback;

				// Save the callback name for future use
				oldCallbacks.push( callbackName );
			}

			// Call if it was a function and we have a response
			if ( responseContainer && isFunction( overwritten ) ) {
				overwritten( responseContainer[ 0 ] );
			}

			responseContainer = overwritten = undefined;
		} );

		// Delegate to script
		return "script";
	}
} );


// Support: Safari 8 only
// In Safari 8 documents created via document.implementation.createHTMLDocument
// collapse sibling forms: the second one becomes a child of the first one.
// Because of that, this security measure has to be disabled in Safari 8.
// https://bugs.webkit.org/show_bug.cgi?id=137337
support.createHTMLDocument = ( function() {
	var body = document.implementation.createHTMLDocument( "" ).body;
	body.innerHTML = "<form></form><form></form>";
	return body.childNodes.length === 2;
} )();


// Argument "data" should be string of html
// context (optional): If specified, the fragment will be created in this context,
// defaults to document
// keepScripts (optional): If true, will include scripts passed in the html string
jQuery.parseHTML = function( data, context, keepScripts ) {
	if ( typeof data !== "string" ) {
		return [];
	}
	if ( typeof context === "boolean" ) {
		keepScripts = context;
		context = false;
	}

	var base, parsed, scripts;

	if ( !context ) {

		// Stop scripts or inline event handlers from being executed immediately
		// by using document.implementation
		if ( support.createHTMLDocument ) {
			context = document.implementation.createHTMLDocument( "" );

			// Set the base href for the created document
			// so any parsed elements with URLs
			// are based on the document's URL (gh-2965)
			base = context.createElement( "base" );
			base.href = document.location.href;
			context.head.appendChild( base );
		} else {
			context = document;
		}
	}

	parsed = rsingleTag.exec( data );
	scripts = !keepScripts && [];

	// Single tag
	if ( parsed ) {
		return [ context.createElement( parsed[ 1 ] ) ];
	}

	parsed = buildFragment( [ data ], context, scripts );

	if ( scripts && scripts.length ) {
		jQuery( scripts ).remove();
	}

	return jQuery.merge( [], parsed.childNodes );
};


/**
 * Load a url into a page
 */
jQuery.fn.load = function( url, params, callback ) {
	var selector, type, response,
		self = this,
		off = url.indexOf( " " );

	if ( off > -1 ) {
		selector = stripAndCollapse( url.slice( off ) );
		url = url.slice( 0, off );
	}

	// If it's a function
	if ( isFunction( params ) ) {

		// We assume that it's the callback
		callback = params;
		params = undefined;

	// Otherwise, build a param string
	} else if ( params && typeof params === "object" ) {
		type = "POST";
	}

	// If we have elements to modify, make the request
	if ( self.length > 0 ) {
		jQuery.ajax( {
			url: url,

			// If "type" variable is undefined, then "GET" method will be used.
			// Make value of this field explicit since
			// user can override it through ajaxSetup method
			type: type || "GET",
			dataType: "html",
			data: params
		} ).done( function( responseText ) {

			// Save response for use in complete callback
			response = arguments;

			self.html( selector ?

				// If a selector was specified, locate the right elements in a dummy div
				// Exclude scripts to avoid IE 'Permission Denied' errors
				jQuery( "<div>" ).append( jQuery.parseHTML( responseText ) ).find( selector ) :

				// Otherwise use the full result
				responseText );

		// If the request succeeds, this function gets "data", "status", "jqXHR"
		// but they are ignored because response was set above.
		// If it fails, this function gets "jqXHR", "status", "error"
		} ).always( callback && function( jqXHR, status ) {
			self.each( function() {
				callback.apply( this, response || [ jqXHR.responseText, status, jqXHR ] );
			} );
		} );
	}

	return this;
};


// Attach a bunch of functions for handling common AJAX events
jQuery.each( [
	"ajaxStart",
	"ajaxStop",
	"ajaxComplete",
	"ajaxError",
	"ajaxSuccess",
	"ajaxSend"
], function( i, type ) {
	jQuery.fn[ type ] = function( fn ) {
		return this.on( type, fn );
	};
} );


jQuery.expr.pseudos.animated = function( elem ) {
	return jQuery.grep( jQuery.timers, function( fn ) {
		return elem === fn.elem;
	} ).length;
};


jQuery.offset = {
	setOffset: function( elem, options, i ) {
		var curPosition, curLeft, curCSSTop, curTop, curOffset, curCSSLeft, calculatePosition,
			position = jQuery.css( elem, "position" ),
			curElem = jQuery( elem ),
			props = {};

		// Set position first, in-case top/left are set even on static elem
		if ( position === "static" ) {
			elem.style.position = "relative";
		}

		curOffset = curElem.offset();
		curCSSTop = jQuery.css( elem, "top" );
		curCSSLeft = jQuery.css( elem, "left" );
		calculatePosition = ( position === "absolute" || position === "fixed" ) &&
			( curCSSTop + curCSSLeft ).indexOf( "auto" ) > -1;

		// Need to be able to calculate position if either
		// top or left is auto and position is either absolute or fixed
		if ( calculatePosition ) {
			curPosition = curElem.position();
			curTop = curPosition.top;
			curLeft = curPosition.left;

		} else {
			curTop = parseFloat( curCSSTop ) || 0;
			curLeft = parseFloat( curCSSLeft ) || 0;
		}

		if ( isFunction( options ) ) {

			// Use jQuery.extend here to allow modification of coordinates argument (gh-1848)
			options = options.call( elem, i, jQuery.extend( {}, curOffset ) );
		}

		if ( options.top != null ) {
			props.top = ( options.top - curOffset.top ) + curTop;
		}
		if ( options.left != null ) {
			props.left = ( options.left - curOffset.left ) + curLeft;
		}

		if ( "using" in options ) {
			options.using.call( elem, props );

		} else {
			curElem.css( props );
		}
	}
};

jQuery.fn.extend( {

	// offset() relates an element's border box to the document origin
	offset: function( options ) {

		// Preserve chaining for setter
		if ( arguments.length ) {
			return options === undefined ?
				this :
				this.each( function( i ) {
					jQuery.offset.setOffset( this, options, i );
				} );
		}

		var rect, win,
			elem = this[ 0 ];

		if ( !elem ) {
			return;
		}

		// Return zeros for disconnected and hidden (display: none) elements (gh-2310)
		// Support: IE <=11 only
		// Running getBoundingClientRect on a
		// disconnected node in IE throws an error
		if ( !elem.getClientRects().length ) {
			return { top: 0, left: 0 };
		}

		// Get document-relative position by adding viewport scroll to viewport-relative gBCR
		rect = elem.getBoundingClientRect();
		win = elem.ownerDocument.defaultView;
		return {
			top: rect.top + win.pageYOffset,
			left: rect.left + win.pageXOffset
		};
	},

	// position() relates an element's margin box to its offset parent's padding box
	// This corresponds to the behavior of CSS absolute positioning
	position: function() {
		if ( !this[ 0 ] ) {
			return;
		}

		var offsetParent, offset, doc,
			elem = this[ 0 ],
			parentOffset = { top: 0, left: 0 };

		// position:fixed elements are offset from the viewport, which itself always has zero offset
		if ( jQuery.css( elem, "position" ) === "fixed" ) {

			// Assume position:fixed implies availability of getBoundingClientRect
			offset = elem.getBoundingClientRect();

		} else {
			offset = this.offset();

			// Account for the *real* offset parent, which can be the document or its root element
			// when a statically positioned element is identified
			doc = elem.ownerDocument;
			offsetParent = elem.offsetParent || doc.documentElement;
			while ( offsetParent &&
				( offsetParent === doc.body || offsetParent === doc.documentElement ) &&
				jQuery.css( offsetParent, "position" ) === "static" ) {

				offsetParent = offsetParent.parentNode;
			}
			if ( offsetParent && offsetParent !== elem && offsetParent.nodeType === 1 ) {

				// Incorporate borders into its offset, since they are outside its content origin
				parentOffset = jQuery( offsetParent ).offset();
				parentOffset.top += jQuery.css( offsetParent, "borderTopWidth", true );
				parentOffset.left += jQuery.css( offsetParent, "borderLeftWidth", true );
			}
		}

		// Subtract parent offsets and element margins
		return {
			top: offset.top - parentOffset.top - jQuery.css( elem, "marginTop", true ),
			left: offset.left - parentOffset.left - jQuery.css( elem, "marginLeft", true )
		};
	},

	// This method will return documentElement in the following cases:
	// 1) For the element inside the iframe without offsetParent, this method will return
	//    documentElement of the parent window
	// 2) For the hidden or detached element
	// 3) For body or html element, i.e. in case of the html node - it will return itself
	//
	// but those exceptions were never presented as a real life use-cases
	// and might be considered as more preferable results.
	//
	// This logic, however, is not guaranteed and can change at any point in the future
	offsetParent: function() {
		return this.map( function() {
			var offsetParent = this.offsetParent;

			while ( offsetParent && jQuery.css( offsetParent, "position" ) === "static" ) {
				offsetParent = offsetParent.offsetParent;
			}

			return offsetParent || documentElement;
		} );
	}
} );

// Create scrollLeft and scrollTop methods
jQuery.each( { scrollLeft: "pageXOffset", scrollTop: "pageYOffset" }, function( method, prop ) {
	var top = "pageYOffset" === prop;

	jQuery.fn[ method ] = function( val ) {
		return access( this, function( elem, method, val ) {

			// Coalesce documents and windows
			var win;
			if ( isWindow( elem ) ) {
				win = elem;
			} else if ( elem.nodeType === 9 ) {
				win = elem.defaultView;
			}

			if ( val === undefined ) {
				return win ? win[ prop ] : elem[ method ];
			}

			if ( win ) {
				win.scrollTo(
					!top ? val : win.pageXOffset,
					top ? val : win.pageYOffset
				);

			} else {
				elem[ method ] = val;
			}
		}, method, val, arguments.length );
	};
} );

// Support: Safari <=7 - 9.1, Chrome <=37 - 49
// Add the top/left cssHooks using jQuery.fn.position
// Webkit bug: https://bugs.webkit.org/show_bug.cgi?id=29084
// Blink bug: https://bugs.chromium.org/p/chromium/issues/detail?id=589347
// getComputedStyle returns percent when specified for top/left/bottom/right;
// rather than make the css module depend on the offset module, just check for it here
jQuery.each( [ "top", "left" ], function( i, prop ) {
	jQuery.cssHooks[ prop ] = addGetHookIf( support.pixelPosition,
		function( elem, computed ) {
			if ( computed ) {
				computed = curCSS( elem, prop );

				// If curCSS returns percentage, fallback to offset
				return rnumnonpx.test( computed ) ?
					jQuery( elem ).position()[ prop ] + "px" :
					computed;
			}
		}
	);
} );


// Create innerHeight, innerWidth, height, width, outerHeight and outerWidth methods
jQuery.each( { Height: "height", Width: "width" }, function( name, type ) {
	jQuery.each( { padding: "inner" + name, content: type, "": "outer" + name },
		function( defaultExtra, funcName ) {

		// Margin is only for outerHeight, outerWidth
		jQuery.fn[ funcName ] = function( margin, value ) {
			var chainable = arguments.length && ( defaultExtra || typeof margin !== "boolean" ),
				extra = defaultExtra || ( margin === true || value === true ? "margin" : "border" );

			return access( this, function( elem, type, value ) {
				var doc;

				if ( isWindow( elem ) ) {

					// $( window ).outerWidth/Height return w/h including scrollbars (gh-1729)
					return funcName.indexOf( "outer" ) === 0 ?
						elem[ "inner" + name ] :
						elem.document.documentElement[ "client" + name ];
				}

				// Get document width or height
				if ( elem.nodeType === 9 ) {
					doc = elem.documentElement;

					// Either scroll[Width/Height] or offset[Width/Height] or client[Width/Height],
					// whichever is greatest
					return Math.max(
						elem.body[ "scroll" + name ], doc[ "scroll" + name ],
						elem.body[ "offset" + name ], doc[ "offset" + name ],
						doc[ "client" + name ]
					);
				}

				return value === undefined ?

					// Get width or height on the element, requesting but not forcing parseFloat
					jQuery.css( elem, type, extra ) :

					// Set width or height on the element
					jQuery.style( elem, type, value, extra );
			}, type, chainable ? margin : undefined, chainable );
		};
	} );
} );


jQuery.each( ( "blur focus focusin focusout resize scroll click dblclick " +
	"mousedown mouseup mousemove mouseover mouseout mouseenter mouseleave " +
	"change select submit keydown keypress keyup contextmenu" ).split( " " ),
	function( i, name ) {

	// Handle event binding
	jQuery.fn[ name ] = function( data, fn ) {
		return arguments.length > 0 ?
			this.on( name, null, data, fn ) :
			this.trigger( name );
	};
} );

jQuery.fn.extend( {
	hover: function( fnOver, fnOut ) {
		return this.mouseenter( fnOver ).mouseleave( fnOut || fnOver );
	}
} );


jQuery.fn.extend( {

	bind: function( types, data, fn ) {
		return this.on( types, null, data, fn );
	},
	unbind: function( types, fn ) {
		return this.off( types, null, fn );
	},

	delegate: function( selector, types, data, fn ) {
		return this.on( types, selector, data, fn );
	},
	undelegate: function( selector, types, fn ) {

		// ( namespace ) or ( selector, types [, fn] )
		return arguments.length === 1 ?
			this.off( selector, "**" ) :
			this.off( types, selector || "**", fn );
	}
} );

// Bind a function to a context, optionally partially applying any
// arguments.
// jQuery.proxy is deprecated to promote standards (specifically Function#bind)
// However, it is not slated for removal any time soon
jQuery.proxy = function( fn, context ) {
	var tmp, args, proxy;

	if ( typeof context === "string" ) {
		tmp = fn[ context ];
		context = fn;
		fn = tmp;
	}

	// Quick check to determine if target is callable, in the spec
	// this throws a TypeError, but we will just return undefined.
	if ( !isFunction( fn ) ) {
		return undefined;
	}

	// Simulated bind
	args = slice.call( arguments, 2 );
	proxy = function() {
		return fn.apply( context || this, args.concat( slice.call( arguments ) ) );
	};

	// Set the guid of unique handler to the same of original handler, so it can be removed
	proxy.guid = fn.guid = fn.guid || jQuery.guid++;

	return proxy;
};

jQuery.holdReady = function( hold ) {
	if ( hold ) {
		jQuery.readyWait++;
	} else {
		jQuery.ready( true );
	}
};
jQuery.isArray = Array.isArray;
jQuery.parseJSON = JSON.parse;
jQuery.nodeName = nodeName;
jQuery.isFunction = isFunction;
jQuery.isWindow = isWindow;
jQuery.camelCase = camelCase;
jQuery.type = toType;

jQuery.now = Date.now;

jQuery.isNumeric = function( obj ) {

	// As of jQuery 3.0, isNumeric is limited to
	// strings and numbers (primitives or objects)
	// that can be coerced to finite numbers (gh-2662)
	var type = jQuery.type( obj );
	return ( type === "number" || type === "string" ) &&

		// parseFloat NaNs numeric-cast false positives ("")
		// ...but misinterprets leading-number strings, particularly hex literals ("0x...")
		// subtraction forces infinities to NaN
		!isNaN( obj - parseFloat( obj ) );
};


// Register as a named AMD module, since jQuery can be concatenated with other
// files that may use define, but not via a proper concatenation script that
// understands anonymous AMD modules. A named AMD is safest and most robust
// way to register. Lowercase jquery is used because AMD module names are
// derived from file names, and jQuery is normally delivered in a lowercase
// file name. Do this after creating the global so that if an AMD module wants
// to call noConflict to hide this version of jQuery, it will work.

// Note that for maximum portability, libraries that are not jQuery should
// declare themselves as anonymous modules, and avoid setting a global if an
// AMD loader is present. jQuery is a special case. For more information, see
// https://github.com/jrburke/requirejs/wiki/Updating-existing-libraries#wiki-anon

if ( typeof define === "function" && define.amd ) {
	define( "jquery", [], function() {
		return jQuery;
	} );
}


var

	// Map over jQuery in case of overwrite
	_jQuery = window.jQuery,

	// Map over the $ in case of overwrite
	_$ = window.$;

jQuery.noConflict = function( deep ) {
	if ( window.$ === jQuery ) {
		window.$ = _$;
	}

	if ( deep && window.jQuery === jQuery ) {
		window.jQuery = _jQuery;
	}

	return jQuery;
};

// Expose jQuery and $ identifiers, even in AMD
// (#7102#comment:10, https://github.com/jquery/jquery/pull/557)
// and CommonJS for browser emulators (#13566)
if ( !noGlobal ) {
	window.jQuery = window.$ = jQuery;
}


return jQuery;
} );
```
