## Supplemental Code S1 for "Accurate reconstruction of bacterial pan- and core- genomes with PEPPAN": index.html

Welcome to PEPPA’s documentation! — PEPPA 1.0 documentation

### Welcome to PEPPA’s documentation!¶

Contents:

- installation
- quickstart
- parameters
- inputs
- outputs
  - inference=ortholog\_group:<source\_genome>:<exemplar\_gene>:<allele\_ID>:<start & end coordinates of alignment in the exemplar gene>:<start & end coordinates of alignmenet in the genome>

#### About PEPPA¶

PEPPA (Phylogeny Enhanced Pipeline for PAn-genome) is a pipeline that can construct a pan-genome from thousands of genetically diversified bacterial genomes.
PEPPA implements a combination of tree- and synteny-based approaches to identify and exclude paralogous genes,
as well as similarity-based gene predictions that support consistent annotations of genes and pseudogenes in individual genomes.

#### Citation¶

If you use GrapeTree please cite the pre-print in BioRxiv:

Z Zhou, M Achtman (2020) “Accurate reconstruction of the pan- and core- genomes of bacteria with PEPPA”
bioRxiv, doi: [https://doi.org/10.1101/2020.01.03.894154](https://doi.org/10.1101/2020.01.03.894154)

### Indices and tables¶

- Index
- Module Index
- Search Page

### PEPPA

##### Navigation

Contents:

- installation
- quickstart
- parameters
- inputs
- outputs

##### Related Topics

- Documentation overview
  - Next: installation

##### Quick search

©2020, Zhemin Zhou.
|
Powered by Sphinx 3.0.2
& Alabaster 0.7.12
|
Page source
