## Supplemental Code S1 for "Accurate reconstruction of bacterial pan- and core- genomes with PEPPAN": search.html

Search — PEPPA 1.0 documentation

### Search

Please activate JavaScript to enable the search
functionality.

Searching for multiple words only shows matches that contain
all words.

### PEPPA

##### Navigation

Contents:

- installation
- quickstart
- parameters
- inputs
- outputs

##### Related Topics

- Documentation overview

©2020, Zhemin Zhou.
|
Powered by Sphinx 3.0.2
& Alabaster 0.7.12
