## Supplemental Code S1 for "Accurate reconstruction of bacterial pan- and core- genomes with PEPPAN": searchindex.js

```
Search.setIndex({docnames:["index","usage/inputs","usage/installation","usage/outputs","usage/parameters","usage/quickstart"],envversion:{"sphinx.domains.c":2,"sphinx.domains.changeset":1,"sphinx.domains.citation":1,"sphinx.domains.cpp":2,"sphinx.domains.index":1,"sphinx.domains.javascript":2,"sphinx.domains.math":2,"sphinx.domains.python":2,"sphinx.domains.rst":2,"sphinx.domains.std":1,sphinx:56},filenames:["index.rst","usage/inputs.rst","usage/installation.rst","usage/outputs.rst","usage/parameters.rst","usage/quickstart.rst"],objects:{},objnames:{},objtypes:{},terms:{"default":4,"final":3,"long":4,"short":4,CDS:3,The:[3,4],There:3,Use:4,absenc:3,absent:4,accur:0,achtman:0,across:3,addit:4,align:[0,4],alignmenet:0,all:[3,4],allel:[3,4],allele_id:0,allow:4,allowed_sigma:4,also:4,amount:4,analys:[3,4],analysi:[3,4],ani:3,annot:[0,3,4],appli:4,approach:0,argument:4,assign:3,attribut:3,back:4,bacteri:0,bacteria:0,base:[0,3,4],bash:5,becaus:3,between:4,biorxiv:0,blast:4,block:4,both:4,built:3,can:[0,4],cat:5,cds_cgav:3,cds_content:3,cdss:[3,4,5],cgav:[3,4],cgmlst:[3,4],check:4,cite:0,clust_ident:4,clust_match_prop:4,cluster:4,code:4,codon:4,combin:[0,5],comma:4,compar:3,consid:4,consist:0,construct:0,contain:[3,4],content:0,coordin:0,core:[0,3,4],cover:4,creat:4,curv:[3,4],dataset:4,defin:4,delimit:4,depend:3,describ:[3,4],diamond:4,differ:[3,4],disabl:4,discard:4,disrupt:3,diversifi:0,doi:0,draw:4,each:[3,4],end:0,enhanc:0,entri:3,equal:4,exampl:5,exclud:0,exemplar:0,exemplar_gen:0,exit:4,extrem:4,fals:4,fast:4,fasta:4,field:3,file:[3,4],filenam:4,flag:4,fna:3,folder:4,form:4,format:[3,4],fraction:3,fragment:4,frameshift:4,from:[0,4],gcf_000010485:5,gene:[0,4],gene_cgav:3,gene_cont:3,gener:[3,4,5],genet:0,genom:[0,4,5],gff3:3,gff:[3,4,5],grapetre:0,greater:4,group:[3,4],gtabl:4,had:4,have:4,help:4,http:0,ident:4,identifi:0,ignor:4,imperfect:4,implement:0,includ:[3,4,5],incompletecd:[4,5],index:0,individu:[0,4],infer:0,inparalog:4,input:[0,4],instal:0,intact:[3,4],integ:4,its:4,kept:4,keyword:4,larg:4,larger:4,length:4,less:4,link:4,link_diff:4,link_gap:4,list:4,map:4,match:4,match_frag_len:4,match_frag_prop:4,match_ident:4,match_len1:4,match_len2:4,match_len:4,match_prop1:4,match_prop2:4,match_prop:4,matrix:[3,4],messag:4,metagenom:4,method:4,min:4,min_cd:[4,5],minimum:4,misc_featur:3,mmseq:4,mode:4,modul:0,more:4,multipl:4,mycoplasma:4,n_thread:4,name:3,neighborhood:4,noneighborcheck:4,normal:4,nucl:4,nucleotid:4,number:4,nwk:3,old:3,old_locus_tag:3,one:3,onli:[4,5],option:4,order:4,org:0,origin:3,ortholog:[3,4],ortholog_group:0,other:4,output:[0,4],overlap:[3,4],page:0,pan:[0,3,5],paralog:[0,4],paramet:0,peppa:[3,4,5],peppa_out:5,peppa_pars:[3,4,5],phylogeni:0,pipelin:0,pleas:0,posit:4,potenti:3,pre:0,predict:[0,3,5],prefer:4,prefix:[3,4],presenc:[3,4],present:4,print:0,prior:4,prioriti:4,proport:4,pseudogen:[0,3,4,5],psueogen:3,put:4,python:[4,5],queri:4,quick:[],quickstart:0,random:4,rare:3,rarefract:4,read:4,reconstruct:0,refer:4,region:4,reliabl:4,remain:4,remov:3,report:4,requir:4,result:3,retiev:4,same:4,sbh:4,scheme:[3,4],score:4,search:[0,4],sensit:4,separ:4,sequenc:[3,4],set:4,shorter:4,should:3,show:4,sife:4,sigma:4,similar:[0,3,4],small:4,source_genom:0,split:4,st131:5,stage:4,start:[0,4],stop:4,summari:5,support:[0,4],suspici:3,synteni:0,tabl:4,than:4,thei:4,thi:[3,4],thousand:0,thread:4,togeth:4,translat:4,treat:3,tree:[0,3,4],two:[3,4],type:4,uniqu:3,univers:4,unrecogn:4,untrust:4,usag:4,use:[0,4],used:4,using:[4,5],valu:4,variat:[3,4],well:0,when:4,which:4,within:4,without:4,xxx:4,you:[0,4],zhou:0},titles:["Welcome to PEPPA\u2019s documentation!","inputs","installation","Outputs","Parameters","Quick Start"],titleterms:{about:0,align:3,alignmenet:3,allele_id:3,citat:0,coordin:3,document:0,end:3,exemplar:3,exemplar_gen:3,gene:3,genom:3,indic:0,infer:3,input:1,instal:2,ortholog_group:3,output:3,paramet:4,peppa:0,quick:5,source_genom:3,start:[3,5],tabl:0,welcom:0}})
```
