## Supplemental Code S1 for "Accurate reconstruction of bacterial pan- and core- genomes with PEPPAN": inputs.html

inputs — PEPPA 1.0 documentation

### inputs¶

### PEPPA

##### Navigation

Contents:

- installation
- quickstart
- parameters
- inputs
- outputs

##### Related Topics

- Documentation overview
  - Previous: Parameters
  - Next: Outputs

##### Quick search

©2020, Zhemin Zhou.
|
Powered by Sphinx 3.0.2
& Alabaster 0.7.12
|
Page source
