## Supplemental Code S1 for "Accurate reconstruction of bacterial pan- and core- genomes with PEPPAN": outputs.html

Outputs — PEPPA 1.0 documentation

### Outputs¶

#### Outputs for PEPPA.py
There are two final outputs for PEPPA.py:

1. &lt;prefix&gt;.PEPPA.gff

   This file includes all pan-genes predicted by PEPPA in GFF3 format. Intact CDSs are assigned as “CDS”, disrupted genes (potential pseudogenes) are assigned as “pseudogene” and suspicious annotations that are removed are described as “misc\_feature” entries.

- If any of the predicted CDSs and psueogenes overlaps with old gene predictions in the original GFF files, the old gene is described in an attribute named “old\_locus\_tag” of the entry.
- Each gene and pseudogene is assigned into one of the orthologous groups. This orthologous group is described in “inference” field in a format of:

#### inference=ortholog\_group:<source\_genome>:<exemplar\_gene>:<allele\_ID>:<start & end coordinates of alignment in the exemplar gene>:<start & end coordinates of alignmenet in the genome>¶

2. &lt;prefix&gt;.alleles.fna

   This file contains all the unique alleles of all pan genes predicted by PEPPA.

#### Outputs for PEPPA\_parse.py
PEPPA\_parse.py generates:

1. &lt;prefix&gt;.gene\_content.matrix or &lt;prefix&gt;.CDS\_content.matrix

A matrix of gene presence/absence in all genomes.

2. &lt;prefix&gt;.gene\_content.nwk or &lt;prefix&gt;.CDS\_content.nwk

A tree built based on gene presence/absence in all genomes.

3. &lt;prefix&gt;.gene\_content.curve or &lt;prefix&gt;.CDS\_content.curve

The rare-fraction curves for the pan-genome and core-genome

4. &lt;prefix&gt;.gene\_CGAV.tree or &lt;prefix&gt;.CDS\_CGAV.tree

Core Genome Allelic Variation trees based on the sequence differences of the core genes. This is similar to but should not be treated as a cgMLST scheme, because the genes included in the analysis depend on the genomes. The result of CGAV analysis is not comparable across different analyses.

### PEPPA

##### Navigation

Contents:

- installation
- quickstart
- parameters
- inputs
- outputs
  - inference=ortholog\_group:<source\_genome>:<exemplar\_gene>:<allele\_ID>:<start & end coordinates of alignment in the exemplar gene>:<start & end coordinates of alignmenet in the genome>

##### Related Topics

- Documentation overview
  - Previous: inputs

##### Quick search

©2020, Zhemin Zhou.
|
Powered by Sphinx 3.0.2
& Alabaster 0.7.12
|
Page source
