## Supplemental Code S1 for "Accurate reconstruction of bacterial pan- and core- genomes with PEPPAN": parameters.html

Parameters — PEPPA 1.0 documentation

### Parameters¶

#### Usage for PEPPA.py
```
$ python PEPPA.py -h
usage: PEPPA.py [-h] [-p PREFIX] [-g GENES] [-P PRIORITY] [-t N\_THREAD]

> [-o ORTHOLOGY] [-n] [–min\_cds MIN\_CDS]
> [–incompleteCDS INCOMPLETECDS] [–gtable GTABLE]
> [–clust\_identity CLUST\_IDENTITY]
> [–clust\_match\_prop CLUST\_MATCH\_PROP] [–nucl]
> [–match\_identity MATCH\_IDENTITY] [–match\_prop MATCH\_PROP]
> [–match\_len MATCH\_LEN] [–match\_prop1 MATCH\_PROP1]
> [–match\_len1 MATCH\_LEN1] [–match\_prop2 MATCH\_PROP2]
> [–match\_len2 MATCH\_LEN2] [–match\_frag\_prop MATCH\_FRAG\_PROP]
> [–match\_frag\_len MATCH\_FRAG\_LEN] [–link\_gap LINK\_GAP]
> [–link\_diff LINK\_DIFF] [–allowed\_sigma ALLOWED\_SIGMA]
> [–pseudogene PSEUDOGENE] [–untrusted UNTRUSTED]
> [–metagenome]
> [N [N …]]

PEPPA.py
(1) Retieves genes and genomic sequences from GFF files and FASTA files.
(2) Groups genes into clusters using mmseq.
(3) Maps gene clusters back to genomes.
(4) Discard paralogous alignments.
(5) Discard orthologous clusters if they had regions which overlapped with the regions within other sets that had greater scores.
(6) Re-annotate genomes using the remained of orthologs.

positional arguments:
:   N [REQUIRED] GFF files containing both annotations and sequences.
    :   If you have sequences and GFF annotations in separate files,
        they can also be put in as: <GFF>,<fasta>

optional arguments:
:   `-h, --help`
    :   show this help message and exit

    `-p PREFIX, --prefix PREFIX`
    :   [Default: PEPPA] prefix for the outputs.

    `-g GENES, --genes GENES`
    :   [optional] Comma delimited filenames that contain fasta of additional genes.

    `-P PRIORITY, --priority PRIORITY`
    :   [optional] Comma delimited, ordered list of GFFs or gene fasta files that are more reliable than others.
        Genes contained in these files are preferred in all stages.

    `-t N_THREAD, --n_thread N_THREAD`
    :   [Default: 20] Number of threads to use. Default: 20

    `-o ORTHOLOGY, --orthology ORTHOLOGY`
    :   [Default: nj] Method to define orthologous groups.
        nj [default], ml (for small dataset) or sbh (extremely large datasets)

    `-n, --noNeighborCheck`
    :   [Default: False] Flag to disable checking of neighborhood for paralog splitting.

    `--min_cds MIN_CDS`
    :   [Default: 150] Minimum length for a gene to be used in similarity searches.

    `--incompleteCDS INCOMPLETECDS`
    :   [Default: ‘’] Allowed types of imperfection for reference genes.
        ‘s’: allows unrecognized start codon.
        ‘e’: allows unrecognized stop codon.
        ‘i’: allows stop codons in the coding region.
        ‘f’: allows frameshift in the coding region.
        Multiple keywords can be used together. e.g., use ‘sife’ to allow random sequences.

    `--gtable GTABLE`
    :   [Default: 11] Translate table to Use. Only support 11 and 4 (for Mycoplasma)

    `--clust_identity CLUST_IDENTITY`
    :   minimum identities of mmseqs clusters. Default: 0.9

    `--clust_match_prop CLUST_MATCH_PROP`
    :   minimum matches in mmseqs clusters. Default: 0.9

    `--nucl`
    :   disable Diamond search. Fast but less sensitive when nucleotide identities < 0.9

    `--match_identity MATCH_IDENTITY`
    :   minimum identities in BLAST search. Default: 0.5

    `--match_prop MATCH_PROP`
    :   minimum match proportion for normal genes in BLAST search. Default: 0.6

    `--match_len MATCH_LEN`
    :   minimum match length for normal genes in BLAST search. Default: 250

    `--match_prop1 MATCH_PROP1`
    :   minimum match proportion for short genes in BLAST search. Default: 0.8

    `--match_len1 MATCH_LEN1`
    :   minimum match length for short genes in BLAST search. Default: 100

    `--match_prop2 MATCH_PROP2`
    :   minimum match proportion for long genes in BLAST search. Default: 0.4

    `--match_len2 MATCH_LEN2`
    :   minimum match length for long genes in BLAST search. Default: 400

    `--match_frag_prop MATCH_FRAG_PROP`
    :   Min proportion of each fragment for fragmented matches. Default: 0.3

    `--match_frag_len MATCH_FRAG_LEN`
    :   Min length of each fragment for fragmented matches. Default: 50

    `--link_gap LINK_GAP`
    :   Consider two fragmented matches within N bases as a linked block. Default: 300

    `--link_diff LINK_DIFF`
    :   Form a linked block when the covered regions in the reference gene
        and the queried genome differed by no more than this value. Default: 1.2

    `--allowed_sigma ALLOWED_SIGMA`
    :   Allowed number of sigma for paralogous splitting.
        The larger, the more variations are kept as inparalogs. Default: 3.

    `--pseudogene PSEUDOGENE`
    :   A match is reported as pseudogene if its coding region is less than this amount of the reference gene. Default: 0.8

    `--untrusted UNTRUSTED`
    :   FORMAT: l,p; A gene is not reported if it is shorter than l and present in less than p of prior annotations. Default: 300,0.3

    `--metagenome`
    :   Set to metagenome mode. equals to
        “–nucl –incompleteCDS sife –clust\_identity 0.99 –clust\_match\_prop 0.8 –match\_identity 0.98 –orthology sbh”

```

#### Usage for PEPPA\_parser.py
```
$ python PEPPA\_parser.py -h
usage: PEPPA\_parser.py [-h] -g GFF [-p PREFIX] [-s SPLIT] [-P] [-m] [-t]

> [-a CGAV] [-c]

PEPPA\_parser.py
(1) reads xxx.PEPPA.gff file
(2) split it into individual GFF files
(3) draw a present/absent matrix
(4) create a tree based on gene presence
(5) draw rarefraction curves of all genes and only intact CDSs

optional arguments:
:   `-h, --help`
    :   show this help message and exit

    `-g GFF, --gff GFF`
    :   [REQUIRED] generated PEPPA.gff file from PEPPA.py.

    `-p PREFIX, --prefix PREFIX`
    :   [Default: Same prefix as GFF input] Prefix for all outputs.

    `-s SPLIT, --split SPLIT`
    :   [optional] A folder for splitted GFF files.

    `-P, --pseudogene`
    :   [Default: Use Pseudogene] Flag to ignore pseudogenes in all analyses.

    `-m, --matrix`
    :   [Default: False] Flag to generate the gene present/absent matrix

    `-t, --tree`
    :   [Default: False] Flag to generate the gene present/absent tree

    `-a CGAV, --cgav CGAV`
    :   [Default: -1] Set to an integer between 0 and 100 to apply a Core Gene Allelic Variation tree.
        The value describes % of presence for a gene to be included in the analysis.
        This is similar to cgMLST tree but without an universal scheme.

    `-c, --curve`
    :   [Default: False] Flag to generate a rarefraction curve.

```

### PEPPA

##### Navigation

Contents:

- installation
- quickstart
- parameters
- inputs
- outputs

##### Related Topics

- Documentation overview
  - Previous: Quick Start
  - Next: inputs

##### Quick search

©2020, Zhemin Zhou.
|
Powered by Sphinx 3.0.2
& Alabaster 0.7.12
|
Page source
