## Supplemental Code S1 for "Accurate reconstruction of bacterial pan- and core- genomes with PEPPAN": quickstart.html

Quick Start — PEPPA 1.0 documentation

### Quick Start¶

#### Quick Start (included in example.bash)
```
$ cat example.bash
# generate pan-genome prediction using PEPPA
python PEPPA.py -P examples/GCF\_000010485.combined.gff.gz –min\_cds 60 –incompleteCDS s -p examples/ST131 examples/\*.gff.gz

# generate summaries for PEPPA predicted CDSs and pseudogenes
python PEPPA\_parser.py -g examples/ST131.PEPPA.gff -s examples/PEPPA\_out -m -t -c -a 95

# generate summaries for PEPPA predicted CDSs only
python PEPPA\_parser.py -g examples/ST131.PEPPA.gff -s examples/PEPPA\_out -m -t -c -a 95 -P
```

### PEPPA

##### Navigation

Contents:

- installation
- quickstart
- parameters
- inputs
- outputs

##### Related Topics

- Documentation overview
  - Previous: installation
  - Next: Parameters

##### Quick search

©2020, Zhemin Zhou.
|
Powered by Sphinx 3.0.2
& Alabaster 0.7.12
|
Page source
