## Supplemental Code S1 for "Accurate reconstruction of bacterial pan- and core- genomes with PEPPAN": make.bat

```
@ECHO OFF

REM Command file for Sphinx documentation

pushd %~dp0

set PDFLATEX=latexmk -pdf -dvi- -ps-

set "LATEXOPTS= "

if "%1" == "" goto all-pdf

if "%1" == "all-pdf" (
	:all-pdf
	for %%i in (*.tex) do (
		%PDFLATEX% %LATEXMKOPTS% %%i
	)
	goto end
)

if "%1" == "all-pdf-ja" (
	goto all-pdf
)

if "%1" == "clean" (
	del /q /s *.dvi *.log *.ind *.aux *.toc *.syn *.idx *.out *.ilg *.pla *.ps *.tar *.tar.gz *.tar.bz2 *.tar.xz *.fls *.fdb_latexmk
	goto end
)

:end
popd
```
