## Supplemental Code S1 for "Accurate reconstruction of bacterial pan- and core- genomes with PEPPAN": Makefile

```
# Makefile for Sphinx LaTeX output

ALLDOCS = $(basename $(wildcard *.tex))
ALLPDF = $(addsuffix .pdf,$(ALLDOCS))
ALLDVI = $(addsuffix .dvi,$(ALLDOCS))
ALLXDV =
ALLPS  = $(addsuffix .ps,$(ALLDOCS))

# Prefix for archive names
ARCHIVEPREFIX =
# Additional LaTeX options (passed via variables in latexmkrc/latexmkjarc file)
export LATEXOPTS ?=
# Additional latexmk options
LATEXMKOPTS ?=
# format: pdf or dvi (used only by archive targets)
FMT = pdf

LATEX = latexmk -dvi
PDFLATEX = latexmk -pdf -dvi- -ps-


%.dvi: %.tex FORCE_MAKE
	$(LATEX) $(LATEXMKOPTS) '$<'

%.ps: %.dvi
	dvips '$<'

%.pdf: %.tex FORCE_MAKE
	$(PDFLATEX) $(LATEXMKOPTS) '$<'

all: $(ALLPDF)

all-dvi: $(ALLDVI)

all-ps: $(ALLPS)

all-pdf: $(ALLPDF)

zip: all-$(FMT)
	mkdir $(ARCHIVEPREFIX)docs-$(FMT)
	cp $(ALLPDF) $(ARCHIVEPREFIX)docs-$(FMT)
	zip -q -r -9 $(ARCHIVEPREFIX)docs-$(FMT).zip $(ARCHIVEPREFIX)docs-$(FMT)
	rm -r $(ARCHIVEPREFIX)docs-$(FMT)

tar: all-$(FMT)
	mkdir $(ARCHIVEPREFIX)docs-$(FMT)
	cp $(ALLPDF) $(ARCHIVEPREFIX)docs-$(FMT)
	tar cf $(ARCHIVEPREFIX)docs-$(FMT).tar $(ARCHIVEPREFIX)docs-$(FMT)
	rm -r $(ARCHIVEPREFIX)docs-$(FMT)

gz: tar
	gzip -9 < $(ARCHIVEPREFIX)docs-$(FMT).tar > $(ARCHIVEPREFIX)docs-$(FMT).tar.gz

bz2: tar
	bzip2 -9 -k $(ARCHIVEPREFIX)docs-$(FMT).tar

xz: tar
	xz -9 -k $(ARCHIVEPREFIX)docs-$(FMT).tar

clean:
	rm -f *.log *.ind *.aux *.toc *.syn *.idx *.out *.ilg *.pla *.ps *.tar *.tar.gz *.tar.bz2 *.tar.xz $(ALLPDF) $(ALLDVI) $(ALLXDV) *.fls *.fdb_latexmk

.PHONY: all all-pdf all-dvi all-ps clean zip tar gz bz2 xz
.PHONY: FORCE_MAKE
```
