## Supplemental Code S1 for "Accurate reconstruction of bacterial pan- and core- genomes with PEPPAN": peppa.pdf

#### *Release 1.0*

**Zhemin Zhou**

**Apr 22, 2020**

### CONTENTS:


**1 installation** **1**

**2 Quick Start** **3**

**3 Parameters** **5**

**4 inputs** **9**

**5 Outputs** **11**

5.1 inference=ortholog_group:<source_genome>:<exemplar_gene>:<allele_ID>:<start & end coordinates of alignment in the exemplar gene>:<start & end coordinates of alignmenet in the genome> . . 11

**6 About PEPPA** **13**

**7 Citation** **15**

**8 Indices and tables** **17**

CHAPTER

**-c, --curve** [Default: False] Flag to generate a rarefraction curve.

```
```

CHAPTER

### FOUR

### INPUTS

CHAPTER

FIVE

### OUTPUTS

#### Outputs for PEPPA.py There are two final outputs for PEPPA.py:

1. <prefix>.PEPPA.gff

   This file includes all pan-genes predicted by PEPPA in GFF3 format. Intact CDSs are assigned as "CDS", disrupted genes (potential pseudogenes) are assigned as "pseudogene" and suspicious annotations that are removed are described as "misc_feature" entries.

- If any of the predicted CDSs and psueogenes overlaps with old gene predictions in the original GFF files, the old gene is described in an attribute named "old_locus_tag" of the entry.
- Each gene and pseudogene is assigned into one of the orthologous groups. This orthologous group is described in "inference" field in a format of:

#### 5.1 inference=ortholog_group:<source_genome>:<exemplar_gene>:<allele_ID> [illegible] & end coordinates of alignment in the exemplar gene>:<start & end coordinates of alignmenet in the genome>

2. <prefix>.alleles.fna

   This file contains all the unique alleles of all pan genes predicted by PEPPA.

#### Outputs for PEPPA_parse.py PEPPA_parse.py generates:

1. <prefix>.gene_content.matrix or <prefix>.CDS_content.matrix

A matrix of gene presence/absence in all genomes.

2. <prefix>.gene_content.nwk or <prefix>.CDS_content.nwk

A tree built based on gene presence/absence in all genomes.

3. <prefix>.gene_content.curve or <prefix>.CDS_content.curve

The rare-fraction curves for the pan-genome and core-genome

4. <prefix>.gene_CGAV.tree or <prefix>.CDS_CGAV.tree

Core Genome Allelic Variation trees based on the sequence differences of the core genes. This is similar to but should not be treated as a cgMLST scheme, because the genes included in the analysis depend on the genomes. The result of CGAV analysis is not comparable across different analyses.

CHAPTER

SIX

### ABOUT PEPPA

PEPPA (Phylogeny Enhanced Pipeline for PAn-genome) is a pipeline that can construct a pan-genome from thousands of genetically diversified bacterial genomes. PEPPA implements a combination of tree- and synteny-based approaches to identify and exclude paralogous genes, as well as similarity-based gene predictions that support consistent annotations of genes and pseudogenes in individual genomes.

CHAPTER

SEVEN

### CITATION

If you use GrapeTree please cite the pre-print in BioRxiv:

Z Zhou, M Achtman (2020) "Accurate reconstruction of the pan- and core- genomes of bacteria with PEPPA" bioRxiv, doi: [https://doi.org/10.1101/2020.01.03.894154{]}(https://doi.org/10.1101/2020.01.03.894154)

CHAPTER

EIGHT

### INDICES AND TABLES

- genindex
- modindex
- search
