## Supplemental Code S1 for "Accurate reconstruction of bacterial pan- and core- genomes with PEPPAN": peppa.tex

%% Generated by Sphinx.
\def\sphinxdocclass{report}
\documentclass[letterpaper,10pt,english]{sphinxmanual}
\ifdefined\pdfpxdimen
   \let\sphinxpxdimen\pdfpxdimen\else\newdimen\sphinxpxdimen
\fi \sphinxpxdimen=.75bp\relax

\PassOptionsToPackage{warn}{textcomp}
\usepackage[utf8]{inputenc}
\ifdefined\DeclareUnicodeCharacter
% support both utf8 and utf8x syntaxes
  \ifdefined\DeclareUnicodeCharacterAsOptional
    \def\sphinxDUC#1{\DeclareUnicodeCharacter{"#1}}
  \else
    \let\sphinxDUC\DeclareUnicodeCharacter
  \fi
  \sphinxDUC{00A0}{\nobreakspace}
  \sphinxDUC{2500}{\sphinxunichar{2500}}
  \sphinxDUC{2502}{\sphinxunichar{2502}}
  \sphinxDUC{2514}{\sphinxunichar{2514}}
  \sphinxDUC{251C}{\sphinxunichar{251C}}
  \sphinxDUC{2572}{\textbackslash}
\fi
\usepackage{cmap}
\usepackage[T1]{fontenc}
\usepackage{amsmath,amssymb,amstext}
\usepackage{babel}


\usepackage{times}
\expandafter\ifx\csname T@LGR\endcsname\relax
\else
% LGR was declared as font encoding
  \substitutefont{LGR}{\rmdefault}{cmr}
  \substitutefont{LGR}{\sfdefault}{cmss}
  \substitutefont{LGR}{\ttdefault}{cmtt}
\fi
\expandafter\ifx\csname T@X2\endcsname\relax
  \expandafter\ifx\csname T@T2A\endcsname\relax
  \else
  % T2A was declared as font encoding
    \substitutefont{T2A}{\rmdefault}{cmr}
    \substitutefont{T2A}{\sfdefault}{cmss}
    \substitutefont{T2A}{\ttdefault}{cmtt}
  \fi
\else
% X2 was declared as font encoding
  \substitutefont{X2}{\rmdefault}{cmr}
  \substitutefont{X2}{\sfdefault}{cmss}
  \substitutefont{X2}{\ttdefault}{cmtt}
\fi


\usepackage[Bjarne]{fncychap}
\usepackage{sphinx}

\fvset{fontsize=\small}
\usepackage{geometry}


% Include hyperref last.
\usepackage{hyperref}
% Fix anchor placement for figures with captions.
\usepackage{hypcap}% it must be loaded after hyperref.
% Set up styles of URL: it should be placed after hyperref.
\urlstyle{same}
\addto\captionsenglish{\renewcommand{\contentsname}{Contents:}}

\usepackage{sphinxmessages}
\setcounter{tocdepth}{1}


\title{PEPPA}
\date{Apr 22, 2020}
\release{1.0}
\author{Zhemin Zhou}
\newcommand{\sphinxlogo}{\vbox{}}
\renewcommand{\releasename}{Release}
\makeindex
\begin{document}

\pagestyle{empty}
\sphinxmaketitle
\pagestyle{plain}
\sphinxtableofcontents
\pagestyle{normal}
\phantomsection\label{\detokenize{index::doc}}


\chapter{installation}
\label{\detokenize{usage/installation:installation}}\label{\detokenize{usage/installation::doc}}

\chapter{Quick Start}
\label{\detokenize{usage/quickstart:quick-start}}\label{\detokenize{usage/quickstart::doc}}
\#\# Quick Start (included in example.bash)
{\color{red}\bfseries{}\textasciigrave{}\textasciigrave{}}\textasciigrave{}
\$ cat example.bash
\# generate pan\sphinxhyphen{}genome prediction using PEPPA
python PEPPA.py \sphinxhyphen{}P examples/GCF\_000010485.combined.gff.gz \textendash{}min\_cds 60 \textendash{}incompleteCDS s \sphinxhyphen{}p examples/ST131 examples/{\color{red}\bfseries{}*}.gff.gz

\# generate summaries for PEPPA predicted CDSs and pseudogenes
python PEPPA\_parser.py \sphinxhyphen{}g examples/ST131.PEPPA.gff \sphinxhyphen{}s examples/PEPPA\_out \sphinxhyphen{}m \sphinxhyphen{}t \sphinxhyphen{}c \sphinxhyphen{}a 95

\# generate summaries for PEPPA predicted CDSs only
python PEPPA\_parser.py \sphinxhyphen{}g examples/ST131.PEPPA.gff \sphinxhyphen{}s examples/PEPPA\_out \sphinxhyphen{}m \sphinxhyphen{}t \sphinxhyphen{}c \sphinxhyphen{}a 95 \sphinxhyphen{}P
{\color{red}\bfseries{}\textasciigrave{}\textasciigrave{}}{\color{red}\bfseries{}\textasciigrave{}}


\chapter{Parameters}
\label{\detokenize{usage/parameters:parameters}}\label{\detokenize{usage/parameters::doc}}
\#\# Usage for PEPPA.py
{\color{red}\bfseries{}\textasciigrave{}\textasciigrave{}}\textasciigrave{}
\$ python PEPPA.py \sphinxhyphen{}h
usage: PEPPA.py {[}\sphinxhyphen{}h{]} {[}\sphinxhyphen{}p PREFIX{]} {[}\sphinxhyphen{}g GENES{]} {[}\sphinxhyphen{}P PRIORITY{]} {[}\sphinxhyphen{}t N\_THREAD{]}
\begin{quote}

{[}\sphinxhyphen{}o ORTHOLOGY{]} {[}\sphinxhyphen{}n{]} {[}\textendash{}min\_cds MIN\_CDS{]}
{[}\textendash{}incompleteCDS INCOMPLETECDS{]} {[}\textendash{}gtable GTABLE{]}
{[}\textendash{}clust\_identity CLUST\_IDENTITY{]}
{[}\textendash{}clust\_match\_prop CLUST\_MATCH\_PROP{]} {[}\textendash{}nucl{]}
{[}\textendash{}match\_identity MATCH\_IDENTITY{]} {[}\textendash{}match\_prop MATCH\_PROP{]}
{[}\textendash{}match\_len MATCH\_LEN{]} {[}\textendash{}match\_prop1 MATCH\_PROP1{]}
{[}\textendash{}match\_len1 MATCH\_LEN1{]} {[}\textendash{}match\_prop2 MATCH\_PROP2{]}
{[}\textendash{}match\_len2 MATCH\_LEN2{]} {[}\textendash{}match\_frag\_prop MATCH\_FRAG\_PROP{]}
{[}\textendash{}match\_frag\_len MATCH\_FRAG\_LEN{]} {[}\textendash{}link\_gap LINK\_GAP{]}
{[}\textendash{}link\_diff LINK\_DIFF{]} {[}\textendash{}allowed\_sigma ALLOWED\_SIGMA{]}
{[}\textendash{}pseudogene PSEUDOGENE{]} {[}\textendash{}untrusted UNTRUSTED{]}
{[}\textendash{}metagenome{]}
{[}N {[}N …{]}{]}
\end{quote}

PEPPA.py
(1) Retieves genes and genomic sequences from GFF files and FASTA files.
(2) Groups genes into clusters using mmseq.
(3) Maps gene clusters back to genomes.
(4) Discard paralogous alignments.
(5) Discard orthologous clusters if they had regions which overlapped with the regions within other sets that had greater scores.
(6) Re\sphinxhyphen{}annotate genomes using the remained of orthologs.
\begin{description}
\item[{positional arguments:}] \leavevmode\begin{description}
\item[{N                     {[}REQUIRED{]} GFF files containing both annotations and sequences.}] \leavevmode
If you have sequences and GFF annotations in separate files,
they can also be put in as: \textless{}GFF\textgreater{},\textless{}fasta\textgreater{}

\end{description}

\item[{optional arguments:}] \leavevmode\begin{optionlist}{3cm}
\item [\sphinxhyphen{}h, \sphinxhyphen{}\sphinxhyphen{}help]  
show this help message and exit
\item [\sphinxhyphen{}p PREFIX, \sphinxhyphen{}\sphinxhyphen{}prefix PREFIX]  
{[}Default: PEPPA{]} prefix for the outputs.
\item [\sphinxhyphen{}g GENES, \sphinxhyphen{}\sphinxhyphen{}genes GENES]  
{[}optional{]} Comma delimited filenames that contain fasta of additional genes.
\item [\sphinxhyphen{}P PRIORITY, \sphinxhyphen{}\sphinxhyphen{}priority PRIORITY]  
{[}optional{]} Comma delimited, ordered list of GFFs or gene fasta files that are more reliable than others.
Genes contained in these files are preferred in all stages.
\item [\sphinxhyphen{}t N\_THREAD, \sphinxhyphen{}\sphinxhyphen{}n\_thread N\_THREAD]  
{[}Default: 20{]} Number of threads to use. Default: 20
\item [\sphinxhyphen{}o ORTHOLOGY, \sphinxhyphen{}\sphinxhyphen{}orthology ORTHOLOGY]  
{[}Default: nj{]} Method to define orthologous groups.
nj {[}default{]}, ml (for small dataset) or sbh (extremely large datasets)
\item [\sphinxhyphen{}n, \sphinxhyphen{}\sphinxhyphen{}noNeighborCheck]  
{[}Default: False{]} Flag to disable checking of neighborhood for paralog splitting.
\item [\sphinxhyphen{}\sphinxhyphen{}min\_cds MIN\_CDS]  
{[}Default: 150{]} Minimum length for a gene to be used in similarity searches.
\item [\sphinxhyphen{}\sphinxhyphen{}incompleteCDS INCOMPLETECDS]  
{[}Default: ‘’{]} Allowed types of imperfection for reference genes.
‘s’: allows unrecognized start codon.
‘e’: allows unrecognized stop codon.
‘i’: allows stop codons in the coding region.
‘f’: allows frameshift in the coding region.
Multiple keywords can be used together. e.g., use ‘sife’ to allow random sequences.
\item [\sphinxhyphen{}\sphinxhyphen{}gtable GTABLE]  
{[}Default: 11{]} Translate table to Use. Only support 11 and 4 (for Mycoplasma)
\item [\sphinxhyphen{}\sphinxhyphen{}clust\_identity CLUST\_IDENTITY]  
minimum identities of mmseqs clusters. Default: 0.9
\item [\sphinxhyphen{}\sphinxhyphen{}clust\_match\_prop CLUST\_MATCH\_PROP]  
minimum matches in mmseqs clusters. Default: 0.9
\item [\sphinxhyphen{}\sphinxhyphen{}nucl]  
disable Diamond search. Fast but less sensitive when nucleotide identities \textless{} 0.9
\item [\sphinxhyphen{}\sphinxhyphen{}match\_identity MATCH\_IDENTITY]  
minimum identities in BLAST search. Default: 0.5
\item [\sphinxhyphen{}\sphinxhyphen{}match\_prop MATCH\_PROP]  
minimum match proportion for normal genes in BLAST search. Default: 0.6
\item [\sphinxhyphen{}\sphinxhyphen{}match\_len MATCH\_LEN]  
minimum match length for normal genes in BLAST search. Default: 250
\item [\sphinxhyphen{}\sphinxhyphen{}match\_prop1 MATCH\_PROP1]  
minimum match proportion for short genes in BLAST search. Default: 0.8
\item [\sphinxhyphen{}\sphinxhyphen{}match\_len1 MATCH\_LEN1]  
minimum match length for short genes in BLAST search. Default: 100
\item [\sphinxhyphen{}\sphinxhyphen{}match\_prop2 MATCH\_PROP2]  
minimum match proportion for long genes in BLAST search. Default: 0.4
\item [\sphinxhyphen{}\sphinxhyphen{}match\_len2 MATCH\_LEN2]  
minimum match length for long genes in BLAST search. Default: 400
\item [\sphinxhyphen{}\sphinxhyphen{}match\_frag\_prop MATCH\_FRAG\_PROP]  
Min proportion of each fragment for fragmented matches. Default: 0.3
\item [\sphinxhyphen{}\sphinxhyphen{}match\_frag\_len MATCH\_FRAG\_LEN]  
Min length of each fragment for fragmented matches. Default: 50
\item [\sphinxhyphen{}\sphinxhyphen{}link\_gap LINK\_GAP]  
Consider two fragmented matches within N bases as a linked block. Default: 300
\item [\sphinxhyphen{}\sphinxhyphen{}link\_diff LINK\_DIFF]  
Form a linked block when the covered regions in the reference gene
and the queried genome differed by no more than this value. Default: 1.2
\item [\sphinxhyphen{}\sphinxhyphen{}allowed\_sigma ALLOWED\_SIGMA]  
Allowed number of sigma for paralogous splitting.
The larger, the more variations are kept as inparalogs. Default: 3.
\item [\sphinxhyphen{}\sphinxhyphen{}pseudogene PSEUDOGENE]  
A match is reported as pseudogene if its coding region is less than this amount of the reference gene. Default: 0.8
\item [\sphinxhyphen{}\sphinxhyphen{}untrusted UNTRUSTED]  
FORMAT: l,p; A gene is not reported if it is shorter than l and present in less than p of prior annotations. Default: 300,0.3
\item [\sphinxhyphen{}\sphinxhyphen{}metagenome]  
Set to metagenome mode. equals to
“\textendash{}nucl \textendash{}incompleteCDS sife \textendash{}clust\_identity 0.99 \textendash{}clust\_match\_prop 0.8 \textendash{}match\_identity 0.98 \textendash{}orthology sbh”
\end{optionlist}

\end{description}

{\color{red}\bfseries{}\textasciigrave{}\textasciigrave{}}{\color{red}\bfseries{}\textasciigrave{}}

\#\# Usage for PEPPA\_parser.py
{\color{red}\bfseries{}\textasciigrave{}\textasciigrave{}}\textasciigrave{}
\$ python PEPPA\_parser.py \sphinxhyphen{}h
usage: PEPPA\_parser.py {[}\sphinxhyphen{}h{]} \sphinxhyphen{}g GFF {[}\sphinxhyphen{}p PREFIX{]} {[}\sphinxhyphen{}s SPLIT{]} {[}\sphinxhyphen{}P{]} {[}\sphinxhyphen{}m{]} {[}\sphinxhyphen{}t{]}
\begin{quote}

{[}\sphinxhyphen{}a CGAV{]} {[}\sphinxhyphen{}c{]}
\end{quote}

PEPPA\_parser.py
(1) reads xxx.PEPPA.gff file
(2) split it into individual GFF files
(3) draw a present/absent matrix
(4) create a tree based on gene presence
(5) draw rarefraction curves of all genes and only intact CDSs
\begin{description}
\item[{optional arguments:}] \leavevmode\begin{optionlist}{3cm}
\item [\sphinxhyphen{}h, \sphinxhyphen{}\sphinxhyphen{}help]  
show this help message and exit
\item [\sphinxhyphen{}g GFF, \sphinxhyphen{}\sphinxhyphen{}gff GFF]  
{[}REQUIRED{]} generated PEPPA.gff file from PEPPA.py.
\item [\sphinxhyphen{}p PREFIX, \sphinxhyphen{}\sphinxhyphen{}prefix PREFIX]  
{[}Default: Same prefix as GFF input{]} Prefix for all outputs.
\item [\sphinxhyphen{}s SPLIT, \sphinxhyphen{}\sphinxhyphen{}split SPLIT]  
{[}optional{]} A folder for splitted GFF files.
\item [\sphinxhyphen{}P, \sphinxhyphen{}\sphinxhyphen{}pseudogene]  
{[}Default: Use Pseudogene{]} Flag to ignore pseudogenes in all analyses.
\item [\sphinxhyphen{}m, \sphinxhyphen{}\sphinxhyphen{}matrix]  
{[}Default: False{]} Flag to generate the gene present/absent matrix
\item [\sphinxhyphen{}t, \sphinxhyphen{}\sphinxhyphen{}tree]  
{[}Default: False{]} Flag to generate the gene present/absent tree
\item [\sphinxhyphen{}a CGAV, \sphinxhyphen{}\sphinxhyphen{}cgav CGAV]  
{[}Default: \sphinxhyphen{}1{]} Set to an integer between 0 and 100 to apply a Core Gene Allelic Variation tree.
The value describes \% of presence for a gene to be included in the analysis.
This is similar to cgMLST tree but without an universal scheme.
\item [\sphinxhyphen{}c, \sphinxhyphen{}\sphinxhyphen{}curve]  
{[}Default: False{]} Flag to generate a rarefraction curve.
\end{optionlist}

\end{description}

{\color{red}\bfseries{}\textasciigrave{}\textasciigrave{}}{\color{red}\bfseries{}\textasciigrave{}}


\chapter{inputs}
\label{\detokenize{usage/inputs:inputs}}\label{\detokenize{usage/inputs::doc}}

\chapter{Outputs}
\label{\detokenize{usage/outputs:outputs}}\label{\detokenize{usage/outputs::doc}}
\#\# Outputs for PEPPA.py
There are two final outputs for PEPPA.py:
\begin{enumerate}
\sphinxsetlistlabels{\arabic}{enumi}{enumii}{}{.}%
\item {} 
\&lt;prefix\&gt;.PEPPA.gff

This file includes all pan\sphinxhyphen{}genes predicted by PEPPA in GFF3 format. Intact CDSs are assigned as “CDS”, disrupted genes (potential pseudogenes) are assigned as “pseudogene” and suspicious annotations that are removed are described as “misc\_feature” entries.

\end{enumerate}
\begin{itemize}
\item {} 
If any of the predicted CDSs and psueogenes overlaps with old gene predictions in the original GFF files, the old gene is described in an attribute named “old\_locus\_tag” of the entry.

\item {} 
Each gene and pseudogene is assigned into one of the orthologous groups. This orthologous group is described in “inference” field in a format of:

\end{itemize}


\section{inference=ortholog\_group:\textless{}source\_genome\textgreater{}:\textless{}exemplar\_gene\textgreater{}:\textless{}allele\_ID\textgreater{}:\textless{}start \& end coordinates of alignment in the exemplar gene\textgreater{}:\textless{}start \& end coordinates of alignmenet in the genome\textgreater{}}
\label{\detokenize{usage/outputs:inference-ortholog-group-source-genome-exemplar-gene-allele-id-start-end-coordinates-of-alignment-in-the-exemplar-gene-start-end-coordinates-of-alignmenet-in-the-genome}}\begin{enumerate}
\sphinxsetlistlabels{\arabic}{enumi}{enumii}{}{.}%
\setcounter{enumi}{1}
\item {} 
\&lt;prefix\&gt;.alleles.fna

This file contains all the unique alleles of all pan genes predicted by PEPPA.

\end{enumerate}

\#\# Outputs for PEPPA\_parse.py
PEPPA\_parse.py generates:
\begin{enumerate}
\sphinxsetlistlabels{\arabic}{enumi}{enumii}{}{.}%
\item {} 
\&lt;prefix\&gt;.gene\_content.matrix or \&lt;prefix\&gt;.CDS\_content.matrix

\end{enumerate}

A matrix of gene presence/absence in all genomes.
\begin{enumerate}
\sphinxsetlistlabels{\arabic}{enumi}{enumii}{}{.}%
\setcounter{enumi}{1}
\item {} 
\&lt;prefix\&gt;.gene\_content.nwk or \&lt;prefix\&gt;.CDS\_content.nwk

\end{enumerate}

A tree built based on gene presence/absence in all genomes.
\begin{enumerate}
\sphinxsetlistlabels{\arabic}{enumi}{enumii}{}{.}%
\setcounter{enumi}{2}
\item {} 
\&lt;prefix\&gt;.gene\_content.curve or \&lt;prefix\&gt;.CDS\_content.curve

\end{enumerate}

The rare\sphinxhyphen{}fraction curves for the pan\sphinxhyphen{}genome and core\sphinxhyphen{}genome
\begin{enumerate}
\sphinxsetlistlabels{\arabic}{enumi}{enumii}{}{.}%
\setcounter{enumi}{3}
\item {} 
\&lt;prefix\&gt;.gene\_CGAV.tree or \&lt;prefix\&gt;.CDS\_CGAV.tree

\end{enumerate}

Core Genome Allelic Variation trees based on the sequence differences of the core genes. This is similar to  but should not be treated as a cgMLST scheme, because the genes included in the analysis depend on the genomes. The result of CGAV analysis is not comparable across different analyses.


\chapter{About PEPPA}
\label{\detokenize{index:about-peppa}}
PEPPA (Phylogeny Enhanced Pipeline for PAn\sphinxhyphen{}genome) is a pipeline that can construct a pan\sphinxhyphen{}genome from thousands of genetically diversified bacterial genomes.
PEPPA implements a combination of tree\sphinxhyphen{} and synteny\sphinxhyphen{}based approaches to identify and exclude paralogous genes,
as well as similarity\sphinxhyphen{}based gene predictions that support consistent annotations of genes and pseudogenes in individual genomes.


\chapter{Citation}
\label{\detokenize{index:citation}}
If you use GrapeTree please cite the pre\sphinxhyphen{}print in BioRxiv:

Z Zhou, M Achtman (2020) “Accurate reconstruction of the pan\sphinxhyphen{} and core\sphinxhyphen{} genomes of bacteria with PEPPA”
bioRxiv, doi: {[}\sphinxurl{https://doi.org/10.1101/2020.01.03.894154{]}(https://doi.org/10.1101/2020.01.03.894154})


\chapter{Indices and tables}
\label{\detokenize{index:indices-and-tables}}\begin{itemize}
\item {} 
\DUrole{xref,std,std-ref}{genindex}

\item {} 
\DUrole{xref,std,std-ref}{modindex}

\item {} 
\DUrole{xref,std,std-ref}{search}

\end{itemize}


\renewcommand{\indexname}{Index}
\printindex
\end{document}
