## Supplemental Code S1 for "Accurate reconstruction of bacterial pan- and core- genomes with PEPPAN": example.bash

```
# generate pan-genome prediction using PEPPA
PEPPA -p examples/ST131 -P examples/GCF_000010485.combined.gff.gz examples/*.gff.gz

# generate summaries for PEPPA predicted CDSs and pseudogenes
PEPPA_parser -g examples/ST131.PEPPA.gff -s examples/PEPPA_out -t -c -a 95
```
