## Supplemental Code S1 for "Accurate reconstruction of bacterial pan- and core- genomes with PEPPAN": clust.py

```
import argparse, tempfile, glob, os, subprocess, sys, shutil
try:
    from configure import externals, uopen, xrange, logger, transeq
except :
    from .configure import externals, uopen, xrange, logger, transeq

def readFasta(fasta) :
    sequence = []
    with uopen(fasta) as fin :
        for line in fin :
            if line.startswith('>') :
                name = line[1:].strip().split()[0]
                sequence.append([name, []])
            elif len(line) > 0 and not line.startswith('#') :
                sequence[-1][1].extend(line.strip().split())
    for s in sequence :
        s[1] = (''.join(s[1])).upper()
    return sequence


def clust(argv) :
    parser = argparse.ArgumentParser(description='Get clusters and exemplars of clusters from gene sequences using mmseqs linclust.')
    parser.add_argument('-i', '--input', help='[INPUT; REQUIRED] name of the file containing gene sequneces in FASTA format.', required=True)
    parser.add_argument('-p', '--prefix', help='[OUTPUT; REQUIRED] prefix of the outputs.', required=True)
    parser.add_argument('-d', '--identity', help='[PARAM; DEFAULT: 0.9] minimum intra-cluster identity.', default=0.9, type=float)
    parser.add_argument('-c', '--coverage', help='[PARAM; DEFAULT: 0.9] minimum intra-cluster coverage.', default=0.9, type=float)
    parser.add_argument('-t', '--n_thread', help='[PARAM; DEFAULT: 8]   number of threads to use.', default=8, type=int)
    parser.add_argument('-a', '--translate', help='[PARAM; DEFAULT: False] activate to cluster in translated sequence.', default=False, action='store_true')
    args = parser.parse_args(argv)
    exemplar, clust = getClust(args.prefix, args.input, args.__dict__)
    logger('Exemplar sequences in {0}'.format(exemplar))
    logger('Clusters in {0}'.format(clust))
    return exemplar, clust
def getClust(prefix, genes, params) :
    groups = {}
    dirPath = tempfile.mkdtemp(prefix='NS_', dir='.')
    try:
        if not params['translate'] :
            geneFile = genes
        else :
            na_seqs = readFasta(genes)
            aa_seqs = transeq(na_seqs, frame='1', transl_table='starts')
            with open(os.path.join(dirPath, 'seq.aa'), 'w') as fout :
                for n, s in aa_seqs :
                    fout.write('>{0}\n{1}\n'.format(n, s[0]))
            geneFile = os.path.join(dirPath, 'seq.aa')
        seqDb = os.path.join(dirPath, 'seq.db')
        tmpDb = os.path.join(dirPath, 'tmp')
        lcDb = os.path.join(dirPath, 'seq.lc')
        tabFile = os.path.join(dirPath, 'clust.tab')
        refFile = os.path.join(dirPath, 'seq.ref')
        
        nRef = 999999999999999
        for ite in xrange(3) :
            if os.path.isdir(tmpDb) :
                shutil.rmtree(tmpDb)
            os.makedirs(tmpDb)
            if os.path.isfile(seqDb) :
                list(map(os.unlink, glob.glob(seqDb + '*')))
            if os.path.isfile(lcDb) :
                list(map(os.unlink, glob.glob(lcDb + '*')))
            subprocess.Popen('{0} createdb {2} {1} -v 0'.format(externals['mmseqs'], seqDb, geneFile).split()).communicate()
            subprocess.Popen('{0} linclust {1} {2} {3} --min-seq-id {4} -c {5} --threads {6} -v 0'.format( \
                externals['mmseqs'], seqDb, lcDb, tmpDb, params['identity'], params['coverage'], params['n_thread']).split(), stdout=subprocess.PIPE).communicate()
            subprocess.Popen('{0} createtsv {1} {1} {2} {3}'.format(\
                externals['mmseqs'], seqDb, lcDb, tabFile).split(), stdout = subprocess.PIPE).communicate()
            with open(tabFile) as fin :
                for line in fin :
                    part = line.strip().split()
                    groups[part[1]] = part[0]
            tmp = []
            with open(geneFile) as fin :
                toWrite, used_grps = False, {None:1}
                for line in fin :
                    if line.startswith('>') :
                        name = line[1:].strip().split()[0]
                        grp = groups.get(name, None)
                        toWrite = False if grp in used_grps else True
                        if toWrite :
                            used_grps[grp] = name
                    if toWrite :
                        tmp.append(line)
            for gene, grp in groups.items() :
                if grp in used_grps :
                    groups[gene] = used_grps[grp]
            with open(refFile, 'w') as fout :
                for line in tmp :
                    fout.write(line)
            if nRef <= len(used_grps) :
                break
            nRef = len(used_grps)
            geneFile = refFile
        if not params['translate'] :
            shutil.copy2(refFile, '{0}.clust.exemplar'.format(prefix))
        else :
            rSeq = readFasta(refFile)
            na_seqs = dict(na_seqs)
            with open('{0}.clust.exemplar'.format(prefix), 'w') as fout :
                for n, s in rSeq:
                    fout.write('>{0}\n{1}\n'.format(n, na_seqs[n]))
    finally :
        shutil.rmtree(dirPath)
    with open('{0}.clust.tab'.format(prefix), 'w') as fout :
        for gene, grp in sorted(groups.items()) :
            g = gene
            while g != grp :
                g, grp = grp, groups[grp]
            groups[gene] = grp
            fout.write('{0}\t{1}\n'.format(gene, grp))
    
    return '{0}.clust.exemplar'.format(prefix), '{0}.clust.tab'.format(prefix)

if __name__ == '__main__' :
    clust(sys.argv[1:])
```
