## Supplemental Code S1 for "Accurate reconstruction of bacterial pan- and core- genomes with PEPPAN": configure.py

```
import os, sys, subprocess, numpy as np, gzip, io, re, shutil

from datetime import datetime
if sys.version_info[0] < 3:
    from cStringIO import StringIO
    xrange = xrange
    asc2int = np.uint8
else :
    from io import StringIO
    xrange = range
    asc2int = np.uint32

dependency_path = os.path.join(os.path.dirname(os.path.dirname(os.path.realpath(__file__))), 'dependencies')
externals = dict(
    mmseqs = 'mmseqs',
    makeblastdb = 'makeblastdb', 
    diamond = 'diamond', 
    blastn = 'blastn', 
    fasttree = ['FastTreeMP', 'FastTree'], 
    rapidnj = 'rapidnj', 
    pigz = ['pigz', 'gzip'], 
)

def checkExecutable(commands) :
    try :
        if not os.path.exists(commands[-1]) :
            return False
        return subprocess.Popen(commands+['-h'], stdout=subprocess.PIPE, stderr=subprocess.PIPE).wait() <= 1 or subprocess.Popen(commands, stdout=subprocess.PIPE, stderr=subprocess.PIPE).wait() <= 1
    except  :
        return False

def checkDependencies() :
    global externals
    for k, v in externals.items() :
        vs = v if isinstance(v, (tuple, list)) else [v]
        for v in vs :
            fname = os.path.join(dependency_path, v)
            if checkExecutable([fname]) :
                externals[k] = fname
                break
            else :
                externals[k] = shutil.which(v)
                if externals[k] is not None :
                    break
        assert externals[k] is not None, 'Required dependency {0} is not executable and has not been installed in environmental PATH variable'.format(' or '.join(vs))
checkDependencies()

# * is designated as U; index is : (ord(r)-65)*32 + ord(q)-65
blosum62 = np.array([  4., -2.,  0., -2., -1., -2.,  0., -2., -1.,  0., -1., -1., -1., -2.,  0., -1., -1., -1.,  1.,  0., -4.,  0.,
                       -3.,  0., -2., -1.,  0.,  0.,  0.,  0.,  0.,  0., -2., 4., -3.,  4.,  1., -3., -1.,  0., -3.,  0.,  0., -4.,
                       -3.,  3.,  0., -2.,  0., -1.,  0., -1., -4., -3., -4., -1., -3.,  1.,  0.,  0.,  0.,  0.,  0.,  0.,  0., -3.,
                       9., -3., -4., -2., -3., -3., -1.,  0., -3., -1., -1., -3.,  0., -3., -3., -3., -1., -1., -4., -1., -2., -2.,
                       -2., -3.,  0.,  0.,  0.,  0.,  0.,  0., -2.,  4., -3., 6.,  2., -3., -1., -1., -3.,  0., -1., -4., -3.,  1.,
                       0., -1.,  0., -2.,  0., -1., -4., -3., -4., -1., -3., 1.,  0.,  0.,  0.,  0.,  0.,  0., -1.,  1., -4.,  2.,
                       5., -3., -2.,  0., -3.,  0.,  1., -3., -2.,  0.,  0., -1.,  2.,  0.,  0., -1., -4., -2., -3., -1., -2.,  4.,
                       0.,  0.,  0.,  0.,  0.,  0., -2., -3., -2., -3., -3., 6., -3., -1.,  0.,  0., -3.,  0.,  0., -3.,  0., -4.,
                       -3., -3., -2., -2., -4., -1.,  1., -1.,  3., -3.,  0., 0.,  0.,  0.,  0.,  0.,  0., -1., -3., -1., -2., -3.,
                       6., -2., -4.,  0., -2., -4., -3.,  0.,  0., -2., -2., -2.,  0., -2., -4., -3., -2., -1., -3., -2.,  0.,  0.,
                       0.,  0.,  0.,  0., -2.,  0., -3., -1.,  0., -1., -2., 8., -3.,  0., -1., -3., -2.,  1.,  0., -2.,  0.,  0.,
                       -1., -2., -4., -3., -2., -1.,  2.,  0.,  0.,  0.,  0., 0.,  0.,  0., -1., -3., -1., -3., -3.,  0., -4., -3.,
                       4.,  0., -3.,  2.,  1., -3.,  0., -3., -3., -3., -2., -1., -4.,  3., -3., -1., -1., -3.,  0.,  0.,  0.,  0.,
                       0.,  0.,  0.,  0.,  0.,  0.,  0.,  0.,  0.,  0.,  0., 0.,  0.,  0.,  0.,  0.,  0.,  0.,  0.,  0.,  0.,  0.,
                       0.,  0.,  0.,  0.,  0.,  0.,  0.,  0.,  0.,  0.,  0., 0., -1.,  0., -3., -1.,  1., -3., -2., -1., -3.,  0.,
                       5., -2., -1.,  0.,  0., -1.,  1.,  2.,  0., -1., -4., -2., -3., -1., -2.,  1.,  0.,  0.,  0.,  0.,  0.,  0.,
                       -1., -4., -1., -4., -3.,  0., -4., -3.,  2.,  0., -2., 4.,  2., -3.,  0., -3., -2., -2., -2., -1., -4.,  1.,
                       -2., -1., -1., -3.,  0.,  0.,  0.,  0.,  0.,  0., -1., -3., -1., -3., -2.,  0., -3., -2.,  1.,  0., -1.,  2.,
                       5., -2.,  0., -2.,  0., -1., -1., -1., -4.,  1., -1., -1., -1., -1.,  0.,  0.,  0.,  0.,  0.,  0., -2.,  3.,
                       -3.,  1.,  0., -3.,  0.,  1., -3.,  0.,  0., -3., -2., 6.,  0., -2.,  0.,  0.,  1.,  0., -4., -3., -4., -1.,
                       -2.,  0.,  0.,  0.,  0.,  0.,  0.,  0.,  0.,  0.,  0., 0.,  0.,  0.,  0.,  0.,  0.,  0.,  0.,  0.,  0.,  0.,
                       0.,  0.,  0.,  0.,  0.,  0.,  0.,  0.,  0.,  0.,  0., 0.,  0.,  0.,  0.,  0.,  0.,  0., -1., -2., -3., -1.,
                       -1., -4., -2., -2., -3.,  0., -1., -3., -2., -2.,  0., 7., -1., -2., -1., -1., -4., -2., -4., -2., -3., -1.,
                       0.,  0.,  0.,  0.,  0.,  0., -1.,  0., -3.,  0.,  2., -3., -2.,  0., -3.,  0.,  1., -2.,  0.,  0.,  0., -1.,
                       5.,  1.,  0., -1., -4., -2., -2., -1., -1.,  3.,  0., 0.,  0.,  0.,  0.,  0., -1., -1., -3., -2.,  0., -3.,
                       -2.,  0., -3.,  0.,  2., -2., -1.,  0.,  0., -2.,  1., 5., -1., -1., -4., -3., -3., -1., -2.,  0.,  0.,  0.,
                       0.,  0.,  0.,  0.,  1.,  0., -1.,  0.,  0., -2.,  0., -1., -2.,  0.,  0., -2., -1.,  1.,  0., -1.,  0., -1.,
                       4.,  1., -4., -2., -3.,  0., -2.,  0.,  0.,  0.,  0., 0.,  0.,  0.,  0., -1., -1., -1., -1., -2., -2., -2.,
                       -1.,  0., -1., -1., -1.,  0.,  0., -1., -1., -1.,  1., 5., -4.,  0., -2.,  0., -2., -1.,  0.,  0.,  0.,  0.,
                       0.,  0., -4., -4., -4., -4., -4., -4., -4., -4., -4., 0., -4., -4., -4., -4.,  0., -4., -4., -4., -4., -4.,
                       1., -4., -4., -4., -4., -4.,  0.,  0.,  0.,  0.,  0., 0.,  0., -3., -1., -3., -2., -1., -3., -3.,  3.,  0.,
                       -2.,  1.,  1., -3.,  0., -2., -2., -3., -2.,  0., -4., 4., -3., -1., -1., -2.,  0.,  0.,  0.,  0.,  0.,  0.,
                       -3., -4., -2., -4., -3.,  1., -2., -2., -3.,  0., -3., -2., -1., -4.,  0., -4., -2., -3., -3., -2., -4., -3.,
                       11., -2.,  2., -3.,  0.,  0.,  0.,  0.,  0.,  0.,  0., -1., -2., -1., -1., -1., -1., -1., -1.,  0., -1., -1.,
                       -1., -1.,  0., -2., -1., -1.,  0.,  0., -4., -1., -2., -1., -1., -1.,  0.,  0.,  0.,  0.,  0.,  0., -2., -3.,
                       -2., -3., -2.,  3., -3.,  2., -1.,  0., -2., -1., -1., -2.,  0., -3., -1., -2., -2., -2., -4., -1.,  2., -1.,
                       7., -2.,  0.,  0.,  0.,  0.,  0.,  0., -1.,  1., -3., 1.,  4., -3., -2.,  0., -3.,  0.,  1., -3., -1.,  0.,
                       0., -1.,  3.,  0.,  0., -1., -4., -2., -3., -1., -2., 4.,  0.,  0.,  0.,  0.,  0.,  0.,  0.,  0.,  0.,  0.,
                       0.,  0.,  0.,  0.,  0.,  0.,  0.,  0.,  0.,  0.,  0., 0.,  0.,  0.,  0.,  0.,  0.,  0.,  0.,  0.,  0.,  0.])


class uopen(object) :
    def __init__(self, fname, label='r') :
        self.fout = None
        if label.find('r')>=0 :
            self.fstream = subprocess.Popen([externals['pigz'], '-cd', fname], stdout=subprocess.PIPE, stderr=subprocess.PIPE, universal_newlines=True).stdout if fname.lower().endswith('gz') else open(fname)
        elif label.find('w') >= 0 :
            if sys.version.startswith('3') :
                self.fout = gzip.open(fname, 'wb')
                self.fstream = io.TextIOWrapper(self.fout, encoding='utf-8')
            else :
                self.fstream = gzip.open(fname, 'wb')
    def __enter__(self) :
        return self.fstream
    def __exit__(self, type, value, traceback) :
        self.fstream.close()
        if self.fout :
            self.fout.close()
        return 
    def __iter__(self) :
        return self.fstream
    def __next__(self) :
        return self.fstream
    def write(self, doc) :
        self.fstream.write(doc)
    def close(self) :
        self.fstream.close()


def readFasta(fasta, headOnly=False) :
    sequence = {}
    with uopen(fasta) as fin :
        for line in fin :
            if line.startswith('>') :
                name = line[1:].strip().split()[0]
                sequence[name] = []
            elif len(line) > 0 and not line.startswith('#') and not headOnly :
                sequence[name].extend(line.strip().split())
    for s in sequence :
        sequence[s] = (''.join(sequence[s])).upper()
    return sequence
def readFastq(fastq) :
    sequence, qual = {}, {}
    with uopen(fastq) as fin :
        line = fin.readline()
        if not line.startswith('@') :
            sequence = readFasta(fastq)
            return sequence, { n: re.sub(r'[^!]', 'I', re.sub(r'[^ACGTacgt]', '!', s)) for n, s in sequence.items() }
    with uopen(fastq) as fin :
        for lineId, line in enumerate(fin) :
            if lineId % 4 == 0 :
                name = line[1:].strip().split()[0]
                sequence[name] = []
                qual[name] = []
            elif lineId % 4 == 1 :
                sequence[name].extend(line.strip().split())
            elif lineId % 4 == 3 :
                qual[name].extend(line.strip().split())
    for s in sequence :
        sequence[s] = (''.join(sequence[s])).upper()
        qual[s] = ''.join(qual[s])
    return sequence, qual

complement = {'A':'T', 'T':'A', 'G':'C', 'C':'G', 'N':'N'}
def rc(seq, missingValue='N') :
    return ''.join([complement.get(s, missingValue) for s in reversed(seq.upper())])


baseConv = np.empty(255, dtype=int)
baseConv.fill(-100)
baseConv[(np.array(['-', 'A', 'C', 'G', 'T']).view(asc2int),)] = (-100000, 0, 1, 2, 3)
def transeq(seq, frame=7, transl_table=None, markStarts=False) :
    frames = {'F': [1,2,3],
              'R': [4,5,6],
              '7': [1,2,3,4,5,6]}.get( str(frame).upper() , None)
    if frames is None :
        frames = [int(f) for f in str(frame).split(',')]
    
    if transl_table == 4 :
        gtable = np.array(list('KNKNTTTTRSRSIIMIQHQHPPPPRRRRLLLLEDEDAAAAGGGGVVVVXYXYSSSSWCWCLFLF-'))
    else :
        gtable = np.array(list('KNKNTTTTRSRSIIMIQHQHPPPPRRRRLLLLEDEDAAAAGGGGVVVVXYXYSSSSXCWCLFLF-'))
    if markStarts :
        gtable[(np.array([46, 62]),)] = 'M'

    seqs = seq.items() if isinstance(seq, dict) else seq
    nFrame = (max(frames) > 3)
    trans_seq = []
    for n,s in seqs :
        s = baseConv[np.array(list(s.upper())).view(asc2int)]
        if nFrame :
            rs = (3 - s)[::-1]
            rs[rs >= 100] *= -1            
        sf = s.size % 3
        tseq = []
        for f in frames :
            codons = s[f-1:] if f <= 3 else rs[f-4:]
            if codons.size % 3 :
                codons = np.concatenate([codons, [-100]*(3 - codons.size % 3)])
            codons = codons.reshape(-1, 3)
            codon2 = np.sum(codons << [4, 2, 0], 1)
            codon2[codon2 < -50000] = 64
            codon2[codon2 < 0] = 50
            tseq.append(''.join(gtable[codon2].tolist()))
        trans_seq.append([n, tseq])
    return dict(trans_seq) if isinstance(seq, dict) else trans_seq


def logger(log, pipe=sys.stderr) :
    pipe.write('{0}\t{1}\n'.format(str(datetime.now()), log))
    pipe.flush()
```
