## Supplemental Code S1 for "Accurate reconstruction of bacterial pan- and core- genomes with PEPPAN": uberBlast.py

```
import os, sys, tempfile, time, shutil, numpy as np, pandas as pd, re
from numba import jit
from subprocess import Popen, PIPE
from multiprocessing.pool import ThreadPool, Pool
from operator import itemgetter
try:
    from .configure import externals, logger, xrange, readFastq, transeq, blosum62, rc, asc2int
except :
    from configure import externals, logger, xrange, readFastq, transeq, blosum62, rc, asc2int

makeblastdb = externals['makeblastdb']
blastn = externals['blastn']
diamond = externals['diamond']


def parseDiamond(data):
    fn, refseq, qryseq, min_id, min_cov, min_ratio = data
    blastab = []
    with open(fn) as fin :
        for line in fin:
            if line.startswith('@'):
                continue
            part = line.strip().split('\t')
            if part[2] == '*': continue
            qn, qf = part[0].rsplit(':', 1)
            rn, rf, rx = part[2].rsplit(':', 2)
            rs = int(part[3]) + int(rx)
            ql, rl = len(qryseq[str(qn)]), len(refseq[str(rn)])
            qm = len(part[9])
            if qm * 3 < min_cov: continue
            cov_ratio = qm * 3. / ql
            if cov_ratio < min_ratio: continue
            cigar = [[int(n) * 3, t] for n, t in re.findall(r'(\d+)([A-Z])', part[5])]
            cl = np.sum([c[0] for c in cigar])
            variation = float(part[12][5:]) * 3 if part[12].startswith('NM:') else float(
                re.findall('NM:i:(\d+)', line)[0]) * 3

            iden = 1 - round(variation / cl, 3)
            if iden < min_id: continue
            qf, rf = int(qf), int(rf)
            qs = int(part[18][5:]) if part[18].startswith('ZS:') else int(re.findall('ZS:i:(\d+)', line)[0])

            rm = int(np.sum([c[0] for c in cigar if c[1] in {'M', 'D'}]) / 3)
            if rf <= 3:
                rs, r_e = rs * 3 + rf - 3, (rs + rm - 1) * 3 + rf - 1
            else:
                rs, r_e = rl - (rs * 3 + rf - 6) + 1, rl - ((rs + rm - 1) * 3 + rf - 4) + 1
            if qf <= 3:
                qs, qe = qs * 3 + qf - 3, (qs + qm - 1) * 3 + qf - 1
            else:
                qs, qe = ql - (qs * 3 + qf - 6) + 1, ql - ((qs + qm - 1) * 3 + qf - 4) + 1
                qs, qe, rs, r_e = qe, qs, r_e, rs
                cigar = list(reversed(cigar))

            cd = [c[0] for c in cigar if c[1] != 'M']
            score = int(part[14][5:]) if part[14].startswith('ZR:') else int(re.findall('ZR:i:(\d+)', line)[0])
            blastab.append(
                [qn, rn, iden, cl, int(variation - sum(cd)), len(cd), qs, qe, rs, r_e, 0.0, score, ql, rl, cigar])
    try:
        os.unlink(fn)
    except :
        pass

    blastab = pd.DataFrame(blastab)
    if blastab.size > 0:
        blastab[[0, 1]] = blastab[[0, 1]].astype(str)
        np.save(fn+'.match.npy', blastab.values, allow_pickle=True)
        return fn + '.match.npy'
    else :
        return None


@jit(nopython=True)
def tab2overlaps(tabs, ovl_l, ovl_p, nTab, overlaps) :
    ovlId = 0
    for i1 in xrange(overlaps[-1, 0], nTab) :
        t1 = tabs[i1]
        ovl_l2 = min(ovl_l, ovl_p*(t1[3]-t1[2]+1))
        if i1 > overlaps[-1, 0] :
            i2r = xrange(i1+1, nTab)
        else :
            i2r = xrange(overlaps[-1, 1], nTab)
        for i2 in i2r :
            t2 = tabs[i2]
            if t1[0] != t2[0] or t2[2] > t1[3] : break
            ovl = min(t1[3], t2[3]) - t2[2] + 1
            if ovl >= ovl_l2 or ovl >= ovl_p*(t2[3]-t2[2]+1) :
                overlaps[ovlId, :] = [t1[1], t2[1], ovl]
                ovlId += 1
                if ovlId == 1000000 :
                    overlaps[-1, :2] = [i1, i2]
                    break
        if ovlId == 1000000 :
            break
    if ovlId < 1000000 :
        overlaps[-1, :] = -1
    return overlaps


def _linearMerge(data) :
    matches, params = data
    grpCol = pd.Series(data= [[]] * matches.shape[0])
    matches = np.hstack([matches, grpCol.values[:, np.newaxis]])
    gapDist, lenDiff = params[1:]
    gene, geneLen = matches[0][0], matches[0][12]
    tailing = 20
    
    def resolve_edges(edges) :
        grps = []
        for id, m1 in edges[0] :
            for jd, m2 in edges[1] :
                if (m1[1] == m2[1] and max(abs(m1[8]), abs(m1[9])) > min(abs(m2[8]), abs(m2[9])) ) or \
                   abs(m1[2]-m2[2]) > 0.3 or m1[6] >= m2[6] or m1[7] >= m2[7] or m2[6]-m1[7]-1 >= gapDist:
                    continue
                rLen = m2[7] - m1[6] + 1
                g1 = -m1[9]-1 if m1[9] < 0 else m1[13] - m1[9]
                g2 =  m2[8]-1 if m2[8] > 0 else m2[13] + m2[8]
                qLen = m1[9]-m1[8]+1 + m2[9]-m2[8]+1 + g1 + g2
                if g1+g2 >= gapDist or min(rLen, qLen)*lenDiff < max(rLen, qLen) :
                    continue
                overlap = sorted([m1[7] - m2[6] + 1, -g1-g2], reverse=True)

                rLen1, rLen2 = m1[7] - m1[6] + 1, m2[7] - m2[6] + 1
                if overlap[0] > 0 :
                    score = m1[11] + m2[11] - overlap[0] * min( float(m1[11])/rLen1, float(m2[11])/rLen2 )
                    ident = (m1[2]*rLen1 + m2[2]*rLen2 - overlap[0] * min(m1[2], m2[2]))/(rLen1 + rLen2 - overlap[0])
                else :
                    score = m1[11] + m2[11]
                    ident = (m1[2]*rLen1 + m2[2]*rLen2)/(rLen1 + rLen2)
                if overlap[1] < 0 :
                    score +=  overlap[1]/3.
                if score > m1[11] and score > m2[11] :
                    grps.append( [ score, ident, rLen, 1, id, jd ] )
        return grps
    
    groups = []
    prev, edges = matches[0][1], [[], []]
    nSave = len(matches)
    
    for id, m1 in enumerate(matches) :
        rLen1 = m1[7] - m1[6] + 1
        groups.append([ m1[11], m1[2], rLen1, 0, id ])
        if m1[6] > tailing and ((m1[8] > 0 and m1[8] - 1 <= gapDist) or (m1[8] < 0 and m1[13] + m1[8] < gapDist)) :   # any hit within the last 300 bps to either end of a scaffold is a potential fragmented gene
            edges[1].append([id, m1])
        if m1[7] <= m1[12] - tailing :
            if (m1[8] > 0 and m1[13]-m1[9] <= gapDist) or (m1[8] < 0 and -1-m1[9] < gapDist) :
                edges[0].append([id, m1])
            for jd in xrange(id+1, nSave) :
                m2 = matches[jd]
                if m1[1] != m2[1] or (m1[8] < 0 and m2[8] > 0) or m2[8] - m1[9] -1 >= gapDist :    # maximum 600bps between two continuous hits in the same scaffold
                    break
                rLen, qLen = m2[7]-m1[6]+1, m2[9]-m1[8]+1
                if abs(m1[2]-m2[2]) > 0.3 or m1[8]+3 >= m2[8] or m1[9]+3 >= m2[9] or m1[6]+3 >= m2[6] or m1[7]+3 >= m2[7] or m2[6] - m1[7] -1 >= gapDist \
                   or min(rLen, qLen)*lenDiff < max(rLen, qLen) :
                    continue
                rLen2 = m2[7] - m2[6] + 1
                overlap = sorted([m1[7]-m2[6]+1, m1[9]-m2[8]+1], reverse=True)
                if overlap[0] > 0 :
                    score = m1[11] + m2[11] - overlap[0] * min( float(m1[11])/rLen1, float(m2[11])/rLen2 )
                    ident = (m1[2]*rLen1 + m2[2]*rLen2 - overlap[0]*min(m1[2], m2[2]))/(rLen1 + rLen2 - overlap[0])
                else :
                    score = m1[11] + m2[11]
                    ident = (m1[2]*rLen1 + m2[2]*rLen2)/(rLen1 + rLen2)
                if overlap[1] < 0 :
                    score +=  overlap[1]/3.
                if score > m1[11] and score > m2[11] :
                    groups.append( [ score, ident, rLen, 0, id, jd ] )
    if len(edges[0]) and len(edges[1]) :
        groups.extend(resolve_edges(edges))
    if len(groups) > len(matches) :
        groups.sort(reverse=True)
        usedMatches, usedGroups = {}, []
        for grp in groups :
            if (grp[4], 4) in usedMatches or (grp[-1], 5) in usedMatches :
                continue
            if grp[3] > 0 :
                if (grp[4], 5) in usedMatches or (grp[-1], 4) in usedMatches :
                    continue
            if grp[4] != grp[-1] :
                lMat, rMat = matches[grp[4]], matches[grp[-1]]
                il, im = sorted([grp[4], grp[-1]])
                skp = 0
                for i in xrange(il+1, im) :
                    if matches[i][1] in {lMat[1], rMat[1]} :
                        if (i, 4) in usedMatches or (i, 5) in usedMatches :
                            skp = 1
                            break
                if skp :
                    continue
                for i in xrange(il+1, im) :
                    if matches[i][1] in {lMat[1], rMat[1]} :
                        usedMatches[(i, 4)] = usedMatches[(i, 5)] = 0
            usedGroups.append(grp)
            usedMatches[(grp[4], 4)] = usedMatches[(grp[-1], 5)] = 1
            if grp[3] > 0 :
                usedMatches[(grp[4], 5)] = usedMatches[(grp[-1], 4)] = 1

        usedGroups.sort(key=itemgetter(4), reverse=True)
        for gId in xrange(len(usedGroups)-1) :
            g1, g2 = usedGroups[gId:gId+2]
            if g1[4] == g2[-1] :
                m = matches[g1[4]]
                score = g1[0] + g2[0] - m[11]
                length = g1[2] + g2[2] - (m[7]-m[6]+1)
                iden = (g1[1]*g1[2] + g2[1]*g2[2] - min(g1[1],g2[1])*(m[7]-m[6]+1))/length
                usedGroups[gId+1] = [score, iden, length, 0, g2[4]] + g1[4:]
                g1[1] = -1
    else :
        usedGroups = groups
        usedMatches = {(k, k): 1 for k in np.arange(matches.shape[0])}
    for g in usedGroups :
        if g[1] >= 0 :
            ids = [matches[i][15] for i in g[4:]]
            for i in g[4:] :
                matches[i, -1] = g[:3] + ids
    ids = { k[0] for k, v in usedMatches.items() if v == 1 }
    matches = matches[np.array(list(ids))]
    return matches


def cigar2score(data) :
    cigar, rSeq, qSeq, frame, mode, gapOpen, gapExtend, table_id = data
    if table_id == 4 :
        gtable[56] = 22
    frame = (frame-1) % 3
    gap, rBlk, qBlk = [], [], []
    qId, rId = 0, 0
    for n, t in cigar :
        if t == 'M' :
            rBlk.append(rSeq[rId:rId+n])
            qBlk.append(qSeq[qId:qId+n])
            rId, qId = rId + n, qId + n
        else :
            if t == 'D' :
                gap.append(n) 
                rId += n
            elif t == 'I' :
                gap.append(n) 
                if mode > 1 :
                    qBlk.append(qSeq[qId:qId+n])
                    rBlk.append([-1]*n)
                qId += n
    nGap, bGap, mGap = len(gap), np.sum(gap), np.sum([g for g in gap if g > 3])
    qAln = np.concatenate(qBlk)
    rAln = np.concatenate(rBlk)
    if mode == 1 :
        nMatch = np.sum(qAln == rAln)
        nMismatch = qAln.size - nMatch
        return float(nMatch)/(nMatch + nMismatch+bGap-mGap), nMatch*3 - nMismatch*1 - nGap*(gapOpen-gapExtend) - (bGap)*gapExtend
    else :
        qAln, rAln = qAln[frame:], rAln[frame:]
        if qAln.size % 3 :
            qAln, rAln = qAln[:-(qAln.size % 3)], rAln[:-(qAln.size % 3)]
        qAln, rAln = qAln.reshape(-1, 3), rAln.reshape(-1, 3)
        if mode == 3 :
            match = (qAln == rAln)
            nMatch = np.sum(np.sum(match, 0) * (9./7., 9./7., 3./7.))
            nMismatch = np.sum(rAln >= 0) - nMatch
            return float(nMatch)/(nMatch + nMismatch+bGap-mGap), nMatch*3 - nMismatch*1 - nGap*(gapOpen-gapExtend) - (bGap)*gapExtend
        else :
            s = (~np.any(rAln < 0, 1), )
            qAln, rAln = qAln[s], rAln[s]
            qCodon = np.sum(qAln * (25,5,1), 1)
            rCodon = np.sum(rAln * (25,5,1), 1)
            qAA, rAA = gtable[qCodon], gtable[rCodon]
            nMatch = np.sum(qAA == rAA)*3.
            nTotal = qAA.size * 3. + bGap-mGap
            score = np.sum(blosum62[(qAA << 5) + rAA])
            return nMatch/nTotal, score - nGap*(gapOpen-gapExtend) - (bGap)*gapExtend
nucEncoder = np.repeat(2, 255).astype(int)
nucEncoder[(np.array(['A', 'C', 'G', 'T']).view(asc2int),)] = (0, 1, 3, 4)
gtable = np.array(list('KNXKNTTXTTXXXXXRSXRSIIXMIQHXQHPPXPPXXXXXRRXRRLLXLLXXXXXXXXXXXXXXXXXXXXXXXXXEDXEDAAXAAXXXXXGGXGGVVXVVXYXXYSSXSSXXXXXXCXWCLFXLF')).view(asc2int).astype(int)-65

def poolBlast(params) :
    def parseBlast(fn, min_id, min_cov, min_ratio) :
        try:
            blastab = pd.read_csv(fn, sep='\t',header=None, dtype=str)
        except :
            return None
        blastab[[2, 10]] = blastab[[2, 10]].astype(float)
        blastab[[3, 4, 5, 6, 7, 8, 9, 11, 12, 13]] = blastab[[3, 4, 5, 6, 7, 8, 9, 11, 12, 13]].astype(int)
        blastab[2] /= 100.
        blastab = blastab[(blastab[2] >= min_id) & (blastab[7]-blastab[6]+1 >= min_cov) & (blastab[7]-blastab[6]+1 >= min_ratio*blastab[12]) ]
        if blastab.shape[0] <= 0 :
            return None
        else :
            blastab[14] = list(map(getCIGAR, zip(blastab[15], blastab[14])))
        blastab = blastab.drop(columns=[15])
        #blastab[[0, 1]] = blastab[[0, 1]].astype(str)
        return blastab
    
    blastn, refDb, qry, min_id, min_cov, min_ratio = params
    outfile = '{0}.bsn'.format(qry)
    blast_cmd = '{blastn} -db {refDb} -query {qry} -word_size 17 -out {qry}.bsn -perc_identity {min_id} -outfmt "6 qseqid sseqid pident length mismatch gapopen qstart qend sstart send evalue score qlen slen qseq sseq" -qcov_hsp_perc {min_ratio} -num_alignments 1000 -task blastn -evalue 1e-2 -dbsize 5000000 -reward 2 -penalty -3 -gapopen 6 -gapextend 2'.format(
        blastn=blastn, refDb=refDb, qry=qry, min_id=min_id*100, min_ratio=min_ratio*100)
    Popen(blast_cmd, stdout=PIPE, shell=True, universal_newlines=True).communicate()
    if os.path.getsize(outfile) > 0 :
        blastab = parseBlast(outfile, min_id, min_cov, min_ratio)
        os.unlink(outfile)
    else :
        blastab = None
    if blastab is None :
        return None

    blastab[14] = [[list(t) for t in tab] for tab in blastab[14].tolist()]
    np.save(qry+'.match.npy', blastab.values, allow_pickle=True)
    return qry + '.match.npy'


def getCIGAR(data) :
    ref, qry = data
    if qry.find('-') < 0 and ref.find('-') < 0 :
        cigar = [[len(qry), 'M']] 
    else :
        tag = np.array(['M', 'I', 'D'])
        cigar = np.concatenate([[-1], (np.array(list(qry)) == '-')*2 + (np.array(list(ref)) == '-'), [-1]])
        pos = np.where(np.diff(cigar) != 0)[0]
        cigar = [ list(v) for v in zip(np.diff(pos), tag[cigar[pos[:-1]+1]]) ]
    return cigar


class RunBlast(object) :
    def __init__(self) :
        self.qrySeq = self.refSeq = None
    def run(self, ref, qry, methods, min_id, min_cov, min_ratio, table_id=11, n_thread=8, useProcess=False, re_score=0, filter=[False, 0.9, 0.], linear_merge=[False, 300.,1.2], return_overlap=[True, 300, 0.6], fix_end=[6., 6.]) :
        tools = dict(blastn=self.runBlast, diamond=self.runDiamond, diamondself=self.runDiamondSELF)
        self.min_id = min_id
        self.min_cov = min_cov
        self.min_ratio = min_ratio
        self.table_id = table_id
        self.n_thread = n_thread
        if useProcess == True :
            self.pool = Pool(n_thread)
        elif useProcess == False :
            self.pool = ThreadPool(n_thread)
        else :
            self.pool = useProcess

        blastab = []
        self.dirPath = tempfile.mkdtemp(prefix='NS_', dir='.')
        try :
            for method in methods :
                if method.lower() in tools :
                    blastab.append(tools[method.lower()](ref, qry))
            blastab = [b for b in blastab if b.shape[0] > 0]
        except :
            import traceback
            print(traceback.print_exc())
        finally :
            shutil.rmtree(self.dirPath)
            if blastab :
                blastab = np.vstack(blastab)
                blastab = np.hstack([blastab, np.arange(blastab.shape[0], dtype=int)[:, np.newaxis]])
            else :
                if return_overlap[0] :
                    return np.empty([0, 16], dtype=object), np.empty([0, 3], dtype=int)
                else :
                    return np.empty([0, 16], dtype=object)
        if useProcess != self.pool :
            self.pool.close()
        
        if re_score :
            blastab=self.reScore(ref, qry, blastab, re_score, self.min_id, self.table_id)
        if filter[0] :
            blastab=self.ovlFilter(blastab, filter)
        if linear_merge[0] :
            blastab=self.linearMerge(blastab, linear_merge)
        self.fixEnd(blastab, *fix_end)
        if return_overlap[0] :
            overlap = self.returnOverlap(blastab, return_overlap)
            blastab = pd.DataFrame(blastab).sort_values([0,1,11]).values
            return blastab, overlap
        else :
            blastab = pd.DataFrame(blastab).sort_values([0,1,11]).values
            return blastab
            
    def returnOverlap(self, blastab, param) :
#        logger('Calculate overlaps.')
        
        ovl_l, ovl_p = param[1:]
        contigs = { tab[1]:id for id, tab in enumerate(blastab) }
        tabs = [ [contigs[tab[1]], tab[15]] + sorted( [tab[8], tab[9]] ) for tab in blastab ]
        tabs = np.array(sorted(tabs, key=itemgetter(0, 2, 3)), dtype=int)
        overlaps = np.empty(shape=[1000001, 3], dtype=int)
        overlaps[-1, :] = [0, 1, -1]
        res = []
        while overlaps[-1, 0] >= 0 :
            #logger('Searching {0} / {1} tabs'.format(overlaps[-1, 0], len(tabs)))
            overlaps[:-1, :] = -1
            overlaps = tab2overlaps(tabs, ovl_l, ovl_p, len(tabs), overlaps)
            res.append(overlaps[overlaps.T[2] > 0][:])
        res = np.vstack(res)
 #       logger('Identified {0} overlaps.'.format(len(res)))
        return res
    
    def reScore(self, ref, qry, blastab, mode, min_id, table_id=11, perBatch=10000) :
        if not self.qrySeq :
            self.qrySeq, self.qryQual = readFastq(qry)
        if not self.refSeq :
            self.refSeq, self.refQual = readFastq(ref)
        for k, v in self.qrySeq.items() :
            self.qrySeq[k] = nucEncoder[np.array(list(v)).view(asc2int)]
        for k, v in self.refSeq.items() :
            self.refSeq[k] = nucEncoder[np.array(list(v)).view(asc2int)]
        
        nTab = len(blastab)
        for bId in xrange(0, blastab.shape[0], perBatch) :
            #logger('Update scores: {0} / {1}'.format(bId, nTab))
            tabs = blastab[bId:bId+perBatch]
            #scores = np.array([ cigar2score([t[14], self.refSeq[str(t[1])][t[8]-1:t[9]] if t[8] < t[9] else 4 - self.refSeq[str(t[1])][t[9]-1:t[8]][::-1], self.qrySeq[str(t[0])][t[6]-1:t[7]], t[6], mode, 6, 1]) for t in tabs ])
            scores = np.array(list(map(cigar2score, ( [t[14], self.refSeq[str(t[1])][t[8]-1:t[9]] if t[8] < t[9] else 4 - self.refSeq[str(t[1])][t[9]-1:t[8]][::-1], self.qrySeq[str(t[0])][t[6]-1:t[7]], t[6], mode, 6, 1, table_id] for t in tabs ))))
            tabs.T[2], tabs.T[11] = np.round(scores.T, 3)
        blastab = blastab[blastab.T[2] >= min_id]
        return blastab

    def ovlFilter(self, blastab, params) :
        coverage, delta = params[1:]
#        logger('Run filtering. Start with {0} hits.'.format(len(blastab)))
        blastab[blastab.T[8] > blastab.T[9], 8:10] *= -1

        blastab = pd.DataFrame(blastab).sort_values(by=[1,0,8,6]).values
        for i, t1 in enumerate(blastab) :
            if t1[2] < 0 : continue
            toDel = []
            for j in xrange(i+1, blastab.shape[0]) :
                t2 = blastab[j]
                if t2[2] < 0 : continue
                if np.any(t1[:2] != t2[:2]) or t1[9] < t2[8] :
                    break
                c = min(t1[9], t2[9]) - t2[8] + 1
                if (c >= coverage*(t1[9]-t1[8]+1) and t2[11] - t1[11] >= delta) :
                    t1[2] = -1.
                    break
                elif (c >= coverage*(t2[9]-t2[8]+1) and t1[11] - t2[11] >= delta) :
                    toDel.append(j)
                elif c >= (t1[9]-t1[8]+1) and c < coverage*(t2[9]-t2[8]+1) :
                    c2 = min(t1[7], t2[7]) - max(t2[6], t1[6]) + 1
                    if c2 >= (t1[7]-t1[6]+1) and c2 < coverage*(t2[7]-t2[6]+1) :
                        t1[2] == -1
                        break
                elif c >= (t2[9]-t2[8]+1) and c < coverage*(t1[9]-t1[8]+1) :
                    c2 = min(t1[7], t2[7]) - max(t2[6], t1[6]) + 1
                    if c2 >= (t2[7]-t2[6]+1) and c2 < coverage*(t1[7]-t1[6]+1) :
                        toDel.append(j)
            if t1[2] >= 0 :
                for j in toDel :
                    blastab[j][2] = -1.
        blastab = blastab[blastab.T[2] >= 0]
        blastab[blastab.T[8] < 0, 8:10] *= -1
#        logger('Done filtering. End with {0} hits.'.format(blastab.shape[0]))
        return blastab
    def linearMerge(self, blastab, params) :
#        logger('Start merging neighboring regions.')
        blastab[blastab.T[8] > blastab.T[9], 8:10] *= -1
        blastab = pd.DataFrame(blastab).sort_values([0,1,8,6]).values
        blastab = np.vstack(list(map(_linearMerge, [[matches, params] for matches in np.split(blastab, np.where(np.diff(np.unique(blastab.T[0], return_inverse=True)[1]))[0]+1 )])))
        blastab[blastab.T[8] < 0, 8:10] *= -1
 #       logger('Finish merging neighboring regions.')
        return blastab

    def fixEnd(self, blastab, se, ee) :
        for p in blastab :
            e1, e2 = p[6] - 1, p[12] - p[7]
            cigar = p[14]
            if p[9] > p[8] :
                if 0 < e1 <= se :
                    d = min(p[6]-1, p[8]-1)
                    p[6], p[8], cigar[0][0] = p[6]-d, p[8]-d, cigar[0][0]+d
                if 0 < e2 <= ee :
                    d = min(p[12]-p[7], p[13]-p[9])
                    p[7], p[9], cigar[-1][0] = p[7]+d, p[9]+d, cigar[-1][0]+d
            else :
                if 0 < e1 <= se :
                    d = min(p[6]-1, p[13]-p[8])
                    p[6], p[8], cigar[0][0] = p[6]-d, p[8]+d, cigar[0][0]+d
                if 0 < e2 <= ee :
                    d = min(p[12]-p[7], p[9]-1)
                    p[7], p[9], cigar[-1][0] = p[7]+d, p[9]-d, cigar[-1][0]+d
            p[14] = ''.join( '{0}{1}'.format(n, t) for n, t in cigar )

    def runBlast(self, ref, qry) :
        logger('Run BLASTn starts')
        if not self.qrySeq :
            self.qrySeq, self.qryQual = readFastq(qry)
        if not self.refSeq :
            self.refSeq, self.refQual = readFastq(ref)
        refDb = refNA = os.path.join(self.dirPath, 'refNA')
        with open(refNA, 'w') as fout :
            for n,s in self.refSeq.items() :
                fout.write('>{0}\n{1}\n'.format(n, s))
        Popen('{makeblastdb} -dbtype nucl -in {refNA} -out {refDb}'.format(makeblastdb=makeblastdb, refNA=refNA, refDb = refDb).split(), stderr=PIPE, stdout=PIPE, universal_newlines=True).communicate()
        qrySeq = sorted(list(self.qrySeq.items()), key=lambda s:-len(s[1]))
        qrys = [ os.path.join(self.dirPath, 'qryNA.{0}'.format(id)) for id in range(min(len(qrySeq), self.n_thread))]
        for id, q in enumerate(qrys) :
            with open(q, 'w') as fout :
                for n, s in qrySeq[id::self.n_thread] :
                    fout.write('>{0}\n{1}\n'.format(n, s))
        blastab = []
        for r in self.pool.imap_unordered(poolBlast, [ [blastn, refDb, q, self.min_id, self.min_cov, self.min_ratio] for q in qrys ]) :
            if r is not None :
                blastab.append(np.load(r, allow_pickle=True))
                os.unlink(r)
        if len(blastab) :
            blastab = np.vstack(blastab)
        else :
            blastab = np.empty([0, 15], dtype=object)
        logger('Run BLASTn finishes. Got {0} alignments'.format(blastab.shape[0]))
        return blastab

    def runDiamondSELF(self, ref, qry) :
        return self.runDiamond(ref, qry, nhits=200, frames='F')
    def runDiamond(self, ref, qry, nhits=10, frames='7') :
        logger('Run diamond starts')

        refAA = os.path.join(self.dirPath, 'refAA')
        qryAA = os.path.join(self.dirPath, 'qryAA')
        aaMatch = os.path.join(self.dirPath, 'aaMatch')
        
        if not self.qrySeq :
            self.qrySeq, self.qryQual = readFastq(qry)
        if not self.refSeq :
            self.refSeq, self.refQual = readFastq(ref)

        qryAASeq = transeq(self.qrySeq, frame='F', transl_table=self.table_id)
        with open(qryAA, 'w') as fout :
            for n, ss in sorted(qryAASeq.items()) :
                _, id, s = min([ (len(s[:-1].split('X')), id, s) for id, s in enumerate(ss) ])
                fout.write('>{0}:{1}\n{2}\n'.format(n, id+1, s))
        
        diamond_fmt = '{diamond} makedb --db {qryAA} --in {qryAA}'.format(
            diamond=diamond, qryAA=qryAA)
        p = Popen(diamond_fmt.split(), stderr=PIPE, stdout=PIPE, universal_newlines=True).communicate()
        
        refAASeq = transeq(self.refSeq, frames, transl_table=self.table_id)
        toWrite = []
        for n, ss in sorted(refAASeq.items()) :
            for id, s in enumerate(ss) :
                cdss = re.findall('.{1000,}?X|.{1,1000}$', s + 'X')
                cdss[-1] = cdss[-1][:-1]
                cdsi = np.cumsum([0]+list(map(len, cdss[:-1])))
                for ci, cs in zip(cdsi, cdss) :
                    if len(cs) :
                        toWrite.append('>{0}:{1}:{2}\n{3}\n'.format(n, id+1, ci, cs))
        
        for id in xrange(5) :
            with open('{0}.{1}'.format(refAA, id), 'w') as fout :
                for line in toWrite[id::5] :
                    fout.write(line)
            diamond_cmd = '{diamond} blastp --no-self-hits --threads {n_thread} --db {refAA} --query {qryAA} --out {aaMatch} --id {min_id} --query-cover {min_ratio} --evalue 1 -k {nhits} --dbsize 5000000 --outfmt 101'.format(
                diamond=diamond, refAA='{0}.{1}'.format(refAA, id), qryAA=qryAA, aaMatch='{0}.{1}'.format(aaMatch, id), n_thread=self.n_thread, min_id=self.min_id*100., nhits=nhits, min_ratio=self.min_ratio*100.)
            Popen(diamond_cmd.split(), stdout=PIPE, stderr=PIPE, universal_newlines=True).communicate()
        blastab = []
        for r in self.pool.imap_unordered(parseDiamond, [ ['{0}.{1}'.format(aaMatch, id), self.refSeq, self.qrySeq, self.min_id, self.min_cov, self.min_ratio] for id in xrange(5) ]) :
            if r is not None :
                blastab.append(np.load(r, allow_pickle=True))
                os.unlink(r)
        blastab = np.vstack(blastab)
        logger('Run diamond finishes. Got {0} alignments'.format(blastab.shape[0]))
        return blastab


def uberBlast(args, extPool=None) :
    import argparse
    parser = argparse.ArgumentParser(description='Five different alignment methods. ')
    parser.add_argument('-r', '--reference',     help='[INPUT; REQUIRED] filename for the reference. This is normally a genomic assembly. ', required=True)
    parser.add_argument('-q', '--query',  help='[INPUT; REQUIRED] filename for the query. This can be short-reads or genes or genomic assemblies. ', required=True)
    parser.add_argument('-o', '--output', help='[OUTPUT; Default: None] save result to a file or to screen (stdout). Default do nothing. ', default=None)
    parser.add_argument('--blastn',       help='Run BLASTn. Slowest. Good for identities between [70, 100]', action='store_true', default=False)
    parser.add_argument('--diamond',      help='Run diamond on tBLASTn mode. Fast. Good for identities between [30-100]', action='store_true', default=False)
    parser.add_argument('--diamondSELF',      help='Run diamond on tBLASTn mode. Fast. Good for identities between [30-100]', action='store_true', default=False)
    parser.add_argument('--gtable',       help='[DEFAULT: 11] genetic table to use. 11 for bacterial genomes and 4 for Mycoplasma', default=11, type=int)
    
    parser.add_argument('--min_id', help='[DEFAULT: 0.3] Minimum identity before reScore for an alignment to be kept', type=float, default=0.3)
    parser.add_argument('--min_cov', help='[DEFAULT: 40] Minimum length for an alignment to be kept', type=float, default=40.)
    parser.add_argument('--min_ratio', help='[DEFAULT: 0.05] Minimum length for an alignment to be kept, proportional to the length of the query', type=float, default=0.05)

    parser.add_argument('-s', '--re_score', help='[DEFAULT: 0] Re-interpret alignment scores and identities. 0: No rescore; 1: Rescore with nucleotides; 2: Rescore with amino acid; 3: Rescore with codons', type=int, default=0)
    parser.add_argument('-f', '--filter', help='[DEFAULT: False] Remove secondary alignments if they overlap with any other regions', default=False, action='store_true')
    parser.add_argument('--filter_cov', help='[DEFAULT: 0.9] ', default=0.9, type=float)
    parser.add_argument('--filter_score', help='[DEFAULT: 0] ', default=0., type=float)
    parser.add_argument('-m', '--linear_merge', help='[DEFAULT: False] Merge consective alignments', default=False, action='store_true')
    parser.add_argument('--merge_gap', help='[DEFAULT: 600] ', default=600., type=float)
    parser.add_argument('--merge_diff', help='[DEFAULT: 1.5] ', default=1.5, type=float)
    parser.add_argument('-O', '--return_overlap', help='[DEFAULT: False] Report overlapped alignments', default=False, action='store_true')
    parser.add_argument('--overlap_length', help='[DEFAULT: 300] Minimum overlap to report', default=300, type=float)
    parser.add_argument('--overlap_proportion', help='[DEFAULT: 0.6] Minimum overlap proportion to report', default=0.6, type=float)
    parser.add_argument('-e', '--fix_end', help='[FORMAT: L,R; DEFAULT: 0,0] Extend alignment to the edges if the un-aligned regions are <= [L,R] basepairs.', default='0,0')
    parser.add_argument('-t', '--n_thread', help='[DEFAULT: 8] Number of threads to use. ', type=int, default=1)
    parser.add_argument('-p', '--process', help='[DEFAULT: False] Use processes instead of threads. ', action='store_true', default=False)
    
    args = parser.parse_args(args)
    if extPool is not None :
        args.process =extPool
    methods = []
    for method in ('blastn', 'diamond', 'diamondSELF') :
        if args.__dict__[method] :
            methods.append(method)
    for opt in ('fix_end',) :
        args.__dict__[opt] = args.__dict__[opt].split(',')
        args.__dict__[opt][-2:] = list(map(float, args.__dict__[opt][-2:]))
    data = RunBlast().run(args.reference, args.query, methods, args.min_id, args.min_cov, args.min_ratio, args.gtable, args.n_thread, args.process, args.re_score, \
                             [args.filter, args.filter_cov, args.filter_score], \
                             [args.linear_merge, args.merge_gap, args.merge_diff], \
                             [args.return_overlap, args.overlap_length, args.overlap_proportion], \
                             args.fix_end)
    if args.output :
        fout = sys.stdout if args.output.upper() == 'STDOUT' else open(args.output, 'w')
        for t in data :
            fout.write ('\t'.join([str(tt) for tt in t ]) + '\n')
        fout.close()
    return data

if __name__ == '__main__' :
    blastab = uberBlast(sys.argv[1:])
```
