## Supplemental Code S1 for "Accurate reconstruction of bacterial pan- and core- genomes with PEPPAN": PEPPAN.py

```
#!/usr/bin/env python
import os, re, sys, shlex, ete3, tempfile, hashlib, shutil
import subprocess, numpy as np, pandas as pd, numba as nb
from operator import itemgetter
import zipfile, io
from multiprocessing import Pool, Manager, Process
from collections import defaultdict
try:
    from modules.configure import externals, logger, rc, transeq, readFasta, uopen, xrange, asc2int
    from modules.clust import getClust
    from modules.uberBlast import uberBlast
except:
    from .modules.configure import externals, logger, rc, transeq, readFasta, uopen, xrange, asc2int
    from .modules.clust import getClust
    from .modules.uberBlast import uberBlast

params = dict(
    ml = '{fasttree} -nt -gtr -pseudo {0}', 
    nj = '{rapidnj} -i fa -t d {0}', 
)

def in1d(arr1, arr2, invert=False) :
    darr2 = set(arr2)
    res = np.array([n in darr2 for n in np.array(arr1).flatten()])
    return ~res if invert else res

class MapBsn(object) :
    def __init__(self, fname, mode='r') :
        self.fname = fname
        self.mode = mode
        self.conn = zipfile.ZipFile(self.fname, mode=mode, compression=zipfile.ZIP_DEFLATED,allowZip64=True)
        self.namelist = set(self.conn.namelist())
    def __enter__(self) :
        return self
    def __exit__(self, type, value, traceback) :
        self.conn.close()
    def get(self, key, default=[]) :
        key = str(key)
        if self.exists(key) :
            buf = io.BytesIO(self.conn.read(key))
            return np.lib.npyio.format.read_array(buf, allow_pickle=True)
        else :
            return default
        
    def __getitem__(self, key) :
        return self.get(key)

    def exists(self, key) :
        return str(key) in self.namelist

    def keys(self) :
        return self.namelist

    def items(self) :
        for key in self.namelist :
            yield key, self.get(key)

    def values(self) :
        for key in self.namelist :
            yield self.get(key)

    def delete(self, key) :
        self.namelist -= {str(key)}

    def delete_real(self, key) :
        key = str(key)
        if key in self.namelist :
            self.namelist -= {key}
            self.conn.close()
            subprocess.Popen(['zip', '-d', self.fname, key]).wait()
            self.conn = zipfile.ZipFile(self.fname, mode='w', compression=zipfile.ZIP_DEFLATED,allowZip64=True)
        
    def pop(self, key, default=[]) :
        val = self.get(key, default)
        self.delete(key)
        return val
    
    def size(self) :
        return len(self.namelist)
        
    def save(self, key, val) :
        key = str(key)
        self.delete_real(key)
        self._save(self.conn, key, val)
        self.namelist |= {key}

    def _save(self, db, key, val) :
        buf = io.BytesIO()
        np.lib.npyio.format.write_array(buf,
                                        np.asanyarray(val),
                                        allow_pickle=True)
        db.writestr(key, buf.getvalue())
        
    def update(self, dataset) :
        newList = set([])
        tmpName = self.fname[:-4] + '.tmp.npz'
        tmp = zipfile.ZipFile(tmpName, mode='w', compression=zipfile.ZIP_DEFLATED,allowZip64=True)
        for d in dataset :
            key = str(d[0][0])
            newList.add(key)
            oldData = self.get(key)
            data = np.vstack([oldData, d]) if len(oldData) else d
            self._save(tmp, key, data)
        for key in self.keys() :
            if key not in newList :
                data = self.get(key)
                if len(data) :
                    newList.add(key)
                    self._save(tmp, key, data)
        tmp.close()
        self.conn.close()
        self.namelist = newList
        os.rename(tmpName, self.fname)
        self.conn = zipfile.ZipFile(self.fname, mode='a', compression=zipfile.ZIP_DEFLATED,allowZip64=True)
        

def iter_readGFF(data) :
    fname, feature, gtable = data
    seq, cds = {}, {}
    names = {}
    fnames = fname.split(',')
    fname = fnames[0]
    fprefix = os.path.basename(fname).split('.')[0]

    for fn in fnames :
        with uopen(fn) as fin :
            sequenceMode = False
            for line in fin :
                if line.startswith('#') : 
                    continue
                elif line.startswith('>') :
                    sequenceMode = True
                    name = line[1:].strip().split()[0]
                    cname = '{0}:{1}'.format(fprefix, name)
                    assert cname not in seq, logger('Error: duplicated sequence name {0}'.format(name))
                    seq[cname] = [fname, []]
                elif sequenceMode :
                    seq[cname][1].extend(line.strip().split())
                else :
                    part = line.strip().split('\t')
                    if len(part) > 2 :
                        name = re.findall(r'locus_tag=([^;]+)', part[8])
                        if len(name) == 0 :
                            parent = re.findall(r'Parent=([^;]+)', part[8])
                            if len(parent) and parent[0] in names :
                                name = names[parent[0]]
                        if len(name) == 0 :
                            name = re.findall(r'Name=([^;]+)', part[8])
                        if len(name) == 0 :
                            name = re.findall(r'ID=([^;]+)', part[8])
    
                        if part[2] == feature :
                            assert len(name) > 0, logger('Error: CDS has no name. {0}'.format(line))
                            #          source_file, seqName, Start,       End,      Direction, hash, Sequences
                            gname = '{0}:{1}'.format(fprefix, name[0])
                            if gname not in cds :
                                cds[gname] = [fname, '{0}:{1}'.format(fprefix, part[0]), int(part[3]), int(part[4]), part[6], 0, [], int(part[3]), int(part[4])]
                            elif part[0] == cds[gname][1] and fname == cds[gname][0] :
                                cds[gname].extend([int(part[3]), int(part[4])])
                                cds[gname][3] = max(cds[gname][3], int(part[4]))
                        else :
                            ids = re.findall(r'ID=([^;]+)', part[8])
                            if len(ids) :
                                names[ids[0]] = name

    for n in seq :
        seq[n][1] = ''.join(seq[n][1]).upper()
    for n, c in cds.items() :
        try:
            for i in np.arange(7, len(c), 2) :
                s = seq[c[1]][1][(c[i]-1) : c[i+1]]
                c[6].append(s)
            c[6] = ''.join([ rc(s) for s in reversed(c[6]) ]) if c[4] == '-' else ''.join(c[6])
            pcode = checkPseu(n, c[6], gtable)
            if pcode :
                c[5], c[6] = pcode, ''
            else :
                c[5] = int(hashlib.sha1(c[6].encode('utf-8')).hexdigest(), 16)
        except :
            c[5], c[6] = 6, ''
        cds[n][:] = c[:7]
    return seq, cds

def readGFF(fnames, feature, gtable) :
    if not isinstance(fnames, list) : fnames = [fnames]
    seq, cds = {}, {}
    #for ss, cc in map(iter_readGFF, [[fn, feature, gtable] for fn in fnames]) :
    for ss, cc in pool.imap_unordered(iter_readGFF, [[fn, feature, gtable] for fn in fnames]) :
        seq.update(ss)
        cds.update(cc)
    return seq, cds


def get_similar_pairs(clust, priorities, params) :
    def get_similar(bsn, ortho_pairs) :
        key = tuple(sorted([bsn[0][0], bsn[0][1]]))
        if key in ortho_pairs :
            return
        if min(int(bsn[0][13]), int(bsn[0][12])) * 20 <= max(int(bsn[0][13]), int(bsn[0][12])) :
            return
        matched_aa = {}
        for part in bsn :
            s_i, e_i, s_j, e_j = [ int(x) for x in part[6:10] ]
            for s, t in re.findall(r'(\d+)([A-Z])', part[14]) :
                s, frame_i, frame_j = int(s), s_i % 3, s_j % 3
                if t == 'M' :
                    if frame_i == frame_j or 'f' in params['incompleteCDS'] :
                        matched_aa.update({ (s_i+x): part[2] for x in xrange( (3 - (frame_i - 1))%3, s )})
                    s_i += s
                    s_j += s
                    if len(matched_aa)*3 >= min(params['match_len2'], params['match_len'], params['match_len1']) and len(matched_aa)*3 >= min(params['match_prop'], params['match_prop1'], params['match_prop2']) * int(bsn[0][12]) :
                        ave_iden = int(np.mean(list(matched_aa.values()))*10000)
                        if ave_iden >= params['match_identity']*10000 :
                            if len(matched_aa)*3 >= min(max(params['match_len'], params['match_prop']* min(int(bsn[0][13]), int(bsn[0][12]))), 
                                                        max(params['match_len1'], params['match_prop1']* min(int(bsn[0][13]), int(bsn[0][12]))), 
                                                        max(params['match_len2'], params['match_prop2']* min(int(bsn[0][13]), int(bsn[0][12]))) ) :
                                ortho_pairs[key] = ave_iden
                            else :
                                ortho_pairs[key] = 0
                            return
                elif t == 'I' :
                    s_i += s
                else :
                    s_j += s
    if params['noDiamond'] :
        self_bsn = uberBlast('-r {0} -q {0} --blastn --min_id {1} --min_cov {2} -t {3} --min_ratio {4} -e 3,3 -p --gtable {5}'.format(\
            clust, params['match_identity'] - 0.05, params['match_frag_len'], params['n_thread'], params['match_frag_prop'], params['gtable']).split(), pool)
    else :
        self_bsn = uberBlast('-r {0} -q {0} --blastn --diamondSELF -s 1 --min_id {1} --min_cov {2} -t {3} --min_ratio {4} -e 3,3 -p --gtable {5}'.format(\
            clust, params['match_identity'] - 0.05, params['match_frag_len'], params['n_thread'], params['match_frag_prop'], params['gtable']).split(), pool)
    self_bsn.T[:2] = self_bsn.T[:2].astype(int)
    presence, ortho_pairs = {}, {}
    save = []
    
    cluGroups = []
    for part in self_bsn :
        if part[0] not in presence :
            presence[part[0]] = 1
        elif presence[part[0]] == 0 :
            continue
        iden, qs, qe, ss, se, ql, sl = float(part[2]), float(part[6]), float(part[7]), float(part[8]), float(part[9]), float(part[12]), float(part[13])
        if presence.get(part[1], 1) == 0 or ss >= se :
            continue
        if ss < se and part[0] != part[1] and iden >= params['clust_identity'] and qs%3 == ss%3 and (ql-qe)%3 == (sl-se)%3 :
            if ql <= sl :
                if qe - qs + 1 >= np.sqrt(params['clust_match_prop']) * sl and priorities[part[0]][0] >= priorities[part[1]][0] :
                    cluGroups.append([int(part[1]), int(part[0]), int(iden*10000.)])
                    presence[part[0]] = 0
                    continue
            elif se - ss + 1 >= np.sqrt(params['clust_match_prop']) * ql and priorities[part[0]][0] <= priorities[part[1]][0] :
                cluGroups.append([int(part[0]), int(part[1]), int(iden*10000.)])
                presence[part[1]] = 0
                continue
                
        if len(save) > 0 and np.any(save[0][:2] != part[:2]) :
            if len(save) >= 50 :
                presence[save[0][1]] = 0
            elif save[0][0] != save[0][1] :
                get_similar(save, ortho_pairs)
            save = []
        save.append(part)
    if len(save) > 0 :
        if len(save) >= 50 :
            presence[save[0][1]] = 0
        elif save[0][0] != save[0][1] :
            get_similar(save, ortho_pairs)
    
    toWrite = []
    with uopen(params['clust'], 'r') as fin :
        for line in fin :
            if line.startswith('>') :
                name = line[1:].strip().split()[0]
                write= True if presence.get(int(name), 0) > 0 else False
            if write :
                toWrite.append(line)
    with open(params['clust'], 'w') as fout :
        for line in toWrite :
            fout.write(line)
    if len(cluGroups) :
        clu = np.load(params['clust'].rsplit('.',1)[0] + '.npy', allow_pickle=True)
        clu = np.vstack([clu, cluGroups])
        clu = clu[np.argsort(-clu.T[2])]
        np.save(params['clust'].rsplit('.',1)[0] + '.npy', clu)
    return np.array([[k[0], k[1], v] for k, v in ortho_pairs.items()], dtype=int)

@nb.jit('i8[:,:,:](u1[:,:], i8[:,:,:])', nopython=True)
def compare_seq(seqs, diff) :
    for id in np.arange(seqs.shape[0]) :
        s = seqs[id]
        c = (s > 0) * (seqs[(id+1):] > 0)
        n_comparable = np.sum(c, 1) + 2
        n_diff = np.sum(( s != seqs[(id+1):] ) & c, 1) + 1
        diff[id, id+1:, 0] = n_diff
        diff[id, id+1:, 1] = n_comparable
    return diff

@nb.jit('i8[:,:,:](u1[:,:], i8[:,:,:])', nopython=True)
def compare_seqX(seqs, diff) :
    for id in (0, seqs.shape[0]-1) :
        s = seqs[id]
        c = (s > 0) * (seqs > 0)
        n_comparable = np.sum(c, 1) + 2
        n_diff = np.sum(( s != seqs ) & c, 1) + 1
        diff[id, :, 0] = n_diff
        diff[id, :, 1] = n_comparable
    return diff


def decodeSeq(seqs) :
    ori_seqs = np.zeros([seqs.shape[0], seqs.shape[1]*3], dtype=np.uint8)
    ori_seqs[:, :seqs.shape[1]] = (seqs/25).astype(np.uint8)
    ori_seqs[:, seqs.shape[1]:seqs.shape[1]*2] = (np.mod(seqs, 25)/5).astype(np.uint8)
    ori_seqs[:, seqs.shape[1]*2:seqs.shape[1]*3] = np.mod(seqs, 5)
    return ori_seqs

def filt_per_group(data) :
    mat, inparalog, ref_len, seq_file, global_file = data
    global_differences = dict(np.load(global_file, allow_pickle=True))
    nMat = mat.shape[0]
    with MapBsn(seq_file) as conn :
        seqs = np.array([ conn.get(int(id/1000))[id%1000] for id in mat.T[5].tolist() ])
    seqs = np.array([45, 65, 67, 71, 84], dtype=np.uint8)[decodeSeq(seqs)][:, :ref_len]
    seqs[in1d(seqs, [65, 67, 71, 84], invert=True).reshape(seqs.shape)] = 0

    def checkDiv(mat, diffX, i1) :
        m1 = mat[i1]
        for i2, m2 in enumerate(mat) :
            if i1 != i2 :
                mut, aln = diffX[i1, i2]
                gd = (np.max([params['self_id'], 2.0/aln]), 0.) if m1[1] == m2[1] else global_differences.get(tuple(sorted([m1[1], m2[1]])), (0.5, 0.6))
                dX = mut/aln/(gd[0]*np.exp(gd[1]*np.sqrt(params['allowed_sigma'])))
                if dX > 1 :
                    return True
        return False

    diffX = compare_seqX(seqs, np.zeros(shape=[seqs.shape[0], seqs.shape[0], 2], dtype=int)).astype(float)
    isDiv = False
    for i1 in (0, mat.shape[0]-1) :
        isDiv = checkDiv(mat, diffX, i1)
        if isDiv :
            break
    if inparalog and not isDiv :
        from collections import defaultdict
        dupGenome = defaultdict(list)
        for idx, g in enumerate(mat.T[1]):
            dupGenome[g].append(idx)
        dupGenome = [v for v in dupGenome.values() if len(v) > 1]
        for group in dupGenome :
            m = mat[group]
            diffX = compare_seqX(seqs[group], np.zeros(shape=[len(group), len(group), 2], dtype=int)).astype(float)
            for i1 in (0, m.shape[0] - 1):
                isDiv = checkDiv(m, diffX, i1)
                if isDiv:
                    break
            if isDiv :
                break
    if not isDiv :
        return [mat]
    
    diff = compare_seq(seqs, np.zeros(shape=[seqs.shape[0], seqs.shape[0], 2], dtype=int)).astype(float)
    distances = np.zeros(shape=[mat.shape[0], mat.shape[0], 2], dtype=float)
    for i1, m1 in enumerate(mat) :
        for i2 in xrange(i1+1, nMat) :
            m2 = mat[i2]
            mut, aln = diff[i1, i2]
            
            gd = (np.max([params['self_id'], 2.0/aln]), 0.) if m1[1] == m2[1] else global_differences.get(tuple(sorted([m1[1], m2[1]])), (0.5, 0.6))
            d = mut/aln/(gd[0]*np.exp(gd[1]*params['allowed_sigma']))
            distances[i1, i2, :] = [d/gd[0], 1/gd[0]]
            distances[i2, i1, :] = distances[i1, i2, :]
    
    if np.any(distances[:, :, 0] > distances[:, :, 1]) :
        groups = []
        for j, m in enumerate(mat) :
            novel = 1
            for g in groups :
                if diff[g[0], j, 0] <= 0.01*diff[g[0], j, 1] : 
                    g.append(j)
                    novel = 0
                    break
            if novel :
                groups.append([j])
        group_size = {g[0]:len(g) for g in groups}
        seqs[seqs == 0] = 45
        try :
            tags = {g[0]:seqs[g[0]].tostring().decode('ascii') for g in groups}
        except :
            tags = {g[0]:seqs[g[0]].tostring() for g in groups}
            
        incompatible = np.zeros(shape=distances.shape, dtype=float)
        for i1, g1 in enumerate(groups) :
            for i2 in range(i1+1, len(groups)) :
                g2 = groups[i2]
                incompatible[g2[0], g1[0], :] = incompatible[g1[0], g2[0], :] = np.sum(distances[g1][:, g2, :], (0, 1))
        
        if np.all(incompatible[:,:,0] <= incompatible[:,:,1]) :
            return [mat]

        for ite in xrange(3) :
            try :
                tmpFile = tempfile.NamedTemporaryFile(dir='.', delete=False)
                for n, s in tags.items() :
                    tmpFile.write('>X{0}\n{1}\n{2}'.format(n, s, '\n'*ite).encode('utf-8'))
                tmpFile.close()
                cmd = params[params['orthology']].format(tmpFile.name, **params) if len(tags) < 500 else params['nj'].format(tmpFile.name, **params)
                phy_run = subprocess.Popen(shlex.split(cmd), stdin=subprocess.PIPE, stdout=subprocess.PIPE, stderr=subprocess.PIPE, universal_newlines=True)
                gene_phy = ete3.Tree(phy_run.communicate()[0].replace("'", ''))
                break
            except :
                if ite == 2 :
                    return [mat]
            finally:
                os.unlink(tmpFile.name)
        for n in gene_phy.get_leaves() :
            if len(n.name) :
                n.name = n.name[1:] 
        
        node = gene_phy.get_midpoint_outgroup()
        if node is not None :
            gene_phy.set_outgroup(node)
        gene_phys = [gene_phy]
        id = 0
        while id < len(gene_phys) :
            gene_phy = gene_phys[id]
            for ite in xrange(3000) :
                all_tips = {int(t) for t in gene_phy.get_leaf_names()}

                if np.all(incompatible[list(all_tips)].T[0, list(all_tips)] <= incompatible[list(all_tips)].T[1, list(all_tips)]) :
                    break
                rdist = sum([c.dist for c in gene_phy.get_children()])
                for c in gene_phy.get_children() :
                    c.dist = rdist
                for node in gene_phy.iter_descendants('postorder') :
                    if node.is_leaf() :
                        node.leaves = { int(node.name) } if node.name != 'REF' else set([])
                    else :
                        node.leaves = { n  for child in node.get_children() for n in child.leaves }
                    node.leaf_size = np.sum([group_size[t] for t in node.leaves])
                    if len(node.leaves) : 
                        oleaves = all_tips - node.leaves
                        c = incompatible[list(node.leaves)].T[:, list(oleaves)]
                        ic = np.sum(c, (1,2))
                        node.ic = [ic[0]/max(1., ic[1]), np.sum(c[1, (c[0] > c[1])])]
                    else :
                        node.ic = [0., 0.]
                cut_node = [[n.ic[0]*np.sqrt(n.ic[1]), n.ic[0], n.dist, n] for n in gene_phy.iter_descendants('postorder') if n.ic[0] > 1]
                if len(cut_node) > 0 :
                    cut_node = max(cut_node, key=lambda x:(x[0], x[1], x[2]))[3]
                    prev_node = cut_node.up
                    cut_node.detach()
                    t2 = cut_node
                    if prev_node.is_root() :
                        gene_phy = gene_phy.get_children()[0]
                    else :
                        prev_node.delete(preserve_branch_length=True)
                    if np.min(np.array(gene_phy.get_leaf_names()).astype(int)) > np.min(np.array(t2.get_leaf_names()).astype(int)) :
                        gene_phy, t2 = t2, gene_phy
                    gene_phys[id] = gene_phy
                    if np.max(mat[np.array(t2.get_leaf_names()).astype(int), 4]) >= (params['clust_identity']-0.02)*10000 :
                        gene_phys.append(t2)
                else :
                    break
            id += 1
        mats = []
        for gene_phy in gene_phys :
            if len(gene_phy.get_leaf_names()) < len(tags) :
                g = {str(g[0]):g for g in groups}
                tips = sorted([ nn for n in gene_phy.get_leaf_names() for nn in g.get(n, [])])
                mats.append(mat[tips])
            else :
                mats.append(mat)
        return mats
    else :
        return [mat]

def get_gene(allScores, priorities, ortho_groups, cnt=1) :
    ranking = {gene:priorities.get(gene)[0] for gene in allScores.keys() if gene in priorities}
    if len(ranking) > 0 :
        min_rank = min(ranking.values())
        scores = { gene:allScores[gene] for gene, r in ranking.items() if r == min_rank }
    else :
        min_rank = -1
        scores = {}
    
    genes, all_useds = [], set([])
    nGene = len(scores)
    for id, (gene, score) in enumerate(sorted(scores.items(), key=itemgetter(1), reverse=True)) :
        if score <= 0 : break
        if gene not in all_useds or nGene < 2*cnt :
            genes.append([gene, score, min_rank])
            if len(genes) >= cnt :
                break
            all_useds.update(set(ortho_groups[ortho_groups.T[0] == int(gene), 1]))
    
    if len(genes) <= 0 :
        for gene in scores :
            allScores.pop(gene)
        return []
    return genes

def load_conflict(data) :
    cfl_file, idss = data
    conflicts = []
    with MapBsn(cfl_file) as cfl_conn :
        for ids in idss :
            gid = int(ids[0, 0]/30000)
            d = cfl_conn[gid]
            for id, g in ids :
                idx = id%30000
                idx1, idx2 = d[idx:(idx+2)]
                if idx1 < idx2 :
                    conflicts.append([g, id, d[idx1:idx2]])
    return conflicts

def filt_genes(groups, ortho_groups, global_file, cfl_file, priorities, scores, encodes) :
    ortho_groups = np.vstack([ortho_groups[:, :2], ortho_groups[:, [1,0]]])*1000
    priorities = { k*1000:v for k, v in priorities.items() }
    scores = { k*1000:v for k, v in scores.items() }
    conflicts, new_groups = {}, {}

    encodes = np.array([n for i, n in sorted([[i, n] for n, i in encodes.items()])])
    clust_ref = { int(n)*1000:s for n, s in readFasta(params['clust']).items()}
    used, pangenome, panList = {}, {}, {}
    lowest_p = max([v[0] for v in priorities.values()])    
    while len(scores) > 0 :
        # get top 500 genes
        ortho_groups = ortho_groups[np.all(in1d(ortho_groups, list(scores.keys())).reshape(ortho_groups.shape), 1)]
        genes = get_gene(scores, priorities, ortho_groups, cnt=500)
        if len(genes) <= 0 :
            continue
        to_run, (min_score, min_rank) = [], genes[-1][1:]
        genes = {gene:score for gene, score, min_rank in genes}

        minSet = len(genes)*0.5
        tmpSet = {}
        for gene, score in list(genes.items()) :
            present_iden = 0
            if gene not in new_groups :
                mat = groups.get(int(gene/1000))
                mat.T[4] = (10000 * mat.T[3]/np.max(mat.T[3])).astype(int)
                for m in mat :
                    v = used.get(m[5], None)
                    if v is None :
                        present_iden = max(present_iden, m[4])
                    else :
                        m[3] = 0 if v > 0 else -m[3]
                if present_iden < (params['clust_identity']-0.02)*10000 :
                    genes.pop(gene, None)
                    scores.pop(gene, None)
                    conflicts.pop(gene, None)
                else :
                    tmpSet[gene] = mat[mat.T[3] != 0]
        if len(genes) < minSet :
            continue

        logger('Selected {0} genes after initial checking'.format(len(genes)))
        conflicts.update({gene:{} for gene in tmpSet.keys()})
        if len(tmpSet):
            tab_ids = np.vstack([ mat[:, (5, 0)] for mat in tmpSet.values() ])
            tab_ids = tab_ids[np.argsort(tab_ids.T[0])]
            tab_ids = np.split(tab_ids, np.cumsum(np.unique((tab_ids.T[0]/30000).astype(int), return_counts=True)[1])[:-1])
            chunk_size = float(len(tab_ids))/params['n_thread']
            tab_ids = [tab_ids[int(i):int(i + chunk_size)] for i in np.arange(params['n_thread'], dtype=float) * chunk_size if \
                       int(i) < int(i + chunk_size)]
            for cfl in pool2.imap_unordered(load_conflict, [ [cfl_file, ids] for ids in tab_ids ]) :
                for g, i, c in cfl :
                    conflicts[g*1000][i] = c
        
        used2 = set([])
        for gene, score in sorted(genes.items(), key=lambda x:-x[1]) :
            if gene not in new_groups :
                mat = tmpSet.get(gene)
                unsure = np.sum([ 1 for m in mat if m[3] > 0 and m[5] in used2 ])
                if unsure >= np.sum(mat.T[3]>0) * 0.6 :
                    genes.pop(gene)
                else :
                    cfl = conflicts.get(int(gene), {})
                    for id, m in enumerate(mat) :
                        if m[5] not in used2 :
                            used2.update( { int(k/10) for k in cfl.get(m[5],[]) } )
                    if np.any(mat.T[3] < 0) :
                        mat[np.where(mat.T[3]< 0)[0][::3], 3] *= -1
                        tmpSet[gene] = mat[mat.T[3] > 0]
        
        if params['orthology'] in ('ml', 'nj') :
            for gene, score in genes.items() :
                if gene not in new_groups :
                    mat = tmpSet.get(gene)
                    cfl = conflicts.get(int(gene), {})
                    _, bestPerGenome, matInGenome = np.unique(mat.T[1], return_index=True, return_inverse=True)
                    region_score = mat.T[4]/mat[bestPerGenome[matInGenome], 4]
                    if region_score.size >= bestPerGenome.size * 2 :
                        mat = mat[np.argsort(-mat.T[2])]
                        used2, kept = set([]), np.ones(mat.shape[0], dtype=bool)
                        for id, m in enumerate(mat) :
                            if m[5] in used2 :
                                kept[id] = False
                            else :
                                used2.update( { int(k/10) for k in cfl.get(m[5],[]) } )
                        mat = mat[kept]
                        mat = mat[np.argsort(-mat.T[3])]
                        _, bestPerGenome, matInGenome = np.unique(mat.T[1], return_index=True, return_inverse=True)
                        region_score = mat.T[4]/mat[bestPerGenome[matInGenome], 4]
                    if region_score.size > bestPerGenome.size * 3 and len(region_score) > 1000 :
                        region_score2 = sorted(region_score, reverse=True)
                        cut = region_score2[bestPerGenome.size*3-1]
                        if cut >= params['clust_identity'] :
                            cut = region_score2[bestPerGenome.size*6] if len(region_score) > bestPerGenome.size * 6 else params['clust_identity']
                        mat = mat[region_score>=cut]
                    inparalog = np.max(np.unique(mat.T[1], return_counts=True)[1])>1
                    if (not inparalog and np.min(mat[:, 3]) >= 10000 * params['clust_identity']) or len(mat) <= 1 :
                        new_groups[gene] = mat
                    else :
                        to_run.append([mat, inparalog, len(clust_ref[ mat[0][0]*1000 ]), params['map_bsn']+'.seq.npz', global_file])

            working_groups = pool2.imap_unordered(filt_per_group, sorted(to_run, key=lambda r:(r[1], r[0].shape[0]*r[0].shape[0]*r[2]), reverse=True))
            #working_groups = list(map(filt_per_group, sorted(to_run, key=lambda r:(r[1], r[0].shape[0]*r[0].shape[0]*r[2]), reverse=True)))
            for working_group in working_groups :
                gene = working_group[0][0][0]*1000
                new_groups[gene] = working_group[0]
                if len(working_group) > 1 :
                    cfl = conflicts.get(gene, {})
                    for id, matches in enumerate(working_group[1:]) :
                        ng = gene + (id+2)
                        new_groups[ng] = matches
                        conflicts[ng] = { mid:cfl.pop(mid, {}) for mid in matches.T[5] }
                        scores[ng] = np.sum(np.abs(matches[np.unique(matches.T[1], return_index=True)[1]].T[2]))
                        priorities[ng] = priorities[gene][:]
                        priorities[ng][0] = lowest_p
        else :
            for gene, score in genes.items() :
                if gene not in new_groups :
                    mat = tmpSet.get(gene)
                    cfl = conflicts.get(int(gene), {})
                    _, bestPerGenome, matInGenome = np.unique(mat.T[1], return_index=True, return_inverse=True)
                    region_score = np.min([mat.T[2]/mat[bestPerGenome[matInGenome], 2], mat.T[4]/mat[bestPerGenome[matInGenome], 4]], axis=0)
                    mat = mat[region_score>=params['clust_identity']]
                    used2, kept = set([]), np.ones(mat.shape[0], dtype=bool)
                    for id, m in enumerate(mat) :
                        if m[5] in used2 :
                            kept[id] = False
                        else :
                            used2.update( { int(k/10) for k in cfl.get(m[5],[]) + [m[5]*10] } )
                            
                    mat = mat[kept]
                    new_groups[gene] = mat
        
        if len(genes) :
            for gene in genes :
                matches = new_groups.get(gene)
                scores[gene] = np.sum(np.abs(matches[np.unique(matches.T[1], return_index=True)[1]].T[2]))

        while len(genes) :
            tmp = [ [scores[gene], gene] for gene in genes ]
            score, gene = max(tmp)
            if score < min_score :
                break
            mat = new_groups.pop(gene)
            # third, check its overlapping again
            paralog = False
            supergroup, used2 = {}, {}
            idens = 0.
            for m in mat :
                gid = m[5]
                conflict = used.get(gid, None) if gid in used else used2.get(gid, None)
                if conflict is not None :
                    if conflict < 0 :
                        superC = pangenome[-(conflict+1000)]
                        if superC not in supergroup :
                            supergroup[superC] = 0
                        if m[4] >= params['clust_identity'] * 10000 :
                            supergroup[superC] += 2
                        else :
                            supergroup[superC] += 1
                    elif conflict >0 :
                        paralog = True
                    m[3] = -1
                else :
                    if idens < m[4] : idens = m[4]
                    used2[gid] = 0
                    for gg in conflicts.get(gene, {}).get(gid, []) : 
                        g2, gs = int(gg/10), gg % 10
                        if gs == 1 :
                            if g2 not in used :
                                used2[g2] = -(gene+1000)
                        else :
                            used2[g2] = gene+1000 if gs == 2 else 0
                    
            if idens < (params['clust_identity']-0.02)*10000 :
                scores.pop(gene)
                genes.pop(gene)
                conflicts.pop(gene)
                continue
            mat = mat[mat.T[3] > 0]
            
            superR = [None, -999, 0]
            if len(supergroup) :
                for superC, cnt in sorted(supergroup.items(), key=lambda d:d[1], reverse=True) :
                    if cnt >= mat.shape[0] or cnt >= len(panList[superC]) :
                        gl1, gl2 = panList[superC], set(mat.T[1])
                        s = len( (gl1 | gl2) ^ gl1 ) - 2*len(gl1 & gl2)
                        if s < 0 : 
                            s = -1
                        if [s, cnt] > superR[1:] :
                            superR = [superC, s, cnt]
            if superR[1] > 0 or superR[2] > 0 :
                pangene = superR[0]
            elif paralog :
                new_groups[gene] = mat
                continue
            else :
                pangene = gene
            scores.pop(gene)
            genes.pop(gene)
            conflicts.pop(gene)
            pangenome[gene] = pangene
            used.update(used2)
            
            panList[pangene] = panList.get(pangene, set([])) | set(mat.T[1])

            pangene_name = (encodes[int(pangene/1000)] + '/' + str(pangene % 1000)) if pangene % 1000 > 0 else encodes[int(pangene/1000)]
            gene_name = (encodes[int(gene/1000)] + '/' + str(gene % 1000)) if gene % 1000 > 0 else encodes[int(gene/1000)]
            if len(pangenome) % 100 == 0 :
                logger('{4} / {5}: pan gene "{3}" : "{0}" picked from rank {1} and score {2}'.format(gene_name, min_rank, score/10000., pangene_name, len(pangenome), len(scores)+len(pangenome)))
            mat_out.append([pangene_name, gene_name, min_rank, mat])
    mat_out.append([0, 0, 0, []])
    return 

def load_priority(priority_list, genes, encodes) :
    file_priority = { encodes[fn]:id for id, fnames in enumerate(priority_list.split(',')) for fn in fnames.split(':') }
    unassign_id = max(file_priority.values()) + 1
    priorities = { n:[ file_priority.get(g[0], unassign_id), -len(g[6]), g[5] ] for n, g in genes.items() }
    priorities.update({ g[1]:[file_priority.get(g[1], unassign_id), 0, 0] for g in genes.values() })
    return priorities

def writeGenomes(fname, seqs) :
    with open(fname, 'w') as fout :
        for g, s in seqs.items() :
            fout.write('>{0} {1}\n{2}\n'.format(g, s[0], s[-1]))
    return fname

def iter_map_bsn(data) :
    prefix, clust, id, taxon, seq, orthoGroup, old_prediction, params = data
    stop = ['TAG', 'TAA', 'TGA'] if params['gtable'] != 4 else ['TAA', 'TAG']
    gfile, out_prefix = '{0}.{1}.genome'.format(prefix, id), '{0}.{1}'.format(prefix, id)
    with open(gfile, 'w') as fout :
        for n, s in seq :
            fout.write('>{0}\n{1}\n'.format(n, s) )

    if params['noDiamond'] :
        blastab, overlap = uberBlast('-r {0} -q {1} -f -m -O --blastn --min_id {2} --min_cov {3} --min_ratio {4} --merge_gap {5} --merge_diff {6} -t 1 -e 0,3 --gtable {7}'.format(\
            gfile, clust, params['match_identity']-0.1, params['match_frag_len'], params['match_frag_prop'], params['link_gap'], params['link_diff'], params['gtable'] ).split())
    else :
        blastab, overlap = uberBlast('-r {0} -q {1} -f -m -O --blastn --diamond --min_id {2} --min_cov {3} --min_ratio {4} --merge_gap {5} --merge_diff {6} -t 1 -s 1 -e 0,3 --gtable {7}'.format(\
            gfile, clust, params['match_identity']-0.1, params['match_frag_len'], params['match_frag_prop'], params['link_gap'], params['link_diff'], params['gtable'] ).split())
    os.unlink(gfile)
    blastab.T[:2] = blastab.T[:2].astype(int)
    
    # compare with old predictions
    blastab = compare_prediction(blastab, old_prediction)
    groups, groups2 = [], {}
    ids = np.zeros(np.max(blastab.T[15])+1, dtype=bool)
    for tab in blastab :
        if tab[16][1] >= params['match_identity'] and (tab[16][2] >= max(params['match_prop']*tab[12], params['match_len']) or \
                                                       tab[16][2] >= max(params['match_prop1']*tab[12], params['match_len1']) or \
                                                       tab[16][2] >= max(params['match_prop2']*tab[12], params['match_len2'])) :
            ids[tab[15]] = True
            if len(tab[16]) <= 4 :
                groups.append(tab[:2].tolist() + tab[16][:2] + [None, 0, [tab[:16]]])
            else :
                length = tab[7]-tab[6]+1
                if tab[2] >= params['match_identity'] and (length >= max(params['match_prop']*tab[12], params['match_len']) or \
                                                           length >= max(params['match_prop1']*tab[12], params['match_len1']) or \
                                                           length >= max(params['match_prop2']*tab[12], params['match_len2'])) :
                    groups.append(tab[:2].tolist() + [tab[11], tab[2], None, 0, [tab[:16]]])
                if tab[16][3] not in groups2 :
                    groups2[tab[16][3]] = tab[:2].tolist() + tab[16][:2] + [None, 0, [[]]*(len(tab[16])-3)]
                x = [i for i, t in enumerate(tab[16][3:]) if t == tab[15]][0]
                groups2[tab[16][3]][6][x] = tab[:16]
        else :
            tab[2] = -1
    groups.extend(list(groups2.values()))
    overlap = overlap[ids[overlap.T[0]] & ids[overlap.T[1]], :2]
    convA, convB = np.tile(-1, np.max(blastab.T[15])+1), np.tile(-1, np.max(blastab.T[15])+1)
    seq = dict(seq)
    for id, group in enumerate(groups) :
        group[4] = np.zeros(group[6][0][12], dtype=np.uint8)
        group[4].fill(0)
        group[5] = id
        group[6] = np.array(group[6])
        if group[6].shape[0] == 1 :
            convA[group[6].T[15].astype(int)] = id
        else :
            convB[group[6].T[15].astype(int)] = id
        max_sc = []
        for tab in group[6] :
            matchedSeq = seq[tab[1]][tab[8]-1:tab[9]] if tab[8] < tab[9] else rc(seq[tab[1]][tab[9]-1:tab[8]])
            ms, i, f, sc = [], 0, 0, [0, 0, 0]
            for s, t in re.findall(r'(\d+)([A-Z])', tab[14]) :
                s = int(s)
                if t == 'M' :
                    ms.append(matchedSeq[i:i+s])
                    i += s
                    sc[f] += s
                elif t == 'D' :
                    i += s
                    f = (f-s)%3
                else :
                    ms.append('-'*s)
                    f = (f+s)%3
            ms = ''.join(ms)
            sc = np.max(sc)
            sc2 = np.max(np.diff(np.concatenate([[0], np.where(in1d(re.findall('...', ms), stop))[0]*3, [len(ms)]])))
            sc = np.min([sc, sc2+3])
            x = baseConv[np.array(list(ms)).view(asc2int)]
            group[4][tab[6]-1:tab[6]+len(x)-1] = x
            r = np.sqrt(float(sc)/tab[12] * tab[10]) #|2./(1./(float(sc)/tab[12]) + 1./tab[10]) if float(sc)/tab[12] > tab[10] else float(sc)/tab[12]
            msc = (sc * tab[2])*np.sqrt(sc*r)
            amsc = float(msc)/(tab[7]-tab[6]+1)
            max_sc.append([tab[6], tab[7], amsc, msc])
        for i, c in enumerate(max_sc[1:]) :
            p = max_sc[i]
            if c[0] < p[1] :
                if c[2] > p[2] :
                    p[1] = c[0] - 1
                    p[3] = np.max(p[2] * (p[1]-p[0]+1), 0)
                else :
                    c[0] = p[1] + 1
                    c[3] = np.max(c[2] * (c[1]-c[0]+1), 0)
        group[2] = np.sum([c[3] for c in max_sc])
    overlap = np.vstack([np.vstack([m, n]).T[(m>=0) & (n >=0)] for m in (convA[overlap.T[0]], convB[overlap.T[0]]) \
                         for n in (convA[overlap.T[1]], convB[overlap.T[1]]) ] + [np.vstack([convA, convB]).T[(convA >= 0) & (convB >=0)]])
    bsn=np.array(groups, dtype=object)
    size = np.ceil(np.vectorize(lambda n:len(n))(bsn.T[4])/3).astype(int)
    bsn.T[4] = [ (b[:s]*25+b[s:2*s]*5 + np.concatenate([b, np.zeros(-b.shape[0]%3, dtype=int)])[2*s:]).astype(np.uint8) for b, s in zip(bsn.T[4], size) ]
    
    orthoGroup = np.load(orthoGroup, allow_pickle=True)
    orthoGroup = dict([[(g[0], g[1]), 1] for g in orthoGroup] + [[(g[1], g[0]), 1] for g in orthoGroup])
    if overlap.shape[0] :
        ovl_score = np.vectorize(lambda m,n:0 if m == n else orthoGroup.get((m,n), 2))(bsn[overlap.T[0], 0], bsn[overlap.T[1], 0])
        overlap = np.hstack([overlap, ovl_score[:, np.newaxis]])
    else :
        overlap = np.zeros([0, 3], dtype=np.int64)
    
    #if np.sum(blastab.T[0] == 14994) > 1 :
    #    print('a')
    np.savez_compressed(out_prefix+'.bsn.npz', bsn=bsn, ovl=overlap)
    return out_prefix

def compare_prediction(blastab, old_prediction) :
    blastab = pd.DataFrame(blastab)
    blastab = blastab.assign(s=np.min([blastab[8], blastab[9]], 0)).sort_values(by=[1, 's']).drop('s', axis=1).values
    
    blastab.T[10] = 0.1
    with MapBsn(old_prediction) as op :
        curr = [None, -1, 0]
        for bsn in blastab :
            if curr[1] != bsn[1] :
                curr = [op.get(bsn[1]), bsn[1], 0]
            if bsn[8] < bsn[9] :
                s, e, f = bsn[8], bsn[9], {(bsn[8] - bsn[6]+1)%3+1, (bsn[9] + (bsn[12] - bsn[7])+1)%3+1}
            else :
                s, e, f = bsn[9], bsn[8], {-(bsn[8] - bsn[6]+1)%3-1, -(bsn[9] + (bsn[12] - bsn[7])-1)%3-1}
            while curr[2] < len(curr[0]) and  s > curr[0][curr[2]][2] :
                    curr[2] += 1

            for p in curr[0][curr[2]:] :
                if e < p[1] : 
                    break
                elif p[3] == '+' :
                    if p[1]%3+1 not in f and (p[2]+1)%3+1 not in f :
                        continue
                else :
                    if -(p[1] - 1)%3-1 not in f and -(p[2])%3-1 not in f :
                        continue
                ovl = min(e, p[2]) - max(s, p[1]) + 1.
                
                if ovl >= 0.6*(p[2]-p[1]+1) or ovl >= 0.6*(e-s+1) :
                    ovl = ovl/(p[2] - p[1]+1)
                    if ovl > bsn[10] :
                        bsn[10] = ovl
            
    return pd.DataFrame(blastab).sort_values(by=[0, 1, 11]).values

baseConv = np.zeros(255, dtype=np.uint8)
baseConv[(np.array(['A', 'C', 'G', 'T']).view(asc2int),)] = (1, 2, 3, 4)

def get_map_bsn(prefix, clust, genomes, orthoGroup, old_prediction, conn, seq_conn, mat_conn, clf_conn, saveSeq) :
    if len(genomes) == 0 :
        sys.exit(1)

    taxa = {}
    for g, s in genomes.items() :
        if s[0] not in taxa : taxa[s[0]] = []
        taxa[s[0]].append([g, s[1]])
    
    ids = 0
    
    seqs, seq_cnts = [], 0
    mats, mat_cnts = [], 0
    
    blastab, overlaps = [], {}
    for bId, bsnPrefix in enumerate(pool.imap_unordered(iter_map_bsn, [(prefix, clust, id, taxon, seq, orthoGroup, old_prediction, params) for id, (taxon, seq) in enumerate(taxa.items())])) :
    #for bId, bsnPrefix in enumerate(map(iter_map_bsn, [(prefix, clust, id, taxon, seq, orthoGroup, old_prediction, params) for id, (taxon, seq) in enumerate(taxa.items())])) :
        tmp = np.load(bsnPrefix + '.bsn.npz', allow_pickle=True)
        bsn, ovl = tmp['bsn'], tmp['ovl']
        bsn.T[5] += ids
        ovl[:, :2] += ids
        
        prev_id = ids
        ids += bsn.shape[0]
        bsn.T[1] = genomes.get(bsn[0, 1], [-1])[0]
        
        if ovl.shape[0] :
            overlaps.update({id:[] for id in np.unique((ovl[:, :2]/30000).astype(int)) if id not in overlaps})
            ovl = np.vstack([ovl, ovl[:, (1,0,2)]])
            ovl = ovl[np.argsort(ovl.T[0])]
            ovl = np.hstack([(ovl[:, :1]/30000).astype(int), ovl[:, :1]%30000, ovl[:, 1:2]*10+ovl[:, 2:]])
            ovl = np.split(ovl, np.cumsum(np.unique(ovl.T[0], return_counts=True)[1])[:-1])
            for ovl2 in ovl :
                overlaps[ovl2[0, 0]].append(ovl2[:, 1:])
            for id in np.arange(int(prev_id/30000), int(ids/30000)) :
                if id in overlaps :
                    ovl = np.vstack(overlaps.pop(id))
                    ovl = np.concatenate([np.cumsum(np.concatenate([[0], np.bincount(ovl.T[0], minlength=30000)]))+30001, ovl.T[1]])
                    clf_conn.save(id, ovl)
        del ovl
        
        if saveSeq :
            seqs = np.concatenate([seqs, bsn.T[4]])
            ss = np.split(seqs, np.arange(1000, seqs.shape[0], 1000))
            seqs = ss[-1]
            for s in ss[:-1] :
                seq_conn.save(seq_cnts, s)
                seq_cnts += 1
        bsn.T[4] = bsn.T[3]

        mats = np.concatenate([mats, bsn.T[6]])
        mm = np.split(mats, np.arange(1000, mats.shape[0], 1000))
        mats = mm[-1]
        for m in mm[:-1] :
            mat_conn.save(mat_cnts, m)
            mat_cnts += 1
        bsn.T[6] = np.array([ len(b) for b in bsn.T[6] ], dtype=np.uint8)
        bsn.T[2:5] = bsn.T[2:5] * 10000
        
        bsn = bsn[np.argsort(-bsn.T[2])].astype(int)
        blastab.append(bsn)
        del bsn

        os.unlink(bsnPrefix + '.bsn.npz')
        logger('Merged {0}'.format(bsnPrefix))
        if bId % 500 == 499 or bId == len(taxa) - 1 :
            blastab = np.vstack(blastab)
            blastab = blastab[np.argsort(blastab.T[0], kind='mergesort')]
            blastab = np.split(blastab, np.cumsum(np.unique(blastab.T[0], return_counts=True)[1])[:-1])
            conn.update(blastab)
            del blastab
            blastab = []
            
    pool.close()
    pool.join()
    if saveSeq and seqs.shape[0] :
        seq_conn.save(seq_cnts, seqs)
    if mats.shape[0] :
        mat_conn.save(mat_cnts, mats)
    for id in overlaps.keys() :
        ovl = np.vstack(overlaps.get(id))
        ovl = np.concatenate([np.cumsum(np.concatenate([[0], np.bincount(ovl.T[0], minlength=30000)]))+30001, ovl.T[1]])
        clf_conn.save(id, ovl)
    del ovl, overlaps

def checkPseu(n, s, gtable) :
    if len(s) < params['min_cds'] :
        #logger('{0} len={1} is too short'.format(n, len(s)))
        return 1

    if len(s) % 3 > 0 and 'f' not in params['incompleteCDS'] :
        #logger('{0} is discarded due to frameshifts'.format(n))
        return 2
    aa = transeq({'n':s.upper()}, frame=1, transl_table=gtable, markStarts=True)['n'][0]
    if aa[0] != 'M' and 's' not in params['incompleteCDS'] :
        #logger('{0} is discarded due to lack of start codon'.format(n))
        return 3
    if aa[-1] != 'X' and 'e' not in params['incompleteCDS'] :
        #logger('{0} is discarded due to lack of stop codon'.format(n))
        return 4
    if len(aa[:-1].split('X')) > 1 and 'i' not in params['incompleteCDS'] :
        #logger('{0} is discarded due to internal stop codons'.format(n))
        return 5
    return 0
    

def addGenes(genes, gene_file, gtable) :
    for gfile in gene_file.split(',') :
        if gfile == '' : continue
        gprefix = os.path.basename(gfile).split('.')[0]
        ng = readFasta(gfile)
        for name, s in ng.items() :
            if not checkPseu(name, s, gtable) :
                genes['{0}:{1}'.format(gprefix,name)] = [ gfile, '', 0, 0, '+', int(hashlib.sha1(s.encode('utf-8')).hexdigest(), 16), s]
    return genes

def writeGenes(fname, genes, priority) :
    uniques = {}
    groups = []
    with open(fname, 'w') as fout :
        for n, _ in sorted(priority.items(), key=itemgetter(1)) :
            if n in genes :
                s = genes[n][6]
                len_s, hcode = len(s), genes[n][5]
                if len_s :
                    if len_s not in uniques :
                        uniques = { len_s:{ hcode:n } }
                    elif hcode in uniques[ len_s ] :
                        groups.append([ uniques[len_s][hcode], n, 10000 ])
                        continue
                    uniques[ len_s ][ hcode ] = n
                    fout.write( '>{0}\n{1}\n'.format(n, s) )
    return fname, groups

def determineGroup(gIden, global_differences, min_iden, nSigma) :
    ingroup = np.zeros(gIden.shape[0], dtype=bool)
    ingroup[gIden.T[1] >= (min_iden-0.02)*10000] = True

    for i1, m1 in enumerate(gIden) :
        if m1[1] >= (min_iden-0.02)*10000 :
            m2 = gIden[i1+1:][ingroup[gIden[i1+1:, 2]] != True]
            if m2.size :
                gs = np.vectorize(lambda g1, g2: (params['self_id'], 0.) if g1 == g2 else global_differences.get(tuple(sorted([g1, g2])), (0.5, 0.6) ))(m2.T[0], m1[0])
                sc = (1.-m2.T[1].astype(float)/m1[1])/(gs[0]*np.exp(nSigma*gs[1]))
                ingroup[m2[sc < 1, 2].astype(int)] = True
            else :
                break
    _, tag, idx = np.unique(gIden.T[0], return_inverse=True, return_index=True)
    ingroup[:] = ingroup[tag[idx]]
    return ingroup

def initializing2(data) :
    bsn_file, genes, global_file = data

    global_differences = dict(np.load(global_file, allow_pickle=True))
    outputs = []
    with MapBsn(bsn_file+'.tab.npz') as conn :
        for gene in genes :
            matches = conn.get(gene)
            if len(matches) <= 1 :
                outputs.append( [gene, matches, matches[0, 2]] )
                continue
            matches = matches[np.argsort(-(1000*np.abs(matches.T[2])/np.max(np.abs(matches.T[2])) + matches.T[3]))]
            matches.T[4] = (10000 * matches.T[3]/matches[0, 3]).astype(int)
            gIden = np.hstack([matches[:, [1, 4]], np.arange(matches.shape[0])[:, np.newaxis]])
            ingroup = determineGroup(gIden, global_differences, params['clust_identity'], params['allowed_sigma'])
            matches = matches[ingroup]
            s = np.sum(np.abs(matches[np.unique(matches.T[1], return_index=True)[1]].T[2]))
            outputs.append([ gene, matches, s ])
    return outputs

def initializing(bsn_file, global_file) :
    gene_scores = {}
    with MapBsn(bsn_file + '.tab.npz') as conn, MapBsn(bsn_file + '.tmp.npz', 'w') as conn2 :
        genes = np.array(sorted(conn.keys()))
        for ite in xrange(0, len(genes), 10000) :
            logger('Initializing: {0}/{1}'.format(ite, len(genes)))
            genes2 = genes[ite:ite+10000]
            
            toUpdates = pool2.imap_unordered(initializing2, [(bsn_file, gs, global_file) for gs in np.split(genes2, np.arange(50, genes2.size, 50)) ])
            #toUpdates = map(initializing2, [(bsn_file, gs, global_file) for gs in np.split(genes2, np.arange(50, genes2.size, 50)) ])
            for toUpdate in toUpdates :
                for gene, data, score in toUpdate :
                    gene_scores[int(gene)] = score
                    conn2.save(gene, data)
    shutil.move(bsn_file + '.tmp.npz', bsn_file + '.tab.npz')
    return gene_scores


def ite_synteny_resolver(data) :
    grp_tag, ids, co_genomes, neighbors, nNeighbor = data
    #ids = np.where(orthologs.T[0] == grp_tag)[0]

    distances, conflicts, inConflicts = [], {}, {}
    for m, i in enumerate(ids) :
        for n in np.arange(m+1, ids.size) :
            j = ids[n]
            #ni, nj = np.array(list(neighbors[i]), dtype=int), np.array(list(neighbors[j]), dtype=int)
            #oi, oj = set(orthologs[ni, 0]), set(orthologs[nj, 0])
            oi, oj = neighbors[m], neighbors[n]
            ois, ojs = np.min([6, len(oi)]), np.min([6, len(oj)])
            s = 3 * len(oi & oj) + np.max([6-ois, 6-ojs, 0]) + 1
            d = 3 * nNeighbor - s
            distances.append([d, co_genomes[m] != co_genomes[n], i, j])
            if co_genomes[m] == co_genomes[n] and d > 0 :
                conflicts[(i, j)] = conflicts[(j, i)] = 1
                inConflicts[i] = inConflicts[j] = 1

    if conflicts :
        distances.sort()
        groups, tags = { id:[[id], []] if id in inConflicts else [[], [id]] for id in ids }, { id:id for id in ids }
        for idx, (d, _, i, j) in enumerate(distances) :
            if tags[i] == tags[j] : continue
            if (i, j) in conflicts :
                distances = distances[idx:]
                break
            ti, tj = tags[i], tags[j]
            skip = False
            for m in groups[ti][0] :
                for n in groups[tj][0] :
                    if (m, n) in conflicts :
                        skip = True
                        break
                if skip :
                    break

            if not skip :
                gg = groups.pop(tj)
                for g in gg[0] :
                    tags[g] = tags[i]
                for g in gg[1] :
                    tags[g] = tags[i]
                groups[ti][0].extend(gg[0])
                groups[ti][1].extend(gg[1])
        diffs = {}
        for d, _, i, j in distances :
            if (i, j) in conflicts :
                diffs[tags[j]] = diffs[tags[i]] = 1

        if len(diffs) >= len(groups) :
            return [grp_tag, { id:grp[0]+grp[1] for id, grp in groups.items() }]
        else :
            return [grp_tag, None]
    return [None, None]

def synteny_resolver(prefix, prediction, nNeighbor = 2) :
    prediction = pd.read_csv(prediction, sep='\t', header=None)
    prediction = prediction.assign(s=np.min([prediction[9], prediction[10]], 0)).sort_values(by=[5, 's']).drop('s', axis=1).values
    neighbors = [ set([]) for i in np.arange(np.max(prediction.T[2])+1) ]
    orthologs = np.vstack([['', ''], np.copy(prediction[:, [0,3]])])
    orthologs[prediction.T[2].astype(int)] = prediction[:, [0,3]]
    orthologs = orthologs[:len(neighbors)]
    orthologs2 = np.unique(orthologs, return_inverse=True)[1].reshape([-1, 2])
    
    orth_cnt = dict(zip(*(np.unique(orthologs2.T[0], return_counts=True))))
    
    for pId, predict in enumerate(prediction) :
        nId = np.concatenate([np.arange(pId-1, max(pId-4, -1), -1), np.arange(pId+1, min(pId+4, prediction.shape[0]))])
        nbs = {orthologs2[neighbor[2], 0] for neighbor in prediction[nId] if neighbor[5] == predict[5]} - set([orthologs2[predict[2], 0]])
        neighbors[predict[2]].update(nbs)
    paralog_groups = np.unique(orthologs2, axis=0, return_counts=True)
    paralog_groups = np.unique(paralog_groups[0][paralog_groups[1]>1, 0])

    #outs = list(map(ite_synteny_resolver, [ [grp_tag, orthologs2, neighbors, nNeighbor] for grp_tag in paralog_groups ]))
    toRun = [ [grp_tag, np.where(orthologs2.T[0] == grp_tag)[0], [], [], nNeighbor] for grp_tag in sorted(paralog_groups, key=lambda p:orth_cnt.get(p, 0)) ]
    for t in toRun :
        genomes, co_genomes = np.unique(orthologs2[t[1], 1], return_inverse=True)
        if len(genomes) > 1 :
            t[2] = co_genomes
            t[3] = [ neighbors[i] for i in t[1] ]
    outs = pool2.imap_unordered(ite_synteny_resolver, [t for t in toRun if len(t[2])>0])
    for grp_tag, groups in outs :
        if groups is not None :
            for id, i in enumerate(sorted(groups.values(), key=lambda v:[-len(v), v])) :
                if id > 0 :
                    if len(re.findall(b'/\d+$', orthologs[i, 0])) > 0 :
                        orthologs[i, 0] = orthologs[i[0], 0] + '.{0}'.format(id)
                    else :
                        orthologs[i, 0] = orthologs[i[0], 0] + '/0.{0}'.format(id)

    prediction.T[0] = orthologs[prediction.T[2].astype(int), 0]
    prediction = pd.DataFrame(prediction).sort_values(by=[0, 2, 7])
    prediction.to_csv(prefix+'.synteny.Prediction', sep='\t', index=False, header=False)
    return prefix+'.synteny.Prediction'

def determineGeneStructure(data) :
    pid, pred, seq, s, e, s2, e2, lp, allowed_vary, gtable = data
    cds, cdss = 'CDS', []
    for frame, aa_seq in zip(pred[14], transeq({'n':seq}, transl_table=gtable, markStarts=True, frame=','.join([str(f+1) for f in pred[14]]))['n']) :
        if (len(seq) - frame) % 3 > 0 :
            aa_seq = aa_seq[:-1]
        cds = 'CDS'
        
        s0, s1 = aa_seq.find('M', int(lp/3), int((lp+allowed_vary)/3)), aa_seq.rfind('M', 0, int(lp/3))
        start = s0 if s0 >= 0 else s1
        if start < 0 :
            cds, start = 'nostart', int(lp/3)
        stop = aa_seq.find('X', start)
        while 0 <= stop < int((lp+allowed_vary)/3) :
            s0 = aa_seq.find('M', stop, int((lp+allowed_vary)/3))
            if s0 >= 0 :
                start = s0
                stop = aa_seq.find('X', start)
            else :
                break
        if stop < 0 :
            cds = 'nostop'
        elif (stop - start + 1)*3 < pred[12] - allowed_vary :
            cds = 'premature_stop:{0:.2f}%'.format((stop - start + 1)*300/pred[12])
            
        if cds == 'CDS' :
            if pred[11] == '+' :
                start, stop = s2 + start*3 + frame, s2 + stop*3 + 2 + frame
            else :
                start, stop = e2 - stop*3 - 2 - frame, e2 - start*3 - frame
            break
        else :
            start, stop = s, e
            cdss.append(cds)
            if frame > 0 :
                cds = cdss[0].replace('premature_stop', 'frameshift') if cdss[0].find('premature_stop') >= 0 else 'frameshift'
    return pid, cds, start, stop

def write_output(prefix, prediction, genomes, clust_ref, encodes, old_prediction, pseudogene, untrusted, gtable, clust=None, orthoPair=None) :
    def addOld(opd, gsize) :
        # old genes in the predictions are not all included in the predicted pan-genome due to varies reasons. They are added in here
        if opd[4] == 0 :
            return [opd[0], -1, -1, op[0], opd[0], op[0], 1., 1, opd[2]-opd[1]+1, opd[1], opd[2], opd[3], opd[2]-opd[1]+1, gsize, [0], 'CDS', '{0}:{1}-{2}'.format(opd[0].split(':', 1)[1], opd[1], opd[2])]
        else :
            if opd[4] < 7 :
                reason = ['', 'Too_short', 'Pseudogene:Frameshift', 'Pseudogene:No_start', 'Pseudogene:No_stop', 'Pseudogene:Premature', 'Error_in_sequence'][opd[4]]
            else :
                reason = 'Overlap_with:{0}_g_{1}'.format(prefix, int(opd[4]/10))
            return [opd[0], -1, -1, op[0], opd[0], op[0], 1., 1, opd[2]-opd[1]+1, opd[1], opd[2], opd[3], opd[2]-opd[1]+1, gsize, -1, 'misc_feature', reason]
    def setInFrame(part) :
        # re-locate start and end points of the alignment, making sure that it is in the coding frame of the representative gene
        if part[9] < part[10] :
            l, r, d = min(part[7]-1, part[9]-1), min(part[12]-part[8], part[13]-part[10]), 1
        else :
            l, r, d = min(part[7]-1, part[13]-part[9]), min(part[12]-part[8], part[10]-1), -1
        if l <= 9 and part[7] - l == 1 :
            part[7], part[9] = part[7]-l, part[9]-l*d
        else :
            ll = (part[7]-1) % 3
            if ll > 0 :
                part[7], part[9] = part[7]+3-ll, part[9]+(3-ll)*d
        if r <= 9 and part[8] + r == part[12] :
            part[8], part[10] = part[8]+r, part[10]+r*d
        else :
            rr = (part[12] - part[8]) % 3
            if rr > 0 :
                part[8], part[10] = part[8]-3+rr, part[10]-(3-rr)*d

        part[9:12] = (part[9], part[10], '+') if d > 0 else (part[10], part[9], '-')

    prediction = pd.read_csv(prediction, sep='\t', header=None)
    prediction = prediction.assign(cds=np.repeat('CDS', prediction.shape[0]), old_tag=np.repeat('New_prediction', prediction.shape[0]), s=np.min([prediction[9], prediction[10]], 0)).sort_values(by=[5, 's']).drop('s', axis=1).values

    for part in prediction :
        setInFrame(part)

    for id, part in enumerate(prediction[1:]) :
        # if two sequential alignments are from the same representative gene, try to merge them together
        prev = prediction[id]
        if part[2] == prev[2] and part[11] == prev[11] and prev[5] == part[5] and part[9] - prev[10] < 500 :
            if part[11] == '+' and part[7] - prev[8] < 500 :
                diff = (part[7]-prev[8]) - (part[9]-prev[10])
                diff = '' if diff == 0 else ('{0}I'.format(diff) if diff > 0 else '{0}D'.format(-diff))
                part[7], part[9] = prev[7], prev[9]
                part[14] = prev[14] + diff + part[14]
                prev[0] = ''
            elif part[11] == '-' and prev[7] - part[8] < 500 :
                diff = (prev[7]-part[8]) - (part[9]-prev[10])
                diff = '' if diff == 0 else ('{0}I'.format(diff) if diff > 0 else '{0}D'.format(-diff))                    
                part[8], part[9] = prev[8], prev[9]
                part[14] = part[14] + diff + prev[14]
                prev[0] = ''
    
    prediction = prediction[prediction.T[0] != '']  # remove redundant alignment
    _, gTag, gIdx, gCnt = np.unique(prediction.T[2], return_counts=True, return_inverse=True, return_index=True)
    prediction.T[1] = gCnt[gIdx]
    prediction.T[2] = gTag[gIdx]+1

    # compare with old annotation and add in old genes if they are not included in the new annotation
    op, old_to_add = ['', 0, []], []
    for pred in prediction :
        pred[14] = sorted(np.unique(np.cumsum([0]+[int(n) if t == 'D' else -int(n) for n, t in re.findall(r'(\d+)([ID])', pred[14])])%3))
        if pred[5] != op[0] :
            if len(op[2]) :
                for k in xrange(op[1], len(op[2])) :
                    opd = op[2][k]
                    if opd[4] != 7 :
                        old_to_add.append(addOld(opd, len(genomes[encodes[op[0]]][1])))
                    
            op = [pred[5], 0, old_prediction.get(pred[5], [])]
        old_tag = []
        s, e = pred[9], pred[10]
        for k in xrange(op[1], len(op[2])) :
            opd = op[2][k]
            if opd[2] < s :
                if opd[4] != 7 :
                    old_to_add.append(addOld(opd, len(genomes[encodes[op[0]]][1])))
                op[1] = k + 1
                continue
            elif opd[1] > e :
                break
            ovl = min(opd[2], e) - max(opd[1], s) + 1
            if ovl >= 300 or ovl >= 0.6 * (opd[2]-opd[1]+1) or ovl >= 0.6 * (e - s + 1) :
                if opd[4] < 7 :
                    opd[4] = 10 * pred[2]
                if opd[3] != pred[11]:
                    continue
                if pred[11] == '+' :
                    f2 = np.unique([(opd[1] - pred[9])%3, (opd[2]+1 - pred[9])%3])
                else :
                    f2 = np.unique([(pred[10] - opd[1]+1)%3, (pred[10] - opd[2])%3])
                if np.any(in1d(f2, pred[14])) :
                    old_tag.append('{0}:{1}-{2}'.format(opd[0].split(':', 1)[1], opd[1], opd[2]))
                    opd[4] = 7

        pred[16] = ','.join(old_tag)
    if len(op[2]) :
        for k in xrange(op[1], len(op[2])) :
            opd = op[2][k]
            if opd[4] != 7 :
                old_to_add.append(addOld(opd, len(genomes[encodes[op[0]]][1])))
    maxTag = np.max(gTag)+1
    for g in old_to_add :
        maxTag += 1
        g[2] = maxTag
        
    # build orthologous groups for missing genes
    old_genes = [ encodes[g[0]] for g in old_to_add if g[15] == 'CDS' ]
    if len(old_genes) :
        queries = set(old_genes)
        groups = { g:[g] for g in old_genes }
        tags = None
        if clust and os.path.exists(clust) :
            clust = np.load(clust.rsplit('.', 1)[0] + '.npy', allow_pickle=True)
            while len(queries) :
                c = clust[in1d(clust.T[1], list(queries))]
                queries = set(c.T[0]) - queries
                for grp, gene, _ in c :
                    if grp not in groups :
                        groups[grp] = groups.pop(gene)
                    else :
                        groups[grp].extend(groups.pop(gene, {}))
            tags = {ggg: g for g, gg in groups.items() for ggg in gg}
        if orthoPair and os.path.exists(orthoPair) :
            clust = np.load(orthoPair, allow_pickle=True)
            clust = clust[np.all(in1d(clust, queries).reshape(clust.shape), 1)]
            for g1, g2, _ in clust :
                t1, t2 = tags[g1], tags[g2]
                for g in groups[t2] :
                    tags[g] = t1
                groups[t1].extend(groups.pop(t2))
            groups = {g:sorted(gg) for g, gg in groups.items()}
            tags = {ggg:gg[0] for g, gg in groups.items() for ggg in gg}
        if tags :
            decodes = {v:k for k, v in encodes.items()}
            for g in old_to_add :
                if g[15] == 'CDS' :
                    g[0] = decodes[tags[encodes[g[0]]]]

    # reorder after adding old genes
    try :
        prediction = pd.DataFrame(np.vstack([prediction, np.array([g for g in old_to_add], dtype=object)])).sort_values(by=[5,9]).values
    except :
        prediction = pd.DataFrame(prediction).sort_values(by=[5,9]).values
    # if the secondary repr gene is the same as the primary gene, remove it
    prediction[np.array([p.rsplit('/', 2)[0] for p in prediction.T[4]]) == np.array([p.rsplit('/', 2)[0] for p in prediction.T[0]]), 4] = ''

    # add representative genes as allele 1
    alleles = {}
    for part in prediction :
        if part[0] not in alleles and part[15] != 'misc_feature' :
            alleles[part[0]] = {}
            if part[0] in encodes :
                gId = encodes[part[0]]
                if gId in clust_ref :
                    alleles[part[0]] = {clust_ref[gId]:str(1)}


    for ppid in np.arange(0, len(prediction), 10000) :
        toRun = []
        for xid, pred in enumerate(prediction[ppid:ppid+10000]) :
            pid = ppid + xid
            if pred[15] == 'misc_feature' or pred[0] == '' : #
                pred[13] = '{0}:{1}:{2}-{3}:{4}-{5}'.format(pred[0], 't0', pred[7], pred[8], pred[9], pred[10])
                continue
            allowed_vary = int(pred[12]*(1-pseudogene)+0.01)
            if pred[4] :
                map_tag = '{0}:({1}){2}:{3}-{4}:{5}-{6}'.format(pred[0], pred[4].rsplit(':', 1)[-1], '{0}', pred[7], pred[8], pred[9], pred[10])
            else :
                map_tag = '{0}:{1}:{2}-{3}:{4}-{5}'.format(pred[0], '{0}', pred[7], pred[8], pred[9], pred[10])
                
            pred2 = None
            if pred[1] > 1 or (pred[10]-pred[9]+1) < pred[12] - allowed_vary :
                cds = 'fragment:{0:.2f}%'.format((pred[10]-pred[9]+1)*100/pred[12])
                s, e = pred[9:11]
                seq2 = genomes[encodes[pred[5]]][1][(s-1):e] if pred[11] == '+' else rc(genomes[encodes[pred[5]]][1][(s-1):e])
                if seq2 not in alleles[pred[0]] :
                    alleles[pred[0]][seq2] = 't{0}'.format(len(alleles[pred[0]])+1)
                pred[13] = map_tag.format(alleles[pred[0]][seq2])
            else :
                s, e = pred[9:11]
                if pred[11] == '+' :
                    for i, pp in enumerate(prediction[pid+1:pid+5]) :
                        if pp[5] != pred[5] : break
                        elif pp[15] != 'misc_feature' :
                            pred2 = pp
                            #break
                    if pred2 is not None:
                        e2 = e + min(3*int((pred[13] - e)/3), 3*int((pred2[10] + 300 - e)/3))
                    else :
                        e2 = e + min(3*int((pred[13] - e)/3), 600+pred[12] - pred[8])
                    s2 = s - min(3*int((s - 1)/3), 60+pred[7]-1)
                    seq = genomes[encodes[pred[5]]][1][(s2-1):e2]
                    lp, rp = s - s2, e2 - e
                else :
                    for i, pp in enumerate(reversed(prediction[max(pid-5, 0):pid])) :
                        if pp[5] != pred[5] : break
                        elif pp[15] != 'misc_feature' :
                            pred2 = pp
                            #break
                    if pred2 is not None :
                        s2 = s - min(3*int((s - 1)/3), 3*int((s - pred2[9] + 300)/3))
                    else :
                        s2 = s - min(3*int((s - 1)/3), 600+pred[12]-pred[8])
    
                    e2 = e + min(3*int((pred[13] - e)/3), 60+pred[7]-1)
                    seq = rc(genomes[encodes[pred[5]]][1][(s2-1):e2])
                    rp, lp = s - s2, e2 - e

                seq2 = seq[(lp):(len(seq)-rp)]
                if seq2 not in alleles[pred[0]] :
                    if pred[4] == '' and pred[7] == 1 and pred[8] == pred[12] :
                        alleles[pred[0]][seq2] = str(len(alleles[pred[0]])+1)
                    else :
                        alleles[pred[0]][seq2] = 't{0}'.format(len(alleles[pred[0]])+1)
                pred[13] = map_tag.format(alleles[pred[0]][seq2])
                
                p = pred[0].rsplit('/', 2)[0]
                if encodes[p] in clust_ref and len(seq2) < pseudogene*len(clust_ref[encodes[p]]) :
                    cds = 'structural_variation:{0:.2f}%'.format(100.*len(seq2)/len(clust_ref[encodes[p]]))
                else :
                    cds = 'CDS'
                    toRun.append([pid, pred, seq, s, e, s2, e2, lp, allowed_vary, gtable])
            if cds != 'CDS' :
                pred[15] = 'pseudogene=' + cds
        for pid, cds, start, stop in pool2.imap_unordered(determineGeneStructure, toRun) :
            pred = prediction[pid]
            pred[9:11] = start, stop
            if cds != 'CDS' :
                pred[15] = 'pseudogene=' + cds
    prediction = pd.DataFrame(prediction).sort_values(by=[5,9]).values
    toMerge, unreliable = {}, {}
    for pid, pred in enumerate(prediction) :
        if pred[15] == 'misc_feature' or pid in toMerge :
            continue
        for pidx, pred2 in enumerate(prediction[pid+1:]) :
            pid2 = pid+pidx+1
            if pred2[15] == 'misc_feature' or pid2 in toMerge :
                continue
            if pred[5] == pred2[5] :
                ovl = (pred[10] - pred2[9]+1)
                if ovl >= 0.8*(pred[10]-pred[9]+1) or ovl >= 0.8*(pred2[10]-pred2[9]+1) :
                    if pred[11] == pred2[11] and (pred[9]%3 == pred2[9]%3 or pred[10]%3 == pred2[10]%3) :
                        pids = [pid, pid2] if pred[10]-pred[9] >= pred2[10] - pred2[9] else [pid2, pid]
                        toMerge[pids[1]] = pids[0]
                        if pred[8] - pred[7] >= pred2[8] - pred2[7] :
                            if pid2 not in unreliable: unreliable[pid2] = 1
                        else :
                            if pid not in unreliable: unreliable[pid] = 1
                        if pids[0] > pids[1] :
                            break
                    else :
                        if pred[8] - pred[7] >= pred2[8] - pred2[7] :
                            unreliable[pid2] = 2
                        else :
                            unreliable[pid] = 2
                elif pred2[9] > pred[10] :
                    break
            else :
                break
    cdss = {}
    for pid, pred in enumerate(prediction) :
        if pred[15] == 'misc_feature' :
            continue
        if pred[0] not in cdss :
            cdss[pred[0]] = [0, 0, 0, 0]
        cdss[pred[0]][2] += 1
        if pred[16] != '' and pid not in unreliable :
            cdss[pred[0]][3] += 1.
        if pred[0] != '' and pred[15] == 'CDS' :
            cdss[pred[0]][0] += 1.
            if pred[16] != '' and pid not in unreliable:
                cdss[pred[0]][1] += 1.

    removed = {'':1}
    for gene, stat in cdss.items() :
        glen = np.mean([len(s) for s in alleles[gene]]) if len(alleles[gene]) else 0
        if glen < params['min_cds'] :
            removed[gene] = 1
        elif (stat[1]+0.5)*(stat[3]+0.5) < (stat[0]+1)*untrusted[1]*(stat[2]+1)*untrusted[1] and \
                glen <= untrusted[0] :
            removed[gene] = 1
        elif stat[3] == 0 and len(re.findall(r'/[\.\d]+$', gene)) :
            removed[gene] = 1
    for pid2, pid1 in sorted(toMerge.items()) :
        p1, p2 = prediction[pid1], prediction[pid2]
        if p1[0] not in removed and p2[0] not in removed :
            p1[9] = min(p1[9], p2[9])
            p1[10] = max(p1[10], p2[10])
            p1[13] = ','.join(sorted([p1[13], p2[13]]))
            p1[4] = ','.join(sorted([x for x in [p1[4], p2[4]] if x]))
            p1[16] = ','.join(sorted(set(p1[16].split(',') + p2[16].split(','))-{''}))
        elif p1[0] in removed :
            p1[:] = p2[:]
        p2[0] = ''

    with open('{0}.allele.fna'.format(prefix), 'w') as allele_file :
        for gene, seqs in alleles.items() :
            if gene not in removed :
                for seq, id in sorted(seqs.items(), key=lambda s:s[1]) :
                    allele_file.write('>{0}_{1}\n{2}\n'.format(gene, id, seq))

    #prediction = prediction[prediction.T[0] != '']
    with open('{0}.PEPPAN.gff'.format(prefix), 'w') as fout :
        fout.write('#!gff-version 3\n#!annotation-source PEPPAN from enterobase.warwick.ac.uk\n')
        for pid, pred in enumerate(prediction) :
            if pred[0] == '' : continue
            if pred[15] == 'misc_feature' :
                fout.write('{0}\t{1}\tPEPPAN\t{2}\t{3}\t.\t{4}\t.\tID={5};{7}inference={6}\n'.format(
                    pred[5], 'misc_feature', pred[9], pred[10], pred[11],
                    '{0}_g_{1}'.format(prefix.rsplit('/', 1)[-1], pred[2]), pred[16],
                    'old_locus_tag={0}:{1}-{2};'.format(pred[0].split(':', 1)[1], pred[9], pred[10]),
                ))
            else :
                if pred[0] in removed :
                    fout.write('{0}\t{1}\tPEPPAN\t{2}\t{3}\t.\t{4}\t.\tID={5};{8}inference=ortholog_group:{6}{7}\n'.format(
                        pred[5], 'misc_feature',
                        pred[9], pred[10], pred[11],
                        '{0}_g_{1}'.format(prefix.rsplit('/', 1)[-1], pred[2]), pred[13],
                        ';{0}'.format(pred[15]) if pred[15].startswith('pseudogen') else '',
                        'note=Removed_untrusted_prediction;' if pred[16] == '' else 'note=Removed_untrusted_prediction;old_locus_tag={0};'.format(pred[16]),
                    ))

                elif unreliable.get(pid, 1) == 2 :
                    fout.write('{0}\t{1}\tPEPPAN\t{2}\t{3}\t.\t{4}\t.\tID={5};{8}inference=ortholog_group:{6}{7}\n'.format(
                        pred[5], 'pseudogene',
                        pred[9], pred[10], pred[11],
                        '{0}_g_{1}'.format(prefix.rsplit('/', 1)[-1], pred[2]), pred[13],
                        ';{0}'.format(pred[15]) if pred[15].startswith('pseudogen') else '',
                        'note=Overlapped_prediction;' if pred[16] == '' else 'note=Overlapped_prediction;old_locus_tag={0};'.format(pred[16]),
                    ))

                else :
                    fout.write('{0}\t{1}\tPEPPAN\t{2}\t{3}\t.\t{4}\t.\tID={5};{8}inference=ortholog_group:{6}{7}\n'.format(
                        pred[5], 'pseudogene' if pred[15].startswith('pseudogen') else pred[15],
                        pred[9], pred[10], pred[11],
                        '{0}_g_{1}'.format(prefix.rsplit('/', 1)[-1], pred[2]), pred[13],
                        ';{0}'.format(pred[15]) if pred[15].startswith('pseudogen') else '',
                        '' if pred[16] == '' else 'old_locus_tag={0};'.format(pred[16]),
                    ))

    allele_file.close()
    logger('Pan genome annotations have been saved in {0}'.format('{0}.PEPPAN.gff'.format(prefix)))
    logger('Gene allelic sequences have been saved in {0}'.format('{0}.allele.fna'.format(prefix)))
    return


def get_gene_group(cluFile, bsnFile) :
    clu = np.load(cluFile.rsplit('.',1)[0] + '.npy', allow_pickle=True)
    bsn = np.load(bsnFile, allow_pickle=True)
    bsn = bsn[bsn.T[2] > 0]
    # score genes
    geneGroups = {}
    pairTag =  {}

    for matrix in (clu, bsn) :
        for r, q, i in matrix :
            q2 = pairTag.get(q, q)
            r2 = pairTag.get(r, r)
            if q2 != r2 :
                rr = geneGroups.get(r2, [r2])
                qq = geneGroups.pop(q2, [q2])
                for q in qq :
                    pairTag[q] = r2
                geneGroups[r2] = rr + qq
    # get one-to-one hits
    return geneGroups

def get_global_difference(geneGroups, cluFile, bsnFile, geneInGenomes, nGene = 1000) :
    groupPresences = {}
    for grp, genes in geneGroups.items() :
        gg = np.array([geneInGenomes[g] for g in genes])
        presence = np.unique(gg[gg>=0], return_counts=True)
        groupPresences[grp] = [np.sum(presence[1] == 1), gg.size]
    groupScore = sorted([ [-p[0], p[1]-p[0], int(hashlib.sha1(bytes(g)).hexdigest(), 16), g] for g, p in groupPresences.items() if p[0] > 1 ])
    selectedGroups = [g[3] for g in groupScore[:nGene]]
    selectedGenes = []
    for g in selectedGroups :
        selectedGenes.extend(geneGroups[g])
    selectedGenes = np.array(selectedGenes)
    
    clu = np.load(cluFile.rsplit('.',1)[0] + '.npy', allow_pickle=True)
    bsn = np.load(bsnFile, allow_pickle=True)
    bsn = bsn[bsn.T[2] > 0]
    
    selectedClu = clu[in1d(clu.T[0], selectedGenes)]
    selectedBsn = bsn[in1d(bsn.T[0], selectedGenes)]
    # get global
    global_differences = {}
    geneGroups = {}
    
    for r, q, i in selectedClu :
        rr = geneGroups.get(r, [r])
        qq = geneGroups.pop(q, [q])
        for r2 in rr :
            g1 = geneInGenomes[r2]
            for q2 in qq :
                g2 = geneInGenomes[q2]
                key = tuple(sorted([g1, g2]))
                if key in global_differences :
                    global_differences[key].append(i)
                else :
                    global_differences[key] = [i]
        geneGroups[r] = rr + qq
    for r, q, i in selectedBsn :
        rr = geneGroups.get(r, [r])
        qq = geneGroups.pop(q, [q])
        for r2 in rr :
            g1 = geneInGenomes[r2]
            for q2 in qq :
                g2 = geneInGenomes[q2]
                if g1 != g2 : 
                    key = tuple(sorted([g1, g2]))
                    if key in global_differences :
                        global_differences[key].append(i)
                    else :
                        global_differences[key] = [i]
    for pair, data in global_differences.items() :
        diff = np.log(1.005-np.array(data)/10000.)
        mean_diff2 = np.mean(diff)
        mean_diff = min(max(mean_diff2, np.log(0.02)), np.log(0.5))
        sigma = min(max(np.sqrt(np.mean((diff - mean_diff2)**2)), np.log(1.6)), np.log(2.4))
        global_differences[pair] = (np.exp(mean_diff), sigma)
    return pd.DataFrame(list(global_differences.items())).values

def add_args(a) :
    import argparse
    parser = argparse.ArgumentParser(description='''
PEPPAN
(1) Retieve genes and genomic sequences from GFF files and FASTA files.
(2) Group genes into clusters using mmseq.
(3) Map gene clusters back to genomes. 
(4) Discard paralogous alignments.
(5) Discard orthologous clusters if they had regions which overlapped with the regions within other sets that had greater scores.
(6) Re-annotate genomes using the remained of orthologs. 
''', formatter_class=argparse.RawTextHelpFormatter)
    parser.add_argument('GFFs', metavar='GFF', help='[REQUIRED] GFF files containing both annotations and sequences. \nIf you have sequences and GFF annotations in separate files, \nthey can also be defined as: <GFF>,<fasta>', nargs='*')
    parser.add_argument('-p', '--prefix', help='[Default: PEPPAN] prefix for the outputs. ', default='PEPPAN')
    parser.add_argument('-g', '--genes', help='[optional] comma delimited filenames of fasta files containing additional genes. ', default='')
    parser.add_argument('-P', '--priority', help='[optional] comma delimited, ordered list of GFFs or gene fasta files that are more reliable than others. \nGenes contained in these files are preferred in all stages.', default='')
    parser.add_argument('-t', '--n_thread', help='[Default: 8] Number of threads to use. Default: 8', default=8, type=int)
    
    parser.add_argument('-o', '--orthology', metavar='nj,ml,sbh', help='[Default: nj] Method to define orthologous groups. \nnj [default], ml (for small dataset) or sbh (extremely large datasets)', default='nj')
    parser.add_argument('-n', '--noNeighborCheck', help='[Default: False] Flag to disable checking of neighborhood for paralog splitting. ', default=False, action='store_true')

    parser.add_argument('--min_cds', help='[Default: 120] Minimum length for a gene to be used in similarity searches.', default=120., type=float)
    parser.add_argument('--incompleteCDS', help="[Default: ''] Allowed types of imperfection for reference genes. \n's': allows unrecognized start codon. \n'e': allows unrecognized stop codon. \n'i': allows stop codons in the coding region. \n'f': allows frameshift in the coding region. \nMultiple keywords can be used together. e.g., use 'sife' to allow random sequences.", default='')
    parser.add_argument('--gtable', help='[Default: 11] Translate table to Use. Only support 11 and 4 (for Mycoplasma)', default=11, type=int)

    parser.add_argument('--clust_identity', help='minimum identities of mmseqs clusters. Default: 0.9', default=0.9, type=float)
    parser.add_argument('--clust_match_prop', help='minimum matches in mmseqs clusters. Default: 0.8', default=0.8, type=float)

    parser.add_argument('--nucl', dest='noDiamond', help='disable Diamond search. Fast but less sensitive when nucleotide identities < 0.9', default=False, action='store_true')
    parser.add_argument('--match_identity', help='minimum identities in BLAST search. Default: 0.5', default=0.5, type=float)
    parser.add_argument('--match_prop', help='minimum match proportion for normal genes in BLAST search. Default: 0.5', default=0.5, type=float)
    parser.add_argument('--match_len', help='minimum match length for normal genes in BLAST search. Default: 250', default=250., type=float)
    parser.add_argument('--match_prop1', help='minimum match proportion for short genes in BLAST search. Default: 0.8', default=0.8, type=float)
    parser.add_argument('--match_len1', help='minimum match length for short genes in BLAST search. Default: 100', default=100., type=float)
    parser.add_argument('--match_prop2', help='minimum match proportion for long genes in BLAST search. Default: 0.4', default=0.4, type=float)
    parser.add_argument('--match_len2', help='minimum match length for long genes in BLAST search. Default: 400', default=400., type=float)
    parser.add_argument('--match_frag_prop', help='minimum proportion of each fragment for fragmented matches. Default: 0.25', default=0.25, type=float)
    parser.add_argument('--match_frag_len', help='minimum length of each fragment for fragmented matches. Default: 50', default=50., type=float)
    
    parser.add_argument('--link_gap', help='consider two fragmented matches within N bases as a linked block. Default: 600', default=600., type=float)
    parser.add_argument('--link_diff', help='form a linked block when the covered regions in the reference gene \nand the queried genome differed by no more than this value. Default: 1.5', default=1.5, type=float)

    parser.add_argument('--allowed_sigma', help='Allowed number of sigma for paralogous splitting. \nThe larger, the more variations are kept as inparalogs. Default: 3.', default=3., type=float)
    parser.add_argument('--pseudogene', help='A match is reported as a pseudogene if its coding region is less than a proportion of the reference gene. Default: 0.8', default=.8, type=float)
    parser.add_argument('--untrusted', help='FORMAT: l,p; A gene is not reported if it is not greater than "l" and present in less than "p" of GFF files. Default: 450,0.35', default='450,0.35')
    parser.add_argument('--continue', help='continue from a previously stopped run.', default=False, action='store_true')

    #parser.add_argument('--intron', help="Enable this to allow multiple CDSs being concatenated if they are under the same name. This is still under development. ", default=False, action='store_true')
    parser.add_argument('--feature', help='feature to extract. Be cautious to change this value. DEFAULT: CDS', default='CDS')
    parser.add_argument('--noncoding', help='Set to noncoding mode. This is still under development. Equals to \n"--nucl --incompleteCDS sife"', default=False, action='store_true')
    parser.add_argument('--metagenome', help='Set to metagenome mode. This is still under development. Equals to \n"--nucl --incompleteCDS sife --clust_identity 0.99 --clust_match_prop 0.8 --match_identity 0.98 --orthology sbh"', default=False, action='store_true')
    parser.add_argument('--testunit', help='download four E. coli ST131 genomes for testing of PEPPAN.', default=False, action='store_true')

    params = parser.parse_args(a)
    if params.testunit :
        prepare_testunit()
        sys.exit(0)
    elif len(params.GFFs) == 0 :
        parser.print_help()
        sys.exit(0)
    params.match_frag_len = min(params.min_cds, params.match_frag_len)
    params.match_len1 = min(params.min_cds, params.match_len1)
    params.clust_match_prop = max(params.pseudogene, params.clust_match_prop, params.match_prop, params.match_prop1, params.match_prop2)
    if params.noncoding :
        params.noDiamond = True
        params.incompleteCDS = 'sife'
    elif params.metagenome :
        params.noDiamond = True
        params.incompleteCDS = 'sife'
        params.clust_identity = 0.99
        params.clust_match_prop = 0.8
        params.match_identity = 0.98
        params.orthology = 'sbh'
    params.untrusted = [float(p) for p in params.untrusted.split(',')]
    params.incompleteCDS = params.incompleteCDS.lower()
    if params.noNeighborCheck :
        params.self_id = 0.002
    else :
        params.self_id = 0.005
    return params

def prepare_testunit() :
    assert not os.path.exists('examples'), 'You have already got something named "examples". Remove it or go to a different folder for the testing.'
    os.makedirs('examples')
    import urllib.request
    urllib.request.urlretrieve('https://github.com/zheminzhou/PEPPA/blob/master/examples/GCF_000010485.combined.gff.gz?raw=true', \
                               'examples/GCF_000010485.combined.gff.gz')
    urllib.request.urlretrieve('https://github.com/zheminzhou/PEPPA/blob/master/examples/GCF_000214765.combined.gff.gz?raw=true', \
                               'examples/GCF_000214765.combined.gff.gz')
    urllib.request.urlretrieve('https://github.com/zheminzhou/PEPPA/blob/master/examples/GCF_001566635.combined.gff.gz?raw=true', \
                               'examples/GCF_001566635.combined.gff.gz')
    urllib.request.urlretrieve('https://github.com/zheminzhou/PEPPA/blob/master/examples/GCF_001577325.combined.gff.gz?raw=true', \
                               'examples/GCF_001577325.combined.gff.gz')
    sys.stderr.write('Folder "examples" has been created with four GFF files downloaded. \nRun:\n')
    sys.stderr.write('$ PEPPAN -p examples/ST131 -P examples/GCF_000010485.combined.gff.gz examples/*.gff.gz\n')
    sys.stderr.write('To test the main program. And then run:\n')
    sys.stderr.write('$ PEPPAN_parser -g examples/ST131.PEPPAN.gff -s examples/PEPPAN_out -t -c -a 95\n')
    sys.stderr.write('To test PEPPAN_parser\n')

def encodeNames(genomes, genes, geneFiles, labelFile, reuse=False) :
    taxon = {g[0] for g in genomes.values()}
    if reuse and os.path.exists(labelFile) :
        labels = dict(pd.read_csv(labelFile, header=None, na_filter=False).values.tolist())
    else :
        labels = {label:labelId for labelId, label in enumerate( sorted(set(list(taxon) + list(genomes.keys()) + list(genes.keys()) + geneFiles.split(',')) ) )}
        pd.DataFrame(sorted(labels.items(), key=lambda v:v[1])).to_csv(labelFile, header=False, index=False)
    genes = { labels[gene]:[labels.get(info[0], -1), labels.get(info[1], -1)] + info[2:] for gene, info in genes.items() }
    genomes = { labels[genome]:[labels[info[0]]] + info[1:] for genome, info in genomes.items() }
    return genomes, genes, labels

def iterClust(prefix, genes, geneGroup, params) :
    identity_target = params['identity']
    g = genes
    iterIden = np.round(np.arange(1., identity_target-0.005, -0.01), 5)
    for iden in iterIden :
        params.update({'identity':iden, 'coverage':np.round(params['coverage'], 2)})
        iden2 = min(1., iden+0.005)
        g, clust = getClust(prefix, g, params)
        exemplarNames = readFasta(g, headOnly=True)
        gp = pd.read_csv(clust, sep='\t').values
        logger('Iterative clustering. {0} exemplars left with identity = {1}'.format(len(exemplarNames), iden))
        for g1, g2 in gp[gp.T[0]!=gp.T[1]] :
            r, q = (g1, g2) if str(g1) in exemplarNames else (g2, g1)
            geneGroup.append([r, q, int(iden2*10000)])
    np.save('{0}.clust.npy'.format(prefix), np.array(geneGroup, dtype=int))
    return g

def async_writeOut(mat_out, matFile, outFile, labelFile) :
    encodes = pd.read_csv(labelFile, header=None, na_filter=False).values.tolist()
    encodes = np.array([n for i, n in sorted([[i, n] for n, i in encodes])])
    outPos = np.ones(16, dtype=bool)
    outPos[[0,3,4,5,10,15]] = False
    
    mat_id, group_id = 0, 0
    mat_conn = None
    import time
    with open(outFile, 'w') as fout :
        while True :
            while mat_id < len(mat_out) :
                if not mat_conn :
                    mat_conn = MapBsn(matFile)
                mat_id2 = min(len(mat_out), mat_id + 1000)
                mat_out2 = mat_out[mat_id:mat_id2]
                for i in range(mat_id, mat_id2) :
                    mat_out[i] = None
                gids = {grp[5]:None for pangene, gene, min_rank, mat in mat_out2 for grp in mat}
                p = [-1, None]
                for gid in sorted(gids) :
                    if p[0] != int(gid/1000) :
                        p = [int(gid/1000), mat_conn.get(int(gid/1000))]
                    gids[gid] = p[1][gid%1000]
                mat_id = mat_id2
                for pangene, gene, min_rank, mat in mat_out2 :
                    if len(mat) == 0 :
                        return
                    for grp in mat :
                        group_id += 1
                        m = gids[int(grp[5])]
                        m.T[1] = encodes[m.T[1].astype(int)]
                        for g in m :
                            gg = g[outPos].astype(str).tolist()
                            fout.write('{0}\t{1}\t{2}\t{3}\t{4}\t{5}\n'.format(pangene, min_rank, group_id, encodes[grp[1]], gene, '\t'.join(gg)))
                    del mat
            time.sleep(1)
    return

pool, pool2, mat_out = None, None, None
def ortho() :
    global params
    logger("COMMAND: {0}".format(' '.join(sys.argv)))
    params.update(add_args(sys.argv[1:]).__dict__)
    params.update(externals)

    global pool, pool2
    pool = Pool(params['n_thread'])
    pool2 = Pool(params['n_thread'])
    
    genomes, genes = readGFF(params['GFFs'], params['feature'], params['gtable'])
    genes = addGenes(genes, params['genes'], params['gtable'])
    
    params['encode'] = params['prefix']+'.encode.csv'
    genomes, genes, encodes = encodeNames(genomes, genes, params['genes'], params.get('encode', None), params['continue'])
    priorities = load_priority(params.get('priority', ''), genes, encodes)
    geneInGenomes = { g:i[0] for g, i in genes.items() }
    
    if len(genes) == 0 :
        logger('Did not find any gene from the inputs. Please check the format of the GFF files.')
        sys.exit(1)

    params['old_prediction'] = params['prefix']+'.old_prediction.npz'
    if not params['continue'] or not os.path.isfile(params['old_prediction']) :
        params['continue'] = False
        old_predictions = {}
        for n, g in genes.items() :
            if '' not in encodes or g[1] != encodes[''] :
                contig = str(g[1])
                if contig not in old_predictions :
                    old_predictions[contig] = []
                old_predictions[contig].append([n, g[2], g[3], g[4], g[5] if not len(g[6]) else 0 ])
        for gene, g in old_predictions.items() :
            old_predictions[gene] = np.array(sorted(g, key=lambda x:x[1]), dtype=object)
        with MapBsn(params['old_prediction'], 'w') as op :
            for n, g in old_predictions.items() :
                op.save(n, g)
        old_predictions.clear()
        del old_predictions, n, g
    
    params['prediction'] = params['prefix'] + '.Prediction'
    if not params['continue'] or not os.path.isfile(params['prediction']) :

        params['clust'] = params['prefix'] + '.clust.exemplar'
        if not params['continue'] or not os.path.isfile(params['clust']) :
            params['continue'] = False
            params['genes'], groups = writeGenes('{0}.genes'.format(params['prefix']), genes, priorities)
            genes.clear()
            del genes
            logger('Run MMSeqs linclust to get exemplar sequences. Params: {0} identities and {1} align ratio'.format(params['clust_identity'], params['clust_match_prop']))
            params['clust'] = iterClust(params['prefix'], params['genes'], groups, dict(identity=params['clust_identity'], coverage=params['clust_match_prop'], n_thread=params['n_thread'], translate=False))
        
        params['self_bsn'] = params['prefix']+'.self_bsn.npy'
        if not params['continue'] or not os.path.isfile(params['self_bsn']) :
            params['continue'] = False
            np.save(params['self_bsn'], get_similar_pairs(params['clust'], priorities, params))
        genes = { int(n):s for n, s in readFasta(params['clust']).items()}
        logger('Obtained {0} exemplar gene sequences from {1}'.format(len(genes), params['clust']))

        params['global'] = params['prefix']+'.global.npy'
        if not params['continue'] or not os.path.isfile(params['global']) :
            params['continue'] = False
            np.save(params['global'], \
                    get_global_difference(get_gene_group(params['clust'], params['self_bsn']), \
                                          params['clust'], params['self_bsn'], geneInGenomes, nGene=1000) )
            
        params['map_bsn']= params['prefix']+'.map_bsn'
        if not params['continue'] or not os.path.isfile(params['map_bsn']+'.tab.npz') :            
            params['continue'] = False
            with MapBsn(params['map_bsn']+'.tab.working.npz', 'w') as tab_conn, MapBsn(params['map_bsn']+'.seq.npz', 'w') as seq_conn, MapBsn(params['map_bsn']+'.mat.npz', 'w') as mat_conn, MapBsn(params['map_bsn']+'.conflicts.npz', 'w') as clf_conn :
                get_map_bsn(params['prefix'], params['clust'], genomes, params['self_bsn'], params['old_prediction'], tab_conn, seq_conn, mat_conn, clf_conn, params.get('orthology', 'sbh') != 'sbh')
            shutil.move(params['map_bsn']+'.tab.working.npz', params['map_bsn']+'.tab.npz')
        pool.close()
        pool.join()

        global mat_out
        #mat_out = []
        mat_out = Manager().list([])
        writeProcess = Process(target=async_writeOut, args=(mat_out, params['map_bsn']+'.mat.npz', params['prediction']+'.working', params['encode']))
        writeProcess.start()
        gene_scores = initializing(params['map_bsn'], params.get('global', None))
        with MapBsn(params['map_bsn']+'.tab.npz') as tab_conn :
            filt_genes(tab_conn, np.load(params['self_bsn'], allow_pickle=True), params['global'], params['map_bsn']+'.conflicts.npz', priorities, gene_scores, encodes)
        writeProcess.join()
        shutil.move(params['prediction']+'.working', params['prediction'])
    else :
        params['clust'] = params['prefix'] + '.clust.exemplar'
        if os.path.isfile(params['clust']) :
            genes = { int(n):s for n, s in readFasta(params['clust']).items()}
        else :
            genes = {n:s[-1] for n,s in genes.items() }
        
    if not params['noNeighborCheck'] :
        logger('Neigbhorhood based paralog splitting starts')
        params['prediction'] = synteny_resolver(params['prefix'], params['prediction'], 2)
        logger('Neigbhorhood based paralog splitting finishes')
    if os.path.isfile(params['old_prediction']) :
        old_predictions = dict(np.load(params['old_prediction'], allow_pickle=True)) if 'old_prediction' in params else {}
    revEncode = {e:d for d, e in encodes.items()}
    old_predictions = { revEncode[int(contig)]:[np.concatenate([ [revEncode[g[0]]], g[1:]]) for g in genes ] for contig, genes in old_predictions.items() if int(contig)>=0 and genes[0][0] >= 0}
    
    params['clust'] = params['prefix'] + '.clust.exemplar'
    params['self_bsn'] = params['prefix']+'.self_bsn.npy'
    write_output(params['prefix'], params['prediction'], genomes, genes, encodes, old_predictions, params['pseudogene'], params['untrusted'], params['gtable'], params.get('clust', None), params.get('self_bsn', None))
    pool2.close()
    pool2.join()
    
if __name__ == '__main__' :
    ortho()
```
