## Supplemental Code S1 for "Accurate reconstruction of bacterial pan- and core- genomes with PEPPAN": PEPPAN_parser.py

```
#!/usr/bin/env python
import sys, os, re, numpy as np, subprocess
from ete3 import Tree
from collections import defaultdict
from scipy.optimize import curve_fit
from scipy.stats.distributions import t
try:
    from .modules.configure import xrange, uopen, externals, logger
except :
    from modules.configure import xrange, uopen, externals, logger

def func_powerlaw(x, m, c ):
    return x**m * c

def splitGFF(gff, folder, prefix) :
    if not os.path.isdir(folder) :
        try :
            os.makedirs(folder)
        except FileExistsError :
            raise FileExistsError('Fail to create the output folder for GFF files')
    prev, fout = None, None
    with uopen(gff) as fin :
        for line in fin :
            if line.startswith('#') :
                continue
            header = line.split(':', 1)[0]
            if header != prev :
                prev = header
                if fout :
                    fout.close()
                fout = uopen(os.path.join(folder, '{0}.{1}.gff.gz'.format(prefix.rsplit('/', 1)[-1], header)), 'w')
                fout.write('#!gff-version 3\n#!annotation-source PEPPA from enterobase.warwick.ac.uk\n')
            fout.write(line)
    fout.close()
    logger('GFF files are saved under folder {0}'.format(folder))

def getOrtho(fname, noPseudogene=False) :
    ids = set([])
    groups = defaultdict(dict)
    with uopen(fname) as fin :
        for line in fin :
            if line.startswith('#') :
                continue            
            part = line.strip().split('\t')
            if part[1] == 'CDS' or (part[1] == 'pseudogene' and not noPseudogene) :
                ID = re.findall(r'ID=([^;]+);', part[8])[0]
                if ID in ids : continue
                
                genome = part[0].split(':', 1)[0]
                grp = groups[genome]
                ortho = re.findall(r'inference=ortholog_group:([^;]+)', part[8])[0]
                orthos = [ [o, int(aId) if aId.isdigit() else -1] for o, aId in [ o.split(':')[1:3] for o in ortho.split(',') ]]

                for ortho, aId in orthos :
                    if ortho not in grp :
                        grp[ortho] = {ID:aId}
                    else :
                        grp[ortho][ID] = aId
                #grp.update({ ortho:([aId, ID] if ortho not in grp else -2) for ortho, aId in orthos })
                ids.add(ID)
    return groups

def writeCurve(prefix, groups, pseudogene=True, n_iter=1000) :
    gtype = ['CDSs', 'genes'][int(pseudogene)]
    encode = {}
    for grp in groups.values() :
        for g in grp.keys() :
            if g not in encode :
                encode[g] = len(encode)
    mat = [ set( encode[g] for g, c in grp.items() ) for grp in groups.values() ]
    mat = [np.array(list(m)) for m in mat]
    x = np.arange(len(mat))+1

    curves = np.zeros([len(mat), n_iter, 2], dtype=int)
    for ite in np.arange(n_iter) :
        np.random.shuffle(mat)
        genes = np.zeros(len(encode), dtype=int)
        for id, m in enumerate(mat) :
            genes[m] += 1
            curves[id, ite, :] = (np.sum(genes>=1), np.sum(genes>=(id+1)) )
    y = np.mean(curves, 1)

    yopt1, ycov1 = curve_fit(func_powerlaw, x, y[:, 0], maxfev=3000)
    tval1 = t.ppf(1.0 - .05/2.0, max(0, y.shape[0] - 2))
    yci1 = np.diag(ycov1)**0.5*tval1

    yopt2, ycov2 = curve_fit(func_powerlaw, x[1:], y[1:, 0] - y[:-1, 0], maxfev=3000)
    tval2 = t.ppf(1.0 - .05/2.0, max(0, y.shape[0] - 1 - 2))
    yci2 = np.diag(ycov2)**0.5*tval2

    yopt3, ycov3 = curve_fit(func_powerlaw, x, y[:, 1], maxfev=3000)
    tval3 = t.ppf(1.0 - .05/2.0, max(0, y.shape[0] - 2))
    yci3 = np.diag(ycov3)**0.5*tval3

    with open('{0}_content.curve'.format(prefix), 'w') as fout :
        fout.write('#! No. genomes: {0}\n'.format(len(groups)))
        fout.write('#! Ave. {1} per genome: {0:.03f}\n'.format(np.mean([m.size for m in mat]), gtype))
        fout.write('#! No. pan {1}: {0}\n'.format(len(encode), gtype))
        fout.write('#! No. core {1}: {0}\n'.format(curves[-1, 0, 1], gtype))
        fout.write('#! Heaps\' law model in DOI: 10.1016/j.mib.2008.09.006:\tGamma={0:.03f} +/- {1:.03f}, Kappa={2:.03f} +/- {3:.03f}, ~{4:.03f} new genes per new genome.\n'.format(\
            yopt1[0], yci1[0], yopt1[1], yci1[1], (x[-1] + 1) ** yopt1[0] * yopt1[1] - x[-1] ** yopt1[0] * yopt1[1]))
        fout.write('#! Power law model in DOI: 10.1016/j.mib.2008.09.006:\tAlpha={0:.03f} +/- {1:.03f}, Kappa={2:.03f} +/- {3:.03f}, ~{4:.03f} new genes per new genome.\n'.format( \
            -yopt2[0], yci2[0], yopt2[1], yci2[1], (x[-1] + 1) ** yopt2[0] * yopt2[1]))
        fout.write('#! Power law model for the core genome:              \tAlpha={0:.03f} +/- {1:.03f}, Kappa={2:.03f} +/- {3:.03f}, ~{4:.03f} fewer core genes per new genome.\n'.format( \
            -yopt3[0], yci3[0], yopt3[1], yci3[1], -(x[-1] + 1) ** yopt3[0] * yopt3[1] + x[-1] ** yopt3[0] * yopt3[1]))
        fout.write('#No. genome\t(Pan-genome)Median\t2.5%\t97.5%\t|\t(Core-genome)Median\t2.5%\t97.5%\n')
        summary = np.zeros([len(mat), 2, 5], dtype=int)
        for id, curve in enumerate(curves) :
            pan  = np.sort(curve.T[0])
            core = np.sort(curve.T[1])
            pan_s =  [  pan[int(pan.size*0.025)],   pan[int(pan.size*0.5)],   pan[int(pan.size*0.975)]  ]
            core_s = [ core[int(core.size*0.025)], core[int(core.size*0.5)], core[int(core.size*0.975)] ]
            fout.write('{0}\t{1[1]}\t{1[0]}\t{1[2]}\t|\t{2[1]}\t{2[0]}\t{2[2]}\n'.format(id+1, pan_s, core_s))
    logger('Curves for {1} are saved in {0}_content.curve'.format(prefix, ['CDS', 'all genes'][int(pseudogene)]))
    return summary

def writeMatrix(prefix, ortho) :
    genomes = sorted(ortho.keys())
    genes = defaultdict(int)
    for gs in ortho.values() :
        for g, i in gs.items() :
            genes[g] += 1
    presences = np.array(list(genes.values()))
    n = len(genomes)
    with open('{0}_content.summary_statistics.txt'.format(prefix), 'w') as fout :
        fout.write('Strict core genes\t(strains = 100%)\t{0}\n'.format(np.sum(presences >= n)))
        fout.write('Core genes\t(99% <= strains < 100%)\t{0}\n'.format(np.sum((1.*n > presences) & (presences >= 0.99*n))))
        fout.write('Soft core genes\t(95% <= strains < 99%)\t{0}\n'.format(np.sum((0.99*n > presences) & (presences >= 0.95*n))))
        fout.write('Shell genes\t(15% <= strains < 95%)\t{0}\n'.format(np.sum((0.95*n > presences) & (presences >= 0.15*n))))
        fout.write('Cloud genes\t(0% <= strains < 15%)\t{0}\n'.format(np.sum((0.15*n > presences) & (presences >= 0.0*n))))
        fout.write('Total genes\t(0% <= strains <= 100%)\t{0}\n'.format(presences.size))
    logger('Summary of the pan-genome is saved in {0}_content.summary_statistics.txt'.format(prefix))
    genes = [ g[0] for g in sorted(genes.items(), key=lambda x:(-x[1], x[0])) ]
    with open('{0}_content.Rtab'.format(prefix), 'w') as fout, open('{0}_content.csv'.format(prefix), 'w') as fout2 :
        fout.write('\t'.join(['Gene']+genomes)+'\n')
        fout2.write(','.join(['Gene']+genomes)+'\n')
        for g in genes :
            mat = [ str(len(ortho[genome].get(g, {}))) for genome in genomes ]
            fout.write('\t'.join([g] + mat)+'\n')
            mat2 = [ ';'.join(sorted(ortho[genome].get(g, {}).keys())) for genome in genomes ]
            fout2.write(','.join([g] + mat2)+'\n')
    logger('Gene content matrix is saved in {0}_content.csv'.format(prefix))
    logger('Gene presence matrix is saved in {0}_content.Rtab'.format(prefix))

def writeCGAV(prefix, ortho, pCore) :
    genomes = sorted(ortho.keys())
    minPresence = len(ortho) * pCore/100.
    genes = defaultdict(int)
    for gs in ortho.values() :
        for g, i in gs.items() :
            if len(i) == 1 and list(i.values())[0] > 0 :
                genes[g] += 1
    genes = sorted([ gene for gene, presence in genes.items() if presence >= minPresence ])
    profiles = []
    with open('{0}_CGAV.profile'.format(prefix), 'w') as fout :
        fout.write('\t'.join(['#Genome'] + genes) + '\n')
        for genome in genomes :
            grp = ortho[genome]
            profiles.append([ ( max(list(grp[g].values())[0], 0) if len(grp.get(g, {})) == 1 else 0) for g in genes ] )
            fout.write('{0}\t{1}\n'.format( genome, '\t'.join([ str(allele) for allele in profiles[-1] ]) ))
    
    distances = getDistance( np.array(profiles) )
    with open('{0}_CGAV.dist'.format(prefix), 'w') as fout :
        fout.write('    {0}\n'.format(len(genomes)))
        for genome, dist in zip(genomes, distances) :
            fout.write('{0} {1}\n'.format(genome, ' '.join(dist.astype(str))))
    
    subprocess.Popen('''{rapidnj} -i pd {0}_CGAV.dist | sed "s/'//g"  > {0}_CGAV.nwk'''.format(prefix, **externals), stderr=subprocess.PIPE, shell=True).wait()
    logger('Core gene allelic variation profile is saved in {0}_CGAV.profile'.format(prefix))
    logger('Core gene allelic variation tree is saved in {0}_CGAV.nwk'.format(prefix))
    
def getDistance(profiles) :
    distances = np.zeros([profiles.shape[0], profiles.shape[0]], dtype=float)
    contents = (profiles > 0)
    for i, p in enumerate(profiles) :
        c, j = contents[i], i+1
        shared, presence = np.sum((p == profiles[j:]) & (contents[j:]*c), 1)+0.5, np.sum(contents[j:]*c, 1)+1.
        distances[i, j:] = distances[j:, i] = -np.log(shared/presence)
    return distances

def writeTree(prefix, ortho) :
    genes = sorted({g for gs in ortho.values() for g in gs})
    with open('{0}_content.fas'.format(prefix), 'w') as fout :
        for genome, content in ortho.items() :
            fout.write('>{0}\n{1}\n'.format(genome, ''.join([['A','T'][int(g in content)] for g in genes ])))
    tree = subprocess.Popen('{fasttree} -quiet -nt {0}_content.fas'.format(prefix, **externals).split(), stdout=subprocess.PIPE, universal_newlines=True).communicate()[0]

    tree = Tree(tree, format=1)
    for n in tree.traverse() :
        n.dist *= len(genes)
    tree.write(outfile='{0}_content.nwk'.format(prefix), format=1)
    logger('Gene content tree is saved in {0}_content.nwk'.format(prefix))

def PEPPA_parser() :
    param = arg_parser(sys.argv[1:])
    if param.split :
        splitGFF(param.gff, param.split, param.prefix)
    if param.matrix or param.tree or param.curve or param.cgav >= 0 :
        target = ['gene', 'CDS'][int(param.pseudogene)]
        prefix = '{0}.{1}'.format(param.prefix, target)
        ortho = getOrtho(param.gff, param.pseudogene)
        if param.matrix :
            writeMatrix(prefix, ortho)
        if param.tree :
            writeTree(prefix, ortho)
        if param.cgav >= 0 :
            writeCGAV(prefix, ortho, param.cgav)
        if param.curve :
            writeCurve(prefix, ortho, not param.pseudogene)
    return
    
def arg_parser(a) :
    import argparse
    parser = argparse.ArgumentParser(description='''
PEPPA_parser.py 
(1) reads <prefix>.PEPPA.gff file
(2) split it into individual GFF files
(3) draw a present/absent matrix
(4) create a tree based on gene presence
(5) draw rarefraction curves of all genes and only intact CDSs
''', formatter_class=argparse.RawTextHelpFormatter)
    parser.add_argument('-g', '--gff', help='[REQUIRED] generated PEPPA.gff file from PEPPA.py.', required=True)
    parser.add_argument('-p', '--prefix', help='[Default: Same prefix as the GFF input] Prefix for all outputs.', default=None)
    parser.add_argument('-s', '--split', help='[optional] A folder for splitted GFF files. ', default=None)
    parser.add_argument('-P', '--pseudogene', help='[Default: Use Pseudogene] Flag to ignore pseudogenes in all analyses. ', default=False, action='store_true')
    parser.add_argument('-m', '--matrix', help='[Default: False] Flag to NOT generate the gene present/absent matrix. ', default=True, action='store_false')
    parser.add_argument('-t', '--tree', help='[Default: False] Flag to generate the gene present/absent tree. ', default=False, action='store_true')
    parser.add_argument('-a', '--cgav', help='[Default: -1] Set to an integer between 0 and 100 to apply a Core Gene Allelic Variation tree. \nThe value describes %% of presence for a gene to be included in the analysis. \nThis is similar to cgMLST tree but without an universal scheme. ', default=-1, type=int)
    parser.add_argument('-c', '--curve', help='[Default: False] Flag to generate a rarefraction curve. ', default=False, action='store_true')
    params = parser.parse_args(a)
    
    assert params.split or params.matrix or params.tree or params.curve or params.cgav >= 0, 'At least one type of output needs to be specified. '
    if not params.prefix : 
        params.prefix = params.gff.rsplit('.gff', 1)[0]
    
    return params


if __name__ == '__main__' :
    PEPPA_parser()
```
