## Supplemental Code S1 for "Accurate reconstruction of bacterial pan- and core- genomes with PEPPAN": setup.py

```
import os, sys
from setuptools import setup, find_packages
#from PEPPA import __VERSION__
__VERSION__ = '1.0.5'

with open('README.rst', encoding='utf-8') as f:
    long_description = f.read()

setup(
    name='PEPPAN',
    version= __VERSION__,
    #scripts=['PEPPA.py'] ,
    author="Zhemin Zhou",
    author_email="",
    description="Phylogeny Enhanded Prediction of PAN-genome https://doi.org/10.1101/2020.01.03.894154",
    long_description=long_description,
    long_description_content_type="text/markdown",
    url="https://github.com/zheminzhou/PEPPAN",
    packages = ['PEPPAN'],
    package_dir = {'PEPPAN':'.'},
    keywords=['bioinformatics', 'microbial', 'genomics', 'MLST', 'pan-genome'],
    install_requires=['ete3>=3.1.1', 'numba>=0.38.0', 'numpy>=1.18.1', 'pandas>=0.24.2', 'scipy>=1.3.2'],
    include_package_data=True,
    entry_points={
        'console_scripts': [
            'PEPPAN = PEPPAN.PEPPAN:ortho',
            'PEPPAN_parser = PEPPAN.PEPPAN_parser:PEPPA_parser',
    ]},
    package_data={'PEPPAN': ['LICENSE', 'README.*', 'dependencies/*', 'modules/*.py']},
    classifiers=[
        'Development Status :: 5 - Production/Stable',
        "Programming Language :: Python :: 3.5",
        "Programming Language :: Python :: 3.6",
        "Programming Language :: Python :: 3.7",
        "Programming Language :: Python :: 3.8",
        'License :: OSI Approved :: GNU General Public License v3 (GPLv3)',
        "Operating System :: OS Independent",
    ],
 )
```
