## Supplementary figures and images for "Accurate reconstruction of bacterial pan- and core- genomes with PEPPAN"

### file.png

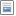

### minus.png

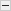

### plus.png

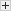
