## Supplemental Code S2 for "Accurate reconstruction of bacterial pan- and core- genomes with PEPPAN": README.md

# SimPan
A simulator of pan-genome for bacterial population. 

* SimPan simulates whole genomic sequences rather than only genes. 
* SimPan implements different evolutionary model for core-genome and accessory genomes. 

SimPan runs in Python with versions >= 3.5 and requires two libraries:

* numpy
* ete3


SimPan also depends on two published simulators:

* [SimBac](https://github.com/tbrown91/SimBac)
* [indelible](http://abacus.gene.ucl.ac.uk/software/indelible/)


Binary files for both dependencies have also been distributed in this repository. Both are compiled in Ubuntu 16.04. 
If they can not run in your system, or if you prefer a different version of these dependencies, please put the binary files in your environment variable PATH. SimPan will find and use them. 

In brief, SimPan 

1. Uses SimBac to simulate a global phylogeny and recombination events of the bacterial genomes.
2. Generate gene contents for both core- and accessory genomes by random indel events. 
3. Uses indelible to fill in sequences for these pan genes. 

Note that indelible can be very slow. If you are only interested in the gene content, use '--noSeq' as an early stop. 

## EXAMPLE:
Use
```
python SimPan.py --aveSize 50 --nBackbone 30 --nMobile 1000 -p test --genomeNum 10
```
To simulate 10 genomes with average of 50 genes each. 

### OUTPUTS:
#### test.gene.content.tsv
```
#ID     0       1       2       3       4       5       6       7       8       9
1       28_0    28_0    28_0    28_0    28_0    28_0    28_0    28_0    28_0    28_0
2       5_0     5_0     5_0     5_0     5_0     5_0     5_0     5_0     5_0     5_0
3       -       -       -       -       -       -       128_0   -       -       -
4       -       -       -       -       -       -       719_0   -       -       -
5       -       -       -       -       -       -       53_0    -       -       -
6       -       -       -       -       -       -       492_0   -       -       -
7       8_0     8_0     8_0     8_0     8_0     8_0     8_0     8_0     8_0     8_0
```
Each column except for the first one shows one simulated genome. Each row shows one gene. "-" are missing genes. The gene names suggest their homologous groups and orthologous sub-groups. e.g., 28_0 and 5_0 are in different homologous groups and thus unlikely to be similar, whereas 28_0 and 28_1 belongs to the same homologous group but in different orthologous sub-group. They are distantly related. 

#### test.aligned.fasta
This is a multi-sequence alignment of all the simulated genomes. 

#### test.aligned.tbl
```
61      120     -       28      0
236     295     -       5       0
416     475     -       128     0
510     569     -       719     0
651     710     +       53      0
774     833     -       492     0
```
This file show the coordinates of genes in the genomic alignment (test.aligned.fasta). The five columns are:

1. start site
2. end site
3. direction
4. homologous group designation
5. orthologous sub-group designation

#### test_0.gff, test_1.gff, test_2.gff, ...
The gene annotation file for simulated genomes in GFF3 format

#### test_0.fna & test_0.tbl
The source files for tbl2asn tool from NCBI. GBK file can be generated using these files. 

## USAGE:
```
$ python SimPan.py -h
usage: SimPan.py [-h] [-p PREFIX] [--genomeNum GENOMENUM] [--geneLen GENELEN]
                 [--igrLen IGRLEN] [--backboneBlock BACKBONEBLOCK]
                 [--mobileBlock MOBILEBLOCK] [--operonBlock OPERONBLOCK]
                 [--aveSize AVESIZE] [--nBackbone NBACKBONE] [--nCore NCORE]
                 [--nMobile NMOBILE] [--pBackbone PBACKBONE]
                 [--pMobile PMOBILE] [--tipAccelerate TIPACCELERATE]
                 [--rec REC] [--recLen RECLEN] [--seqRec SEQREC]
                 [--insRec INSREC] [--delRec DELREC] [--noSeq]
                 [--idenOrtholog IDENORTHOLOG] [--idenParalog IDENPARALOG]
                 [--idenDuplication IDENDUPLICATION] [--indelRate INDELRATE]
                 [--indelLen INDELLEN] [--freqStart FREQSTART]
                 [--freqStop FREQSTOP]

SimPan is a simulator for bacterial pan-genome.
Global phylogeny and tree distortions are derived from SimBac and the gene and intergenic sequences are simulated using indelible.

optional arguments:
  -h, --help            show this help message and exit
  -p PREFIX, --prefix PREFIX
                        prefix for all intermediate files and outputs
  --genomeNum GENOMENUM
                        No of genome in population [DEFAULT: 20]
  --geneLen GENELEN     [negative bionomial with r=2] mean,min,max sizes of genes [DEFAULT: 900,150,6000]
  --igrLen IGRLEN       [negative bionomial] mean,min,max sizes of intergenic regions [DEFAULT: 50,0,300]
  --backboneBlock BACKBONEBLOCK
                        [geometric] mean,min,max number of backbone genes per block [DEFAULT: 3,0,30]
  --mobileBlock MOBILEBLOCK
                        [geometric] mean,min,max number of mobile genes per block [DEFAULT: 10,0,100]
  --operonBlock OPERONBLOCK
                        [geometric] mean,min,max number of continuous genes that share the same coding strand [DEFAULT: 3,0,15]
  --aveSize AVESIZE     average gene number per genome (greater than nBackbone). [DEFAULT: 4500]
  --nBackbone NBACKBONE
                        number of backbone genes (present in common ancestor) per genome. [DEFAULT: 4000]
  --nCore NCORE         sizea of core gene (smaller than the size of backbone genes). [DEFAULT: 3500]
  --nMobile NMOBILE     size of mobile gene pool for accessory genome. [DEFAULT: 20000]
  --pBackbone PBACKBONE
                        propotion of paralogs in backbone (core) genes. [DEFAULT: 0.05]
  --pMobile PMOBILE     propotion of paralogs in mobile (accessory) genes. [DEFAULT: 0.4]
  --tipAccelerate TIPACCELERATE
                        grandient increasing of gene indels in recent times. [DEFAULT: 100]
  --rec REC             expected coverage of homoplastic events in pairwise comparisons. [DEFAULT: 0.05]
  --recLen RECLEN       expected size of homoplastic events. [DEFAULT: 1000]
  --seqRec SEQREC       Use homoplastic events to infer sequences. Use 0 to disable [DEFAULT: 1]
  --insRec INSREC       Use homoplastic events to infer gene insertions. Use 0 to disable [DEFAULT: 1]
  --delRec DELREC       Use homoplastic events to infer gene deletions. Use 0 to disable [DEFAULT: 1]
  --noSeq               Do not infer sequence but only the gene presence/absence. [DEFAULT: False]
  --idenOrtholog IDENORTHOLOG
                        average nucleotide identities for orthologous genes. [DEFAULT: 0.98]
  --idenParalog IDENPARALOG
                        average nucleotide identities for paralogous genes. [DEFAULT: 0.6]
  --idenDuplication IDENDUPLICATION
                        average nucleotide identities for recent gene duplications. [DEFAULT: 0.995]
  --indelRate INDELRATE
                        average frequency of indel events relative to mutation rates. [DEFAULT: 0.01]
  --indelLen INDELLEN   average size of short indel events within each gene. [DEFAULT:130]
  --freqStart FREQSTART
                        frequencies of start codons of ATG,GTG,TTG. DEFAULT: 0.83,0.14,0.03
  --freqStop FREQSTOP   frequencies of stop codons of TAA,TAG,TGA. DEFAULT: 0.63,0.08,0.29
```
