## Supplemental Code S2 for "Accurate reconstruction of bacterial pan- and core- genomes with PEPPAN": SimPan.py

```
import subprocess, sys, os, numpy as np, ete3, re, glob, shutil
import tempfile
from multiprocessing import Pool

complement = {'A':'T', 'T':'A', 'G':'C', 'C':'G', 'N':'N'}
def rc(seq, missingValue='N') :
    return ''.join([complement.get(s, missingValue) for s in reversed(seq.upper())])

externals = {
    'simbac' : shutil.which('SimBac') if shutil.which('SimBac') else os.path.join(os.path.dirname(os.path.realpath(__file__)), 'dependencies', 'SimBac'),
    'indelible' : shutil.which('indelible') if shutil.which('indelible') else os.path.join(os.path.dirname(os.path.realpath(__file__)), 'dependencies', 'indelible'),
}

def getEvent(trees, tipAcc, num) :
    results = []
    w = np.array([t[1] for t in trees]).astype(float)
    eventSources = np.random.choice(len(trees), size=num, p=w/np.sum(w))
    for tId, cnt in zip(*np.unique(eventSources, return_counts=True)) :
        tree = ete3.Tree(trees[tId][2])
        annotateTree(tree)
        indelTree(tree, tipAcc)

        nodes, weights = [], []
        for n in tree.get_descendants() :
            nodes.append(n.descendants)
            weights.append(n.dist)
        weights = np.array(weights)
        picks = np.random.choice(nodes, size=cnt, p=weights/np.sum(weights))
        results.extend(picks)
    return results
def annotateTree(tree) :
    for node in tree.traverse('postorder') :
        if node.is_leaf() :
            node.descendants = [int(node.name)]
            node.height = 0.0
        else :
            node.descendants = [n for c in node.get_children() for n in c.descendants]
            node.height = np.mean([c.height + c.dist for c in node.get_children()])
    return tree

def indelTree(tree, tipAccelerate) :
    lmbda = np.log(tipAccelerate) 
    for node in tree.traverse('postorder') :
        node.dist = lmbda*abs(np.exp(-lmbda*node.height) - np.exp(-lmbda*(node.height+node.dist)))

def borderedGeometric(mean, low, high, size) :
    data = np.random.geometric(p=1./mean, size=size)
    data[data < low] = low
    data[data > high] = high
    return data


def borderedNB(mean, low, high, size, r=2) :
    data = np.random.negative_binomial(n=r, p=(1. - mean/(2.+mean)), size=size)
    data[data < low] = low
    data[data > high] = high
    return data

def setGeneContent(items, itrees, dtrees, tipAccelerate, genomeNum, aveSize, nCore, backboneBlockSize, mobileBlockSize) :
    genomes = np.random.permutation(np.repeat(np.where(items.T[0] == 0)[0], genomeNum).reshape(-1, genomeNum))
    
    events = getEvent(dtrees, tipAccelerate, genomes.shape[0] )
    backBoneSite = np.random.choice(genomes.shape[0], genomes.shape[0])
    backBoneSize = borderedGeometric(backboneBlockSize[0], backboneBlockSize[1], backboneBlockSize[2], genomes.shape[0])
    
    coreIdx = np.ones(genomes.shape[0], dtype=bool)
    for b, s, l in zip(events, backBoneSite, backBoneSize) :
        e = s + l
        c1 = np.sum(coreIdx)
        
        if e <= genomes.shape[0] :
            coreIdx[s:e] = False
            c2 = np.sum(coreIdx)
            if abs(c1-nCore) < abs(c2-nCore) :
                break
            genomes[s:e, b] = -1
        else :
            l1 = genomes.shape[0] - s
            l2 = l - l1
            coreIdx[s:e] = False
            c2 = np.sum(coreIdx)
            if abs(c1-nCore) < abs(c2-nCore) :
                break            
            genomes[s:, b], genomes[:l2, b] = -1, -1
    
    genePool = np.random.permutation(np.where(items.T[0] == 1)[0])
    toAdd = aveSize*genomeNum - np.sum(genomes >= 0)
    events = getEvent(itrees, tipAccelerate, toAdd)
    mobileBlock = borderedGeometric(mobileBlockSize[0], mobileBlockSize[1], mobileBlockSize[2], toAdd)
    gInEvent = np.cumsum([n*len(e) for n, e in zip(mobileBlock, events)])
    
    events = events[:np.argmin(np.abs(gInEvent - toAdd))+1]
    mobiles = []
    for b, s, l in zip(events, np.random.choice(len(genePool), len(events)), mobileBlock) :
        mobile = np.empty([l, genomeNum], dtype=int)
        mobile[:] = -1
        e = s + l
        if e <= len(genePool) :
            mobile[:, b] = genePool[s:e, np.newaxis]
        else :
            l1 = len(genePool) - s
            l2 = l - l1
            mobile[:l1,  b] = genePool[s:,  np.newaxis]
            mobile[-l2:, b] = genePool[:l2, np.newaxis]
        mobiles.append(mobile)
        
    blk = (np.random.dirichlet(np.ones(len(mobiles)+1, dtype=int))*genomes.shape[0]+0.5).astype(int)
    genomes = np.split(genomes, np.cumsum(blk)[:-1])
    genomes = np.vstack([nn for n in zip(genomes[:-1], mobiles) for nn in n] + genomes[-1:])
    
    return genomes

def treeGenerator(prefix, items, genomeNum, rec, recLen) :
    genomeLength = int(np.sum(items[:, 2:]))
    
    r0 = float(recLen)/genomeLength
    r = np.log(1-rec)/np.log(1-r0)/2/genomeLength
    if r <= 0 :
        r = 0
    cmd = '{simbac} -N {0} -T 0 -R {1} -B {2} -c {3}.phylogeny -l {3}.local'.format(\
        genomeNum, r, genomeLength, prefix, **externals
    )

    sys.stdout.write(cmd+'\n')
    p = subprocess.Popen(cmd.split(), universal_newlines=True, stdout=subprocess.PIPE).wait()
    
    globaltrees, localtrees = [], []
    c = 0
    with open('{0}.phylogeny'.format(prefix), 'r') as fin :
        for line in fin :
            globaltrees.append([genomeLength, genomeLength, line.strip()])
    with open('{0}.local'.format(prefix), 'r') as fin :
        for line in fin :
            if line.startswith('[') :
                w, t = line.strip()[1:].split(']')
                w = int(w)
                c += w
                localtrees.append([c, w, t])

    os.unlink('{0}.local'.format(prefix))
    return globaltrees, localtrees
    
def itemShaper(items, geneLen, igrLen) :
    nOrtho = np.max((items.T[1]/10).astype(int))+1
    orthos = borderedNB(geneLen[0], geneLen[1], geneLen[2], nOrtho, r=2.)
    orthos *= 3
    oigrs1 = borderedGeometric(igrLen[0], igrLen[1], igrLen[2], nOrtho)
    oigrs2 = borderedGeometric(igrLen[0], igrLen[1], igrLen[2], nOrtho)
    genes = orthos[(items.T[1]/10).astype(int)]
    igrs1 = oigrs1[(items.T[1]/10).astype(int)]
    igrs2 = oigrs2[(items.T[1]/10).astype(int)]
    return np.hstack([items, igrs1[:, np.newaxis], genes[:, np.newaxis], igrs2[:, np.newaxis]])

def itemGenerator(nBackbone, nMobile, pBackbone, pMobile) :
    types = np.concatenate([np.zeros(nBackbone, dtype=int), np.ones(nMobile, dtype=int)])

    genes = []
    for n, p in zip([nBackbone, nMobile], [pBackbone, pMobile]) :
        p2 = p - np.sqrt(p*(p+3)) + 1
        nOrtho = int(n / (1+2*p2) + 0.5)
        orthoGene = np.arange(nOrtho)
        gene = np.sort(np.concatenate([np.random.choice(np.repeat(orthoGene, 2), n - nOrtho, replace=False), orthoGene]))*10
        for i, g in enumerate(gene[1:]) :
            if int(g/10) == int(gene[i]/10) :
                gene[i+1] = gene[i]+1
        if len(genes) :
            gene += np.max(genes[-1])+10
        genes.append(gene)
    genes = np.concatenate(genes)
    ortho = np.unique(genes)
    return np.vstack([types, genes]).T


def alignTrees(trees) :
    result = [[], [], []]
    for idx, ts in enumerate(zip(*trees)) :
        res = result[idx]
        indices = [ 0 for t in ts ]
        while indices[0] < len(ts[0]) :
            s = np.min([ t[i][0] for t, i in zip(ts, indices) ])
            res.append([s] + [ t[i][1] for t, i in zip(ts, indices) ])
            for t, i in zip(ts, indices) :
                t[i][0] -= s
            for ti, t in enumerate(ts) :
                i = indices[ti]
                if t[i][0] <= 0 :
                    indices[ti] += 1
    return result
def duplicateOrthoTree(trees1, trees2, distDuplication=0.005) :
    result = alignTrees([trees1, trees2])
    tips = np.random.permutation(result[1][0][1].get_leaf_names())
    dists = np.random.exponential(distDuplication, size=len(tips))
    events = [[t.split('_', 1)[0], t, d] for t, d in zip(tips.tolist(), dists)]
    for res in result :
        for r in res :
            w, t1, t2 = r
            tr, ts = t1.copy(), t2.copy()
            n1 = {}
            for n in tr.get_leaves() :
                o = int(n.name.split('_', 1)[0])
                if o not in n1 :
                    n1[o] = [n]
                else :
                    n1[o].append(n)
            n2 = { n.split('_', 1)[0]:n for n in ts.get_leaf_names() }
            for o, cn1, dist in events :
                c2 = ts.get_leaves_by_name(n2[o])
                if not c2 :
                    continue
                c2 = c2[0]
                c1 = tr.get_leaves_by_name(cn1)[0]
                d1 = dist
                while d1 - c1.dist > 0 and c1.up :
                    d1, c1 = d1 - c1.dist, c1.up
                d2 = dist
                while d2 - c2.dist > 0 and c2.up :
                    d2, c2 = d2 - c2.dist, c2.up
                p, l = c1.up, c1.dist
                
                n = ete3.TreeNode()
                if p is not None :
                    p.remove_child(c1)
                n.add_child(c1)
                c1.up, c1.dist = n, d1
                
                if p is not None :
                    p.add_child(n)
                    n.dist = l - c1.dist
                else :
                    tr = n
                    for _, x, _ in events :
                        tr.get_leaves_by_name(x)[0]
                p2 = c2.up
                if p2 is not None :
                    p2.remove_child(c2)
                n.add_child(c2)
                c2.up, c2.dist = n, d2
                if p2 is not None :
                    gp = p2.up
                    if gp :
                        gp.remove_child(p2)
                        for c in p2.get_children() :
                            p2.remove_child(c)
                            gp.add_child(c)
                            c.up, c.dist = gp, c.dist + p2.dist
                    elif len(p2.get_children()) == 1 :
                        ts = p2.get_children()[0]
                        ts.up = None
                else :
                    break
            r[:] = [w, tr]
            
    return result
                

def getOrthoTree(trees, distOrtholog, postfix, strand, s, e, t, v, i0=0) :
    res = [[], [], []]
    while v > trees[-1][0] :
        x = min(s, trees[-1][0])
        s, e, t, v = s-x, e-x, t-x, v-x
    for r, i, j in ((res[0], s, e), (res[1], e, t), (res[2], t, v)) :
        while trees[i0][0] <= i :
            i0 += 1
        for m in range(i0, len(trees)) :
            tree = trees[m]
            i1, j1 = max(tree[0]-tree[1], i), min(tree[0], j)
            if j1 - i1 > 0 :
                tre = ete3.Tree(tree[2])
                for n in tre.get_descendants() :
                    n.dist *= distOrtholog
                for n in tre.get_leaves() :
                    n.name = '{1}_{0}'.format(postfix, n.name)            
                r.append([int(j1-i1), tre])
            else :
                break
    if strand < 0 :
        res = [ r[::-1] for r in res[::-1] ]
    return res, i0

def mergeOrthoTrees(prefix, orthoTrees, distParalog) :
    genes = list(orthoTrees.values())
    trees = alignTrees(genes)

    cmd = '{simbac} -N {0} -T 0 -R 0 -c {1}.gene'.format(\
        len(genes), prefix, **externals )
    subprocess.Popen(cmd.split(), stdout=subprocess.PIPE, universal_newlines=True).communicate()
    paratree = ete3.Tree('{0}.gene'.format(prefix))
    os.unlink('{0}.gene'.format(prefix))
    for n in paratree.get_descendants() :
        n.dist *= distParalog

    results = [[], [], []]
    for res, ts in zip(results, trees) :
        for tt in ts :
            tree = paratree.copy()
            for n in tree.get_leaves() :
                t = tt[int(n.name)+1]
                p = n.up
                p.remove_child(n)
                p.add_child(t)
                t.up, t.dist = p, n.dist
            res.append([tt[0], tree])
    return results

i00 = 0
def getHomologTree(prefix, homologs, trees, distOrtholog, distParalog, distDuplication) :
    orthoTrees = {}
    global i00
    i0s = []
    for gene in homologs :
        if gene[3] not in orthoTrees :
            orthoTrees[gene[3]], i0 = getOrthoTree(trees, distOrtholog, gene[0], gene[1], gene[7], gene[8], gene[9], gene[10], i00)
        else :
            tree, i0 = getOrthoTree(trees, distOrtholog, gene[0], gene[1], gene[7], gene[8], gene[9], gene[10], i00)
            orthoTrees[gene[3]] = duplicateOrthoTree(orthoTrees[gene[3]], tree, distDuplication)
        i0s.append(i0)
    i00 = np.min(i0s)
    if len(orthoTrees) > 1 :
        homoTree = mergeOrthoTrees(prefix, orthoTrees, distParalog)
    else :
        homoTree = list(orthoTrees.values())[0]
    return homoTree

controls = dict(cds='''[TYPE] CODON 1
[MODEL] m1
[submodel] 2.5 0.3
[indelmodel] NB {indelLen} 1
[geneticcode] 11
[indelrate]   {indelRate}

{trees}

[PARTITIONS] seq
{partitions}

[EVOLVE] seq 1 seq
''', 
               igr='''[TYPE] NUCLEOTIDE 1
[MODEL] m1
[submodel] HKY 2.5
[statefreq] 0.25 0.25 0.25 0.25
[indelmodel] NB {indelLen} 1
[indelrate]   {indelRate}

{trees}

[PARTITIONS] seq
{partitions}

[EVOLVE] seq 1 seq
''')

def getHomoSequence_indelible(prefix, homologs, homologTree, indelRate, indelLen, freqStart, freqStop) :
    tips = [int(t.split('_')[0])-1 for t in homologTree[1][0][1].get_leaf_names()]
    results = { homolog:{t:[] for t in tips} for homolog in homologs.T[0] }
    with tempfile.TemporaryDirectory(prefix=prefix, dir='.') as tmpdirname:
        for region, treeBlock in zip(['igr', 'cds', 'igr'], homologTree) :
            trees = [ ['t{0}'.format(id), tre[1].write(format=1, dist_formatter='%0.12f')] for id, tre in enumerate(treeBlock) ]
            partitions = [ ['t{0}'.format(id), tre[0] if region == 'igr' else int(tre[0]/3+0.5)] for id, tre in enumerate(treeBlock) ]
            for pId, p in reversed(list(enumerate(partitions))) :
                if p[1] == 0 :
                    del trees[pId]
                    del partitions[pId]
            if len(trees) :
                control = controls[region].format(indelLen=(1. - 1./indelLen), indelRate=indelRate, \
                                                      trees='\n'.join(['[TREE] {0} {1}'.format(*tree) for tree in trees]), \
                                                      partitions='\n'.join(['  [{0} m1 {1}]'.format(*partition) for partition in partitions]) )
                with open(os.path.join(tmpdirname, 'control.txt'), 'w') as fout :
                    fout.write(control)
                subprocess.Popen([externals['indelible']], stdout=subprocess.PIPE, cwd=tmpdirname).communicate()
                seq = {}
                with open(os.path.join(tmpdirname, 'seq_TRUE.phy')) as fin :
                    fin.readline()
                    for line in fin :
                        part = line.strip().split()
                        if len(part) > 1 :
                            n, s = part
                            go, ge = np.array(n.split('_'), dtype=int)
                            if go-1 not in results[ge] :
                                results[ge][go-1] = [s]
                            else :
                                results[ge][go-1].append(s)
            else :
                for ge in results :
                    for go in results[ge] :
                        results[ge][go].append('')

    starts = np.unique([v[1].replace('-', '')[:3] for gene_seq in results.values() for v in gene_seq.values()], return_counts=True)
    starts = starts[0][np.argsort(-starts[1])]
    n = int(np.ceil(float(len(starts))/len(freqStart[0])))
    if n <= 1 :
        startConvs = np.random.choice(freqStart[0], p=freqStart[1], size = starts.shape[0], replace=False)
    else :
        fs0 = freqStart[0] * n
        fs1 = np.array(freqStart[1].tolist() * n)/n
        startConvs = np.random.choice(fs0, p=fs1, size = starts.shape[0], replace=False)
    startConvs = dict(zip(starts, startConvs))
    
    stops = np.unique([v[1].replace('-', '')[-3:] for gene_seq in results.values() for v in gene_seq.values()], return_counts=True)
    stops = stops[0][np.argsort(-stops[1])]
    n = int(np.ceil(float(len(stops))/len(freqStop[0])))
    if n <= 1 :
        stopConvs = np.random.choice(freqStop[0], p=freqStop[1], size = stops.shape[0], replace=False)
    else :
        fs0 = freqStop[0] * n
        fs1 = np.array(freqStop[1].tolist() * n)/n
        stopConvs = np.random.choice(fs0, p=fs1, size = stops.shape[0], replace=False)
    stopConvs = dict(zip(stops, stopConvs))
    
    neg_strand = set(homologs[homologs.T[1] == -1, 0])    
    for gene, genomes in results.items() :
        for genome, seq in genomes.items() :
            seq[1] = re.sub(r'^(-*)([^-])(-*)([^-])(-*)([^-])', lambda g:startConvs[g.group(2)+g.group(4)+g.group(6)] + g.group(1)+g.group(3)+g.group(5), seq[1])
            seq[1] = re.sub(r'([^-])(-*)([^-])(-*)([^-])(-*)$', lambda g:stopConvs[g.group(1)+g.group(3)+g.group(5)] + g.group(2)+g.group(4)+g.group(6), seq[1])

            if gene in neg_strand :
                seq = [ rc(s, '-') for s in seq[::-1] ]
            m, n, k, m0, n0, k0 = len(seq[0]), len(seq[1]), len(seq[2]), len(seq[0].replace('-', '')), len(seq[1].replace('-', '')), len(seq[2].replace('-', ''))
            genomes[genome] = [''.join(seq), m+1, m+n, m+n+k, m0+1, m0+n0, m0+n0+k0, -1 if gene in neg_strand else 1]
    return results


def getSequences(prefix, genomes, items, operonBlock, trees, idenOrtholog, idenParalog, idenDuplication, indelRate, indelLen, freqStart, freqStop) :
    distOrtholog = -3./4.*np.log(1.-4./3.*(1.-idenOrtholog))/2.
    distParalog = -3./4.*np.log(1.-4./3.*(1.-idenParalog))/2. - distOrtholog
    distDuplication = -3./4.*np.log(1.-4./3.*(1.-idenDuplication))/2.
    genes = np.max(genomes, 1)
    blocks = np.cumsum(borderedGeometric(operonBlock[0], operonBlock[1], operonBlock[2], genes.size))
    blocks = blocks[blocks < genes.size]
    strand = np.random.choice([-1, 1], 1)
    strands = np.zeros(genes.size, dtype=int)
    for s, e in zip([0] + blocks.tolist(), blocks.tolist()+[genes.size]) :
        strands[s:e] = strand
        strand *= -1
    genes = np.hstack([np.arange(genomes.shape[0])[:, np.newaxis], strands[:, np.newaxis], items[genes]])
    genes[genes.T[1] == -1, 4:] = genes[genes.T[1] == -1, 4:][:, ::-1]
    sites = np.cumsum(genes[:, 4:]).reshape(-1, 3)
    genes = np.hstack([genes, np.concatenate([[0], sites[:-1, -1]])[:, np.newaxis], sites])
    
    alignments, genome_seqs, annotations, archives = [ [] for g in np.arange(genomes.shape[1]) ], [ [] for g in np.arange(genomes.shape[1]) ], [ [] for g in np.arange(genomes.shape[1]) ], {}
    for gId, gene in enumerate(genes) :
        print(gId, gene)
        if gId not in archives :
            homologs = genes[(genes.T[3]/10).astype(int) == (gene[3]/10).astype(int)]
            homologTree = getHomologTree(prefix, np.random.permutation(homologs), trees, distOrtholog, distParalog, distDuplication)
            homologSeq = getHomoSequence_indelible('{0}.gene.{1}_'.format(prefix, gId), homologs, homologTree, indelRate, indelLen, freqStart, freqStop)
            archives.update(homologSeq)

        gene_seqs = archives.pop(gId)
        gene_stat = genomes[gId]
        for genomeId, gseq in gene_seqs.items() :
            if gene_stat[genomeId] >= 0 :
                alignments[genomeId].append(gseq[0])
                genome_seqs[genomeId].append(gseq[0].replace('-', ''))
                annotations[genomeId].append([gene[3]]+gseq[1:])
            else :
                alignments[genomeId].extend('-'*len(gseq[0]))
                annotations[genomeId].append([gene[3]] + gseq[1:4] + [ 0, 0, 0, gseq[7]])
    alignments = [ ''.join(aln) for aln in alignments ]
    genome_seqs = [ ''.join(genome) for genome in genome_seqs ]
    annotations = np.array(annotations)
    aSite = np.cumsum(annotations[:, :, 3], axis=1) - annotations[:, :, 3]
    annotations[:, :, 1:4] += aSite[:, :, np.newaxis]
    gSite = np.cumsum(annotations[:, :, 6], axis=1) - annotations[:, :, 6]
    annotations[:, :, 4:7] += gSite[:, :, np.newaxis]
    return alignments, genome_seqs, annotations

def writeOut(prefix, alignments, genome_seqs, annotations) :
    with open(prefix+'.aligned.fasta', 'w') as fout :
        for id, aln in enumerate(alignments) :
            fout.write('>{0}\n{1}\n'.format(id, aln))
    with open(prefix+'.aligned.tbl', 'w') as fout :
        for annotation in annotations[0] :
            fout.write('{0}\t{1}\t{2}\t{3}\t{4}\n'.format(annotation[1], annotation[2], '+' if annotation[7]>0 else '-', int(annotation[0]/10), annotation[0]%10))
    for id, (genome, ann) in enumerate(zip(genome_seqs, annotations)) :
        with open(prefix + '_{0}.gff'.format(id), 'w') as fout :
            fout.write('##gff-version 3\n##sequence-region {0} 1 {1}\n'.format(id, len(genome)))
            for gid, annotation in enumerate(ann) :
                if annotation[5] > annotation[4] :
                    fout.write('{0}\tGenomeSim\tgene\t{1}\t{2}\t.\t{3}\t.\tID={4}_gene;gene={5};locus_tag={4}\n'.format(\
                        id, annotation[4], annotation[5], '+' if annotation[7]>0 else '-', \
                        'G_{0}_{1}'.format(id, gid+1), \
                        '{0}_{1}'.format(int(annotation[0]/10), annotation[0]%10)) )
                    fout.write('{0}\tGenomeSim\tCDS\t{1}\t{2}\t.\t{3}\t0\tID={4};Parent={4}_gene;gene={5};locus_tag={4};transl_table=11;product=simulated_gene\n'.format(\
                        id, annotation[4], annotation[5], '+' if annotation[7]>0 else '-', \
                        'G_{0}_{1}'.format(id, gid+1), \
                        '{0}_{1}'.format(int(annotation[0]/10), annotation[0]%10)) )
            fout.write('##FASTA\n')
            fout.write('>{0}\n{1}\n'.format(id, '\n'.join([genome[i:i+80] for i in range(0, len(genome), 80)]) ))
    for id, genome in enumerate(genome_seqs) :
        with open(prefix+'_{0}.fna'.format(id), 'w') as fout :
            fout.write('>{0} [organism=Simulated Genome] [gcode=11]\n{1}\n'.format(id, '\n'.join([genome[i:i+80] for i in range(0, len(genome), 80)]) ))
    for id, ann in enumerate(annotations) :
        with open(prefix+'_{0}.tbl'.format(id), 'w') as fout :
            fout.write('>Feature {0} Table1\n'.format(id))
            for gid, annotation in enumerate( ann ) :
                if annotation[5] > annotation[4] :
                    if annotation[7] > 0 :
                        fout.write('{0}\t{1}\tgene\n\t\t\tgene\t{2}_{3}\n\t\t\tlocus_tag\t{4}\n{0}\t{1}\tCDS\n\t\t\tproduct\tsimulated_gene\n\t\t\ttransl_table\t11\n'.format(annotation[4], annotation[5], \
                                                                                                                       int(annotation[0]/10), annotation[0]%10, \
                                                                                                                       'G_{0}_{1}'.format(id, gid+1)))
                    else :
                        fout.write('{1}\t{0}\tgene\n\t\t\tgene\t{2}_{3}\n\t\t\tlocus_tag\t{4}\n{1}\t{0}\tCDS\n\t\t\tproduct\tsimulated_gene\n\t\t\ttransl_table\t11\n'.format(annotation[4], annotation[5], \
                                                                                                                       int(annotation[0]/10), annotation[0]%10, \
                                                                                                                       'G_{0}_{1}'.format(id, gid+1)))
                        
    
def runGenomeSim(args) :
    # prepare gene lengths and intergenic region
    items = itemGenerator(args.nBackbone, args.nMobile, 1-args.pBackbone, 1-args.pMobile )
    items = itemShaper(items, args.geneLen, args.igrLen)
    
    # simulate a global tree
    globaltrees, localtrees = treeGenerator(args.prefix, items, args.genomeNum, args.rec, args.recLen)
    seqtrees = localtrees if args.insRec == 1 else globaltrees
    itrees   = localtrees[:10000] if args.insRec == 1 else globaltrees
    dtrees   = itrees     if args.delRec == 1 else globaltrees
    # impose gene content variation on the tree
    genomes = setGeneContent(items, itrees, dtrees, args.tipAccelerate, args.genomeNum, args.aveSize, args.nCore, args.backboneBlock, args.mobileBlock)
    
    # write parameter and gene content
    with open(args.prefix+'.params.log', 'w') as fout :
        fout.write(str(args).replace(', ', ',\n'))
    with open('{0}.gene.content.tsv'.format(args.prefix), 'w') as fout :
        fout.write('#ID\t{0}\n'.format( '\t'.join(np.arange(genomes.shape[1]).astype(str)) ))
        for gId, gene in enumerate(genomes) :
            fout.write('{0}\t{1}\n'.format(gId+1, '\t'.join([ str(items[g, 1])[:-1]+'_'+str(items[g, 1]%10) if g >= 0 else '-' for g in gene ])))
    
    if not args.noSeq :
        # simulate sequences
        alignments, genome_seqs, annotations = getSequences(args.prefix, genomes, items, args.operonBlock, seqtrees, args.idenOrtholog, args.idenParalog, args.idenDuplication, args.indelRate, args.indelLen, args.freqStart, args.freqStop)
        # output
        writeOut(args.prefix, alignments, genome_seqs, annotations)

def genomeSim(a) :
    import argparse
    parser = argparse.ArgumentParser(description='''SimPan is a simulator for bacterial pan-genome. 
Global phylogeny and tree distortions are derived from SimBac and the gene and intergenic sequences are simulated using INDELile.''', formatter_class=argparse.RawTextHelpFormatter)

    parser.add_argument('-p', '--prefix', help='prefix for all intermediate files and outputs. {DEFAULT: SimPan]', default='SimPan')
    parser.add_argument('--genomeNum', help='No of genome in population [DEFAULT: 20]', default=20, type=int)

    parser.add_argument('--geneLen', help='[negative bionomial with r=2] mean,min,max sizes of genes [DEFAULT: 900,150,6000]', default='900,150,6000')
    parser.add_argument('--igrLen', help='[negative bionomial] mean,min,max sizes of intergenic regions [DEFAULT: 50,0,300]', default='50,0,300')
    parser.add_argument('--backboneBlock', help='[geometric] mean,min,max number of backbone genes per block [DEFAULT: 3,0,30]', default='3,0,30')
    parser.add_argument('--mobileBlock', help='[geometric] mean,min,max number of mobile genes per block [DEFAULT: 10,0,100]', default='10,0,100')
    parser.add_argument('--operonBlock', help='[geometric] mean,min,max number of continuous genes that share the same coding strand [DEFAULT: 3,0,15]', default='3,0,15')

    parser.add_argument('--aveSize', help='average gene number per genome (greater than nBackbone). [DEFAULT: 4500]', default=4500, type=int)
    parser.add_argument('--nBackbone', help='number of backbone genes (present in common ancestor) per genome. [DEFAULT: 4000]', default=4000, type=int)
    parser.add_argument('--nCore', help='sizea of core gene (smaller than the size of backbone genes). [DEFAULT: 3500]', default=3500, type=int)
    parser.add_argument('--nMobile', help='size of mobile gene pool for accessory genome. [DEFAULT: 20000]', default=20000, type=int)

    parser.add_argument('--pBackbone', help='propotion of paralogs in backbone (core) genes. [DEFAULT: 0.05]', default=0.05, type=float)
    parser.add_argument('--pMobile', help='propotion of paralogs in mobile (accessory) genes. [DEFAULT: 0.4]', default=0.4, type=float)

    parser.add_argument('--tipAccelerate', help='grandient increasing of gene indels in recent times. [DEFAULT: 100]', default=100., type=float)
    parser.add_argument('--rec', help='expected coverage of homoplastic events in pairwise comparisons. [DEFAULT: 0.05]', default=0.05, type=float)
    parser.add_argument('--recLen', help='expected size of homoplastic events. [DEFAULT: 1000]', default=1000, type=float)
    parser.add_argument('--seqRec', help='Use homoplastic events to infer sequences. Use 0 to disable [DEFAULT: 1]', default=1, type=int)
    parser.add_argument('--insRec', help='Use homoplastic events to infer gene insertions. Use 0 to disable [DEFAULT: 1]', default=1, type=int)
    parser.add_argument('--delRec', help='Use homoplastic events to infer gene deletions. Use 0 to disable [DEFAULT: 1]', default=1, type=int)
    
    parser.add_argument('--noSeq', help='Do not infer sequence but only the gene presence/absence. [DEFAULT: False]', default=False, action='store_true')
    
    parser.add_argument('--idenOrtholog', help='average nucleotide identities for orthologous genes. [DEFAULT: 0.98]', default=0.98, type=float)
    parser.add_argument('--idenParalog', help='average nucleotide identities for paralogous genes. [DEFAULT: 0.6]', default=0.6, type=float)
    parser.add_argument('--idenDuplication', help='average nucleotide identities for recent gene duplications. [DEFAULT: 0.995]', default=0.995, type=float)
    
    parser.add_argument('--indelRate', help='average frequency of indel events relative to mutation rates. [DEFAULT: 0.01]', default=0.01, type=float)
    parser.add_argument('--indelLen', help='average size of short indel events within each gene. [DEFAULT: 10]', default=10, type=float)
    parser.add_argument('--freqStart', help='frequencies of start codons of ATG,GTG,TTG. DEFAULT: 0.83,0.14,0.03', default='0.83,0.14,0.03')
    parser.add_argument('--freqStop', help='frequencies of stop codons of TAA,TAG,TGA. DEFAULT: 0.63,0.08,0.29', default='0.63,0.08,0.29')

    args = parser.parse_args(a)
    args.geneLen = np.round(np.array(args.geneLen.split(',')).astype(float)/3)
    args.igrLen = np.array(args.igrLen.split(',')).astype(float)
    args.backboneBlock = np.array(args.backboneBlock.split(',')).astype(float)
    args.mobileBlock = np.array(args.mobileBlock.split(',')).astype(float)
    args.operonBlock = np.array(args.operonBlock.split(',')).astype(float)
    args.freqStart = ('ATG,GTG,TTG'.split(','), np.array(args.freqStart.split(',')).astype(float))
    args.freqStop = ('TAA,TAG,TGA'.split(','), np.array(args.freqStop.split(',')).astype(float))
    runGenomeSim(args)
    
if __name__ == '__main__' :
    genomeSim(sys.argv[1:])
```
